# Assessment of the evolutionary consequence of putative driver mutations in colorectal cancer with spatial multiomic data

**DOI:** 10.1101/2021.07.14.451265

**Authors:** Timon Heide, Jacob Househam, George D Cresswell, Inmaculada Spiteri, Claire Lynn, Chris Kimberley, Max Mossner, Luis Zapata, Calum Gabbutt, Daniele Ramazzotti, Bingjie Chen, Javier Fernandez-Mateos, Chela James, Alessandro Vinceti, Alison Berner, Melissa Schmidt, Eszter Lakatos, Ann-Marie Baker, Daniel Nichol, Helena Costa, Miriam Mitchinson, Benjamin Werner, Francesco Iorio, Marnix Jansen, Christopher Barnes, Giulio Caravagna, Darryl Shibata, John Bridgewater, Manuel Rodriguez-Justo, Luca Magnani, Trevor A Graham, Andrea Sottoriva

## Abstract

Cancer genomic medicine relies on targeting driver genes. However, current catalogues of cancer drivers are mostly based on indirect measurements of mutation frequencies, positions or types, rather than their effect on clonal expansions *in vivo*. Moreover, non-genetic drivers are largely unknown, as are the epigenetic and transcriptomic effects of genetic drivers. Here we perform spatial computational inference on multiomic data with matched whole-genome sequencing, ATAC-seq and RNA-seq. Using 436 samples, we directly quantify the contribution, or lack thereof, of putative driver genes to subclonal expansions *in vivo* in 30 colorectal carcinomas (4-33 samples per patient, median=15). Although subclonal neutral evolution was widespread (13/26 cases with sufficient data), there were cases with clear evidence of subclonal selection (6/26) in which we measured epigenetic and transcriptomic differences between subclones *in vivo*. In 7/26 cases we could not distinguish between neutral or selective evolution with the available data. We identified expanding subclones that were not driven by known genetic alterations, and propose candidate epigenetic drivers. We identified the distinguishing patterns of genomic heterogeneity produced in fast, exponentially growing tumours (7/26) versus neoplasms growing only at the periphery (19/26), as well as identifying clonally intermixed (16/28 cases with sufficient data) versus segregated malignancies (10/28). Our model-based approach measures genetic and non-genetic subclonal selection, or lack thereof, in space and time and allows *in vivo* comparisons of the emergent phenotypic properties of subclones within human tumours.

## Introduction

Identification of somatic DNA mutations in cancer driver genes is at the core of genomic medicine and precision oncology (Velden et al., 2019). Seminal contributions have led to the generation of compendia of curated cancer drivers (Martínez-Jiménez et al., 2020; Sondka et al., 2018) that inform clinical trials and drug development. Although these efforts are a fundamental part of cancer research, most cancer driver genes are not identified by measuring their functional impact *in vivo* in humans. Instead, sophisticated statistical analyses discover driver gene candidates based on the frequency of genetic mutations within the gene (i.e., if a gene is mutated more often than expected by chance in a cohort), the clustering of mutations along the gene length (i.e., if mutations in the gene accumulate in specific hotspots) and predicted damaging/functional effects on the specific protein that the gene encodes (i.e., if mutations are predicted to substantially change the protein function). So far, relatively few cancer driver genes have been validated functionally in experimental systems (Bailey et al., 2018)], and moreover methods that evaluate what cancer driver gene mutations actually do *in vivo* during human tumourigenesis are lacking. For example, only very recently has the mechanism of selective advantage provided by *APC* mutation – the gatekeeper mutation for colorectal carcinogenesis discovered more than three decades ago (Kinzler et al., 1991) – been elucidated (Flanagan et al., 2021; Neerven et al., 2021). Moreover, potential epigenetic drivers of subclonal expansions have received comparably little attention, and there are calls for the field to move beyond the genome in the study of intra-tumour heterogeneity (Black and McGranahan, 2021).

In many cancer evolution studies, it is typical to specify a list of driver genes *a priori* is applied and annotate alterations to these genes upon phylogenetic trees that give the impression of functional evolution (Gundem et al., 2015; Noorani et al., 2020; Yates et al., 2015, 2017). However, this does not correspond to a direct measurement of the effects of the driver on the cellular population. Convergence of multiple driver mutations in the same gene in a patient are a strong sign of selection (Gerlinger et al., 2012, 2014), but such events are not very common and often arise in the context of therapy, where extremely strong selective pressures are present (Parikh et al., 2019). Moreover, due to the high mutation rate of copy number alterations (CNAs) (Burrell et al., 2013; Dijk et al., 2021), convergence of chromosomal arm events should not be considered a likely sign of selection since the same CNAs could occur multiple times independently (Williams et al., 2021). Therefore, it is likely that a proportion of reported drivers may be false positives, due to incorrect normalisation for confounding factors in genetic analyses, or by assumption that driver status of a particular gene is maintained across tissues. More importantly, some genes may behave as drivers in one patient, but not in another.

We previously demonstrated that colorectal cancers, after malignant transformation, often grow as a single ‘Big Bang’ expansion of clones under weak or without subclonal selection (Sottoriva et al., 2015), an observation that was corroborated in subsequent studies (Cross et al., 2018; Sun et al., 2017). This model also predicts clonal intermixing, where samples in distant regions of the tumour are more closely related to nearby samples due to the effect of early disordered growth. Big Bang dynamics have also been identified in other cancer types (Gaiti et al., 2019; Gao et al., 2016; Ling et al., 2015; Sun et al., 2018). Intra-tumour genetic heterogeneity in ‘Big Bang’ cancers is largely driven by neutral evolution (Kimura, 1984), which can be verified through detection of a characteristic pattern of mutation allele frequencies in bulk sequencing of single samples (Williams et al., 2016, 2018) as well as the pattern of intra-tumour heterogeneity in multi-region and longitudinal sampling (Caravagna et al., 2020a). Notably the Big Bang model, unlike a strict model of neutrality, does allow for the presence of weakly selected subclones that do not grow rapidly enough to substantially change the clonal makeup of the tumour.

Here we use colorectal carcinogenesis as a model system to evaluate the selective advantage of putative driver mutations. We delineate the evolutionary dynamics of primary colorectal malignancies using a multi-region approach based on single-gland sequencing of multiple distant tumour regions, making use of a novel phylogenetic approach that, unlike single bulk sample strategies (Dentro et al., 2021), has good power to detect positively-selected subclonal expansions. Marrying this sensitive detection of subclonal selection together with an assessment of the genetic makeup of subclones, enables us to assess the functional impact of putative cancer driver gene mutations – measured in terms of the fitness advantage bestowed to the mutant clone -*in vivo* in patients. The same approach enables us to detect candidate driver epigenetic modifications. Further, we leverage matched ATAC-seq and RNA-seq data to functionally characterise the epigenetic and transcriptomic identity of positively-selected subclones.

## Results

### Darwinian selection on DNA mutations in cancer driver genes

We implemented a single-clone multiomic profiling protocol that allows concomitant assessment of single nucleotide variants (SNVs), copy number alterations (CNAs), chromatin accessibility changes and transcriptomic profiles in many samples from the same patient (data generation described in our associated paper EPIGENOME). Specifically, from 30 primary colorectal cancers and 8 concomitant adenomas we generated deep whole-genome sequencing (WGS, median depth 35x) in 3-15 samples per patient (median=9), low-pass whole genome sequencing (lpWGS, median depth 1.2x) in 0-22 samples per patient (median=8), and chromatin characterisation with ATAC-seq in 18-61 samples per patient (median=42). We also generated 297 whole-transcriptomes (1-40 samples per patient, median=7) with many overlapping the WGS dataset, the ATAC-seq dataset or both.

We leveraged the large number of samples per patient profiled with whole-genome sequencing to accurately identify clonal and subclonal somatic variants in each case. The IntOGen list (Martínez-Jiménez et al., 2020) of 72 colorectal cancer driver genes is the gold standard compendium of putative drivers in intestinal malignancies. We discarded from the list three genes that are likely false positives, LRP1B, KMT2C and PARP4, due to mappability issues in their highly repetitive regions, passenger hotspots, and dN/dS∼1 (see below) in the TCGA colorectal cancer cohort. In our cohort, we annotated clonal and subclonal coding mutations and indels in the remaining 69 genes (Figure 1A), illustrating an apparent plethora of cancer driver gene mutations in our cohort.

**Figure 1.**
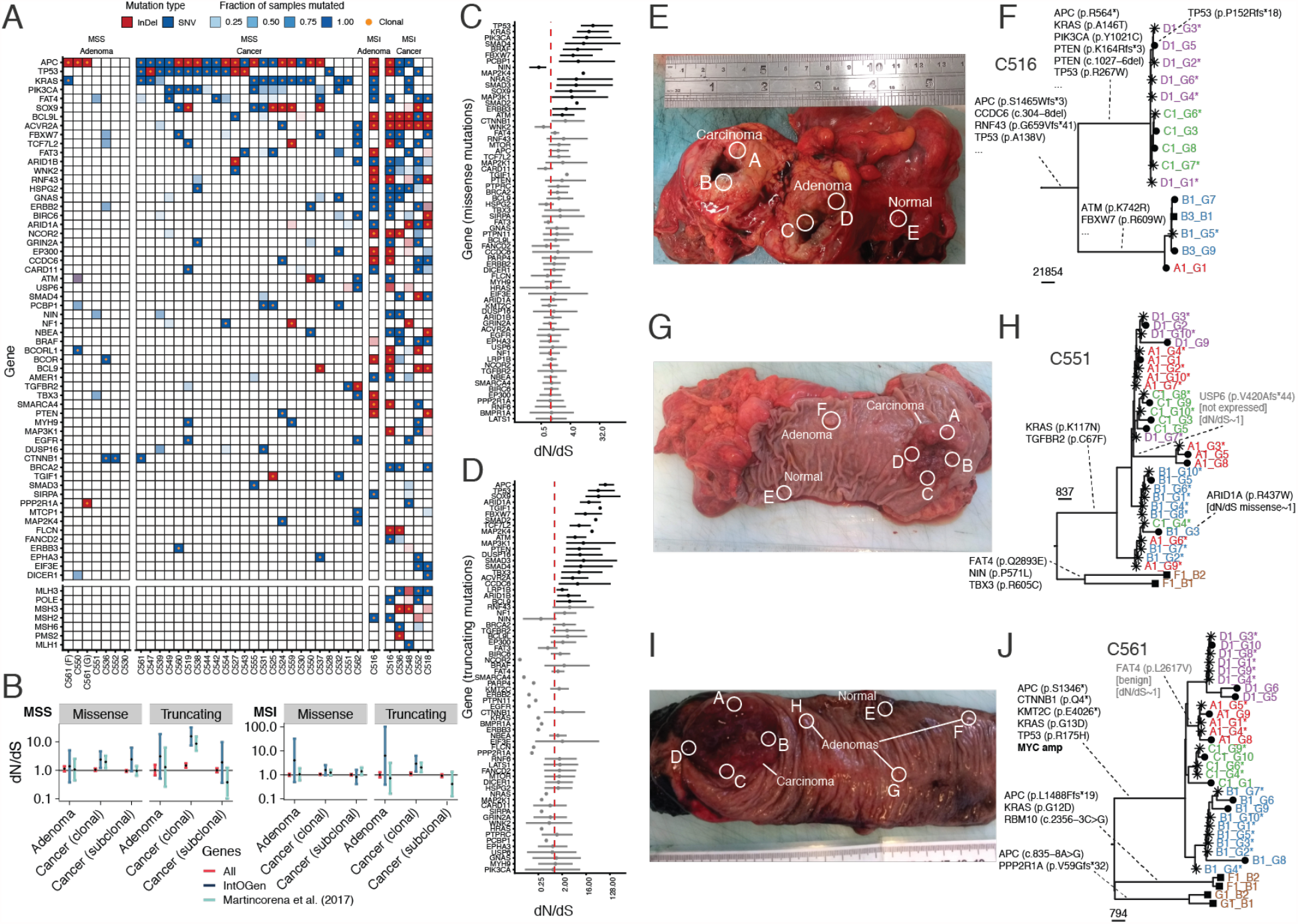
Clonal and subclonal colorectal cancer drivers from spatial genomic profiling. **(A)** Nonsynonymous somatic mutations and indels in IntOGen colorectal cancer driver genes, with clonality status indicated. **(B)** dN/dS analysis on clonal versus subclonal driver gene mutations, divided between MSS (Microsatellite Stability) and MSI (Microsatellite Instability) cases. **(C)** Per-gene missense mutations dN/dS and **(D)** per-gene truncating mutations dN/dS for the 69 IntOGen drivers in TCGA colon and rectal cohorts, many genes have dN/dS value ∼1 indicating lack of evidence for positive selection. **(E)** In this MSI case C516, the cancer (region A and B) and the adenoma (regions C and D) formed a large mass and were physically adjacent to one another. **(F)** The precursor adenoma shared a mismatch repair mutation in MSH2 with the cancer but displayed early divergence. **(G)** Case C551 presented with a cancer and a concomitant adenoma that were very distant, pointing at two independent events. **(H)** The phylogenetic tree was characterised by clonal intermixing of diverging lineages collocated in the same region (e.g., some lineages from regions A, B and C were genetically close). Subclonal drivers of unknown significance were present, including a non-expressed variant in USP6 and an ARID1A mutation. Early divergence between the cancer and the adenoma F was evident, with no shared drivers between the two lesions. **(I)** Case C561 presented with a large cancer mass and multiple small concomitant adenomas. **(J)** Again, there was no significant somatic alteration in common between the different lesions. The cancer showed clonal amplification of MYC and only a benign subclonal mutation in FAT4.

We found that the most frequently mutated drivers in colorectal cancer, such as APC, KRAS, TP53, and SOX9, as well as other known drivers such as PTEN, EGFR, CCDC6, PCBP1, ATM and CTNNB1 are invariably clonal in cancers, with the exception of one single case with a subclonal KRAS mutation. These findings are consistent with previous multi-region sequencing studies (Cross et al., 2018)but contradict claims of frequent subclonality of these genes in single-sample bulk data (Dentro et al., 2021), highlighting the need for properly powered analyses to assess intra-tumour heterogeneity (Caravagna et al., 2020a).

We then assessed the strength of evolutionary selection with dN/dS analysis (Martincorena et al., 2017; Zapata et al., 2018) which quantifies the excess of non-synoymous mutations in driver genes. We compared dN/dS values measured in the whole coding genome (‘all’), in a list of pan-cancer driver genes (Martincorena et al., 2017) and in the IntOGen colorectal cancer genes from Figure 1A. We split the cohort between cases characterized by microsatellite stability (MSS) versus instability (MSI) to account for the very different mutation rates in these two groups (in line with previous analyses (Martincorena et al., 2017). We found clear evidence of selection for clonal missense and truncating mutations in MSS cancers, with dN/dS values significantly larger than one, in both lists of putative cancer drivers (Figure 1B). The dN/dS value is higher for the IngoGen list, confirming the list is highly enriched for genes that are true drivers in this colorectal cancer cohort. We found no evidence of subclonal selection for truncating variants. Moreover, missense mutations showed slightly higher than 1 dN/dS values although non-significant, suggesting that only a small subset of subclonal IntOGen driver mutations are likely to be under positive selection. For the MSI cases selection was less evident from dN/dS, likely due to the higher mutation rate generating a much larger number of neutral mutations in cancer driver genes, thus diluting the dN/dS signal, but nevertheless selection for clonal truncating mutations was significant in MSI cancers (Figure 1B).

We then examined the dN/dS values for each of the colorectal IntOGen drivers, including the three genes we discarded, to assess the evidence of positive selection in the TCGA colon and rectal cancer cohorts combined with additional data(Giannakis et al., 2016; Vasaikar et al., 2019) using SOPRANO (Zapata et al., 2018) for a total of n=1,253 colorectal cancer cases analysed. This large number of cases allows calculating dN/dS per each driver gene, rather than in sets of genes. The results are shown in Figure 1C for missense mutations and in Figure 1D for nonsense variants. Clearly, most genes in the list show no evidence of selection, with the majority of the top genes being the “usual suspect” colorectal cancer drivers (Cross et al., 2018) We confirmed that both missense and nonsense mutations in PARP4 and KMT2C had dN/dS∼1, as did missense mutations in LRP1B (we only find missense subclonal mutations in this gene in our cohort).

The values of dN/dS for subclonal mutations being ∼1 are consistent with extensive neutral evolution ongoing at the subclonal level in these tumours. Although dN/dS measurements are powerful tools to assess selection, both positive and negative (Zapata et al., 2018), they have the limitation that they provide an ‘average’ estimate of selection across a group of patients (Heide et al., 2018), and dN/dS on driver genes cannot be estimated from a single patient. Further, we note that theory predicts dN/dS to be less sensitive to detect subclonal selection than clonal selection (Williams et al., 2020). Subclonal expansions are likely to be few and rare, but nevertheless very important (Turajlic et al., 2019). Recognition that most intra-tumour heterogeneity in colorectal cancers is the consequence of neutral evolution provides a tractable “null hypothesis” against which deviations due to clonal selection can be robustly identified (Williams et al., 2016, 2018) Neutral evolution is a simple evolutionary scenario to simulate, and this simplicity allows for the design of robust statistical methods to detect the action of selection through deviations from the expectation provided by neutral evolution (Caravagna et al., 2020a, 2020b; Durrett, 2013; Kessler and Levine, 2013, 2014; Williams et al., 2016, 2018).

We next sought to identify which subclones were selected, and which ones were not, in our cohort, on a patient by patient basis.

### Measuring evolution with spatial profiling and phylogenetics

Phylogenetics relates the genetic variation amongst individuals to their shared genetic ancestry (Yang and Rannala, 2012). Phylogenetic methods have been extensively applied to cancer data (Schwartz and Schäffer, 2017). Phylodynamics extends this methodology to incorporate “demographic” mathematical models of the underlying stochastic birth and death process of lineage evolution (Stadler et al., 2021). Phylodynamics provides an explicit means to model mutation, trait evolution through Darwinian selection, and spatial processes. The methodology has been extensively applied in animals (Gaggiotti et al., 2002), viruses (Suchard et al., 2018) and cancer (Alves et al., 2019; Lote et al., 2017; Stadler et al., 2021). In the context of cancer, clonal (often also referred as ‘truncal’) mutations have undergone selection and clonal expansion in the past and are now ‘fixed’ in the population. These contain no phylodynamic information on how the population is currently evolving. Instead, intra-tumour heterogeneity data from multi-region profiling documents selection “in action” (Williams et al., 2019). Moreover, when a subclone is “caught in the act” of expanding but it has not yet reached fixation, it provides a unique opportunity to functionally characterise the genetic and epigenetic alterations driving its expansion. Whereas single-sample clonal driver mutations are a black and white ‘photo’ of what happened in the past, multi-region sequencing provides more of a ‘movie’ of ongoing tumour evolution (Swanton, 2012; Turajlic et al., 2019).

Here, we combine multi-region single gland whole-genome sequencing with the spatial information we retained from our sampling strategy (Figure S1) to reconstruct the phylogenetic trees for each patient initially using a maximum parsimony ‘model-free’ approach. Our single-clone tissue collection with monoclonal glands makes the task easier than using large bulks of tissue as subclonal deconvolution is not required (Alves et al., 2017).

We reconstructed the phylogenetic history of each tumour using SNVs from single glands and minibulks. Each sample was imaged (Figure S2) and whole-genome data was used to call copy number alterations (Figure S3), mutations and indels (Figure S4) (see related EPIGENOME manuscript for methods). Histograms of the variant allele frequency (VAF) for each sample is reported in Figure S5. We also developed a method to genotype sets of SNVs present in different branches of the tree on lpWGS samples and hence position those in the tree (see Material and Methods). This allowed us to construct large phylogenetic trees that informed on the population structure of each cancer and concomitant adenoma, and to link clades on the phylogenetic tree to the geographical location where the samples were taken (Figure 1E-J). This was done for the whole cohort (Figure 2, see Figure S6 for bootstrapping values confirming tree robustness).

**Figure 2.**
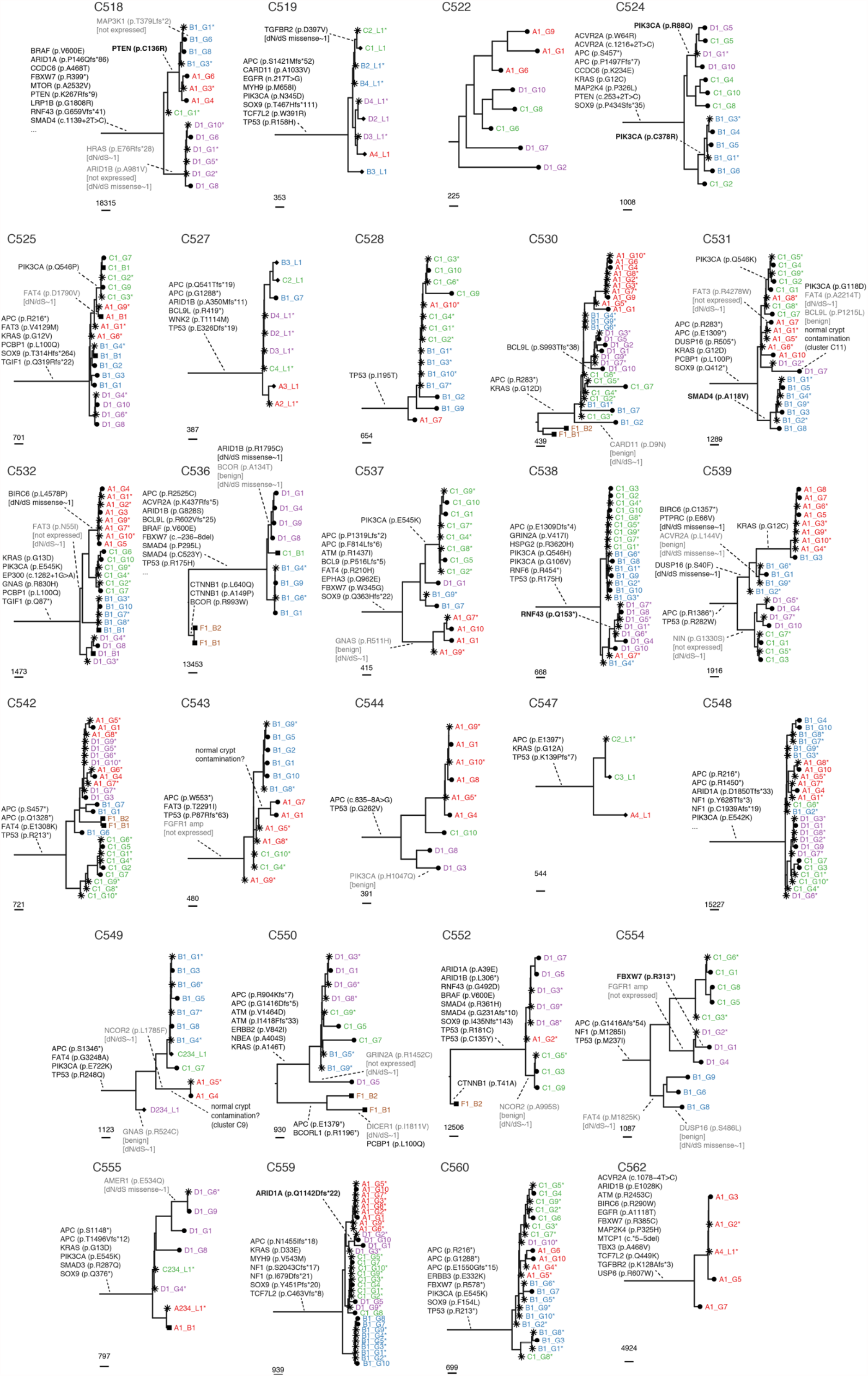
Phylogenetic reconstruction of all cancers and adenomas. Putative cancer driver genes from the IntOGen set were reported in each branch. For MSI cases we report only a subset of most relevant genes (note “…”). For subclonal drivers, we report whether the variant was expressed (bold), not expressed or benign (grey), and if the per-gene dN/dS value was ∼1.

We found that the large majority of the trees were balanced, with similar branch lengths between different samples and regions. A clear outlier was case C539 where the tree contains a particularly large clade that expanded recently and spans multiple geographical regions of the tumour (all A and part of B). These patterns are expected when subclonal selection is active (Chkhaidze et al., 2019). Indeed, the expanded clade contained a KRAS G12C mutation.

Distinctive pattern of clonal intermixing of genetically divergent samples that ended up in geographical proximity was evident in 16/28 (57%) of cases (Figure 1E-J, 2 and 3A). We previously reported the surprising finding that samples from distant tumour regions were more related to each other than nearby samples, and argued through simulations that this was predicted by ‘Big Bang’ dynamics and was a sign of early malignant potential (Sottoriva et al., 2015; Sun et al., 2017). Other cancers were spatially well organised, with all samples from one region being genetically similar (e.g., cases C525 and C532). These patterns of intermixing clearly split cancers into two groups, one characterised by subclonal intermixing, and one by subclonal segregation (Figure 3A).

**Figure 3.**
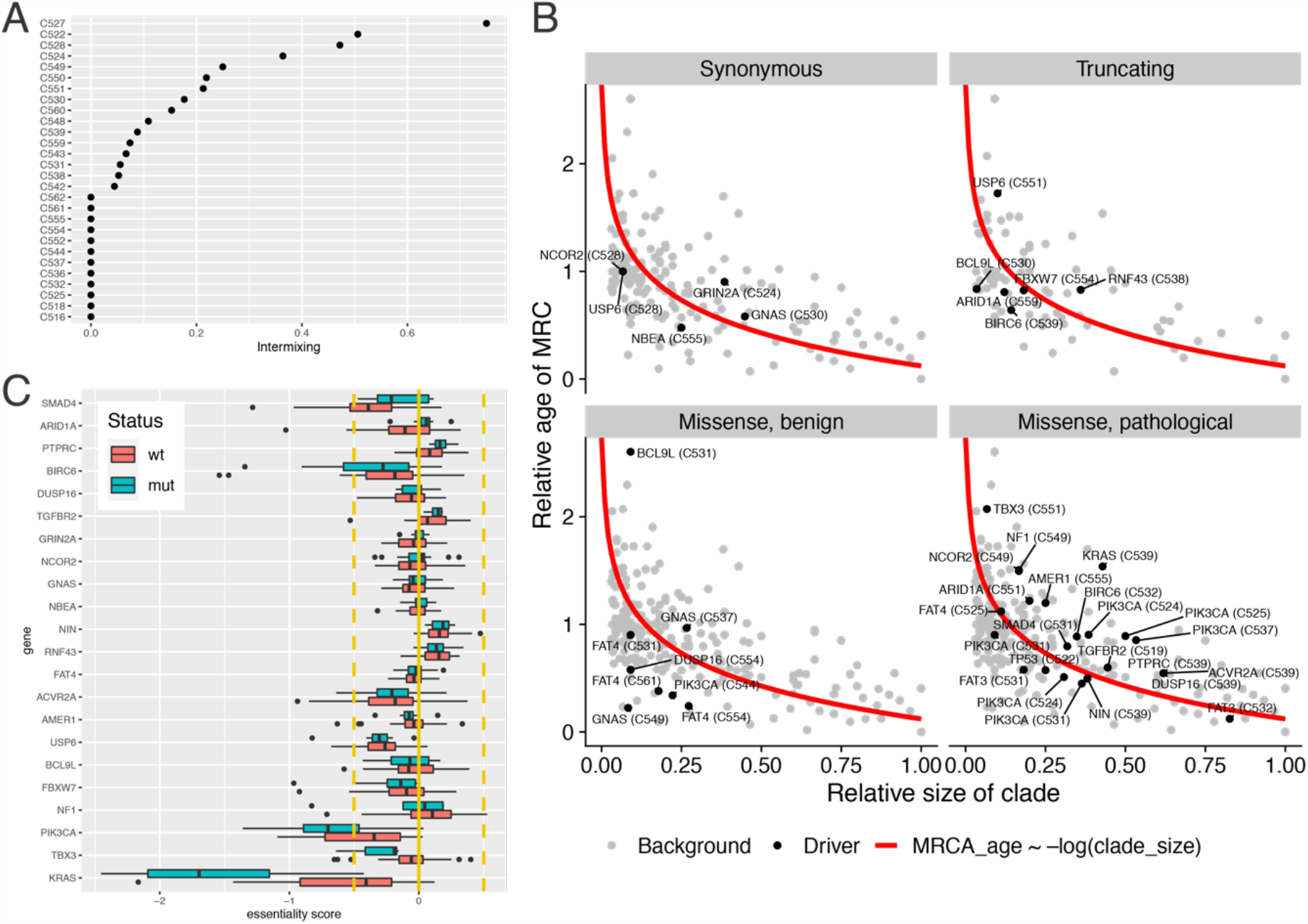
Clonal intermixing, subclone dynamics and essentiality. **(A)** Subclonal intermixing between genetically divergent lineages in the same geographical region was common, with 16/28 cases showing this pattern. **(B)** Relative clade size vs relative age of MRCA for each putative driver variant versus background (non-cancer genes) shows that probably only RNF43 in truncating mutations, and a few variants in KRAS and PIK3CA show clear signs of deviating from the neutral expectation in red. **(C)** Indeed, essentiality scores from the cancer dependency map (DepMap) of many putative driver genes we find subclonal are not significant across hundreds of cancer cell lines, and are not significantly different across mutant versus wild-type subpopulations of cell-lines (which would be consistent with oncogenic function), with the exception of KRAS and PIK3CA. Orange lines denote the significance thresholds of the essentiality score as defined by DepMap, i.e. half of the median essentiality scores observed for prior known essential genes (= −1).

We then annotated the 44 subclonal colorectal cancer driver variants we find in our cohort from IntOGen list from Figure 1A onto the phylogenetic trees (excluding adenomas). We noted that 7/44 subclonal drivers were from MSI cases, which showed dN/dS in subclonal mutations of ∼1 (Figure 1B), suggesting that most the majority of these 7 putative driver mutations were actually neutral.

We further examine the PolyPhen (Adzhubei et al., 2010) functional score of each mutation and found that 10/44 (22.7%) mutations were identified as putatively benign SNVs (marked in grey in Figure 1E-J and 2). We also leveraged on the whole-transcriptome information to verify the expression of mutated transcripts, and we found that for 7/44 (15.9%) driver mutations only wild-type reads were detected (also marked in grey). We could not assess mutant transcript expression for 29/44 mutations (65.9%) because of missing RNA data or lack of reads covering the variant location. Of those, 13/29 (45%) were in genes with dN/dS∼1 in the TCGA COAD and READ cohorts. We also report in bold the 6/44 (13.6%) variants identified as deleterious and also found expressed by RNA-seq.

Together, these analyses revealed that of the large number of putative driver events identified in our cohort (Figure 1A) most showed no evidence of being under selection: 38.6% of variants were either benign or not expressed in the cancer (and we note expression could not be assessed for two thirds of variants), and another 29.5% of variants were in genes with dN/dS∼1.

To further dissect which subclonal cancer driver genes are actually driving a subclonal expansion and which ones are effectively neutral, we perform further analysis of the topology of the reconstructed phylogenies. The trees in Figures 1E-J and 2 provided good estimates of time of emergence of different lineages (their Most Recent Common Ancestor −MRCA) measured relative to the total age of the tumour from trunk to leaves, calculated as the mean number of SNVs per lineage in the clade divided by mean SNVs per from root to leaf for every lineage. Moreover, for each lineage we had a good estimate of its clade size because of the many samples per clade. Under neutral evolution, only early diverging lineages lead to large clade sizes, and in general clade sizes are predicted to follow a power-law ∼1/f distribution generating more and more lineages at smaller and smaller frequency as the tumour expands. In comparison, positive selection causes late emerging lineages to undergo subclonal expansion, reaching clade sizes that are larger than expected under neutral evolution. The clearest example of this phenomenon in our cohort is case C539, where a late arriving subclonal lineage in region A underwent a large expansion (Figure 2). The relationship between MRCA time *t*_*MRCA*_(how far into the tumour’s past that a subclone was generated) and the size of the clade *f* (expressed as the proportion of samples within the mutated clade) should follow under neutrality as:

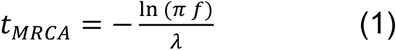

Where *π* is the copy number of the locus and *λ* is the tumour growth rate, assuming exponential growth N(t) = *e*^*λt*^. We calculated *t*_*MRCA*_ vs *f* for every gene mutation, dividing into five categories:

- non-cancer gene mutations (background)
- synonymous mutations driver genes
- missense mutations in driver genes scored as functionally *benign* by PolyPhen
- missense mutations in driver genes scored as functionally deleterious by PolyPhen
- truncating mutations in driver genes

When examining our data, we found that the large majority of mutations in driver genes behaves evolutionarily as the predicted from neutral theory (equation 1; Figure 3B). In particular, mutations in benign missense mutations as expected behave as neutral, as do truncating mutations which were indeed found to have dN/dS∼1. A few noticeable exceptions of functionally deleterious missense mutations (which indeed showed slightly higher dN/dS) show sign of deviating from the neutral expectation, most of all KRAS G12C in C539, but also a subset of the subclonal PIK3CA mutations. Evidently, putative drivers such as FAT3 and FAT4, which are very large genes that show often benign mutations behave largely neutral, are unlikely to be true drivers in colorectal cancer.

We attempted to estimate the functionality of the subclonal mutations in our cohort using an orthogonal data type. To this aim, we mined *gene essentiality scores* assembled within the cancer dependency map (DepMap) project and quantifying the functional consequences (in terms of cellular viability reduction) of CRISPR-Cas9 gene targeting, at a genome wide level, across hundreds of immortalised human cancer cell lines (Pacini 2021). Most CRC candidate drivers showed no evidence of essentiality (scaled essentiality score > −0.5, with −1 = median essentiality score for prior known essential genes) across the CRC cell lines of the DepMap dataset, whereas the two genes most likely under strong selection in our cohort, KRAS and PIK3CA, were significantly essential in many CRC cell lines and found significantly differentially essential when contrasting mutant versus wild-type CRC cell lines (Student’s t-test p-value < 10^−13^, Figure 3C).

These results provide, through statistical measurements on mutations and tree, an intuitive assessment of the evolutionary dynamics individual cases. Further, they illustrate of how “driverness” of candidate driver mutations can be determined by using phylogenetic analysis to assess the evolutionary consequence of driver mutation acquisition.

### Quantification of cancer evolutionary dynamics with spatial inference

The above analyses neglect the potentially confounding influence of the spatial structure of the tumour on our assessments of selection. The trees presented in Figures 1 and 2 were constructed with a maximum parsimony approach that is ‘model-free’. We hypothesised that explicit assessment of spatial structures together with phylogenetic history (a phylodynamics approach) would yield additional information on the birth, death and spatial growth patterns of tumour subclones. The phylogenomics field has developed excellent methods for this task, including Maximum Likelihood (Alves et al., 2019) and Bayesian (Reis et al., 2016) phylogenetic frameworks. Within a Maximum Likelihood framework, the aim is, given a model of evolution *M* with parameters *θ*, and given the data *D*, to find the tree that maximises the likelihood ℒ (*D*| *M, θ*). However, current likelihood/Bayesian methods for phylogenomics like BEAST (Drummond et al., 2012; Suchard et al., 2018), assume well mixed non-spatial population models that do not capture the spatial structure of an expanding solid tumour, and therefore did not make maximal use of the spatial data available in our cohort.

To quantify more accurately the evolutionary dynamics from the genomic data we setup a computational inference framework. We previously presented a spatial model of cancer growth that could be directly employed to perform inference on spatial multi-region sequencing data (Chkhaidze et al., 2019).

We substantially extended this model to incorporate more realistic spatial growth conditions, as well as cell replication, death and branch overdispersion (Figure 4A; Material and Methods). The model was able to produce characteristic clonal patterns both for neutrally expanding tumours (Figure 4B) as well as for malignancies with one (Figure 4C) or two subclones under positive selection (Figure 4D). Our model simulated the accumulation of somatic mutations in each lineage, both neutral and selected, allowing to compare lineages accumulating passenger variants (Figure 4B-D, bottom) versus lineages that were phenotypically distinct in terms of growth (Figure 4B-D, top). In the model we also simulated tumours with spatial constrains where growth could only occur at the periphery, versus malignancies where all cells grow, leading to exponential growth. The balance between these two scenarios was described with a parameter d_push_ that determined the distance which a cell could push neighbouring cells outwards and generate space to divide, even when entirely surrounded by other cells. These different spatial dynamics produced radically distinct subclonal patterns, with peripheral growth (low d_push_) displaying segregating lineages (Figure 4E and S7), and with more increasing clonal intermixing as larger proportions of cells everywhere in the tumour had the ability to grow.

**Figure 4.**
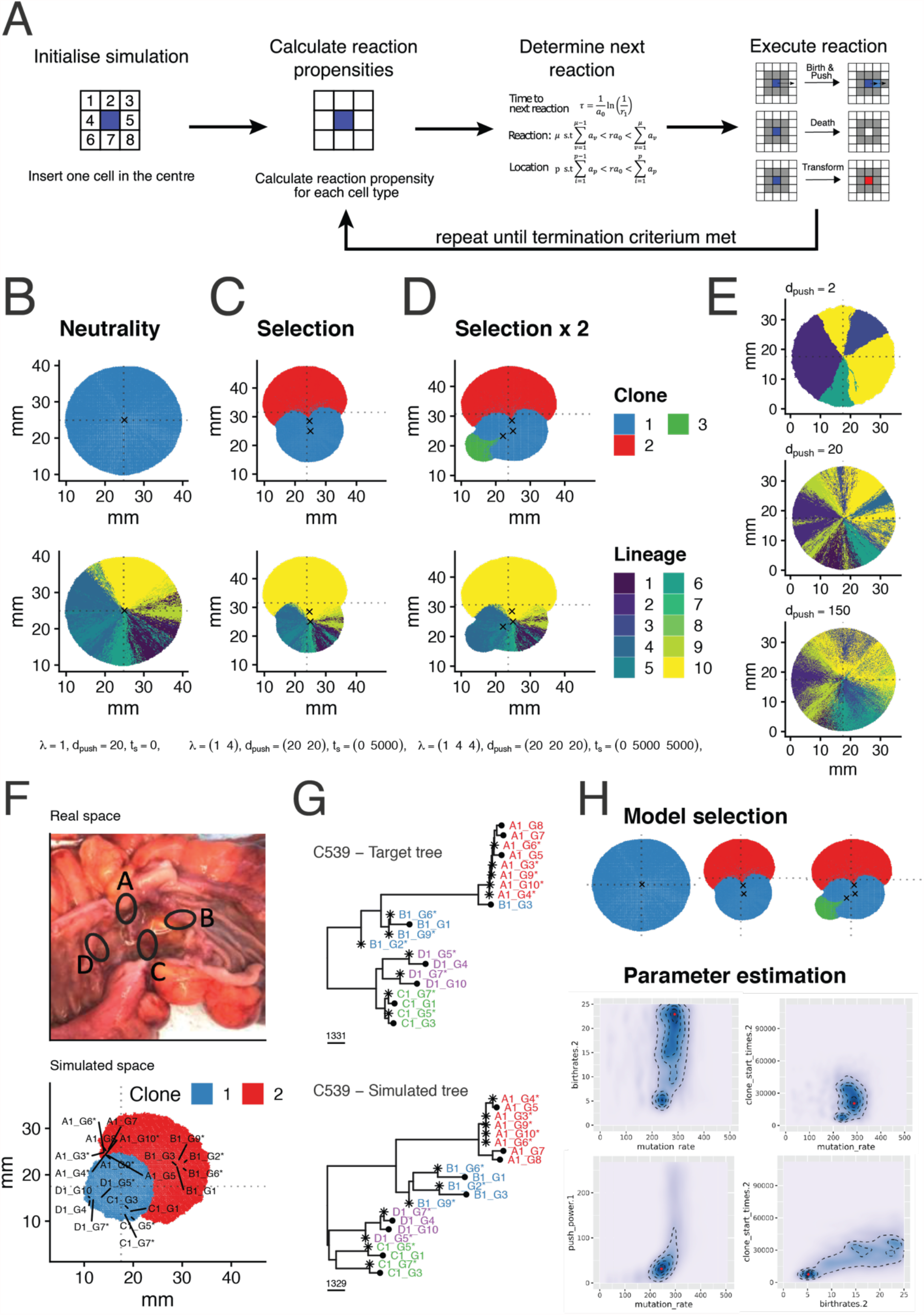
Bayesian inference framework for cancer dynamics in space and time. **(A)** Schematic representation of the spatial cellular automaton model of tumour growth. **(B)** Instance of simulation of a neutrally expanding cancer with a single ‘functional’ clone (blue, top), and corresponding neutral mutation lineages (bottom). **(C)** Simulation of a tumour containing a differentially selected subclone (red, top) and corresponding neutral mutation lineages (bottom). **(D)** Simulation with two branching subclonal selection events. **(E)** In this neutral simulation we illustrate peripheral versus exponential growth and the effects on lineage mixing. **(F)** Spatial sampling annotated during tissue collection for case C539 and corresponding simulated spatial sampling. **(G)** Real data from patient C539 (top) versus simulated data from an instance selected by the inference framework (bottom). **(H)** Inference framework based on Approximate Bayesian Computation -Sequential Monte Carlo (ABC-SMC) allows for model selection and posterior parameter estimation given the data. In this case birthrates.2 is the birth rate of the selected subclone, clone_start_times.2 is the time when the subclone arose during the growth of the tumour, push_power.1 is the coefficient of boundary driven growth and mutation_rate is the rate of accumulation of mutations per genome per division.

We implemented the simulation of our empirical spatial sampling scheme as in Figure 2 and S1, where for every patient we generate synthetic data by sampling the simulated tumour as the real tumour was sampled (Figure 4F). We simulated the accumulation of mutations across the whole-genome in each sampled lineage. This generated realistic appearing whole-genome sequencing data that we then used to reconstruct the a (synthetic) phylogenetic tree to match the real tree (Figure 4G). The model parameters that matched the synthetic trees to the empirical tree were estimated on a patient-by-patient basis using Bayesian inference.

Specifically, we used Approximate Bayesian Computation based on Sequential Monte Carlo (ABC-SMC) (Moral et al., 2012) to infer the model parameters from the data and perform model selection between neutral evolution, selection with one subclone, and selection with two distinct subclones (Figure 4H, see Material and Methods and Figures S8 and S9 for details on inference). In the example of case C539 (Figure 5A), we compared the three models, which have distinct number of parameters *k*, and calculate the negative log-likelihood from the inference. We then regularise with Akaike Information Criterion (AIC) to identify the best model that fit the data. Importantly, we also plot the difference in the AIC values, where for ΔAIC values <4 we considered two models having both substantial evidence for, and hence making it impossible to select one in favour of another (Burnham and Anderson, 2004). We also report the AIC value for increasingly low values of distance between simulation and target data ε performed during the SMC-ABC inference process (Figure 5B), showing the preferred model as the simulations get closer and closer to the data. We then calculated the posterior predictive p-value to assess how often does the selected model predict the data well if we consider stochastic fluctuations (Figure 5C). In this case the inference rejects the neutral model and identifies either selection with one or selection with two subclones as the best model. In Figure 5D we can see the comparison between the phylogenetic tree from the real data and the best inferred tree (Figure 5E). The inference identifies a clearly selected clade in A, where indeed we find a KRAS G12C subclonal mutation. The model with two selected subclones also identifies a possible selected event in the ancestral branch of C and D. Simulation of bulk sequencing of the whole tumour shows the subclonal cluster from selected events as well as the expected neutral tail of passenger mutations within each subclone (Caravagna et al., 2020a, 2020b). Figure 5G shows the spatial sampling of a selected simulation with two subclones coloured in the spatial structure, showing divergent evolution of the two selected events.

**Figure 5.**
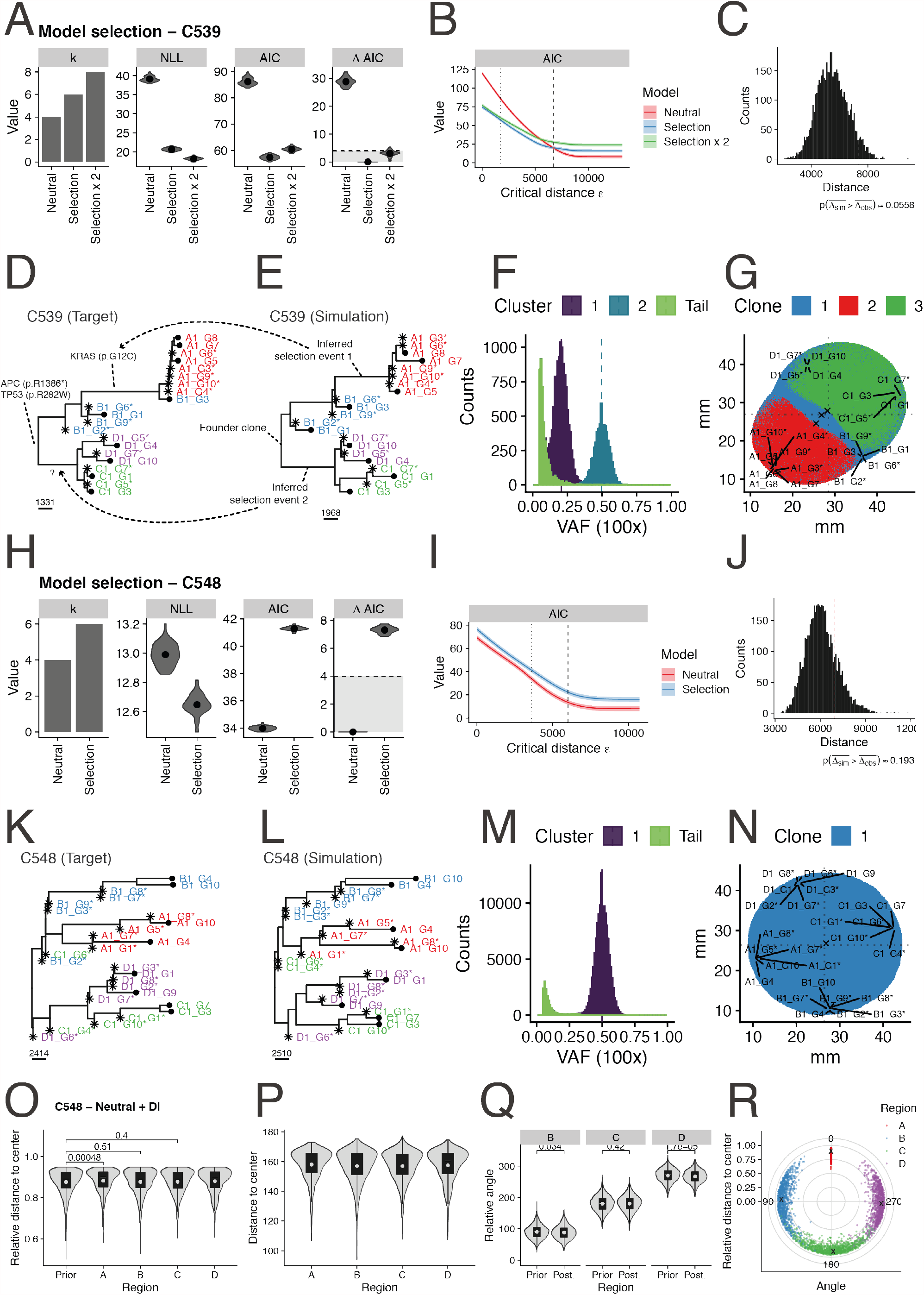
Example results from the inference of evolutionary dynamics. **(A)** Model selection analysis considers the number k of parameters for each model, the negative log-likelihood (NLL) to calculate AIC. Models with AIC differences (ΔAIC) >4 should be considered the preferred model that explains the data. **(B)** AIC value with respect to the distance from the data ε for each of the model. Dotted line is distance used for ABC-SMC. Dashed line is average distance to trees with random uniform noise (* 0.5-2). **(C)** Posterior predictive p-value. Red line is the average distance of the target to simulated trees. **(D)** Target (real) phylogenetic tree for C539. **(E)** Simulated phylogenetic tree for C539 from a simulation identified by the inference. **(F)** Simulated 100x whole-genome sequencing of a hypothetical sampling of the whole tumour. **(G)** Spatial patterns of the simulation that generated the matched data with corresponding sampled regions. **(H)** Model selection for C548. **(I)** AIC value vs ε. **(J)** Posterior predictive p-value. **(K)** Target (real) phylogenetic tree for C539. **(L)** Simulated phylogenetic tree for C548 from a simulation identified by the inference. **(M)** Simulated 100x whole-genome sequencing of a hypothetical sampling of the whole tumour. **(N)** Spatial patterns of the simulation that generated the matched data with corresponding sampled regions. **(O)** Posterior of sampling bias for relative distance from the centre and **(P)** absolute distance from the centre. **(Q)** Posterior of sampling bias for relative angle of sampling. **(R)** Simulated sampling bias in different regions of the tumour.

Differently, for case C548 the inference identified neutral subclonal evolution as the best model (Figure 5H-J). Indeed, the target tree (Figure 5K) and best simulation tree (Figure 5L) are remarkably similar, producing a characteristic neutral bulk sequencing pattern of 1/f tail (Figure 5M). Matched spatial sampling of selected simulation is reported in Figure 5N. Indeed, no subclonal driver events were identified in this case (Figure 2).

In the inference we also incorporate sampling variability to account for sampling biases during tissue collection in terms of relative (Figure 5O) and absolute distance (Figure 5P) from the centre, as well as angle (Figure 5Q) and confirm that prior expected sampling variability was consistent with the data, showing that no unexpected variability in sampling occurred. Sampling points of selected simulations for this case of C548 are reported in Figure 5R.

When we considered the whole cohort (Figure S10), we found strong evidence of subclonal selection by ΔAIC>4 in 6/26 samples (Figure 6A). In 4/6 cases a known subclonal driver mutation was present in the selected clade and the variant was expressed in the RNA (subclone drivers are listed Table SX and highlighted with a box in Figure 2). Those were C525 with subclonal selection in C driven by PIK3CA missense mutation Q546P, C538 with subclonal selection in D driven by RNF43 nonsense mutation Q153*, C539 with subclonal selection in A and part of B driven by KRAS missense mutation G12C, and C531 with subclonal selection in B driven by SMAD4 missense mutation A118V. Corroborating our previous analysis, RNF43 in C538 was the subclonal truncating variant with the highest divergence from the neutral expectation (Figure 3B), and subclonal events in C525 and C539 were also amongst the most divergent from the neutral expectation. Two other cases had strong signs of subclonal selection but did not show evidence of known driver genes.

**Figure 6.**
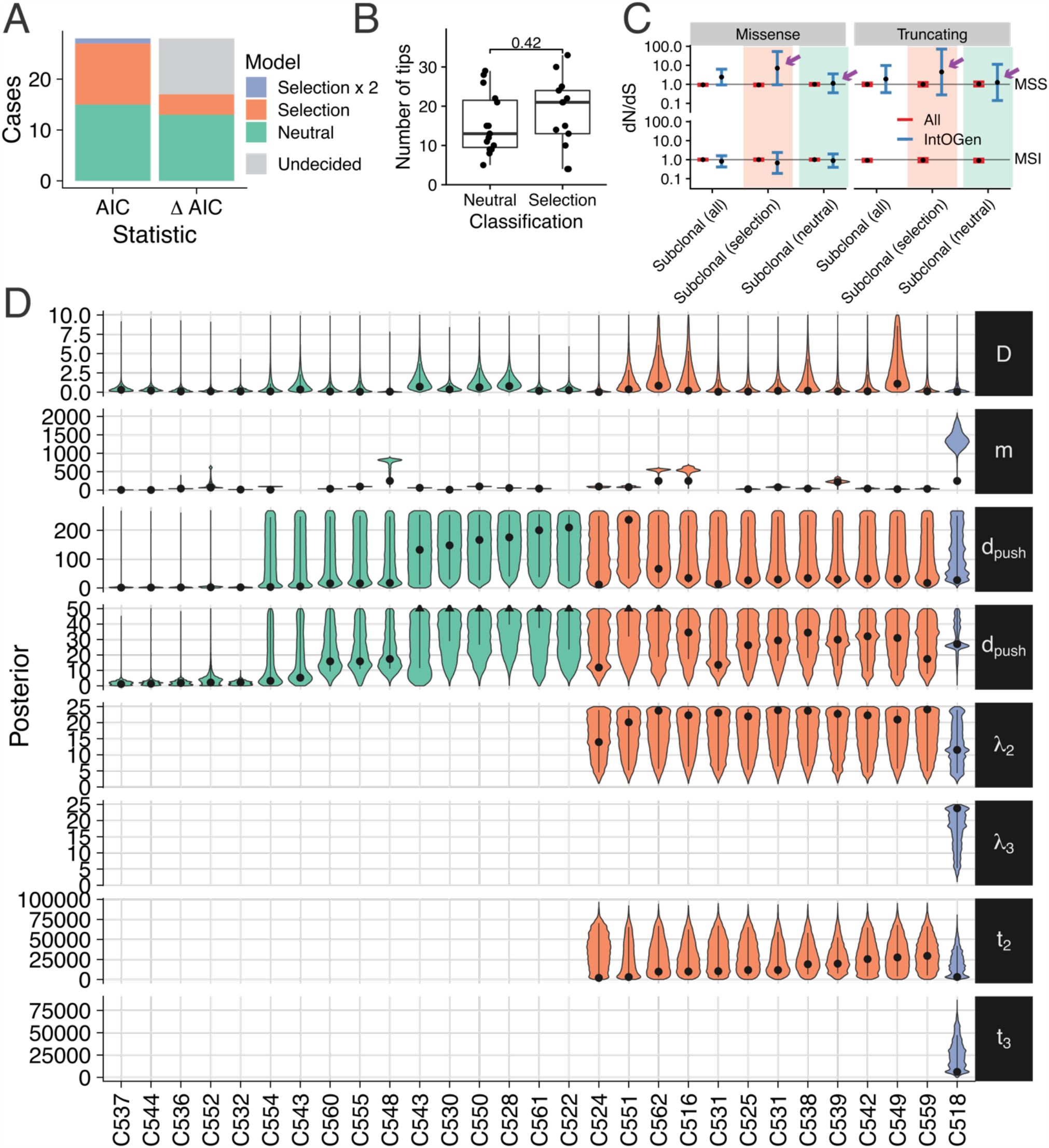
Quantification of evolutionary dynamics across the EPICC cohort. **(A)** Proportion of the cases inferred to contain subclones under selection versus cases explained by neutrality alone. AIC and ΔAIC values are reported, with the latter indicating that a good proportion of cases can be explained by both models. (B) The number of samples per tumour was not associated with a greater frequency of tumours called as having selection. **(C)** Subclonal dN/dS values for the cohort split between selected and neutral by AIC. **(D)** Posterior distributions of the parameters for all cases, split by neutral (green), selected (orange) and selected x 2 (purple).

In other 13/26 cases we found strong evidence for the neutral model to be supportive of the data. In another 7/26 cases ΔAIC could not identify a best model. Considering only AIC, we found 12/26 with selection as the slightly preferred model suggesting that the effects of selection in these cases may be weak. Evidence of selection in the phylogenetic trees included a significantly longer branch containing the selected event (e.g., Figure 5E – selection event 1), or two distinct regions having a more recent common ancestor with respect to the others (e.g., Figure 5E – selection event 2). In 7/12 selected cases, we found a driver event in the corresponding selected clade, although in 3 cases with low ΔAIC the effects on the clonal expansion of the subclonal driver were small. In the other 14/26 cases, the preferred model was neutral (Figure 6A). Number of samples per case (i.e., more extensive tumour sampling) did not confound model selection (Figure 6B).

We then sought orthogonal validation of the inference results. Indeed, splitting the cohort into neutral vs non-neutral based on AIC and recalculating subclonal dN/dS for IntOGen drivers showed increased values of dN/dS in the selected cohort, and values of dN/dS=1 for the neutral cases (Figure 6C). This analysis also supports the absence of subclonal selection even in small clades that may not have undergone sufficient expansion to be detectable by our inference method.

Figure 6D reports the posterior distributions of all the cases, split by neutral vs selected by AIC. We found relatively low overdispersion *D* of branch lengths in the majority of cases. Median inferred mutation rates were 9.8×10^−9^ muts/bp/division in MSS cases and 68.7×10^−9^ muts/bp/division in MSI cases, consistent with previous measurements (Werner et al., 2020).

The d_push_ parameter clearly distinguished fast, exponentially driven tumours that grew throughput their mass, from boundary-driven cancers that only grow at the periphery. The formers were enriched for neutrally expanding cancers. We were able to estimate the increased growth rate of selected subclones, which was at time even 20 times higher than the background clone. The majority of clones also originated relatively early during the expansion of the tumour, when the tumour mass was <50,000 cells.

Hence, selected events significantly driving subclonal expansions seem to arrive early during tumour growth and have high selective coefficients. As a consequence, most bulk sequencing of a single biopsy, generally collected from a single region of the tumour, was unlikely to contain multiple selected subclones and hence neutrality fully explained the data. Moreover, even with considerable spatial sampling, neutral evolution was widespread in colorectal cancer, with half of more cases lacking subclonal selection entirely, and the rest mostly containing a single subclonal driver.

### Non-genetic drivers and *in vivo* characterisation of the epigenome and transcriptome of selected subclones

Subclone evolution within a primary human malignancy is a natural “competition experiment” between human cells with the same genetic background in the same microenvironment. It can be seen as a controlled experiment to assess the phenotypic differences between subclones and the consequences of driver alterations.

We exploited the matched ATAC-seq and RNA-seq data from the same clades to measure the epigenetic and transcriptomic differences between the selected subclone and the background cell population in the 5 cases with sufficient matched ‘omics data where we had inferred subclonal selection was acting, 2 of which had no known driver event.

Enrichment analysis of differentially expressed genes between the subclone and the background clone highlighted consistent dysregulation of focal adhesion pathways for C531, C542 and C559. We also found EMT upregulated in C524, C542 and C551 whereas MYC+E2F targets were upregulated in C524, C531 and C551 (Figure 7A and S11).

**Figure 7.**
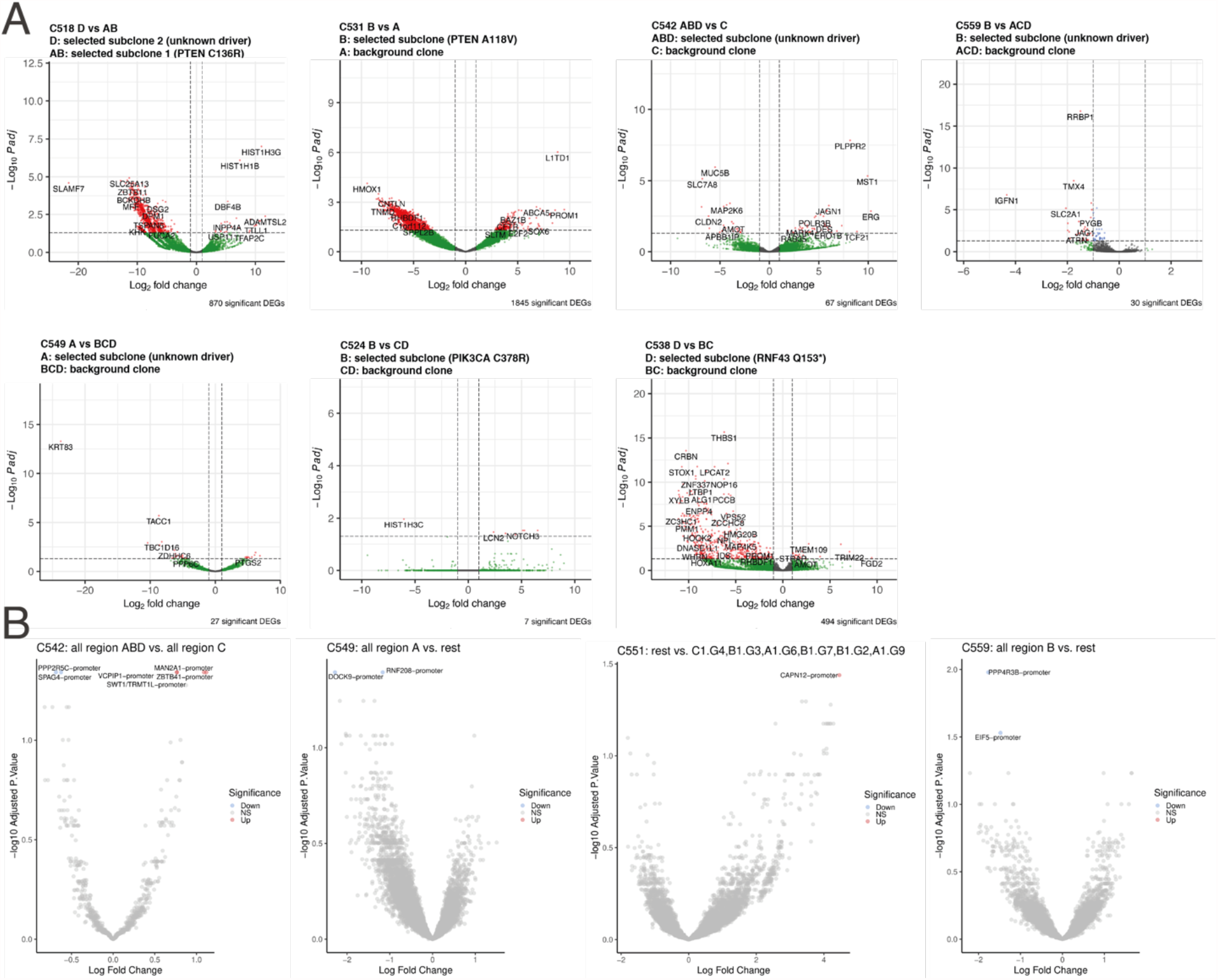
Differential gene expression and chromatin accessibility between detected subclone and backgound clone. **(A)** Differential gene expression between the subclone and background clone for all cases with selection (AIC) for which there was sufficient samples. **(B)** Differential ATAC peak between subclone and background clone for those cases with no detected genetic driver.

We also performed differential ATAC-seq peak analysis between subclone and background clone to identify potential non-genetic drivers, in particular in those cases that lack a subclonal driver event. The results for the four cases with no genetic driver (all but C516 for which we had not enough ATAC samples from the two clones) are represented in Figure 7A. Amongst interesting candidates we found promoter loss of accessibility of PPP2R5C, a regulator of TP53 and ERK.

## Discussion

Here we directly quantify, with spatial multi-region sequencing and inference, the extent of subclonal selection in colorectal cancer. We also measure subclonal characteristics with matched epigenetic and transcriptomic information. We found that colorectal malignancies could be clearly delineated based on their evolutionary dynamics, namely by the presence or absence of subclonal selection, the mode of expansion (peripheral vs exponential growth) and the degree of clonal intermixing. All these are not static molecular features, such as a gene mutation, but instead are measures of the *dynamics* of how the tumour was changing. These may be predictive of unseen metastatic deposits in primary colorectal cancers, as well as being prognostic and predictive evolutionary biomarkers. We are now following these patients longitudinally to prospectively identify those that relapse with metastatic disease within the following five years, versus those that are effectively cured. We will test whether any of the evolutionary measurements above predict such transition.

We show how incorporating an assessment of the evolutionary consequence of candidate driver mutations, through phylogenetic analysis, provides a means of *in vivo* human assessment of a gene’s “driverness”, which could ultimately assist with prioritisation for therapeutic targets. Using this approach, we discovered that many putative driver mutations do not in fact contribute to clonal expansions, and so are unlikely to be drivers of untreated disease. Using matched multi-omic data we also investigated what is the phenotypic change experienced by a subclone with a driver event, both in terms of its epigenome and its transcriptome. We also found that multiple cases of subclonal expansion were not driven by existing candidate genetic drivers, implicating epigenetic events as possible alternative driver alterations.

Overall, we show that assessment of intra-tumour heterogeneity can serve as a ‘controlled experiment’, enabling quantitative measurement of ongoing evolutionary competition within the human body between different lineages with distinct subclonal mutations.

## Materials and Methods

Tumour collection and generation of molecular data is described in the accompanying manuscript (PROTOCOL). Processing of DNA and ATAC-seq data is described in the accompanying manuscript (EPIGENOME), and RNAseq data in (TRANSCRIPTOME).

Inference of evolutionary dynamics using spatially-resolved genomic data was performed by Bayesian fitting of a spatial agent-based model of clonal evolution to the observed molecular data. The model described growth, death, physical dispersion and mutation of individual tumour glands, and was a substantial modification of the framework previously described in (Chkhaidze et al., 2019). Full details are provided in the supplementary mathematical note.

For the gene essentiality analysis, cancer dependency profiles were downloaded from https://depmap.org/broad-sanger/ (version used: CRISPRcleanR_FC.txt) and scaled as described in PMID: 29083409, i.e. making the median essentiality scores of prior known essential and non-essential genes equal to −1 and 0, respectively. The mutational status of selected putative cancer driver genes used to produce the box plots in Figure 3C and to test differential gene essentiality across mutant versus wild-type cell lines, was obtained from the Cell Model Passports (PMID: 30260411).

## Acknowledgments

This study was principally supported by funding from the Medical Research Council (MR/P000789/1 to A.S.) and the Wellcome Trust (202778/Z/16/Z to T.A.G. and 202778/B/16/Z to A.S.). A.S. and T.A.G. were also supported by Cancer Research UK (A22909 and A19771) and the National Institute of Health (NCI U54 CA217376 to D.S., T.A.G. and A.S). This work was also supported by a Wellcome Trust award to the Centre for Evolution and Cancer at the ICR (105104/Z/14/Z). C.P.B. acknowledges funding from the Wellcome Trust (209409/Z/17/Z). B.W. is supported by a Barts Charity Lectureship (grant MGU045). D.R. was partially supported by a Bicocca 2020 Starting Grant and by a Premio Giovani Talenti dell’Università degli Studi di Milano-Bicocca. L.M. is supported by Cancer Research UK (A23110).

## Data availability

Analysed data are available on Mendeley: https://data.mendeley.com/datasets/dvv6kf856g/2. Sequence data have been deposited at the European Genome-phenome Archive (EGA), which is hosted by the EBI and the CRG, under accession number EGAS00001005230. Further information about EGA can be found on https://ega-archive.org.

## Code availability

Complete scripts to replicate all bioinformatic analysis and perform simulations and inference are available at: https://github.com/sottorivalab/EPICC2021_data_analysis.

## Author contributions

T.H. analysed and interpreted the data, designed and implemented the computational inference framework and performed inference analysis. J.H. analysed RNA data. GDC performed copy number analysis. I.S. devise multi-omics protocol, collected the samples and generated the data. C.K. collected the samples and contributed to data generation. H.C., M.M., A.B. and M.S. supported tissue and patient data collection. C.L. contributed to ATAC data analysis. M.M., J.F.M., A.M.B., contributed to data generation. B.J. contributed to phylogenetic analysis. L.Z. contributed to dN/dS data analysis. C.J., E.L., D.N., G.Car. contributed to data analysis. A.B. generated methylation array data. A.V. performed cancer dependency analysis under the supervision of F.I. M.J. contributed to tissue collection. D.S. contributed to experimental design and data interpretation. J.B. contributed to sample collection coordination. M.R.J. supervised sample collection. L.M. contributed to result interpretation. T.A.G. and A.S. conceived and supervised the study, and wrote the manuscript.

