## Supplementary Note for "Assessment of the evolutionary consequence of putative driver mutations in colorectal cancer with spatial multiomic data"

### ABC-SMC inference

In the previous chapters, I have described the mutational landscape of a total of 30 multi-region single glands sequenced colorectal cancers from the EPICC cohort. In this context, I described the rare intermixing of glands from different regions and hypothesised that this might be a phenotypic property of some cancers.

Further, I described the status of driver mutations previously identified in other studies (e.g., Muzny et al. 2012; Martínez-Jiménez et al. 2020; Martincorena et al. 2017). The vast majority of these driver mutations were found to be clonal mutations present in all glands taken from different regions of the tumour. These mutations likely accumulated before the initiation of the growth or have been part of a subclone that effectively swept through the population. Nevertheless, a few examples of potential subclonal driver mutations (e.g., one KRAS p.G12C and seven PIK3CA mutations) were identified. In one of these cases, C539, the mutation was accompanied by an evident elongation of the associated branches of the phylogenetic tree. This branch elongation indicates that the associated glands share a most recent common ancestor that went through many more generations than the background, the 'hallmark' of subclonal selection.

While the presence of a selected subclone, or more broadly speaking, changes of the evolutionary dynamics, seemed rather obvious in this case, it was unclear how to interpret the information contained in the tree shapes of other cases. The same problem exists in many previous studies of the subclonal diversification at primary sites (Yates et al. 2015) or during metastasis (Gundem et al. 2015; Yates et al. 2017;

Noorani et al. 2020). Despite being very impressive, these studies have provided little functional insights into the evolutionary dynamics driving these processes. Specifically, it remains unclear what degree selection of adaptive phenotypic properties plays a role in the later stages of cancer evolution, and the effect occasionally observed subclonal driver mutations have.

The majority of cancer driver genes were previously identified based on their recurrence across patients using complex statistical models (e.g., mutSigCV) to quantify the degree to which mutations are overrepresented (e.g., Lawrence et al. 2014; Martínez-Jiménez et al. 2020). Still, identical mutations might have little or no effect in some genetic or environmental backgrounds, and often complex analysis and tedious experiments are required to uncover these relationships. An excellent example of context-dependent selection can be found for PTEN mutations in prostate cancers and leukaemia (Berger, Knudson, and Pandolfi 2011). In these tissues, the incomplete loss of PTEN in a TP53 wildtype background is tumorigenic, whereas total loss of PTEN leads to the induction of senescence and hence no tumour formation. If complete PTEN loss does instead occur after the loss of TP53 (i.e., in a TP53 mutant context), more aggressive tumour growth is observed. In order to study the effect of such driver alterations *in vivo* mice models are often used, but these are often costly, time-consuming and require specific hypothesis to test.

For self-evident reasons, longitudinal observation of solid tumours in their primary site can normally[1] not be conducted in humans. For this reason, the effects that driver mutations have at the primary site are not well studied. Here I will apply a spatial computational inference framework to single-gland multi-region WGS data. Doing so, I will demonstrate how this approach can indirectly gain insight into the fitness effect of naturally arising somatic mutations in primary CRC. This approach allows identifying relevant alterations occurring in primary tumours, which could be validated subsequently in controlled *in vivo* experiments. As such, this allows prioritisation of relevant alterations observed in primary tumours for valida-

[1]Some cases untreatable tumours or refusal of treatment can give the rare opportunity to do this.

tion. Unlike other studies based on bulk sequencing data (e.g., Dentro et al. 2021), this approach has sufficient power to infer subclonal selection and also allows to characterise the specific genetic background that putative driver mutations occurred in (i.e., their respective lineage).

#### 6.1 Bayesian statistics

In order to understand the evolutionary dynamics observed in cancer genomic data, we can not resort to our intuition or descriptive statistics. Instead, one optimally wants a statistical model that captures the process underlying measured data well enough to allow to gain significant insight into it (Box, Launer, and Wilkinson 1979). Aspects of the model that influence its behaviour, the model parameters, can then provide a more interpretable summary of the data.

Several methods to fit such a model to actual observations (i.e., statistical inference) exist. In order to apply most of these, one needs to be able to calculate the likelihood function $p(D|\theta)$, defining the probability of observing the data $D$ under a given set of parameters $\theta$ from the parameter space $\theta \in \Theta$.

With a likelihood function available classic frequentist methods can be used to identify parameters under which it would be most likely to observe the data, the maximum likelihood estimation (MLE $\hat{\theta} = \arg\max_{\theta \in \Theta} p(D|\theta)$). This ML estimation is possible even if a closed form for the MLE is not available or hard to obtain.

An alternative approach, so called Bayesian inference, is to use Bayes' theorem to instead calculate a probability distribution over the parameter space $p(\theta|D)$ the so called posterior distribution or short posterior. From Bayes' theorem it follows that

$$p(\theta|D) = \frac{p(D|\theta)p(\theta)}{p(D)} = \frac{p(D|\theta)p(\theta)}{\sum_{\theta' \in \Theta} p(D|\theta')p(\theta')}$$

where $p(D|\theta)$ is the likelihood and $p(\theta)$ is a probability distribution over the parameters space, the so called prior likelihood or short prior.

The likelihood of the data $p(D)$ can be interpreted as a normalisation constant. Dropping this results in a distribution that is proportional to the actual posterior likelihood but does not sum to one

$$p(\theta|D) \propto p(D|\theta)p(\theta)$$

##### 6.1.1 Approximate Bayesian Computation

Unfortunately, a likelihood function for the spatial distribution of mutations in a growing tumour, potentially with several differently fast-growing subpopulations (i.e., neutrality vs selection), cell death and various modes of growth (boundary driven growth vs exponential growth) is not available and probably intractable.

Nevertheless, since it is possible to simulate the underlying process, a class of algorithms that allows to perform Bayesian inference without a likelihood function can be used. These Approximate Bayesian Computation (ABC) methods use a generative process to approximate the likelihood conditional on a set of parameters $\theta$ (Karabatsos and Leisen 2018).

For this purpose ABC methods require a model $f(\cdot|\theta)$ from which random realisations can be drawn $D^* \in \mathscr{D}$ a prior distribution $p(\theta)$ on the set of the inferred parameters $\theta \in \Theta$ multiple summary statistics $\eta : \mathscr{D} \rightarrow S$ as well as a distance function $\rho : S \times S \rightarrow \mathbb{R}^+$ and a critical distance $\varepsilon \in \mathbb{R}^+$ below which random observations are assumed to match the observed data $D$. With these we then seek to sample from the marginal posterior distribution

$$p(\theta, D^*|D, \varepsilon) = \frac{p(\theta) f(D^*|\theta) \mathbb{I}_{A_{\varepsilon,D}}}{\int \pi(\theta) f(D^*|\theta)\, dD^* d\theta}$$

where $\mathbb{I}\mathbb{A}_{\varepsilon,\mathbb{D}}(x)$ indicates whether $x$ is an element of the set $A_{\varepsilon,D}$ of observations with

$$A_{\varepsilon,D} = \{z \in \mathscr{D} : \rho(\eta(D), \eta(D^*)) \leq \varepsilon\}$$

ABC methods were pioneered in the field of population genetics by Tavaré et al. (1997) to infer coalescence times from DNA sequence data and (Pritchard et al. 1999) to study the evolution of the Y chromosome. Since then such methods have been used extensively (Underhill et al. 2000; Kaessmann et al. 2001; Glover et al. 2013). Examples of the application of ABC methods in other fields include molecular biology (Woods and Barnes 2016), pharmacology (Picchini 2014), epidemiology (McKinley, Cook, and Deardon 2009; Tanaka et al. 2006) or indeed cancer evolution (Sottoriva et al. 2015; Williams et al. 2018b).

Various extensions of the brute-force accept-reject method used by Tavaré et al. (1997) and (Pritchard et al. 1999) exist, these seek to combine ABC with other algorithms to increase the efficiency of sampling in the parameter space. Examples include ABC-MCMC (Marjoram et al. 2003; Wegmann, Leuenberger, and Excoffier 2009), ABC-SMC (Sisson, Fan, and Tanaka 2007; Del Moral, Doucet, and Jasra 2012; Filippi et al. 2013) or ABC-PMC (Beaumont et al. 2008; Baragatti, Grimaud, and Pommeret 2012; Murakami 2014). Other modifications seek to replacement the rejection based approximation of the likelihood with alternative estimators (see Karabatsos and Leisen 2018, for details). One example of this, which will later be used for the calculation of the expectation of the posterior predictive likelihood, are synthetic likelihoods (SL) similar to the method proposed by (Wood 2010). In the context of ABC, these methods make the assumption that the distribution of summary statistics follow a specific distribution. With this assumptions likelihoods for a critical distances $p(D \leq \varepsilon) \ll 1/N$, that are impermissible to be used with rejection based methods, can be approximated. More detail on SL will be provided below.

Two of these algorithms will be used for the ABC inference of parameters describing the growth dynamics in individual tumours i) rejection sampling described first by Pritchard et al. (1999) (see Chapter 6.2.4.1, page 186 for details) and ii) the ABC-SMC algorithm proposed by Del Moral, Doucet, and Jasra (2012) (see Chapter 6.2.4.2, page 187 for details). In brief, the number of subclones $max(i)$, their relative growth rate $\lambda_i$ compared to the ancestral clone $i = 0$, the respective coalescent population sizes $t_i$ and a global parameter $d_{push}$ describing the distance from the outer rim of the tumour at which glands can grow and "push" outwards will be inferred. It will also explore if a global increase of death rates $\mu$ can explain the observed data better. A overview of all the inferred and constant parameters of the model can be found in Table 6.1 (page 186). A more detailed explanation of the simulation setup and how a simulated sampling scheme equivalent to the used one was generated will be provided in Chapter 6.2.1 and Chapter 6.2.2 (pages 181, pages 182), respectively. A summary of the statistics and distance metrics used to compare the observed data to simulated datasets will be provided in Chapter 6.2.3 (pages 184). A general overview over the inference framework is shown in Figure

6.1.

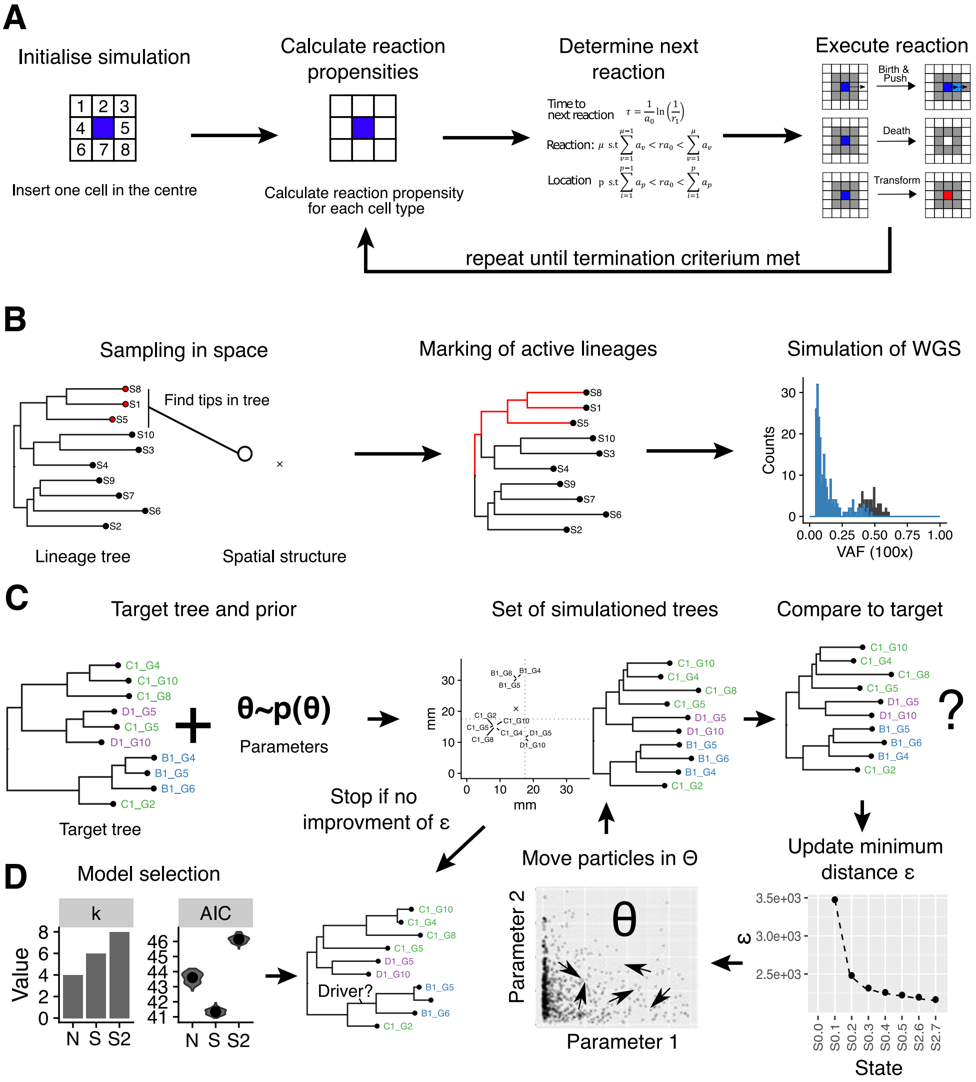


**Figure 6.1:** The ABC-SMC inference framework. A) The spatial simulation of the tumour using the Gillespie algorithm. B) The simulation of WGS sequencing data. For this, a subset of cells are selected in space, then active linages in the tree are marked, and finally, the active part of the tree is traversed to simulate WGS data. C) Using a ABC-SMC algorithm (Del Moral, Doucet, and Jasra 2012) parameters of each model are inferred. D) To the fitted models, a model selection procedure is applied to select the best model.

### 6.2 Methods

#### 6.2.1 Spatial simulations

As outlined above, a model $f(\cdot|\theta)$ from which one can sample simulated observations $D^*$ given a set of parameters $\theta$ is required to apply ABC based inference to the whole-genome sequencing data described in the previous chapter. For this, a slightly modified version of the spatial tumour simulator (Chkhaidze et al. 2019) described before (Chapter 3) will be used and generate synthetic sequencing data according to a spatial sampling scheme that is equivalent to the one used to generate the actual data (Chapter 4). For the inference, simulations of a two-dimensional tumour were used. This choice was made, based on the observation that colorectal cancers grow, at least during the initial stages, primarily in a two-dimensional plane through crypt fission (Greaves et al. 2006; Chen et al. 2005; Shen et al. 2005; Bernstein et al. 2008). While this might be a simplification, the simulation of a tumour in two dimensions should still give some insight into a three-dimensional tumour's general growth dynamics.

In well and moderately differentiated colorectal carcinomas the majority of the tumour consists of glandular structures (Fleming et al. 2012; Nagtegaal et al. 2020). These structures are reminiscent of the crypts, small finger-like invaginations into the underlying tissue that normally form the colorectal epithelium (Humphries and Wright 2008). Similar to normal crypts, these glands are assumed to expand spatially through a process of gland fission (Graham et al. 2011; Garcia et al. 1999; Bruens et al. 2017), which also drives neoplastic growth of colon tumours (Wong et al. 2002; Preston et al. 2003). In summary, individual glands are the "clonal units" of colorectal cancers (Baker et al. 2014), and for this reason, each cell of the spatial simulation will be assumed to represent a single gland. Due to the fast replacement of stem cell lineages compared to the rate of crypt fission, the complexity of the population structure within each gland will also be disregard.

The diameter of colonic crypts in normal colon tissue is about 60 microns, and the colorectal epithelium contains roughly 100 crypts per square millimetre of colon (Nguyen et al. 2010). Assuming a similar number of glands per square millimetre

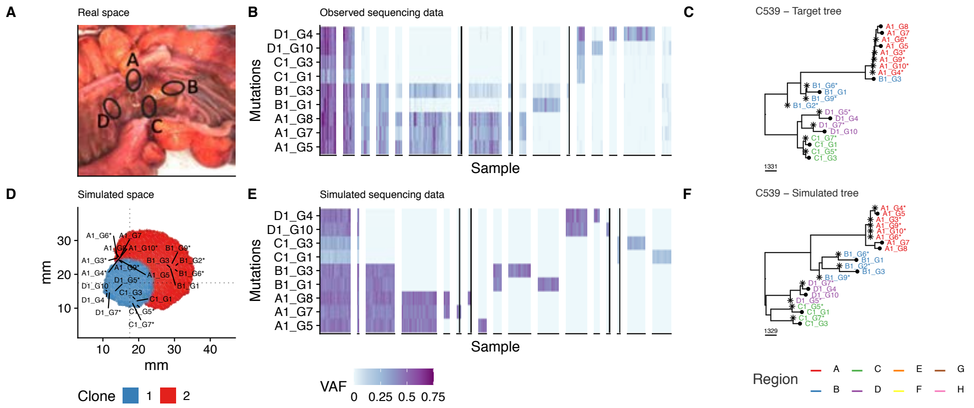


**Figure 6.2:** An illustrative example of simulated and observed WGS sequencing data. **A** and **D** show the spatial layout of the tumour in space and the sampling locations for the actual and simulated tumours, respectively. **B** and **E** summarise the VAF of actual and simulated WGS sequencing data for deep WGS samples, respectively. **C** and **F** show the actual and simulated trees reconstructed from the data, respectively.
VAF: variant allele frequency,

of tumour tissue, a colon tumour with a diameter of $\approx 3.5\,cm$ can be represented by a grid size of $350x350$ (Figure 6.2D). Such a tumour would contain $\approx 96,000$ glands, of which each is consists of $\approx 2,000 - 10,000$ cells. The number of $\approx 10^9$ cells simulated by these $350x350$ simulations are roughly similar to that present in human malignancies (Del Monte 2009) and are still fast enough $\leq 3\,s$ to allow the generation of a sufficient number of simulations for ABC inference.

#### 6.2.2 Equivalent sampling scheme

After the generation of a simulated tumour, I tried to generate a set of samples that reflected the sampling schema used (Figure 6.2A) to generate the single-gland WGS data (Figure 6.2B). For this, a random angle *phi* from the centre of the tumour $O(x_O, y_O)$ along which the first sample region (i.e., the centre of the region 'A') should be placed is generated first:

$$\phi_A \sim U(0, 2\pi)$$

For each region A-D a offset $\phi_i'$ was added to $\phi_A$. Here two different methods were used i) a constant offset where $\phi' = (0, 0.5\pi, \pi, 1.5\pi)$ for the regions A-D, respectively and ii) a randomly varied sampling schema with relative angles between adjacent regions sampled from a Dirichlet distribution $\mathbf{x}_\phi = Dir(K = 4, \alpha)$ with

$\alpha = (0.25m_\phi, ..., 0.25m_\phi)$. Here $m_\phi$ denotes the prior strength of the prior and the angle offsets $\phi'_i$ are given by $\phi'_i = 2\pi \sum_{j=0}^{i} x_{\phi,j}$.

Along each of these vectors the distances to the most distant occupied grid point $\mathbb{I}_o(x,y)$ (i.e., the edge of the tumour) were searched using a half-interval search between $O$ and the edge of the simulated space, to identify:

$$d_{e,i} = \underset{r \in [0, d_max]}{\arg\max}\ r\, \mathbb{I}(r\, cos(\phi_A + \phi_i) + x_O, r\, sin(\phi_A + \phi_i) + y_O))$$

In cases with a high death rate, a high number of grid points within the centre of the tumour are empty. Hence, positions were only assumed to be empty when grid points along the vector defined by $\phi$ up to a distance of 10 from the evaluated position were also unoccupied. At the most extreme values of the parameter range considered ($\mu \leq 0.5$ and $d_{push} = 1$) up to $\approx 8.5\%$ of grid points in the centre of the tumour can be empty, but even at these values, the observation of 7 or more consecutive empty grid points is very unlikely ($\ll 10^-7$).

After the identification of the distance to the edge $d_{e,a}$ the centre of sampling regions were placed at a relative position $x_e \in [0,1]$ along the vector with the coordinates being given by

$$x = [x_d\ d_{e,a}\ cos(\phi_A + \phi_i) + x_O]$$

$$y = [x_d\ d_{e,a}\ sin(\phi_A + \phi_i) + y_O]$$

Similar too the angle offsets $\phi'_i$, two different methods were used to define $x_e$ i) a constant fixed value of $x_e = 0.75$ and ii) a random value sampled from a Beta distribution with a given prior strength $m_d$ and mean $\mu_d$ with $x_e \sim B(\mu_d\, m_d, (1 - \mu_d)\, m_d)$.

After the definition of the centre of the four regions, random grid points within a rectangular area of edge length $d_b$ around these were sampled randomly without replacement until the required number of samples from the region were obtained. Sampled unoccupied grid points were rejected. Figure 6.2A and 6.2D show an illustrative example of the equivalent sampling scheme described above applied to a simulated tumour and the actual macroscopic sampling locations in the real tumour, respectively.

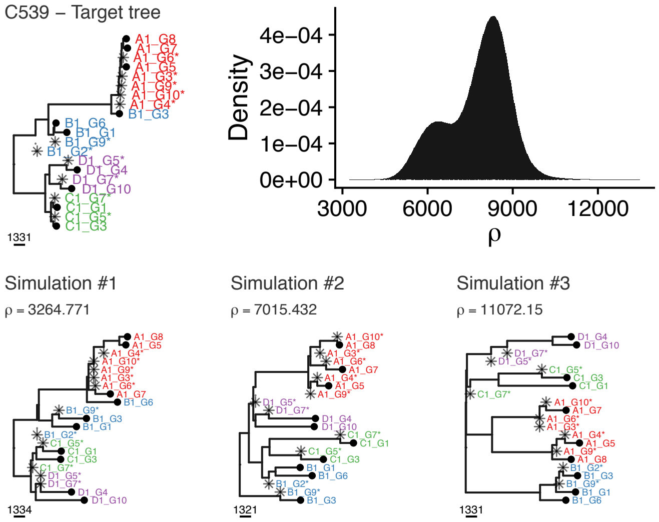


**Figure 6.3:** Simulated trees with different distance $\rho$ to target tree on the top left. On the top right the distribution of distances between the target and simulated trees is shown.

##### 6.2.3 Distance function

Due to spatial information contained in the measured data, a distance metric that is not indifferent to the sample labels itself was used, and labels of samples with the same characteristics were swapped instead (i.e., sequencing type and sample region) to minimise the distance metric. Since clonal mutations do not inform on the subclonal dynamics, these were removed from the trees $T$ and $T'$ prior to the calculation of the distance between them. Next, the patristic distances $d(i,j)$, that is the sum of the lengths of the edges that link two nodes $i$ and $j$ in the tree, for all pairs of tip nodes, was determined and scaled by the tree height, i.e., the maximum distance from the root 0 to a tip.

For a maximum parsimony tree without any homoplasy this is equivalent to the following distance calculated from the mutation data itself

$$d_{j,k} = (|M_j \cup M_k| - |M_j \cap M_k|)/max\{|M_i| : i \in 1,...,n\}$$

where $M_i$ denotes the set of mutations found to be present in the sample associated with the tip $i$ and $n$ is the total number of tips present in the tree.

The distance between two trees $T$ and $T'$ is then calculated from the differences of the scaled patristic distances using the L2-norm:

$$\rho(T,T') = \left( \sum_{i=0}^{|V^1|} \sum_{j=i}^{|V^1|} \left( \frac{d_T(i,j)}{h(T)} - \frac{d_{T'}(i,j)}{h(T')} \right)^2 \right)^{\frac{1}{2}}, h(T) = \max_{i \in V^1} d_T(0,x)$$

This distance takes into account the tip labels of the tree, but samples of the same type (i.e., WGS or LP-WGS) and region (i.e., A-D) can be considered to be equivalent. Accordingly, such equivalent tip labels in the tree $T'$ were swapped until the distance between both trees could not be reduced by swapping any additional labels as outlined in Algorithm 3.

**Algorithm 3:** Label swapping in trees

```
Data: Trees T and T*
Result: Minimised distance between T and T*
Initialise list L of all label pairs in T* of same type and region.;
T*' ← T*;
do
    T* ← T*';
    d ← Δ(T,T*);
    Δd ← 0;
    foreach l ∈ L do
        T_l*' ← T* with labels l swapped;
        d_l ← Δ(T,T*');
        Δd_l ← d_l − d;
        if Δd_l < Δd then
            T*' ← T_l*';
            Δd ← Δd_l;
while Δd < 0;
return (d)
```

It is worth noting that this gradient descent does not necessarily result in the tree with the smallest distance possible, which could only be found by exploring all swaps. As the number of possible ways to label a tree is $\prod_{i=0}^{|N|} = N_i!$ where $N$ are the labels in the label group $i$, this would be infeasible for all but the smallest trees. For the tree shown in Figure 6.3 for example there are $N = (2,2,2,2,2,3,3,5)$ labels per group resulting in $138,240$ trees for which the distance $\rho$ would have to be calculated. This is computationally infeasible and instead, only find the closest local optimum is searched.

In Figure 6.3 a couple of simulated trees with variable distances to a given target are shown to illustrate how changes of the tree topology and branch length are reflected in the distance metric.

##### 6.2.4 ABC algorithms

As mentioned before, two ABC inference algorithms were applied to the datasets to conduct the statistical inference. In the following, the simple ABC rejection sampling algorithm (Pritchard et al. 1999) will be described first. As this method severely suffers from the "curse of dimensionality", a more complex ABC-SMC algorithm (Del Moral, Doucet, and Jasra 2012) that is less affected by this problem will be described following this.

The parameters infer using the ABC algorithms are summarised in Table 6.1. The death rate $\mu$, mutation rate $m$ and "push distance" $d_{push}$ were assumed to be global properties of the tumour, whereas the number of subclones $max(i)$ and the associated birthrate $\lambda_i$ and clone start time $t_i$ are assumed to be clone specific parameters.

**Table 6.1:** Overview over model parameters for the spatial tumour model. Variables with a subclone index $i$ are set individually for each subclone. All other variables are assumed the be constant for the whole tumour.

| Symbol | Name | Description | Limits |
|---|---|---|---|
| $max(i)$ | Subclone number | Number of subclones | [0,2] |
| $\lambda_i$ | Birthrates | Rate of cells division | [1,20] |
| $t_i$ | Clone start times | Population size at introduction | $[1, \lfloor N_{max}/2 \rfloor]$ |
| $a_i$ | Fathers | Index of Ancestor | $i-1$ |
| $\mu$ | Deathrates | Likelihood of death during division | $0 \vee$ [0,0.5] |
| $d_{push}$ | Push distance | Distance from the edge cells grow | $[0{,}x/2]$ |
| $m$ | Mutationrates | Number of mutations during division | $\sim h(T^*)$ |
| $d_b$ | Sample box size | Diameter of the sampling region | [15,25] |

###### 6.2.4.1 ABC rejection sampling

The simplest ABC inference algorithm is that of rejection sampling. While this brute force method can be used the obtain an approximation of the posterior, it is computationally very costly as a time proportional to the density of the prior on $\theta$ is spent on the generation of samples, even in regions with very low probability.

Due to this shortcoming, many more efficient alternative algorithms do exist. Still, due to its simplicity, this algorithm will be used to test the output of the ABC-SMC algorithm described later and provide a short introduction to ABC in general.

The ABC rejection algorithm was first used by Tavaré et al. (1997) and any a more generalised version, introducing the explicit definition of a distance function $\rho(\eta(D^{*}), \eta(D)) \leq \varepsilon)$ by Pritchard et al. (1999). The general procedure used in both papers is principle identical and described by the Algorithm 4.

**Algorithm 4:** ABC rejection sampler

**Data:** Target $y$, distance function $\Delta$, prior distribution $p(\theta)$, simulator $f(\cdot|\theta)$.
**Result:** A set of N particles $P$ approximating the posterior distribution.

```
for i ← 0 to N do
    repeat
        θ ~ π(·); // sample parameter from prior
        z ~ f(·|θ); // generate simulation
    until Δ(y,z) ≤ ε;
    P ← P ∪ {θ}; // append θ to set of particles
return (P)
```

This procedure results in the generation of a set number $N$ of particles for which the distance between the summary statistics of simulated and observed data fall below a predetermined distance threshold $\varepsilon$. These particles can be used to approximate the posterior distribution of the parameters $\theta$.

###### 6.2.4.2 ABC-SMC algorithm

Due to the general shortcomings of the simple rejection ABC algorithm, especially in high dimensional problems with many parameters, a different ABC inference algorithm was used for most cases. This algorithm is an adaptation of sequential Monte Carlo methods for the ABC context and was described by Del Moral, Doucet, and Jasra (2012). It has a number of advantage over previous ABC-SMC methods described by others (e.g., Sisson, Fan, and Tanaka 2007; Toni et al. 2009; Beaumont et al. 2009). First, it has a linear complexity $O(N)$ in the number of particles $N$ instead of $O(N^2)$ for these previously proposed methods. Secondly, the ABC-SMC algorithm by Del Moral, Doucet, and Jasra automatically adjusts the distance threshold $\varepsilon_n$, whereas other methods require the explicit definition of a distance schedule. The careful adjustment of this schedule on $\varepsilon$ is often critical, as a too fast reduction can reduce the performance of the inference or even lead to its collapse (Del Moral, Doucet, and Jasra 2012). The ABC-SMC algorithm conceived by Del Moral, Doucet, and Jasra only requires the definition of one parameter $\alpha$, which controls how fast the critical distance $\varepsilon$ is reduced.

The SMC algorithm consists of a number of steps, in which a set of $N$ particles is updated repeatedly. Each $i = 1,...,N$ of these particles consists of a set of parameters $\theta^{(i)}$, $M$ simulated observations $X^{(i)}_{j=1...M,n=0}$ and a weight $W^{(i)}_{n=0}$. The weight of each particle is proportional to the number of simulated observations $j \in \{1,...,M\}$ that are part of the set $A_{\varepsilon_n,y} = \{z \in D : \rho(y,z) < \varepsilon_n\}$ and can be calculated as:

$$W_n^{(i)} = \frac{\sum_{j=0}^{M} \mathbb{I}_{A_{\varepsilon_n,y}}(X^i_{j,n=0})}{\sum_{i'=0}^{N}\sum_{j=0}^{M} \mathbb{I}_{A_{\varepsilon_n,y}}(X^{i'}_{j,n=0})}$$

where $\mathbb{I}_{A_{\varepsilon,y}}(x)$ indicates whether the observation $x$ is an element of the set $A_{\varepsilon_n,y}$.

The Effective Sample Size ($ESS$) of a set of particles with the set of weights $\{W_n^{(i)}\}$ is:

$$ESS(\{W_n^{(i)}\}) = \frac{1}{\sum_{i=0}^{N} W_n^{(i)^2}}$$

This $ESS$ is a measure of the complexity of the particle set (Liu 2008) and used to update the distance threshold $\varepsilon$ and trigger a resampling of particles in different steps of the SMC. The individual steps of the ABC-SMC algorithm are described in detail below:

0. **Initialisation** At the beginning of the SMC, i.e., at step $n = 0$, a initial set of $N$ particles is generated. For each of these parameters and random, simulated observations are generated:

$$X^{(i)}_{j,n=0} \sim f(\cdot,\theta^{(i)}), \theta^{(i)}_{n=0} \sim p(\theta), \text{for } j = 1,...,M \text{ and } i = 1,...,N$$

   As the distance threshold is initialised as $\varepsilon_{n=0} = \infty$, with $W^{(i)}_{n=0} = 1/N$ and $ESS = N$ at this point.

1. **Update of weights and ESS** At the beginning of each SMC step $n \leftarrow n+1$. Then the distance threshold $\varepsilon_n$ gets updated so that $ESS(W_n^{(i)}) \leq \alpha ESS(W_{n-1}^{(i)})$

for the new value. If the relative reduction of the distance threshold $\Delta\varepsilon_{rel} = \frac{\varepsilon_{n-1}-\varepsilon_n}{\varepsilon_{n-1}}$ falls below a critical value the inference is terminated at this point.

2. **Resampling step** If $ESS(W_n^{(i)}) \leq N_R$ a resampling step is triggered. For this, $N$ particles are sampled from the set of all particles with resampling at a rate proportional to the particle weight. After the resampling all particle weights are set to $W_n^{(i)} = \frac{1}{N}$.

3. **MCMC step** For each particle $i$ with $W_n^{(i)} > 0$ a proposed new parameter set $\theta_i^*$ is sampled from the transition kernel $\theta_i^* \sim K(\theta_i)$. With the $\hat{\Sigma}$ is the empirically estimate of the covariance obtained from the particle parameters $\{\theta_n^{(i)}\}$ weighted by $\{W_n^{(i)}\}$ existing at the time point $n$.

   For each of these proposed new parameter values $\theta^*$, a random set of observations $X_{1:M}^*$ are sampled: $X{*}_j^{(i)} \sim f(\cdot|\theta_i^*)$, where $j = 1,...,M$. Each proposed particle is then accepted with the likelihood given by the Metropolis-Hastings ratio:
$$A(X{*}_j^{(i)}, X_{1:M}^{(i)}) = min\left(1, \frac{\sum_{j=1}^{M} \mathbb{I}_{A_{\varepsilon,y}}(X{*}_j^{(i)})}{\sum_{j=1}^{M} \mathbb{I}_{A_{\varepsilon,y}}(X_j^{(i)})} \frac{q(\theta,\theta^*)}{q(\theta^*,\theta)}\right)$$
   At which point the algorithm continues at step 1.

**Parametrisation of the ABC-SMC** For the purpose of the ABC inference conduced here a value of $\alpha = 0.95$ was used in all cases. For the transition kernel a truncated multivariate normal distribution from the *tmvtnorm* package (Wilhelm and G 2015) was used, giving a proposal of $\theta_i^* \sim MVN(\theta_i, 2\hat{\Sigma}, \mathbf{a}, \mathbf{b})$. With the $\hat{\Sigma}$ being the empirically estimate of the covariance obtained from the particle parameters $\{\theta_n^{(i)}\}$ weighted by $\{W_n^{(i)}\}$ existing at the time point $n$. The upper and lower limits $a$ and $b$ for each parameter set are listed in Table 6.1.

As many different spatial samples from a single simulated tumour can be generated, a second layer of sampling doing this was introduced. The number of multiple observations drawn from a simulation are denoted as $M'$ below. The number of independent realisations of simulations as $M$ instead. In general, two setups were used: i) $N = 500$, $M = 25$ and $M = 100$ for the first set of models with fixed sampling position parameters $x_e = 0.75$, $\phi_i = (0, 0.25, 0.5, 0.75)$ and $d_b = 25$ and ii)

$N = 5000$, $M = 1$ and $M = 100$ for a second set of models with variable sampling positions with prior strength $m_\phi = 50$ and $m_d = 20$ (see Chapter 6.2.2 for details). In all cases the threshold at which resampling was triggered was set to $N_R = 0.75N$.

**Termination criteria** The ABC-SMC algorithm was terminated when the relative distance reduction $\Delta\varepsilon_{rel}$ decreased below 1% for $t_{\Delta\varepsilon} > 3$ steps (assumed convergence) or until a total of $n = 40$ steps were run. In a couple of cases, a larger value for $t_{\Delta\varepsilon}$ was tested to evaluate if, after longer mixing of the chain, a further decrease in $\varepsilon$ could be archived. An example of a distance schedule derived by the algorithm and the associated statistics of the ABC-SMC chain are shown in Figure 5.11. In this example, the algorithm's parameters were $\alpha = 0.95$, $N = 5000$ and $N_T = 3750$ and the inference was terminated after 14 steps.

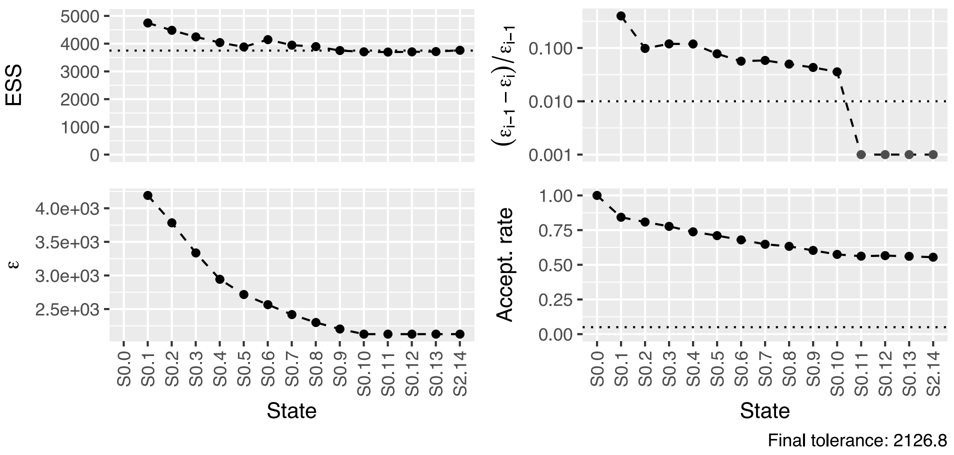


**Figure 6.4:** Example of a ABC-SMC chain. The plot summaries i) the change of the effective sample size (ESS) for the chain with $\alpha = 0.95$, $N = 5000$ and $N_T = 3750$, ii) the resulting schedule of the distance threshold $\varepsilon$, iii) the relative change of the distance $\Delta\varepsilon_{rel}$ and iv) the acceptance rate in the MCMC step for different states of the SMC chain.

#### 6.2.5 Model selection

##### 6.2.5.1 Surrogate likelihood

Generally, the distances under point estimates for neutral simulations were distributed approximately normally, as indicated by quantile-quantile-plots (see Figure 6.5). Further the likelihood to observe a single datum with a distance $\Delta_{D^*}$ below the critical distances $\varepsilon$ was often very low ($\ll 0.1\%$). This makes the rejection based approximation of the likelihood $\tilde{L(\theta)} = p(\rho(\eta(D), \eta(D^*)) \leq \varepsilon)$ computationally very expensive and a synthetic-likelihood approach (Wood 2010) was used instead in cases where the observed fraction of random realisations $D^*$ with distance below $\varepsilon$ was less than 1%. In these cases the likelihood was approximated as

$$\hat{L_s}^N(\theta) = \Phi\left(\frac{\varepsilon - \mu}{\sigma}\right) = \frac{1}{2}\left[1 + erf\left(\frac{\varepsilon - \mu}{\sigma\sqrt{2}}\right)\right]$$

where

$$\hat{\mu}_\theta = \frac{1}{N}\sum_{i=1}^{N} \rho(\eta(D), \eta(D_i^*)),\ \hat{\sigma^2}_\theta = \frac{1}{N-1}\sum_{i=1}^{N}(\rho(\eta(D), \eta(D_i^*)) - \hat{\mu}_\theta)$$

are empirical estimates of the variance $\sigma^2$ and mean $\mu$ calculated from a minimum of $N = 1000$ realisation for each $\theta$.

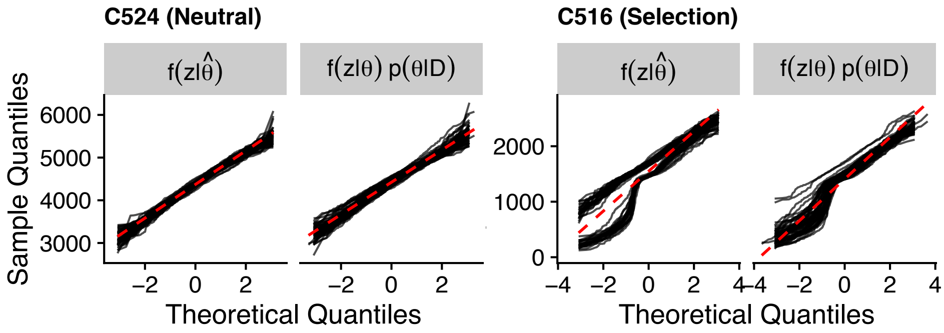


**Figure 6.5:** Quantile–quantile plot demonstrating that the distance between observed and simulated data ($z$) generated under the maximum-a-posteriori ($\hat{\theta}$) as well as the posterior distribution ($p(\theta|D)$) are approximately normal for neutral simulations (left), but not for models with selection (right) .

While it has been shown that the synthetic likelihoods can be fairly robust to violations of normality assumptions (Everitt 2018; Price et al. 2018; Grazian and Fan 2019), improved semi-parametric version have been proposed (An, Nott, and Drovandi 2019). In the context of the ABC-SMC conducted here, distances obtained from neutral models were approximately normally distributed, whereas non-neutral models typically exhibited multiple modes across the observed distance distribution. To these univariate Gaussian mixture models with a variable variance and $K = \arg\min_{K \in 1,...,9} BIC(M_K)$ using the *mclust* R package were fitted instead and estimates of the likelihood were obtained from these (Scrucca et al. 2016; Fraley and Raftery 2002; Fraley et al. 2012).

For each step that involved the estimation of synthetic likelihoods, a simple bootstrap method (Efron 1992) was applied to the observed distances and fitted the Gaussian mixture models to these permutated datasets. This approach allows us to estimate the variability of these estimates and quantify errors arising from the limited Monte-Carlo integration.

###### 6.2.5.2 AIC

With these synthetic likelihoods at hand, the Akaike information criterion (AIC) was used to penalise the different models for the number of free parameters. Here, $\varepsilon = \min_{m \in M} \varepsilon_{end,m}$ was used where $M$ is the set of models on which to perform model selection. Further, a large range of values for $\varepsilon$ was also used to assess the fits. For either of these two options the marginal likelihood $p(D|m,\varepsilon) = \int_\theta p(D|\theta,m,\varepsilon)p(\theta|m)\mathrm{d}\theta$ was estimate for each model $m$ using Monte-Carlo integration across a minimum of 200 parameter sets obtained from the posterior particles weighted by $\{W_n^{(i)}\}$. For $p(D|\theta,m,\varepsilon)$ the SL approximation $\hat{L}^N_{s,\varepsilon}(\theta)$ obtained from the distribution of distances $\Delta(D,D^*)$ of a minimum of 100 simulated datasets $D^* \sim f(\cdot|\theta)$ was used.

We then use the obtained estimate of the marginal likelihood $p(D|m,\varepsilon)$ to calculate $\hat{\mathrm{AIC}}_\varepsilon = 2k - 2lnp(D|m,\varepsilon))$ where $k$ is the number of free model parameters. An example of such estimated marginal likelihoods and AIC values for given values of $\varepsilon$ are shown in Figure 6.8 D&I. Here the negative marginal log-likelihoods (NMLL) of the 'Neutral' and 'Selection' model are very similar, leading to the 'Neutral' mode being preferred over the alternative models due to its lower AIC.

##### 6.2.6 Code availability

The code used to perform the ABC-SMC inference on the EPICC cohort and to create other figures shown here can be found on GitHub: `https://github.com/T-Heide/EPICC_inference`.

#### 6.3 Results

For the ABC-SMC inference, two sets of trees were generally used as input i) the maximum parsimony (MP) trees reconstructed from the mutation data (Figure S.42, page S.42) and ii) the MP trees with assigned LP-WGS samples (Figure 4.11, page 118). The simple accept-reject ABC algorithm was initially applied to a couple of test cases. Two of these — one for which the ABC inferred boundary driven growth and another with non-boundary driven growth — will be shown as a simple example first. Following this, results using the ABC-SMC algorithm from the entire cohort will be summarised.

##### 6.3.1 Rejection ABC

###### 6.3.1.1 Inference of non-boundary driven growth in C561

ABC rejection sampling was, together with a couple of other examples, initially applied to the tree of C561 shown in Figure 6.6B and a total of $1,000,000$ simulated trees from $10,000$ particles were generated. These were filtered to retain $0.05\%$ of simulations with a distance $\varepsilon < 3987.2$. One of the trees from the posterior tree set is shown in Figure 6.6C. This tree and most other trees in the set reflect the general structure of the topology and structure of the target tree, suggesting that the distance method and $\varepsilon$ chosen were able to select trees with a good fit to the target.

Only a single (neutral) model without cell death (i.e., $\mu = 0$) and fixed sampling positions relative to the centre (i.e., $d_b = 25$ and $d_{e,a} = 0.75$) was considered. For this reason, only two parameters, the mutation rate $m$ and pushing distance $d_{push}$ were inferred, with the posterior distribution of both being shown in Figure 6.6A. These suggest that simulations similar to the target tree are more likely to be observed under conditions with weak spatial constrains (i.e., $d_{push} > 20$) at a mutation rate of $m \approx 110$ per gland division. The bottom left and top right grid in Figure 6.6A show the joined posterior probability distribution of $m$ and $d_{push}$, showing only a weak correlation of the two parameters across the posterior distribution.

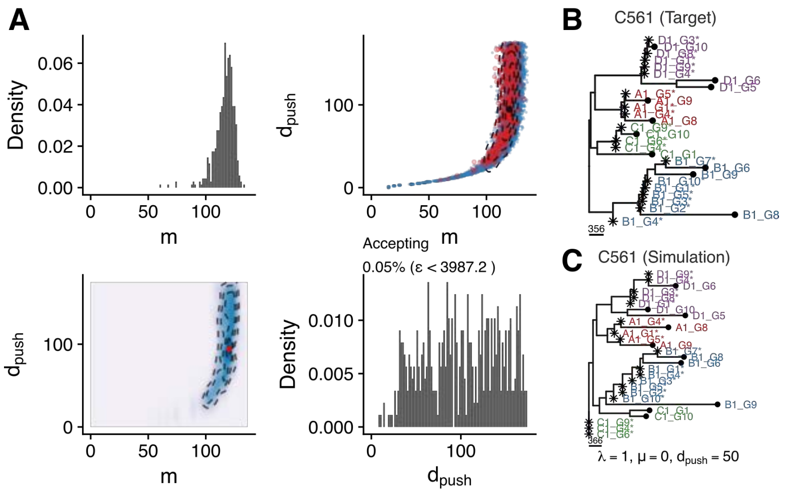


**Figure 6.6:** Results of the ABC rejection algorithm applied to the tree of case C561 (see top of D). A) The estimated marginal and joined posterior density distributions of the $d_{push}$ parameter in the interval $[0,175]$ and corresponding mutations rates, $m$. The marginal densities of the parameters, shown across the diagonal of the plot suggest that $m \approx 110$ and $d_{push} > 20$. A weak correlation of $m$ and $d_{push}$ can be seen in the joined posterior density (bottom left grid). B) Shows the target tree, $y$, and C) a simulated tree, $z$, from the posterior set of trees with $\varepsilon < 39$.

###### 6.3.1.2 Inference of boundary driven growth in case C356

The same ABC rejection sampling was also applied to the tree of case C536 shown in Figure 6.7D. For the distance threshold, the value of $\varepsilon = 760$ — which identical to the final $\varepsilon$ of the ABC-SMC algorithm described below — was used. This allows a direct comparison between the results of the two algorithms. At this $\varepsilon$, all accepted trees were extremely similar to the observed tree. A representative example of a simulated tree from the posterior particle set is shown in Figure 6.7E.

Again only a neutral model with fixed values for $\mu = 0$, $d_b = 25$ and $d_{e,a} = 0.75$ was used, meaning that only two parameters, $m$ and $d_{push}$ were inferred. The posterior distribution of these two parameters is shown in Figure 6.7A. Here the posterior distribution of $d_{push}$ indicates strongly boundary driven growth with $0 \leq d_{push} \leq 5$ and a mutation rate of $m \approx 75$ per gland division.

As seen in the lower right grid of Figure 6.7A the prior distribution for $d_{push}$ was restricted to the interval $[0,25]$ leading to the acceptation of $\approx 0.03\%$ of proposed trees. If proposed particles were instead drawn from the full range of the

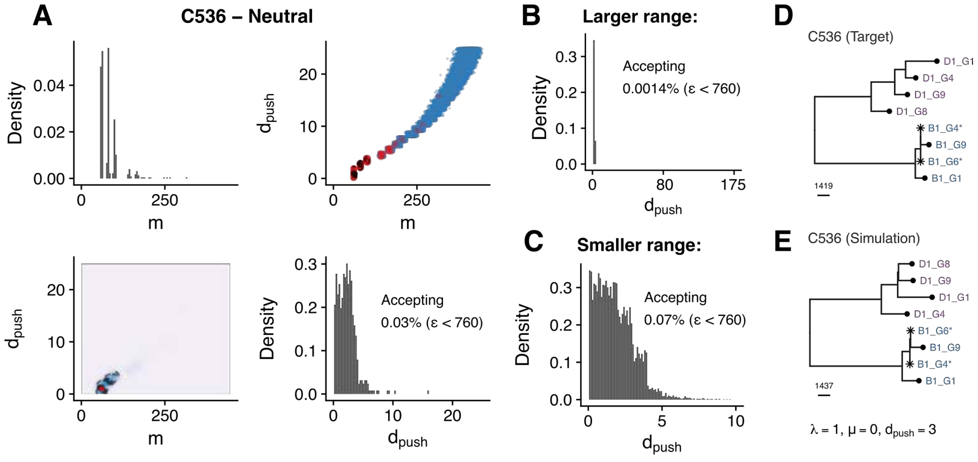


**Figure 6.7:** Results of the ABC rejection algorithm applied to the tree of case C536 (see D). A) The estimated posterior density distribution of the $d_{push}$ parameter in the interval $[0,25]$ and corresponding mutations rates, $m$. B) As A but for the full range of $d_{push}$ ($[0,175]$). C) As A but for a smaller range of $d_{push}$ ($[0,10]$). D) Shows the target tree, $y$. E) Shows one simulated tree, $z$, from the posterior set.

prior (i.e., [0,175]) as shown in Figure 6.7B, the acceptance rate is substantially lower and $\approx 0.0014\%$. Not a single tree with a distance of $\varepsilon < 760$ simulated from particles in the range of $20 < d_{push} < 175$ was observed, hence reducing the total fraction of accepted trees substantially. Likewise a narrower range across the parameter are shown in Figure 6.7C. From the data shown, one can estimate that over the whole distribution of the $d_{push}$ parameter, approximately $1.6 \cdot 10^{-5}$ of proposed simulations are accepted, and to generate 500 accepted trees, one would have to simulate $> 3.2 \cdot 10^7$ trees. In multivariate setups with additional parameters, the number of accepted parameters across the entire parameter space can become computationally infeasible. Even in the above case, the generation of simulations with $d_{push} > 20$ is wasteful, and the low number of accepted simulations would lead to a poor approximation of the posterior. Alternative ABC algorithms — like the ABC-SMC algorithm (Del Moral, Doucet, and Jasra 2012), which will be used below — were developed for this exact reason, and these methods will be applied to the entire cohort instead.

##### 6.3.3 Variable Sampling & Overdispersion

Further evaluation of the posterior trees derived using the fixed sampling layout showed that while the structure of the accepted trees was generally similar to the ones observed, specific aspects of the tree topology and shape were not. For example, compared to internal edges, the length of terminal edges was often consistently too long (see for example Figure 6.9H or 6.9L). I suspected that such differences would be reduced by locating the sampling regions closer to the edge of the simulated tumour, which some simple tests confirmed. Further, questions regarding the effect of the relative sampling position along the edge of the tumour arose, especially whether moving two sampling regions closer to each other would emulate features otherwise attributed to subclonal selection (e.g., clades formed by two adjacent regions).

**Variable sampling model** For this reason, I derived an alternative simulated sampling schema that took the uncertainty of the conducted sampling layout into consideration. The details of this are outlined in Section 6.2.2 above, but in brief, a Dirichlet prior on the angle $\phi$ between sampling regions, a Beta prior on their distance from the edge $d_e$ and a parameter $d_b$ inferred as part of the ABC-SMC, modifying the width of the sampling regions, was added to the model. The average position relative to the edge was assumed to be around 90%, which upon reconsideration of the actual sample collection performed appeared to be more realistic, and the average angular position was again assumed to be at 90, 180, 270 and 360 degree.

In addition to this variation of the sampling positions, I also allowed for increased variability of individual edge length within the tree. This was motivated by the assumption that spatio-temporal variations of the mutation rate or drift within the stem-cell compartment of glands could cause such an overdispersion. Specifically, I replaced the Poisson distributed number of mutations accumulated per generation with a Negative-Binomial distribution. The Negative-Binomial distribution was parametrised so that an additional parameter $\alpha$ would control the amount of overdispersion, which was then inferred as part of the ABC-SMC. For $\alpha = 0$, the distribution of edge length is equivalent to a Poisson distribution, and as $\alpha$ increased, the length of individual edges in the tree gets more varied with the expectation given by the product of mutation rate and the number of generations.

###### 6.3.3.1 Results of model selection

Again the classification and inference framework — this time using the modified version of the simulated sampling — was applied to both WGS and ML LP-WGS trees from the entire cohort. The results of the model selection from this procedure are summarised in Figures 6.15 and 6.16. Summary plots of the results in each case are shown in the Figures S.158-S.186 (page 347-375).

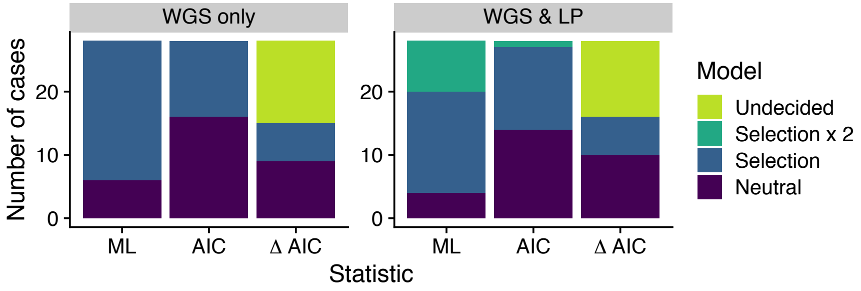


**Figure 6.15:** Summary of model selection results from ABC-SMC with the variable sample setup. Results shown are based on the two tree sets (WGS and LP+WGS), using different model selection criteria. In order of increasing penality these are the marginal likelihood (ML), the AIC and lastly ΔAIC where only models with a minimum difference $\Delta\text{AIC} > 4$ were classified.

When using this alternative setup, a substantially larger fraction of cases were considered to be compatible with the neutral model. Specifically, $16/27$ ($\approx 59\%$) and $15/27$ ($\approx 55\%$) of cases were classified as neutral based on the WGS and ML LP-WGS trees, respectively. A even larger fraction of cases — $10/16$ ($\approx 62\%$) and $13/17$ ($\approx 76\%$) of cases respectively — of those in which the $\Delta\text{AIC} > 4$ suggested sufficient discriminatory power, were classified as neutral.

Comparison of the different classifications of individual cases showed that these were generally consistent with results obtained previously (Figure 6.16). Notably, the six cases[5], which were specifically discussed before (i.e., undecidable with regard to the preferred model), were classified as neutral in this setup. As

[5]C543, C560, C544, C528, C530 and C554

such, no unexpected changes of the general classifications did occur, but the overall structure of the simulated trees was more similar to the observed ones.

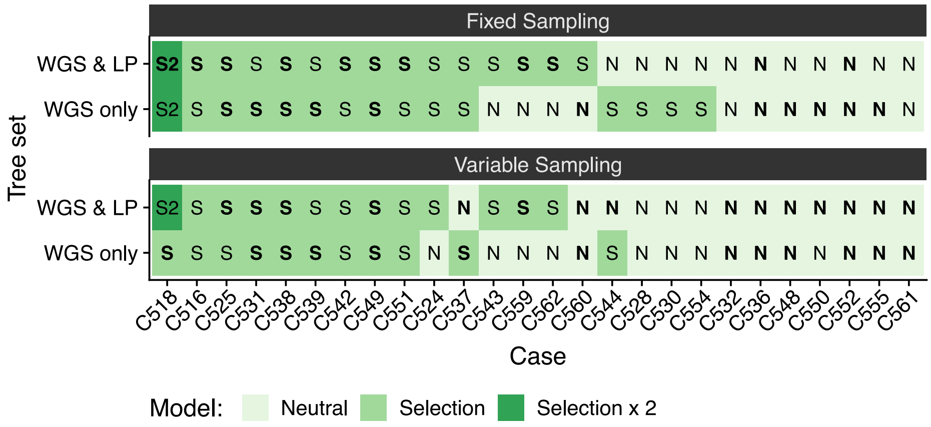


**Figure 6.16:** Agreement of the model selection based on LP+WGS sample and WGS sample trees for both sampling setups. Model sets with ΔAIC > 4 are highlighted by bold text within the tiles.

Finally, I assessed, as before, whether the number of tips differed significantly between cases classified as neutral and non-neutral to elucidate whether the number of tips might have limited the ability to classify cases. As shown in Figure 6.17, this was not the case, indicating that the number of assessed samples did not generally limit the ability to detect selection.

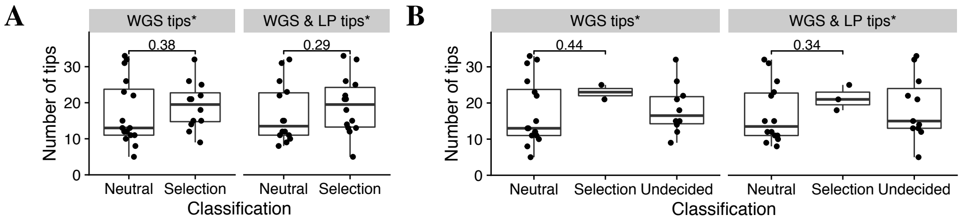


**Figure 6.17:** Relationship of the number of samples on the selected model. A) Shows that for neither of the two datasets a significant difference in the number of available samples existed between the cases classified as neutral ('Neutral') and non-neutral ('Selection') for the inference using the variable sampling scheme. This suggests that insufficient power did not caused the classification of a large number of cases as 'Neutral'. B) The same, but with all samples where discriminatory power was insufficient to choose between two or more alternative model (i.e., ΔAIC ≤ 4) shown as 'Undecided'. Again no difference in the number of samples between cases classified as neutral and non-neutral was evident.
* Cancer samples only.

Summarising the results from the classification obtained using the ABC-SMC inference suggest that the trees observed in $\approx 40\%$ of cases indicated the presence of a selected subclone in at least one of the regions. This is between the estimates of Williams et al. (2016) of $\approx 60\%$ of analysed colon cancers with deviations from the expected VAF-spectrum and the $\approx 20\%$ of analysed cases reported by Williams et al. (2018b). I applied the 1/f test statistic to simulations from the posterior of the ABC-SMC in each case and observed that both cases with boundary driven growth (i.e., $d_{push} < 10$) and subclonal selection, frequently had values of $R^2 < 0.98$ (see Figure S.150A, page 340 and Figure S.151A-B, page 341). As suggested by Wang et al. (2018a), the 1/f statistic used by Williams et al. (2016) might have been more a measure of growth mode than subclonal selection per se. Given the improved power to detect selection events in the single-gland sequencing data of the EPICC cohort, the slightly larger fraction of cases with evidence for the presence of subclonal selection compared to the study by Williams et al. (2018b) appears reasonable.

###### 6.3.3.2 Overdispersion and sampling locations

I next assessed the posterior distribution of the dispersion parameter $\alpha$ and the distribution of the sample locations of the accepted trees. The marginal posterior distribution of $\alpha$ is shown, together with those of the other parameters, in Figure 6.20. As seen here, the posterior distributions of $\alpha$ generally suggested that only a relatively low amount of overdispersion was present in the data (compare Figure S.148 summarising the effect of $\alpha$ on the tree shapes). Notable exceptions from this observation were the cases C516, C549, C538 and C562 (see Figure 6.20, compare Figure S.148, page 338).

In two of these cases, a very small number of samples were obtained (i.e., C516 and C562), and here the larger value of $\alpha$ only improved the fit of the edge to the sample closest to the edge. The remaining two cases with large posterior values of $\alpha$ were still inferred to contain a selected subclone (i.e., not ‘Neutral’), suggesting that some general variability within the tree was insufficiently explained by the introduction of a selected subclone into the model and absorbed by increased variability of the edge length. While the specific reason for this is elusive, it is important to note that the larger degree of dispersion allowed by the model did not cause a different classification of either of these two cases compared to the previous model that did not include such a parameter (see Figure 6.16).

A similar observation was made for the relative positioning of samples within the tumour. Here the variability of $\phi$ and $d_e$ meant that samples could have theoretically been placed much closer to the edge of the tumour or closer to each other. If such changes in the positioning of samples would have consistently lead to improved fits of the model, these patterns should be observable in the marginal distribution of the sample locations within the tumour. In general, little deviation of the sample locations from the prior locations did occur. A representative example of this can be seen in Figure 6.18 (C561).

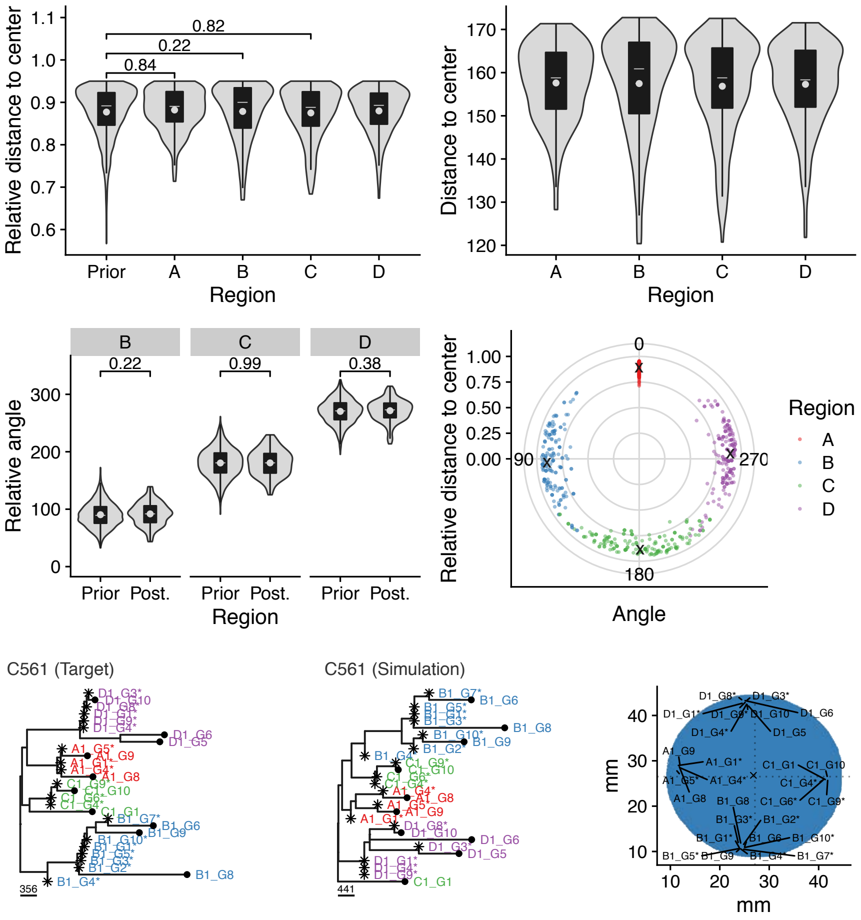


**Figure 6.18:** Regional sample positioning in case C561.

As for the majority of other cases, only limited deviation from the prior did occur in this case, with the mean of the region position being at the average position of the prior. A decent counter-example showing a clear deviation from this pattern can be seen in the fit of the neutral model to the tree of C518 shown in Figure S.149 (page 339). Here, the regions A&B were consistently moved close to each other in space, thus sampling from a clone patch with a more recent MRCA and confirming the initial assumptions. Given the priors defined on the positioning of samples in space, the non-neutral models were still favoured over the neutral model, leading to the selection of the model with two subclones in this specific case.

In summary, these results confirm that while the relative sampling positions were relatively unimportant in many cases, a couple of exceptions did exist. In these, non-neutral were sometimes preferred to explain the observed patterns (e.g., C518), thus leaving the question of whether alternative growth models (e.g., 3D or growth along existing spatial structures) would explain the observed patterns better.

###### 6.3.3.3 Subclonal dN/dS after ABC-SMC classification

While the general consistency of the ABC-SMC across sampling setups and different datasets was on its own reassuring, I next sought to test if an excess of non-synonymous mutations suggested the presence of subclones in those cases classified as neutral. For this, an approach similar to the one used by Tarabichi et al. (2018) in their criticism of the $1/f$ method was followed (Figure 6.19A). In brief, subclonal variants identified in the entire cohort were split based on the microsatellite stability of cases and estimated dN/dS ratios for various set of genes using the *dndscv* package for R (Martincorena et al. 2017). Based on the results of the model selection, cases were divided into those classified as "Neutral" and "Non-Neutral", and the estimation of dN/dS values from subclonal variants detected in these was repeated. The dN/dS ratios estimated in these twelve groups are summarised in Figure 6.19B.

As seen here, subclonal dN/dS estimates of known driver genes were markedly elevated above one (blue arrow in Figure 6.19), indicating the presence of subclonal selection in a subset of cases. Reassuringly, the point estimates of the dN/dS values for which the presence of a subclone was inferred as part of the ABC-SMC inference were even higher, consistent with the presence of a clone with a selec-

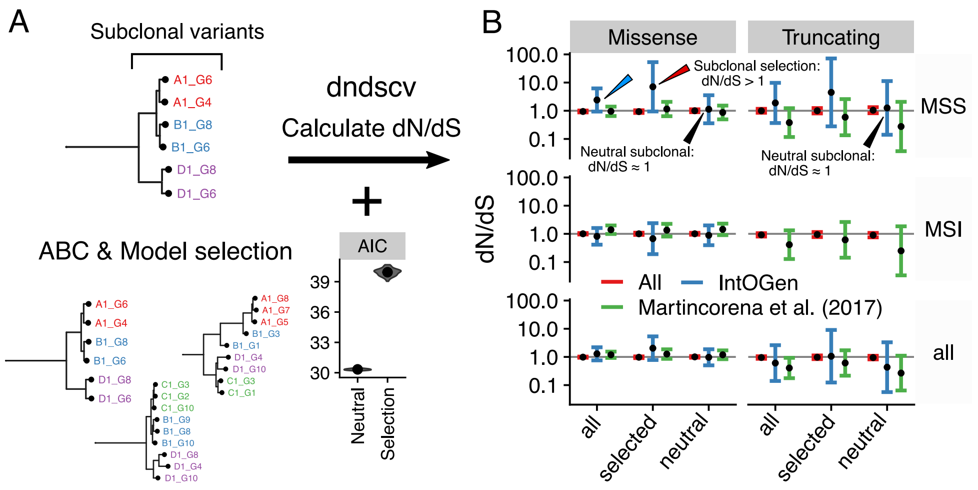


**Figure 6.19:** dN/dS estimates from subclonal variants based on ABC-SMC classification. A) The set of all subclonal variants was split based on the model selection performed by the by the ABC-SMC inference method.B) Shows dN/dS estimates by *dndscv* (Martincorena et al. 2017) from subclonal variants of all, MSI and MSS colorectal carcinomas (y-axis grids) for missense and truncating variants (x-axis grids) in the entire genome ("All"), colorectal driver genes defined by IntOGen (Martínez-Jiménez et al. 2020) and a set of pan-cancer drivers from Martincorena et al. (2017). The results show a general excess of subclonal non-synonymous variants in driver genes for MSS cases (blue arrow). After classification of cases with the ABC-SMC inference method, evidence of positive selection was found in those classified "selected" (red arrow), but not not in those classified as "neutral" (black arrows).

tive advantage due to a somatic mutation (red arrow in Figure 6.19). In contrast the dN/dS point estimates for subclonal variant of the remaining cases (i.e., those classified as neutral), was almost exactly $dN/dS = 1$ (black arrow in Figure 6.19), consistent with neutral evolution in the observed parts of the tumours.

Altogether, this analysis suggests that the ABC-SMC classification was indeed able to discriminate trees for which their structure suggested the presence of subclonal selection in some parts of the tumour. While potentially underpowered due to the relatively small number of analysed cases (27), the majority of data (i.e., $\approx 60\%$) appeared to be better explained by the very simple spatial model without subclonal selection. Consistent with this, the orthogonal dN/dS analysis also did not reveal any excess of non-synonymous mutations. While it is certainly possible that non-genetic drivers could cause widespread subclonal selection, the somewhat limited analysis of chromatin accessibility changes and differential expression[6] conducted as part of the EPICC study did not reveal any evidence for this. Again, alternative models including more complex spatial dynamics, like immunoselection, necrosis or structures of the surrounding environment, should certainly be considered to assess whether these would provide a better fit to the data.

###### 6.3.3.4 Posterior distributions

I next evaluated the posterior distribution of the individual parameters. For this reason, the marginal posterior distribution of all parameters was determined. These are summarised for both datasets and all cases in Figure 6.20. In the following, specific aspects of the marginal posterior distributions will be explained.

**Boundary vs non-boundary driven growth** Consistent with previous results, $5/6$ neutral cases[7] for which the inference previously suggested the presence of boundary driven growth (i.e., $d_{push} \leq 20$). Such a pattern was also observed in the posterior of the inference using the alternative sampling setup (see panels $d_{push}$). Similarly, neutral cases for which the inference previously suggested non-boundary driven growth were generally also so with the new results in $7/8$ cases[8] (87.5%). In the remaining cases, wide posterior intervals suggested that little information on the strength of boundary driven growth was contained in the data. Still, the measurements of such growth properties are rarely possible (Gerlee 2013), and as such, the inference of the strength of boundary driven growth in a subset of cases is interesting in itself.

**Mutation rates** Mutations are expected to accumulate faster at sites with many copy-numbers. For this reason, mutation rates obtained from the posterior distribution of the $m$ parameter, mutation rates were adjusted to account for the differences in the ploidy of individual cases. After this correction, the median of the estimated mutations rates (MAP) in the tumours was 78.6 and 540 per gland division per diploid genome for MSS and MSI cases, respectively. Assuming an effective genome size of $2.9 \cdot 10^9$ base pairs, this corresponds to a mutation rate of $1.35 \cdot 10^{-8}$

[6] This work was done by Jacob Househam.
[7] C536, C532, C544, C552, C554. Not C548.
[8] C522, C528, C530, C550, C555, C560 and C561. Not C548.

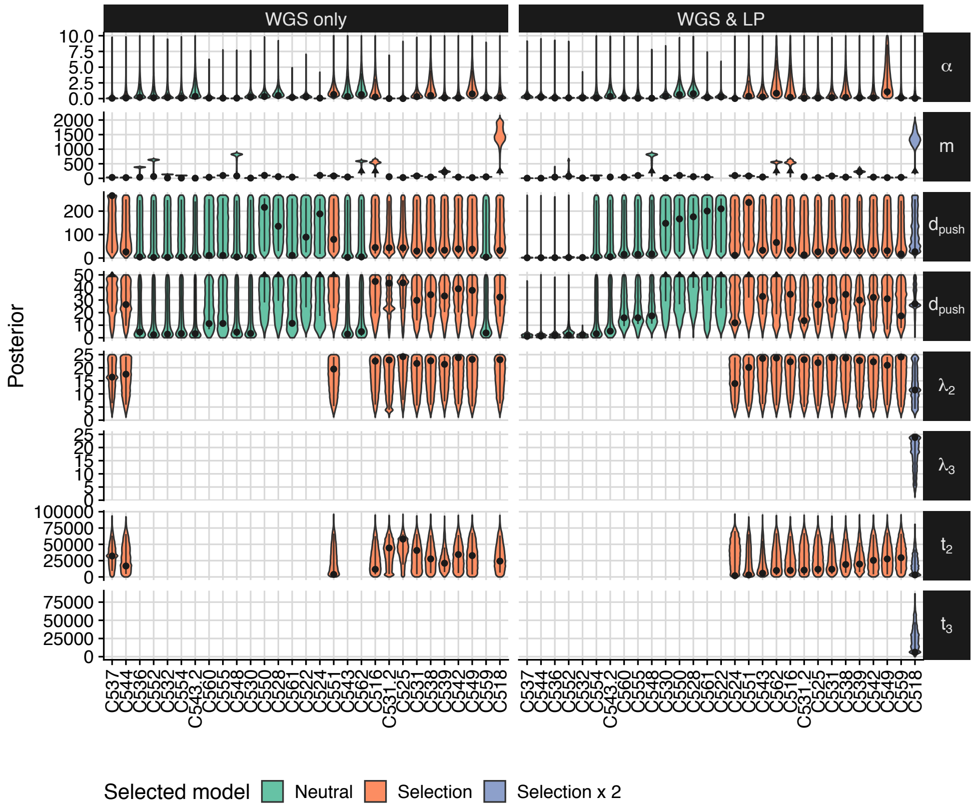


**Figure 6.20:** Marginal posterior distribution of inferred parameters obtained from the ABC-SMC using the variable sampling setup. For the parameter estimates of the width of the growing outer rim $d_{push}$ a smaller window $(0,50)$ of the posterior is shown (fourth row). Mutation rates $m$ with a value over 250 (i.e., MSI cases) are truncated (indicated by a small triangle). Abbreviations: $\alpha$ dispersion parameter, $m$ mutation rate, $d_{push}$ width of growing edge, $\lambda_i$ birthrate of subclone $i$ and $t_i$ start time of subclone $i$. The lines indicate the central 90% intervals of the marginal posterior distribution, dots the multivariate estimates of the maximum a posteriori probability.

and $9.31 \cdot 10^{-8}$ per base and gland division, respectively. The relative difference of a $\approx 6.8$ fold difference between MSI and MSS cases is generally consistent with those reported previously. Still, the rates themselves are lower then those estimated by Williams et al. (2016) and Williams et al. (2018b) from bulk WGS sequencing data.

Previous studies have suggested that boundary driven growth, which is not part of the model used in either of the two studies, would lead to the overestimation of mutation rates using the statistics (Fusco et al. 2016; Schreck et al. 2019). For this reason, I tested the performance of the 1/f test statistic on these cases and found that in those with boundary driven growth, estimates of the mutation rates were indeed larger than the true rates (Figure S.151C and S.150B). Still, as the majority of cases with strong boundary driven growth had a $R^2 < 0.98$ the majority of these would have been expected to be excluded from the analysis performed in Williams et al. (2016) (see Figure S.151A, S.151B and S.150A). I also assessed whether a high false-negative rate during the calling of mutations in single-gland samples with Mutect2 might explain this observed discrepancy (see Figure S.156, page 344), but found that the power to identify clonal mutations in individual samples was generally above $> 90\%$ (i.e., FNR $< 0.1$). It might still be possible that a significant amount of false-positive variant calls, which would be an issue expected to less problematic in the single-gland WGS data analysed here (Salcedo et al. 2020).

**Subclone parameters** Consistent with the previous observations, the posteriors of the clone specific parameters $\lambda_i$ and $t_i$ were very widely distributed. Especially, the birthrate varied over a large range of values. The likely reasons for this were described in detail above, but in brief, very little information on the size of individual clones is contained in the generated data. As this is the main measure that allows estimating $\lambda$ and $t_i$, large posterior intervals are expected. For more precise estimates, a different sampling layout would have to be used.

**Conclusion** Assuming that the inferred estimates of $d_{push}$ are informative of this property of a tumour, one would assume that the absence of spatial limitations on the tumour growth might be a property of generally more invasive phenotype and hence point to a worse prognosis. Likewise, the presence of a faster-growing subclone might be indicative of a worse outcome as well. As the entire study was planned as prospective study data on the outcome will be available in the future.

###### 6.3.3.5 Tree topologies

In Figure 6.21 critical parameters of the ABC-SMC inference, as well as the classification of cases along the trees used for the inference, are shown alongside each other. Edges of the tree that were consistently equivalent to the simulated ‘Driver’ mutation in the 200 best fitting simulated trees are highlighted in colour for the cases classified as non-neutral. In cases where somatic subclonal drivers were identified as part of the previous analysis, these were also added as labels to the tree.

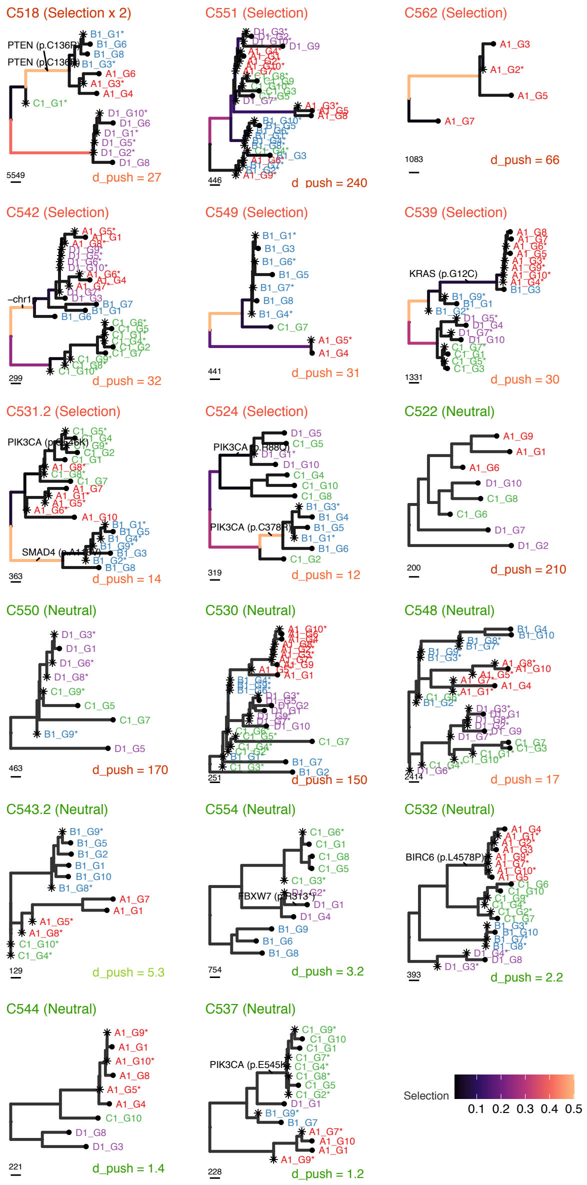


**Figure 6.21:** ML LP-WGS trees and associated results of ABC-SMC inference. The classification of cases based on the AIC (i.e., 'Neutral', 'Selection' and 'Selection x2') are shown above trees. The MAP estimates of the strength of boundary driven growth for the selected model are shown below the trees. The frequency colours edges of the trees that these were associated with the introduced subclonal 'Driver' mutation in the 200 best fitting trees from the posterior distribution (legend in the bottom left).

In a number of these highlighted, selected edges, a subclonal driver mutation was identified. These observations suggest that the subclonal mutations in Table 6.2

had detectable fitness effects in the respective genetic and environmental context in which they occurred. The following paragraphs will discuss each of these putative driver mutations in detail based on the existing literature. In the majority of cases, the identified subclonal driver mutations did indeed provide a reasonable explanation for the observations, thus demonstrating that the inference framework was able to recover a relevant signal from the reconstructed phylogenetic trees.

**Table 6.2:** Likely subclonal driver mutations identified by the ABC-SMC inference on the trees of the EPICC cohort.

| Case | Gene | Mutation | Type | Figure |
|---|---|---|---|---|
| C518 | PTEN | p.C136R | Second hit of TSG | S.134, page 329 |
| C524 | PIK3CA | p.C378R | Oncogene | S.135, page 330 |
| C525 | PIK3CA | p.Q546P | Oncogene | S.136, page 330 |
| C531 | SMAD4 | p.A118V | TSG | S.137, page 331 |
| C538 | RNF43 | p.D153* | TSG | S.138, page 331 |
| C539 | KRAS | p.G12C | Oncogene | S.139, page 332 |
| C542 | chr1p | Loss | CNA | S.101, page 308 |

**Activating KRAS mutation** Activating mutations in classic colorectal oncogenes, like KRAS and PIK3CA, both of which lead to the over-activation of signalling pathways — the RAS and the PI3K/AKT pathway, respectively — generally have a dominant effect. As such, the observation of a fitness altering effect for these is not surprising per se. Especially for the KRAS p.G12C mutation observed in C539, these would certainly be expected.

Alterations of RAS genes are the single most common oncogenic mutation acquired in various human malignancies (Bos 1989). While in colorectal cancers typically p.G12D, p.G12V and p.G13D mutations are found to be present, a smaller subset of these ($\approx 3\%$) harbour p.G12C mutations (Prior, Lewis, and Mattos 2012). The most likely explanation for these tissue-specific differences is that the specific substitutions with which these are associated — a C[C>A]A mutation for the p.G12C KRAS mutation — are more likely to occur under specific mutagenic processes active in these tumour entities (Prior, Lewis, and Mattos 2012; Temko et al. 2018). In the case of C539, the analysis of somatic signatures indicated the presence of a relatively large number of C>A mutations in this case (Figure S.152, page 342), thus providing a reasonable explanation for the presence of this unusual KRAS mutation.

Furthermore, the KRAS mutation arose in the context of a full loss of APC and TP53 activity, thus providing a nice counter-example to the classic adenoma-carcinoma sequence, which suggests that oncogenic KRAS mutations are important for the initiation of adenomas and typically occur before TP53 mutations (Vogelstein et al. 2013). The analysis shows that activating subclonal KRAS mutations lead to substantial fitness effects in an already established colorectal carcinoma.

**PIK3CA mutations** Interestingly, neither of the two PIK3CA mutation for which a significant fitness effect was inferred (i.e., p.C378R and p.Q546P) was one of the most common hotspot mutations (i.e., p.E542K, p.E545K or H1047R). Nevertheless, previous screening of the phenotypic effects of such rarer PIK3CA mutations showed, these can also have growth-promoting activity (Dogruluk et al. 2015) and the obtained data suggest that in the genetic background of KRAS mutations, these two variants cause significant phenotypic effects *in vivo*. Interestingly, a recent paper found co-mutation of KRAS and PIK3CA to be associated with poor overall survival (Luo et al. 2020)

Still, while the lack of phenotypic effects might explain the absence of selection for some of the other rare PIK3CA variants (i.e., p.R88Q and p.G118D), the p.Q546K observed in C531 was previously found to have growth-promoting effects, specifically for one a very similar to the p.Q546P variant identified in C525. While it is certainly possible that the environmental or genetic background of this specific case played a role, it seems to be more likely that the variant arose very recently within the carcinoma and that selection did not have sufficient time to act on this variant. In this case, it would be expected that only a little increase of the edge length would occur, causing problems to identify deviations from the neutral expectation.

A similar lack of power to detect selection might explain why the p.H1047Q and p.545K found in C544 and C537, respectively, were not identified. Alternatively, the absence of co-occurring KRAS mutations might reduce the effect of these

PI3KCA mutations, which would be in line with observations made by others (e.g., Wang et al. 2013; Stintzing and Lenz 2013; Phipps, Makar, and Newcomb 2013; Green, Trejo, and McMahon 2015; Oda et al. 2008; Wang et al. 2018b).

**PTEN mutation in C518** In the case C518, the inference suggested that the presence of a PTEN p.C136R might have caused the significant selection of the mutant subclone. Importantly, this subclonal PTEN mutation occurred in the background of a clonal truncating PTEN mutation p.K267Rfs*9) that likely caused the loss of function of the alternate allele. A separate analysis of the RNA-seq data[9] from the EPICC cohort confirmed that the expression of the alternate allele was completely lost in samples from region A and B of the tumour (see Figure S.153, page 343).

The subclonal variant identified leads to a substitution very close to the $NG_2$-terminal phosphatase domain of the PTEN protein (Han et al. 2000), but not one of the frequently mutated hotspots themself (Bonneau and Longy 2000; Dillon and Miller 2014). Similar PTEN mutations (i.e., p.C136Y) have been found to disrupt the phosphatase activity of the PTEN protein itself (Han et al. 2000) and others have reported association of germline p.C136R mutations with Cowden syndrome — a genetic disease caused by germline PTEN mutations — as well as reduced stability and loss of phosphatase activity of the encoded protein (He et al. 2013). Taken together, this demonstrates that the subclonal variant likely lead to the complete loss of the remaining PTEN activity.

PTEN itself is considered a CRC tumour suppressor gene, which inhibits the conversion of phosphatidylinositol (PI)-4,5-bisphosphate to PI 3,4,5-triphosphate (PIP3) (Molinari and Frattini 2014). Loss of PTEN activity, in turn, causes the accumulation of PIP3 within the cells, leading to the over-activation of downstream PI3K pathway (Song, Salmena, and Pandolfi 2012). This activation of the PI3K pathway, mediated through its various targets, is associated with increased proliferation, inhibition of cell death and stimulation of angiogenesis (Song, Salmena, and Pandolfi 2012; Molinari and Frattini 2014). Summarised, the subclonal PTEN mutation identified in C518 provides another example of subclonal alterations of the

[9]This work was done by Jacob Househam.

PI3K pathway in the analysed CRC.

**SMAD4 mutation in C531** The ABC-SMC inference applied to the tree of C531 suggested the presence of a putatively selected subclone in region B of the tumour. The assessment of potential somatic driver alterations in genes reported in the IntOGen database revealed the presence of s single SMAD4 mutation.

Mutations in SMAD4 typically occur in about 8.5% of CRC (Fleming et al. 2013) and three clonal SMAD4 variants in MSI CRC and the aforementioned subclonal SMAD4 p.A118V mutation in C531 were identified. SMAD4 mutations have been associated with poor outcome in a metastatic setting (Alazzouzi et al. 2005; Mizuno et al. 2018; Kawaguchi et al. 2019) as well as chemo-resistance to 5-fluorouracil (Zhang et al. 2014a) and EMT mediated resistance to Cetuximab (Lin et al. 2019), highlighting the importance of SMAD4 loss in late-stage disease of CRC. The genes of the SMAD are a family of transcription factors that control the expression of genes regulated by the TGF-$\beta$ signalling pathway. Activation of TFG-$\beta$ signalling has both tumour suppressive as well as invasion and metastasis promoting effects (Bierie and Moses 2006). SMAD2-4 complexes are necessary for the transduction of growth-inhibiting effects of TGF-$\beta$ signalling in the nucleus, and their loss appears to cause a shift towards negative effects of TGF-$\beta$ signalling in CRC (Zhang et al. 2010).

The specific subclonal driver mutation observed (i.e., p.A118V) in C531, occurred in a frequently mutated position of the gene and that is likely pathogenic (Iacobuzio-Donahue et al. 2004; Jones et al. 2008; Fleming et al. 2013). Still, the mutation only affected one of the two SMAD4 alleles (see Figure S.137, page 331). Some previous studies indicate that mutations of genes of the SMAD family can act dominantly-negative (Hoodless et al. 1999; Xu et al. 2000; Alberici et al. 2006), but bi-allelic mutation of SMAD4 are frequently observed in cancer genomic data (Fleming et al. 2012), suggesting that the mutation of the second allele is required. For this reason, I next assessed the expression of SMAD4 in general and the p.A118V variant specifically in all the samples of C531. Surprisingly, this analysis of the RNA-seq of the EPICC project showed that virtually no expression of SMAD4 occurred in samples from the mutated region B (see Figure S.154, page 343). This complete loss of SMAD4 expression — potentially due to the down-regulation of expression due to a second independent event — as well as the pathogenic SMAD4 mutation, suggest that a complete loss of SMAD4 activity occurred in the corresponding lineage of the tumour. In summary, this makes the SMAD4 mutation in combination with a potentially independent loss of its expression the most likely explanation for the observed subclonal selection of region B in this case.

**RNF43 mutation in C538** Another example of a subclonal mutation of a known CRC driver gene on an edge for which the ABC-SMC suggested the presence of a positively selected alteration was identified in C538. Here all samples found in region B and one sample from region A (A1_G7) were part of a putatively selected subclone, and an associated truncating mutation of RNF43 p.Q153* was identified. Truncating mutations of RNF43 are relatively frequently observed in colorectal, endometrial, ovarian and pancreatic carcinoma (Jiang et al. 2013; Zou et al. 2013; Ryland et al. 2013; Giannakis et al. 2014).

RNF43, together with ZNRF3, typically plays a vital role in the inhibition of Wnt signalling through the degradation of Wnt receptors of the Frizzled (FZD) family (Koo et al. 2012; Jiang et al. 2015; Tsukiyama et al. 2015). Since the expression of RNF43 and ZNRF3 is induced by Wnt/β-catenin signalling itself, these protein provide a negative feedback loop required for this signalling cascade. The deletion of both RNF43 and ZNRF3 was shown experimentally to induce tumour formation in mice (Koo et al. 2012) in a similar fashion to APC loss. Indeed, APC and RNF43 mutations were found to be mutually exclusive in CRC (Giannakis et al. 2014), which is consistent with their activation of the Wnt pathway by different means. The degradation of FZD receptors by RNF43/ZNRF3 is thought to be caused by the activation of endocytosis through ubiquitination by the RING domain of these proteins (Hao et al. 2012; Koo et al. 2012) and the recognition of FZD receptors is mediated by DVL binding to the DIR domain of RNF43 and ZNRF3 (Jiang et al. 2015).

The variant observed in C531 is predicted to cause the loss of both of these domains (i.e., DIR and RING) as well as the transmembrane domain and various other parts of the protein. While such a mutation certainly leads to the loss of function of the mutated allele, only one of two alleles was found to be mutated in C531 (Figure S.138, page 331), meaning that the other allele could potentially compensate for the loss of RNF43 function. Similarly, it had been suggested previously that ZNRF3 can potentially compensate for the loss of RNF43 and that only loss of both proteins leads to tumour formation and altered Wnt/β-catenin signalling (Koo et al. 2012; Lannagan et al. 2019). Assessment of the mutation and expression status of ZNRF43 revealed no evidence for somatic mutations or altered expression of ZNFRF43 in the corresponding samples. The expression of the mutant RNF43 variant showed that the gene was dominantly expressed in one of the three samples from tumour region B but not in the others.

Interestingly, some previous studies suggested that missense mutations observed in CRC (Koo et al. 2012; Tsukiyama et al. 2015; Yu et al. 2020) and truncated RNF43 variants missing the RING domain (Hao et al. 2012; Tsukiyama et al. 2015) might have a dominant-negative (DN) effect. Still, the overwhelming majority of truncating variants tested in a large screening of RNF43 variants by Yu et al. (2020) and specifically a p.Q152* mutation, i.e., one AA difference to the one observed here, were found to only lead to loss of function and not to be dominant-negative. A single truncating frameshift variant of APC was observed (p.E1309Dfs*4) in this patient. Assuming that a second APC variant affecting the alternate allele was present but not detected, a mutation like that of RNF43, which also activates the Wnt/β-catenin pathway, would be expected to not lead to any or very little fitness increase.

Evidently, the results suggest that the mono-allelic LOF mutation of RNF43 p.Q153* in C538 might have a fitness altering effect, but this appears to be at odds with existing literature and the assumed mode of function of this mutation. Given the APC mutated background and the inconclusive analysis of the gene expression data, it might be possible that a different, possibly undetected, somatic mutation or non-genetic alteration was responsible for the observed effect. Alternatively, while certainly speculative, mono-allelic RNF43 mutations in the APC depleted background present in C538 might be haploinsufficient and hence lead to further activation of the Wnt/β-catenin pathway with corresponding fitness effects.

**Other cases** Similarly, non-genetic or unidentified somatic mutations of driver genes, might provide an explanation for four cases in which no corresponding somatic mutation in a driver gene could be identified. These were C542 (region B&C, Figure S.141, page 332), C542 (region A, B & D, Figure S.141, page 332), C549 (region B&C, Figure S.143, page 333), C551 (Figure S.144, page 334) and C559 (region B, Figure S.145, page 334). In C542 for example, the subclonal loss of chr1p — an alteration that was found to be recurrent in Stage III CRC (Xia et al. 2020) and more frequently mutated in metastatic CRC (Ghadimi et al. 2006; González-González et al. 2014) — could provide an explanation for the subclonal selection of one tumour region. In general, a larger cohort of cases would be needed to perform an assessment of whether such cases in which no subclonal driver mutations could be identified are simply false-positives or if identifiable rare or non-genetic drivers are indeed present in these.

I found no conclusive evidence of subclonal alterations of gene expression using the available RNA-seq data or changes of chromatin accessibility that could explain the observed difference, but future studies powered to perform a comprehensive analysis of such alterations in putatively selected subclones should be considered.

###### 6.3.3.6 Prediction of clone-size distributions

The marginal posterior of the ABC-SMC inference can also be used to estimate the distribution of subclones within the tumour. For this, the 100 simulations that produced trees with the closest distance to the observed data were obtained and used to estimate the marginal distribution of each subclone in space. The location of simulated cells was first rotated by $-\phi_A$, assign each to the closet grid point and then used to calculate the fraction of times specific subclone were observed at these positions (see Figure 6.22A). The results of this procedure are shown for one example (C539) in Figure 6.22B. Here the inference suggested that the subclone (Clone 2) extended over a large area of space.

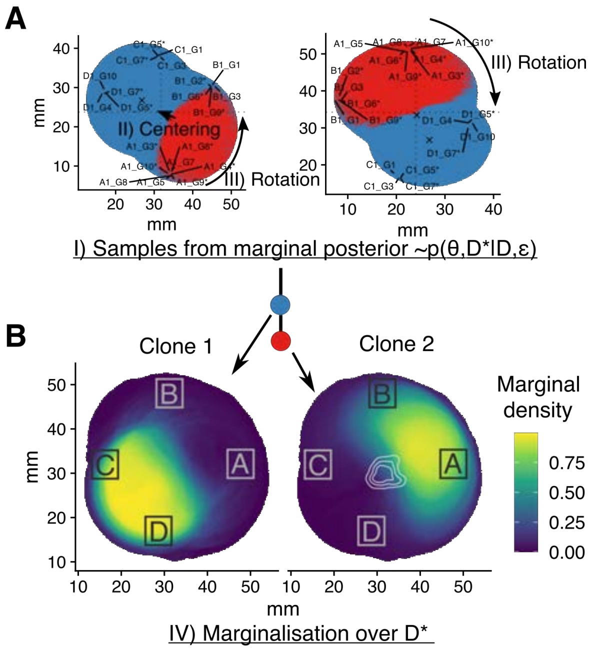


**Figure 6.22:** Marginal clone size estimate for C539. A) Shows two realisations drawn from the set of particles approximating $p(\theta, D^*|D, \varepsilon)$. These are centred and rotate to calculate B) The marginal fraction each grid point contains a specific clone.

Various method to detect and map somatic variants *in situ* have been developed (Bagasra 2007; Larsson et al. 2004). One example of this is the BaseScope assay (Baker et al. 2017), which allows the detection of mutations using RNA *in situ* hybridisation techniques. This method is more robust than similar *in situ* mutation detection methods and can be performed on permeabilised formalin-fixed paraffin-embedded tissue. Such tissue blocks were available as part of this study. In order to validate and potentially refine the prediction of the ABC-SMC on the spatial distribution of individual subclones (e.g., Figure 6.22B), cases with subclonal mutations detectable by the BaseScope assay were determined. A total of two such cases were identified i) the subclonal, selected KRAS p.G12C mutation found in C539 and ii) a subclonal PIK3CA p.E545K mutation in C537, that, at least according to the inference on ML LP-WGS trees, was not positively selected. The basescope assay will be conducted on these two cases to validate the results described here.

##### 6.3.4 Limitations of the current analysis

While the performed inference appeared to generally provide sensible results and able to recover subclonal selection due to somatic mutations of bona fide CRC drivers — examples of this are the activating KRAS mutation in C539 or the PIK3CA mutation in C525 — a couple of limitations of the analysis, which will be discussed in the following, have to be considered.

**Limits of the used sampling schema** First of all, the applied sampling schema provides on its own a minimal amount of information on the spatial extent of selected subclones within the tumours. If, for example, all samples from one region of the tumour would be sampled from a subclone — lets assume that this could be identified by an elongated edge in the reconstructed tree in this case — only very little would be known about the size of the clone itself. From the observed data, it would be reasonable to assume that the subclone covers at least the sampled region (i.e., $d \approx 5mm$) and less than a quarter of the tumour (i.e., from the left to the right neighbouring region).

Since the size of a subclone and the time it arose provide information on the relative strength of selection a subclone is experiencing, large credibility intervals (CI) on $\lambda$ would be expected. These large CI are precisely what was observed from the inference, with potential values of $\lambda$ ranging from little more than the background clone to a, certainly unrealistic, 25-fold increases of the division rate of selected subclones (see Figure 6.20).

Performing systematic spatial sampling while keeping a record of the relative sampling positions in space or deep bulk sequencing of a huge tissue sample would provide much more accurate information on the spatial extend of selected subclones. Other, more practical methods might also allow a better idea of the spatial extent of these. For example, mixing the spatial sampling performed here with several random small punch biopsies along the outer diameter and subjecting these to pooled deep sequencing might also allow a more accurate estimation of relative clone sizes in combination with the spatial model.

Similarly, large parts of each tumour were not observed, potentially reducing the ability to detect expanding subclones in general. The sampling locations themselves only cover a tiny area of the tumour as a whole (i.e., $< 5\%$), and while selected subclones by definition grow to cover an area larger than expected, they could, in principle, still be present between sampled regions. This assumption is inherently encoded in the model itself and hence the inference framework, but a detailed analysis of the likelihood of this under different model parameters might be worth conducting.

**Three-dimensional structures** Furthermore, the three-dimensional structure CRC exhibit was simplified (see Figure 6.2A) into a two-dimensional model. This might be reasonable if cells primarily grew along the horizontal plane, but adenoma and carcinoma frequently exhibit complex three-dimensional structures growing into the colorectal lumen. Whether subclonal dynamics within these are sufficiently recapitulated in the simpler two-dimensional model used here could undoubtedly be questioned. Especially as some conclusions made here are on the strength of boundary driven growth, questions remain if and how these would be recapitulated in three dimensions.

Maybe, alternative growth models should also be considered. The spatial model used here is equivalent to the 'constant crust' model of tumour growth (Mayneord 1932; Conger and Ziskin 1983). Such simple models were suggested to be insufficient to even explain growth dynamics of tumour spheroids in culture (Marušić et al. 1994) or *in vivo* (Marušic et al. 1994). Despite a long-held interest in the growth laws of human malignancies (Steel and Lamerton 1966; Gerlee 2013), the exact nature of these in tumour entities in general (Chignola and Foroni 2005; Talkington and Durrett 2015) and CRC in specific (Burke et al. 2020) are, surprisingly, not fully understood. It would certainly be of interest if genomic measurements could give insight into the growth law (i.e., exponential vs boundary driven growth) on a patient-by-patient basis, especially as these might even predict the growth rate at related metastatic sites, but various other models should be considered as well. For example, desquamation at the surface of individual CRC — which was proposed by Spratt (1961) to explain the slower growth rates of CRC at the primary site compared to the metastatic site — or the inclusion of central growth inhibition (Mayneord 1932; Steel and Lamerton 1966) might significantly alter the predictions made by inference on the genomic data.

**Stem cell dynamics** Another factor that should probably be considered are the stem cell dynamics within single-glands and the heterogeneity of cancer stem cells (CSC) in these. These additional complexities were ignored here, which is a reasonable choice if the replacement rate of stem cells within glands/crypts is swift compared to the division rate of these. Previous studies have shown that this is to be accurate, but this might not generally be the case.

Siegmund et al. (2009b) for example, have suggested that 'palm shaped' phylogenies, which were found to be explained reasonably well by boundary driven growth alone, might arise due to the presence of a few long-lived CSC and that star-shaped phylogenies, here explained by non-boundary driven growth, instead arise in the presence of many CRC.

Not having included these dynamics into the spatial model appears to be one of the major shortcomings of the analysis conducted here. It is unclear whether large stem cell pools and growth dynamics could be distinguished from each other and how data generated under a combination of both of these would behave.

**Changes of mutation rates** Last but not least, spatio-temporal changes of mutation rates might occur during the evolution of individual tumours. The previously described analysis of mutational signatures, as a surrogate of such mutation rates, did not identify any particularly prevalent subclonal changes. While the analysis of mutation signatures did suggest that the relative contribution of the individual process was not altered over time, it could certainly be that the absolute contribution changes over time or that the analysis was simply unable to resolve variations of the different processes sufficiently. Given that one of the signatures of subclonal selection is expected to be the elongation of individual edges compared to the remaining tumour, it might be hard to resolve these two hypotheses.

#### 6.4 Discussion

Here I presented results from statistical inference, which used an extended version of a spatial tumour model previously described by us (Chkhaidze et al. 2019) to perform computational inference on single-gland multi-region sequencing data from a total of 26 CRC. Using this framework, it was possible to quantify tumour-specific properties that describe their evolutionary dynamics. Specifically, cases were distinguishable based on i) the presence of subclonal selection, ii)their growth law (i.e., slow, boundary driven vs fast exponential growth), and iii) their mutation rates. Importantly, these are — unlike a mere collection of features provided by the measurement of gene expression (Uhlen et al. 2017) or somatic mutations (Campbell et al. 2020) — interpretable descriptions of the growth dynamics occurring in individual tumours.

While speculative, these properties might also predict the speed with which already exiting, but still, invisible metastasis grows or be associated with a generally malignant phenotype. As such, the corresponding parameter estimates appear to be a reasonable proposal for an ‘evolutionary biomarker’. The quality of these will be assessed as part of the prospective follow-up of the EPICC study in the near future.

A specific aspect of the inference that appears to be worthwhile to validate further is the prediction that some CRC evolve under boundary driven growth, whereas others did not. This would imply that individual CRCs are growing at a substantially different speed. Measuring such ‘growth laws of CRC in general and especially in individual patients appears to have been challenging (Burke et al. 2020), but of general importance (Friberg and Mattson 1997; Sachs, Hlatky, and Hahnfeldt 2001; Comen, Morris, and Norton 2012; Rodriguez-Brenes, Komarova, and Wodarz 2013).

The presented results also corroborate previous studies on the prevalence of subclonal selection in CRC. In many of the characterised tumour lineages, I was unable to identify any putative driver alterations. These cases were further sufficiently explained by a spatial model without selected subclones, suggesting that the observed parts of the tumours evolved effectively neutral. This is consistent with previous observations made in bulk sequencing data (e.g., Williams et al. 2016; Williams et al. 2018b) and supports the reply to previous criticism (e.g., Williams et al. 2017; Williams et al. 2018a; Heide et al. 2018) of these studies by others (e.g., Balaparya and De 2018; Tarabichi et al. 2018; McDonald, Chakrabarti, and Michor 2018). In a subset of cases, subclonal bona fide driver mutations were identified. Here the ABC-SMC inference framework frequently suggested the presence of associated positively selected subclones. This demonstrates that the method was sufficiently powered to detect deviations from neutrality in general. An orthogonal analysis of dN/dS values, inspired by the criticism of Tarabichi et al. (2018), support these conclusions.

The aforementioned raises questions regarding the interpretation of subclonal tumour evolution from bulk WGS sequencing data done by Dentro et al. (2021). Here the authors found evidence of pervasive subclonal selection,
in colorectal cancers and frequently in the absence of associated subclonal driver mutations. As such, the observations by Dentro et al. (2021) appear at odds with the higher resolution (i.e., single-gland WGS) data obtained by us. We have indeed previously raised concerns (Williams et al. 2018b; Caravagna et al. 2020) regarding the ability to perform such analysis from low-coverage bulk sequencing data with the methods used by the authors (Dentro, Wedge, and Van Loo 2017; Tarabichi et al. 2021).

Our previous assertion that reliable detection of selection from low-coverage single-bulk sequencing data is likely impossible and reconstructed 'clone trees' provide little information on actually present lineages (Caravagna et al. 2020) remains valid in the light of the new results presented here. Generally, the findings highlight the fundamental importance of neutral evolution as a null model in tumour evolution. While these models might have shortcomings, ignoring issues arising from these risks to frustrate future efforts to disentangle the complexity of the evolutionary dynamics in tumours. Potentially, delaying the translation of these into clinically relevant insights.

Last but not least, the presented analysis demonstrates that single-clone multi-region sequencing can provide a general framework to measure the effect of subclonal driver alterations *in vivo*. Due to the ability to fully reconstruct the lineage of individual subclones, this approach allows analysing the effect of driver alterations in the genetic and environmental context they occurred in. Such context-specific effects are thought to be of importance (Berger, Knudson, and Pandolfi 2011) and a large-scale application of the approach used here could be used for the discovery of refined models of *in vivo* driver gene activity.

Due to the scarcity of subclonal selection and the relatively small cohort analysed, a comprehensive analysis of novel driver mutations could not be performed. Still, I am optimistic that in the future, more extensive studies using a similar approach might allow to gain significant insight into the role of genetic and epigenetic alterations in subclonal tumour evolution. Here, similar approaches could distinguish the large amount of 'neutral signal' present in measurements of intra-tumour heterogeneity from that associated with meaningful subclonal selection.

The identification of signal from selected subclones could be especially beneficial for the analysis of non-genetic drivers (Black and McGranahan 2021) that lack appropriate models to describe their dynamics. Indeed, a small subset of cases in which inferred somatic driver mutations did not explain subclonal selection was identified as part of this work. In these cases, non-genetic events could have had a critical role.

#### 6.5 Conclusion

The spatial computational inference on the single-gland sequencing data of the EPICC cohort presented here has allowed to i) predict the presence of selected subclones and ii) estimated growth laws in individual tumours. We propose these as potential evolutionary biomarkers and will assess their predictive ability once sufficient survival data are available. Furthermore, the conducted analysis demonstrated that the developed method allows to identify relevant subclonal selection events and could thus guide the discovery of novel or rare genetic driver alteration in the future.
