## Supplementary Figures for "Assessment of the evolutionary consequence of putative driver mutations in colorectal cancer with spatial multiomic data": figS5.pdf

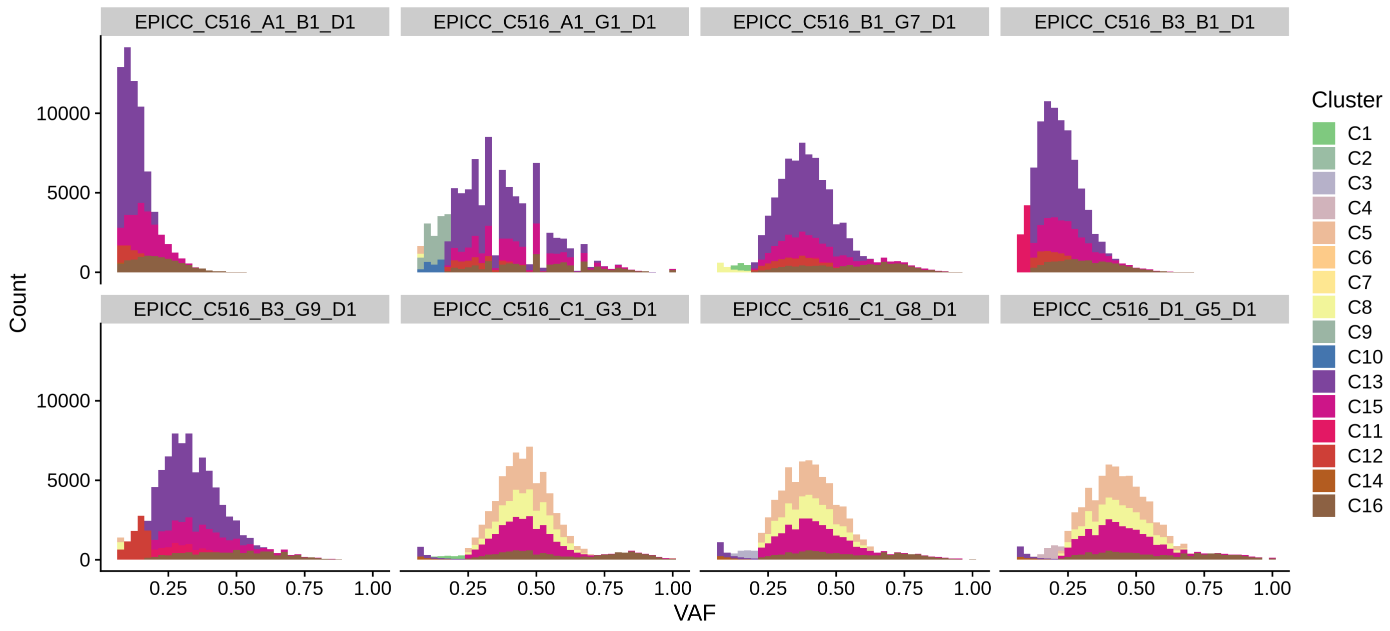
EPICC_C516_A1_B1_D1
EPICC_C516_A1_G1_D1
EPICC_C516_B1_G7_D1
EPICC_C516_B3_B1_D1
EPICC_C516_B3_G9_D1
EPICC_C516_C1_G3_D1
EPICC_C516_C1_G8_D1
EPICC_C516_D1_G5_D1
Count
10000
5000
0
0.25
0.50
0.75
1.00
VAF
Cluster
C1
C2
C3
C4
C5
C6
C7
C8
C9
C10
C13
C15
C11
C12
C14
C16

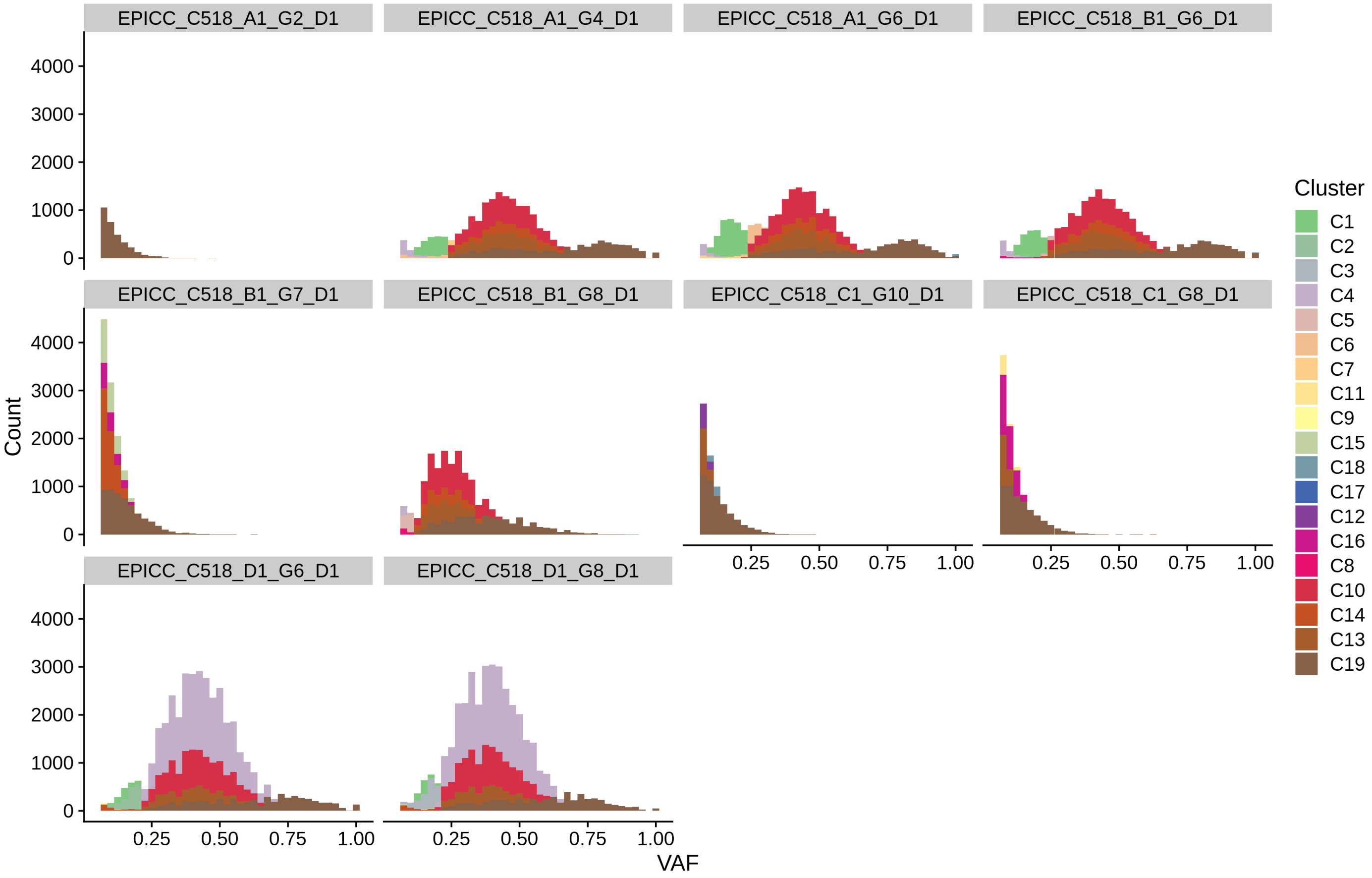
EPICC_C518_A1_G2_D1
EPICC_C518_A1_G4_D1
EPICC_C518_A1_G6_D1
EPICC_C518_B1_G6_D1
EPICC_C518_B1_G7_D1
EPICC_C518_B1_G8_D1
EPICC_C518_C1_G10_D1
EPICC_C518_C1_G8_D1
EPICC_C518_D1_G6_D1
EPICC_C518_D1_G8_D1
Count
0
1000
2000
3000
4000
VAF
0.25
0.50
0.75
1.00
Cluster
C1
C2
C3
C4
C5
C6
C7
C11
C9
C15
C18
C17
C12
C16
C8
C10
C14
C13
C19

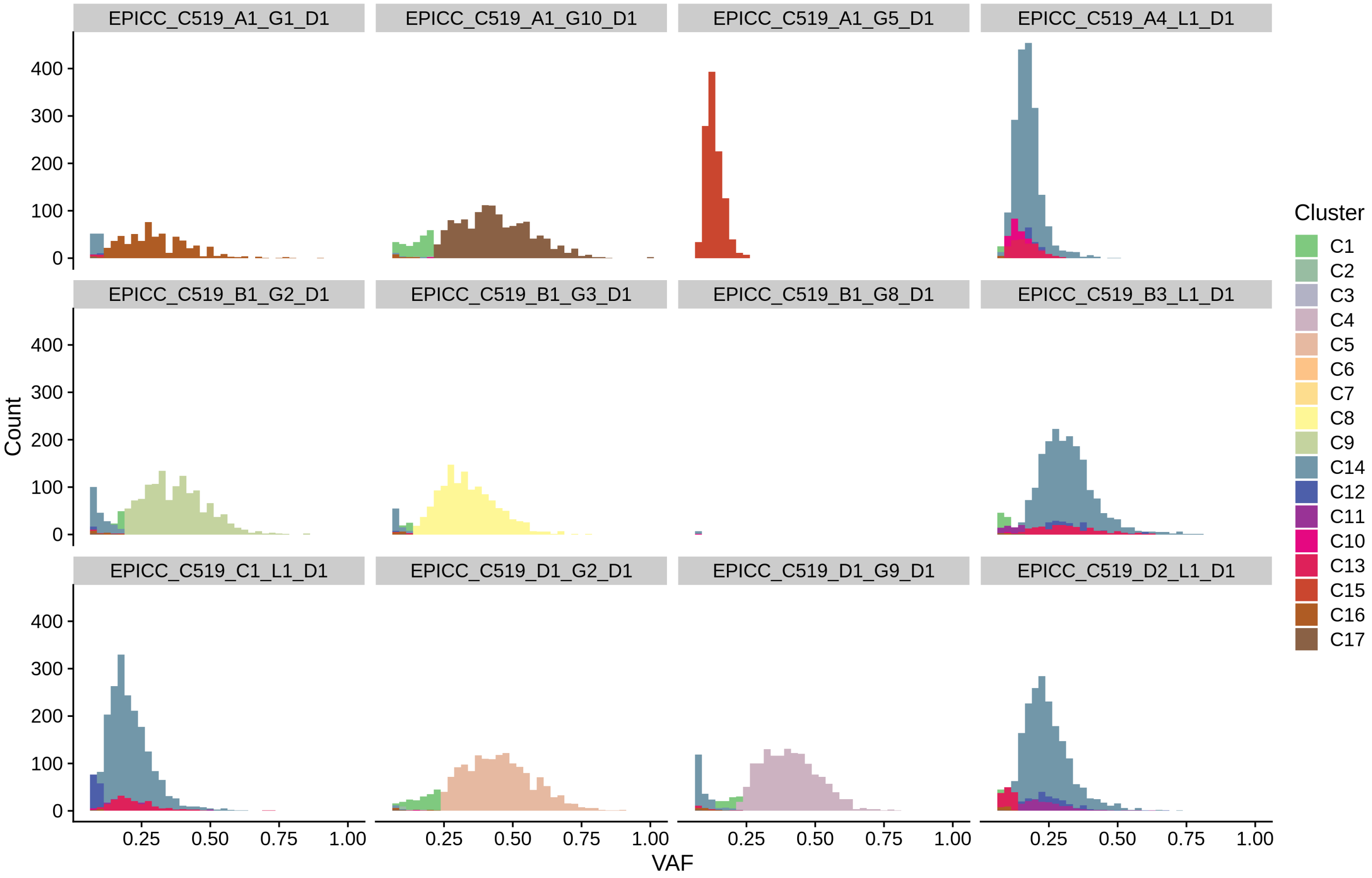

EPICC_C519_A1_G1_D1
EPICC_C519_A1_G10_D1
EPICC_C519_A1_G5_D1
EPICC_C519_A4_L1_D1
EPICC_C519_B1_G2_D1
EPICC_C519_B1_G3_D1
EPICC_C519_B1_G8_D1
EPICC_C519_B3_L1_D1
EPICC_C519_C1_L1_D1
EPICC_C519_D1_G2_D1
EPICC_C519_D1_G9_D1
EPICC_C519_D2_L1_D1
Count
0
100
200
300
400
VAF
0.25
0.50
0.75
1.00
Cluster
C1
C2
C3
C4
C5
C6
C7
C8
C9
C14
C12
C11
C10
C13
C15
C16
C17

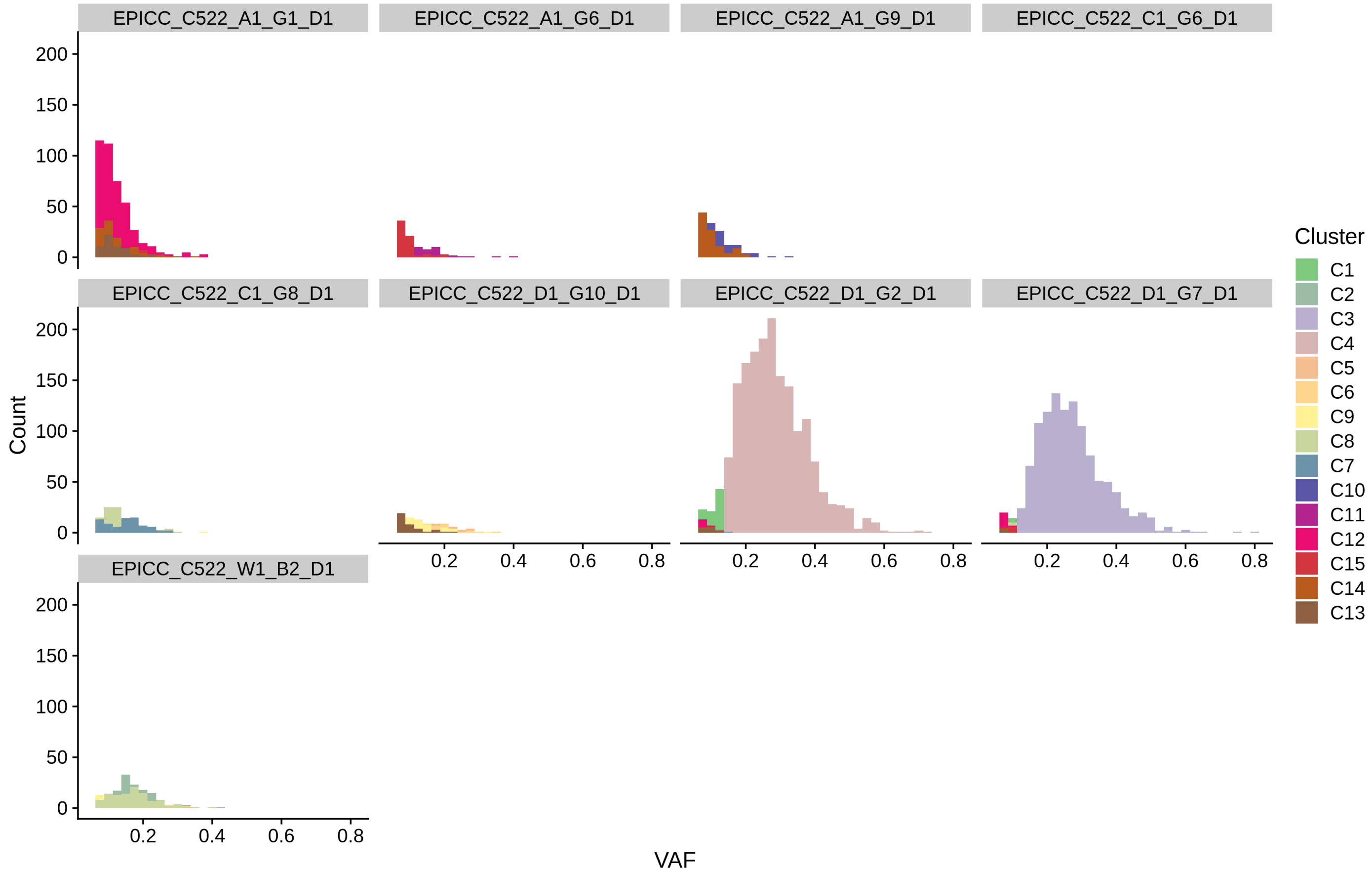

EPICC_C522_A1_G1_D1
EPICC_C522_A1_G6_D1
EPICC_C522_A1_G9_D1
EPICC_C522_C1_G6_D1
EPICC_C522_C1_G8_D1
EPICC_C522_D1_G10_D1
EPICC_C522_D1_G2_D1
EPICC_C522_D1_G7_D1
EPICC_C522_W1_B2_D1
Count
VAF
0
50
100
150
200
0.2
0.4
0.6
0.8
Cluster
C1
C2
C3
C4
C5
C6
C9
C8
C7
C10
C11
C12
C15
C14
C13

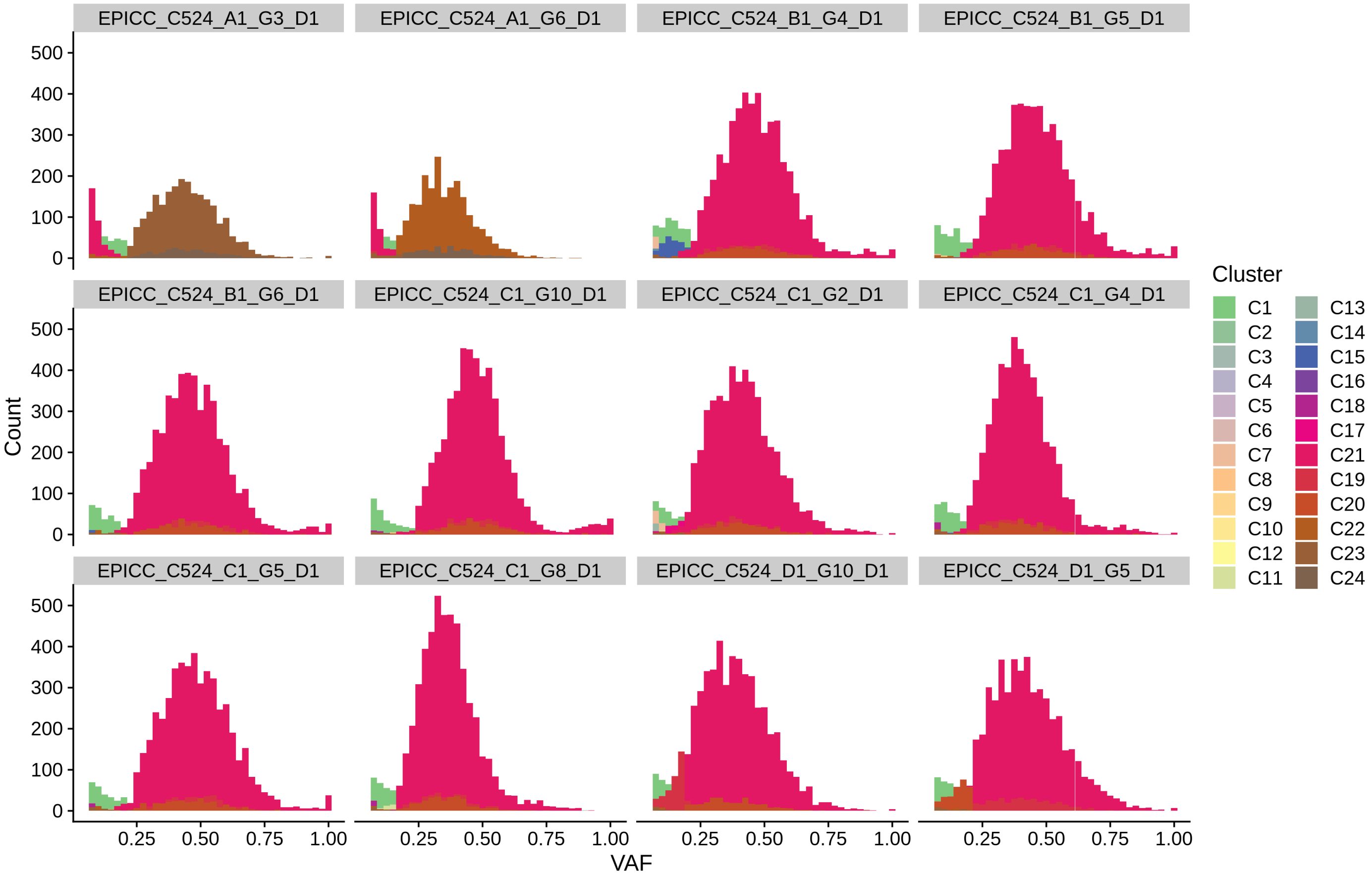
EPICC_C524_A1_G3_D1
EPICC_C524_A1_G6_D1
EPICC_C524_B1_G4_D1
EPICC_C524_B1_G5_D1
EPICC_C524_B1_G6_D1
EPICC_C524_C1_G10_D1
EPICC_C524_C1_G2_D1
EPICC_C524_C1_G4_D1
EPICC_C524_C1_G5_D1
EPICC_C524_C1_G8_D1
EPICC_C524_D1_G10_D1
EPICC_C524_D1_G5_D1
Count
0
100
200
300
400
500
VAF
0.25
0.50
0.75
1.00
Cluster
C1
C2
C3
C4
C5
C6
C7
C8
C9
C10
C12
C11
C13
C14
C15
C16
C18
C17
C21
C19
C20
C22
C23
C24

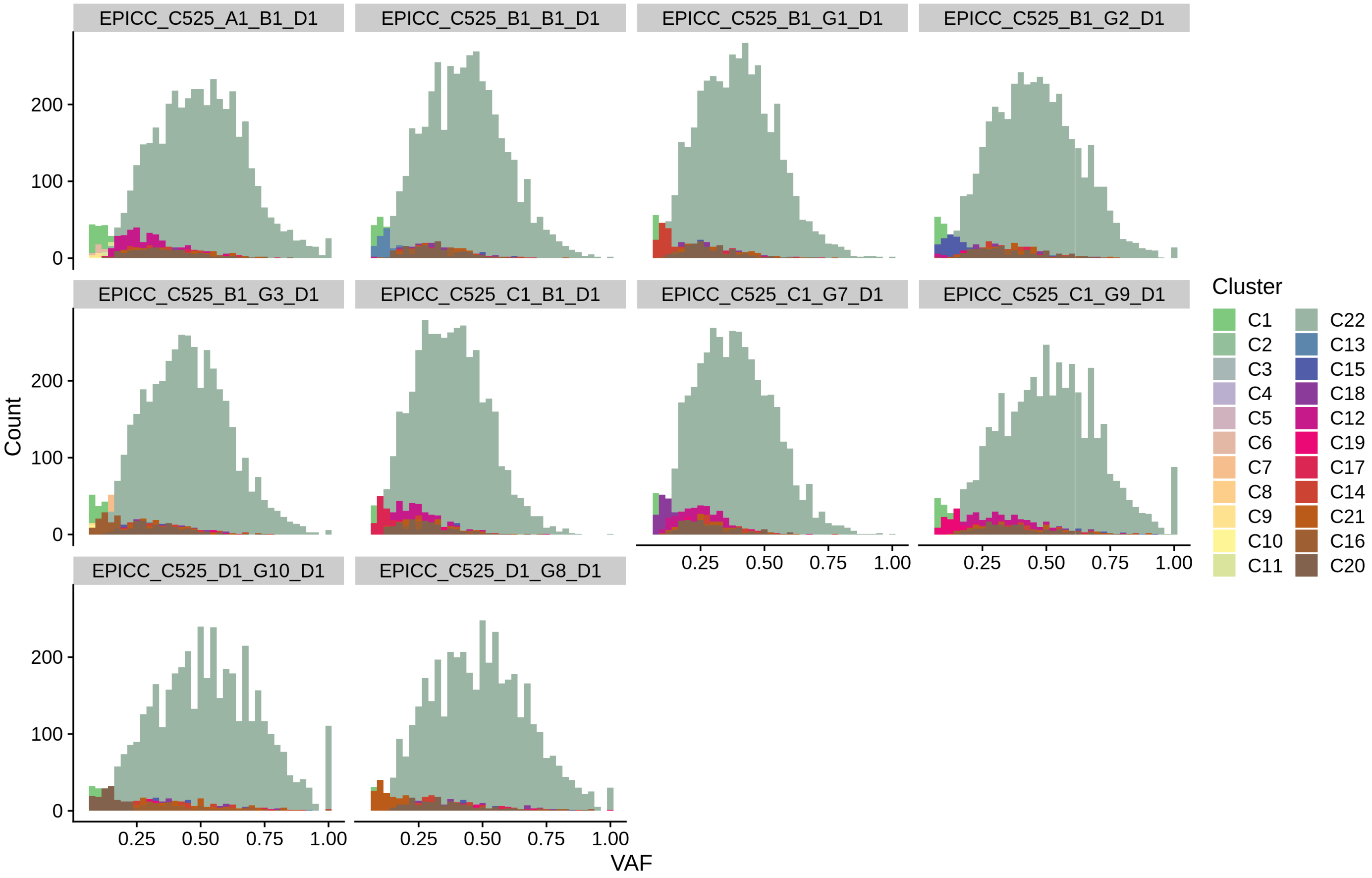

EPICC_C525_A1_B1_D1
EPICC_C525_B1_B1_D1
EPICC_C525_B1_G1_D1
EPICC_C525_B1_G2_D1
EPICC_C525_B1_G3_D1
EPICC_C525_C1_B1_D1
EPICC_C525_C1_G7_D1
EPICC_C525_C1_G9_D1
EPICC_C525_D1_G10_D1
EPICC_C525_D1_G8_D1
Count
0
100
200
VAF
0.25
0.50
0.75
1.00
Cluster
C1
C2
C3
C4
C5
C6
C7
C8
C9
C10
C11
C22
C13
C15
C18
C12
C19
C17
C14
C21
C16
C20

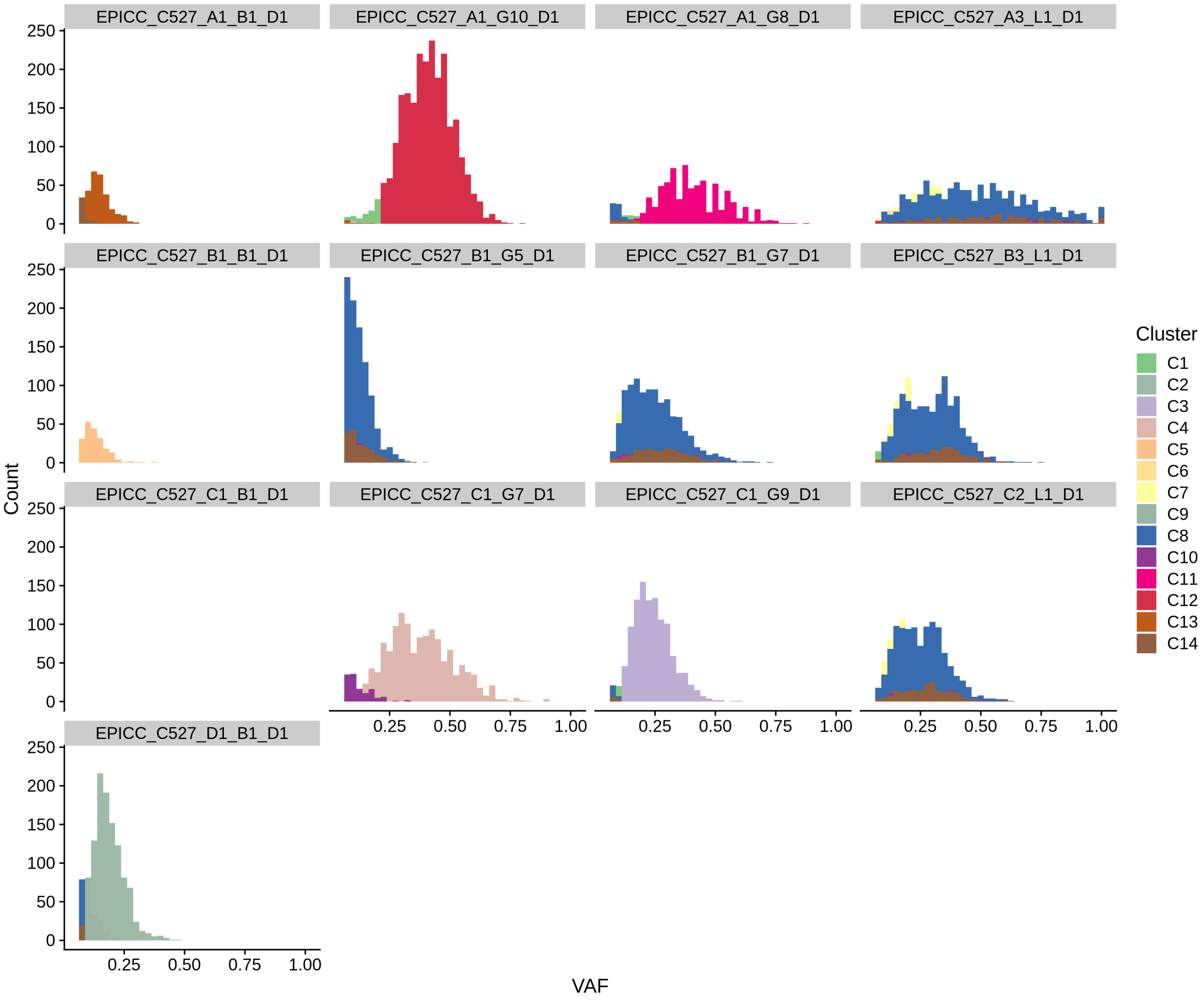
EPICC_C527_A1_B1_D1
EPICC_C527_A1_G10_D1
EPICC_C527_A1_G8_D1
EPICC_C527_A3_L1_D1
EPICC_C527_B1_B1_D1
EPICC_C527_B1_G5_D1
EPICC_C527_B1_G7_D1
EPICC_C527_B3_L1_D1
EPICC_C527_C1_B1_D1
EPICC_C527_C1_G7_D1
EPICC_C527_C1_G9_D1
EPICC_C527_C2_L1_D1
EPICC_C527_D1_B1_D1
Count
0
50
100
150
200
250
VAF
0.25
0.50
0.75
1.00
Cluster
C1
C2
C3
C4
C5
C6
C7
C9
C8
C10
C11
C12
C13
C14

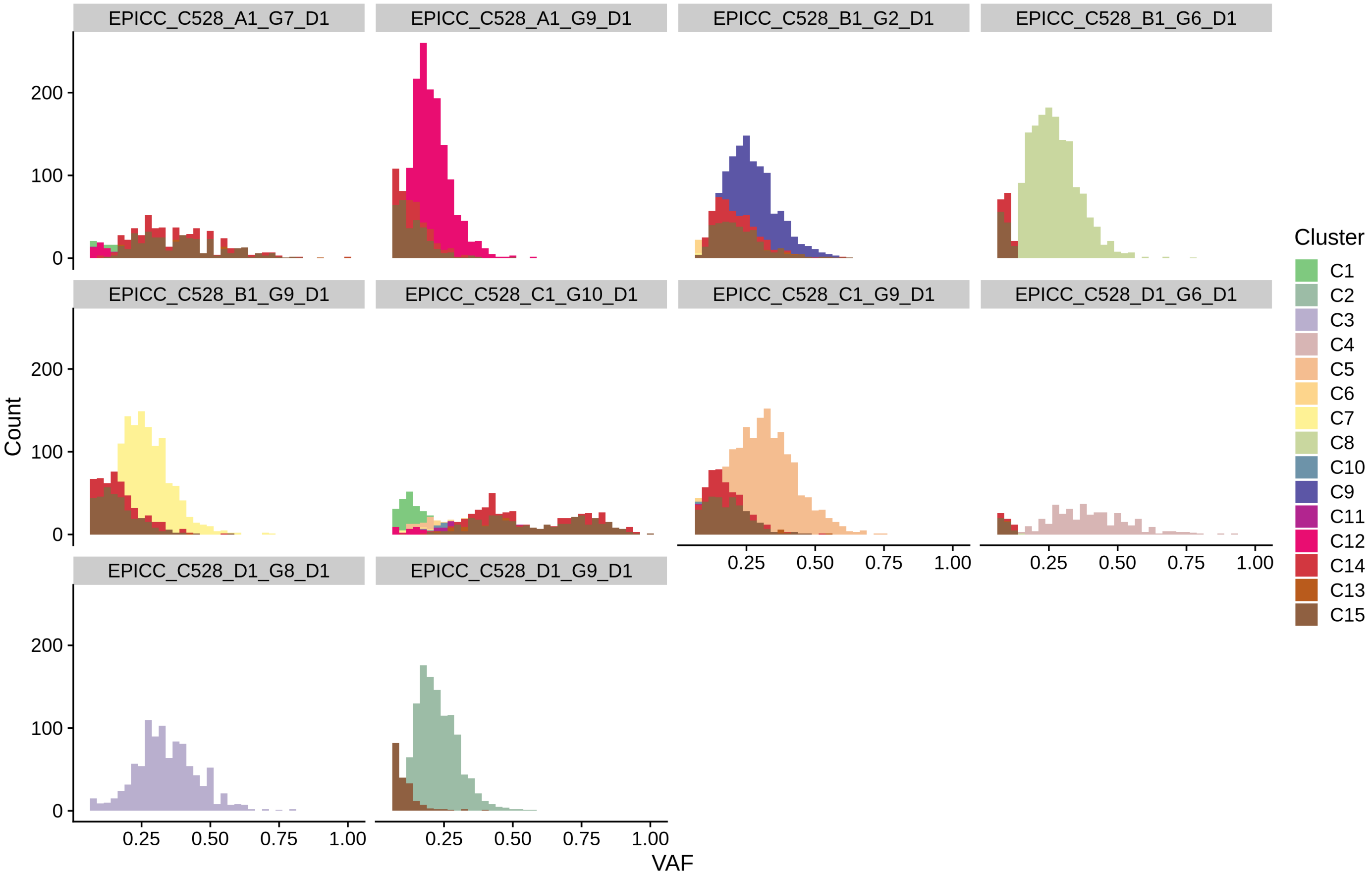

EPICC_C528_A1_G7_D1
EPICC_C528_A1_G9_D1
EPICC_C528_B1_G2_D1
EPICC_C528_B1_G6_D1
EPICC_C528_B1_G9_D1
EPICC_C528_C1_G10_D1
EPICC_C528_C1_G9_D1
EPICC_C528_D1_G6_D1
EPICC_C528_D1_G8_D1
EPICC_C528_D1_G9_D1
Count
0
100
200
VAF
0.25
0.50
0.75
1.00
Cluster
C1
C2
C3
C4
C5
C6
C7
C8
C10
C9
C11
C12
C14
C13
C15

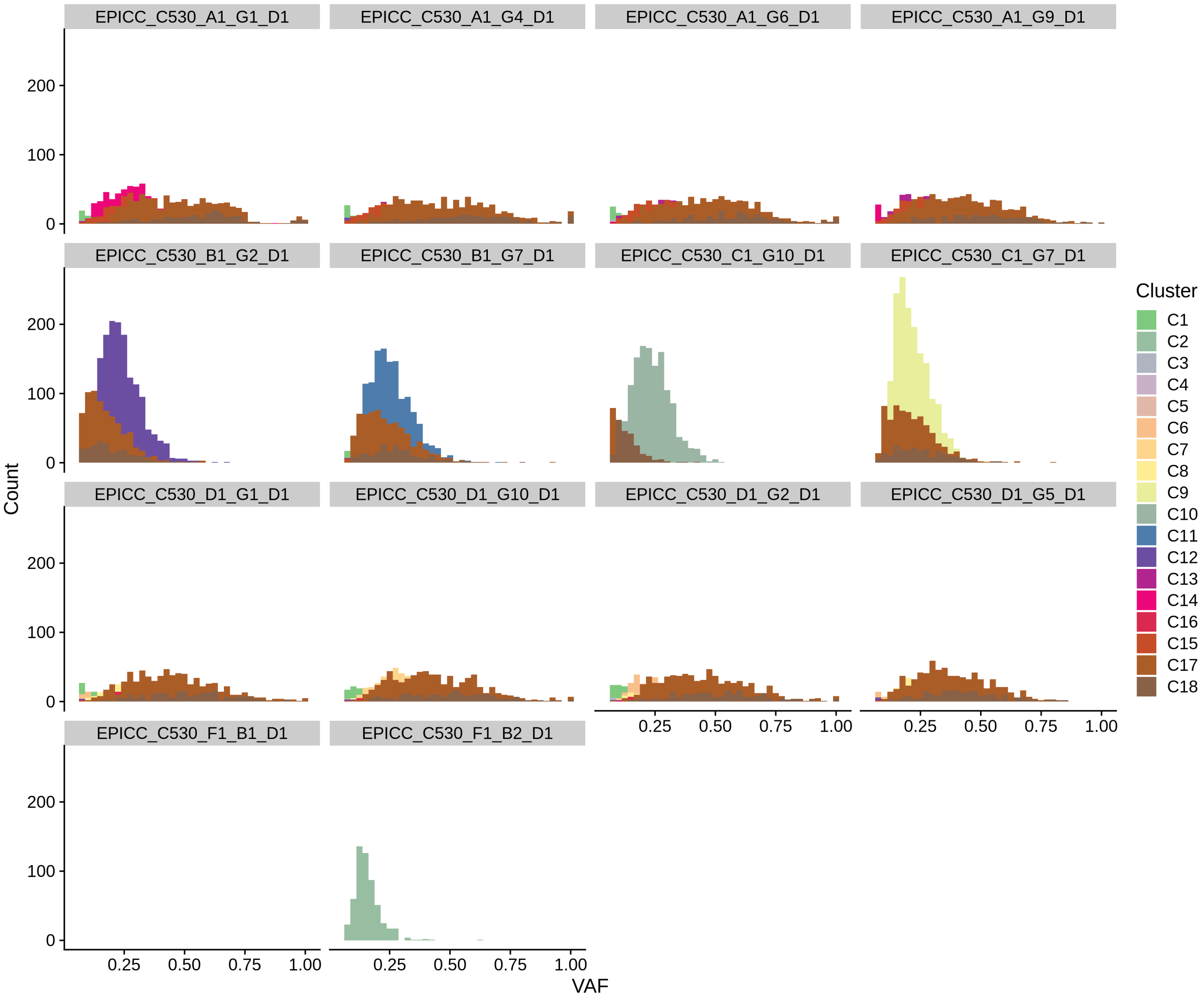

EPICC_C530_A1_G1_D1
EPICC_C530_A1_G4_D1
EPICC_C530_A1_G6_D1
EPICC_C530_A1_G9_D1
EPICC_C530_B1_G2_D1
EPICC_C530_B1_G7_D1
EPICC_C530_C1_G10_D1
EPICC_C530_C1_G7_D1
EPICC_C530_D1_G1_D1
EPICC_C530_D1_G10_D1
EPICC_C530_D1_G2_D1
EPICC_C530_D1_G5_D1
EPICC_C530_F1_B1_D1
EPICC_C530_F1_B2_D1
Count
0
100
200
VAF
0.25
0.50
0.75
1.00
Cluster
C1
C2
C3
C4
C5
C6
C7
C8
C9
C10
C11
C12
C13
C14
C16
C15
C17
C18

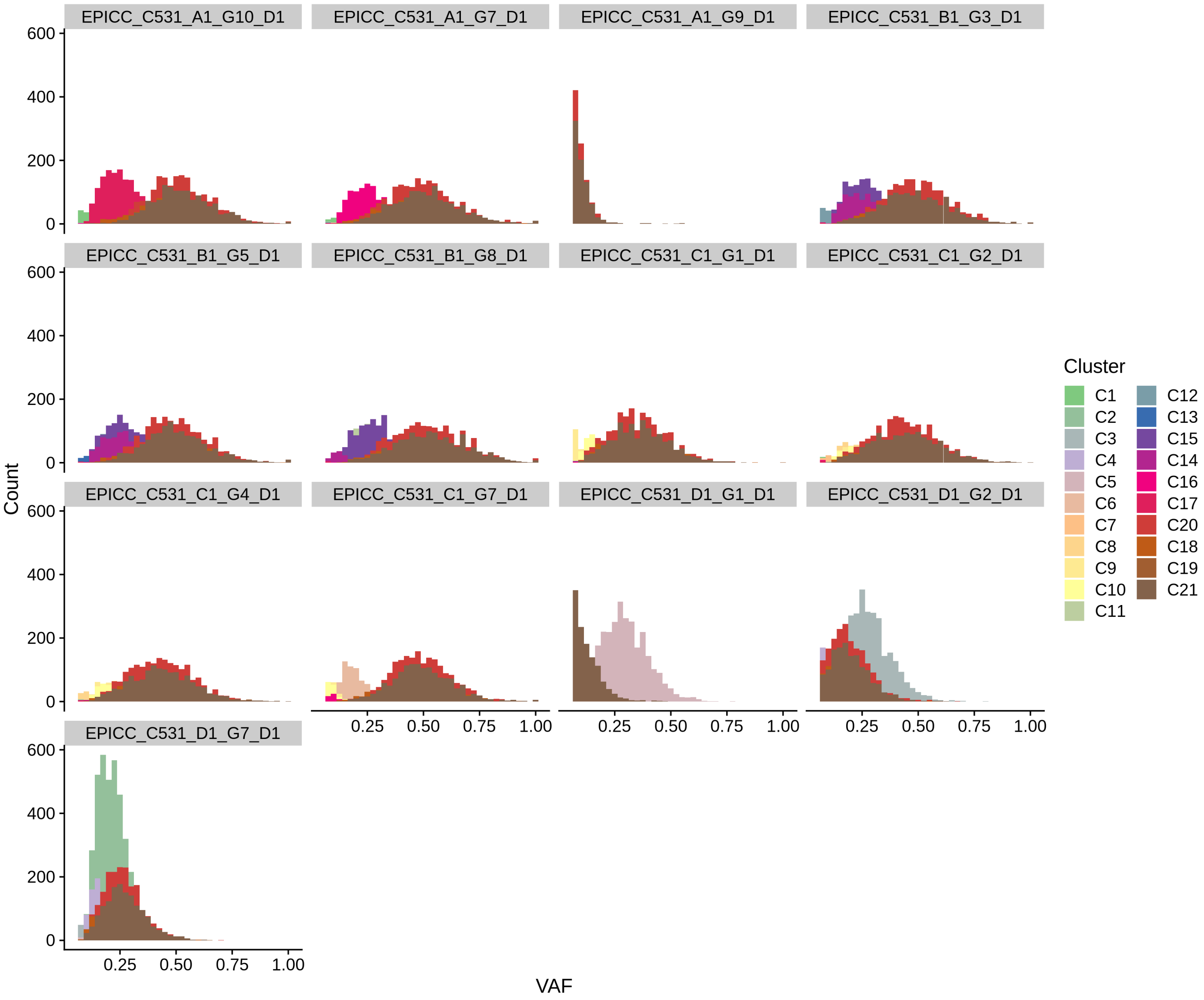
EPICC_C531_A1_G10_D1
EPICC_C531_A1_G7_D1
EPICC_C531_A1_G9_D1
EPICC_C531_B1_G3_D1
EPICC_C531_B1_G5_D1
EPICC_C531_B1_G8_D1
EPICC_C531_C1_G1_D1
EPICC_C531_C1_G2_D1
EPICC_C531_C1_G4_D1
EPICC_C531_C1_G7_D1
EPICC_C531_D1_G1_D1
EPICC_C531_D1_G2_D1
EPICC_C531_D1_G7_D1
Count
0
200
400
600
VAF
0.25
0.50
0.75
1.00
Cluster
C1
C2
C3
C4
C5
C6
C7
C8
C9
C10
C11
C12
C13
C15
C14
C16
C17
C20
C18
C19
C21

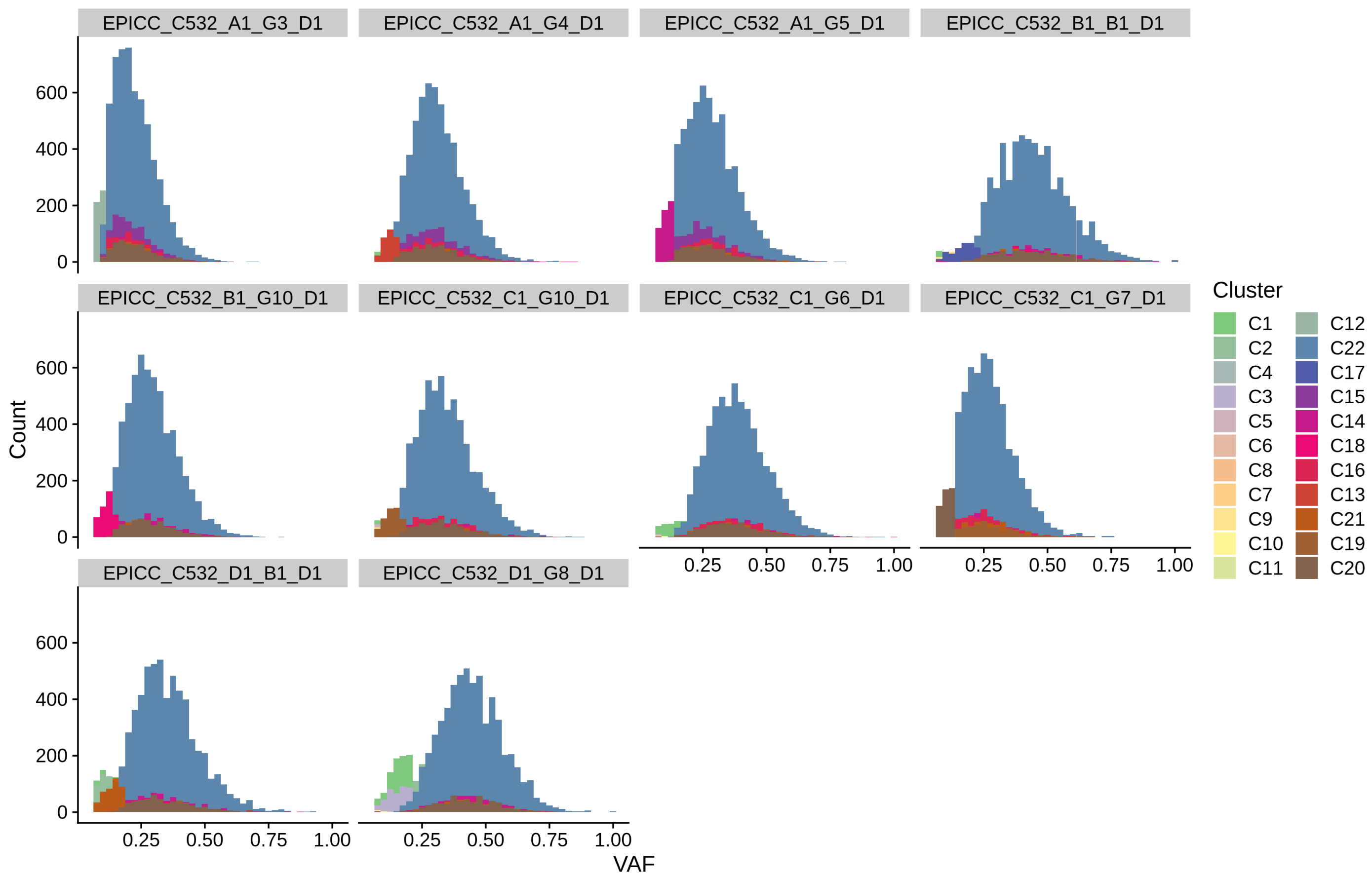

EPICC_C532_A1_G3_D1
EPICC_C532_A1_G4_D1
EPICC_C532_A1_G5_D1
EPICC_C532_B1_B1_D1
EPICC_C532_B1_G10_D1
EPICC_C532_C1_G10_D1
EPICC_C532_C1_G6_D1
EPICC_C532_C1_G7_D1
EPICC_C532_D1_B1_D1
EPICC_C532_D1_G8_D1
Count
0
200
400
600
0.25
0.50
0.75
1.00
VAF
Cluster
C1
C2
C4
C3
C5
C6
C8
C7
C9
C10
C11
C12
C22
C17
C15
C14
C18
C16
C13
C21
C19
C20

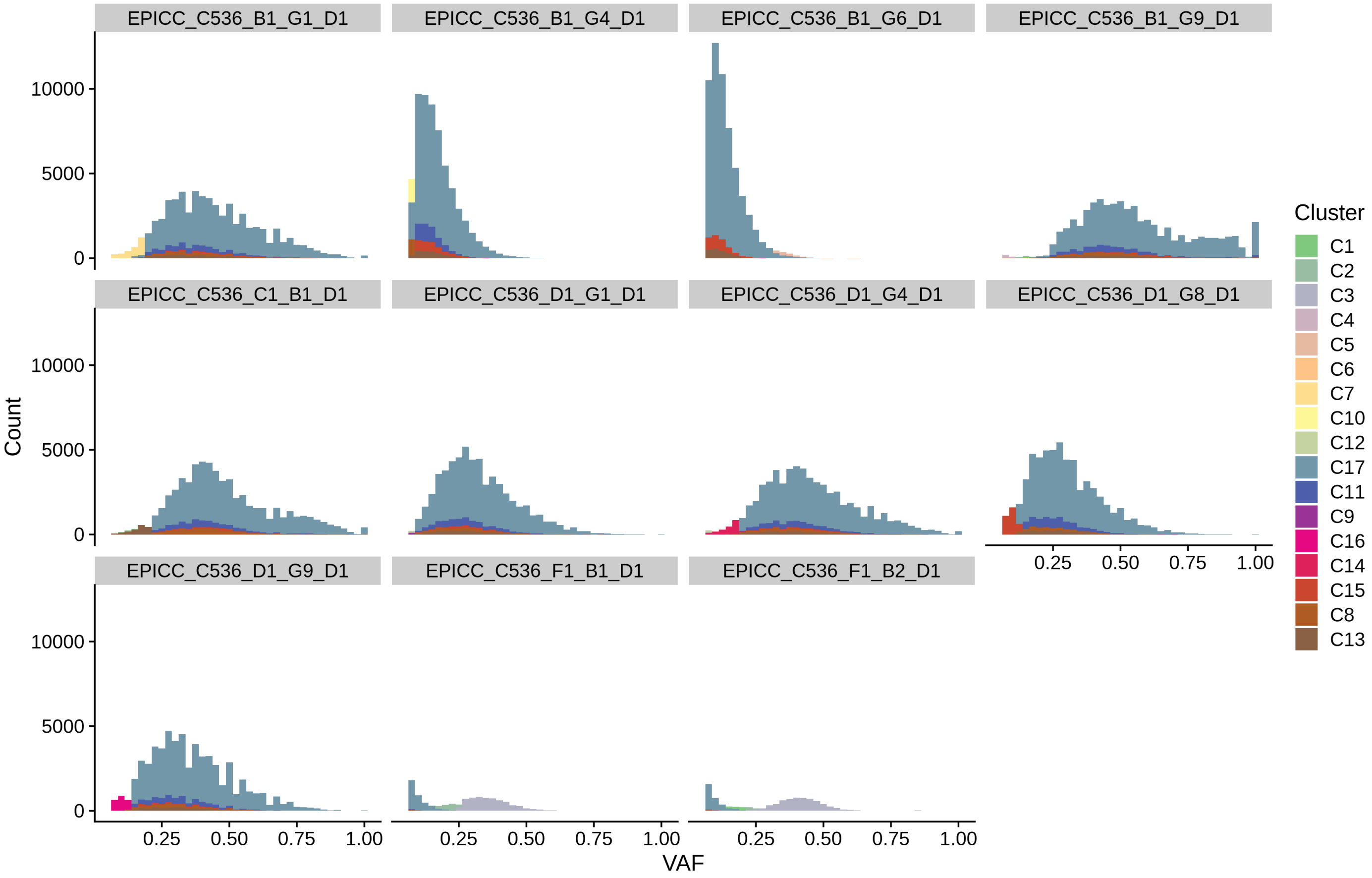

EPICC_C536_B1_G1_D1
EPICC_C536_B1_G4_D1
EPICC_C536_B1_G6_D1
EPICC_C536_B1_G9_D1
EPICC_C536_C1_B1_D1
EPICC_C536_D1_G1_D1
EPICC_C536_D1_G4_D1
EPICC_C536_D1_G8_D1
EPICC_C536_D1_G9_D1
EPICC_C536_F1_B1_D1
EPICC_C536_F1_B2_D1
Count
0
5000
10000
VAF
0.25
0.50
0.75
1.00
Cluster
C1
C2
C3
C4
C5
C6
C7
C10
C12
C17
C11
C9
C16
C14
C15
C8
C13

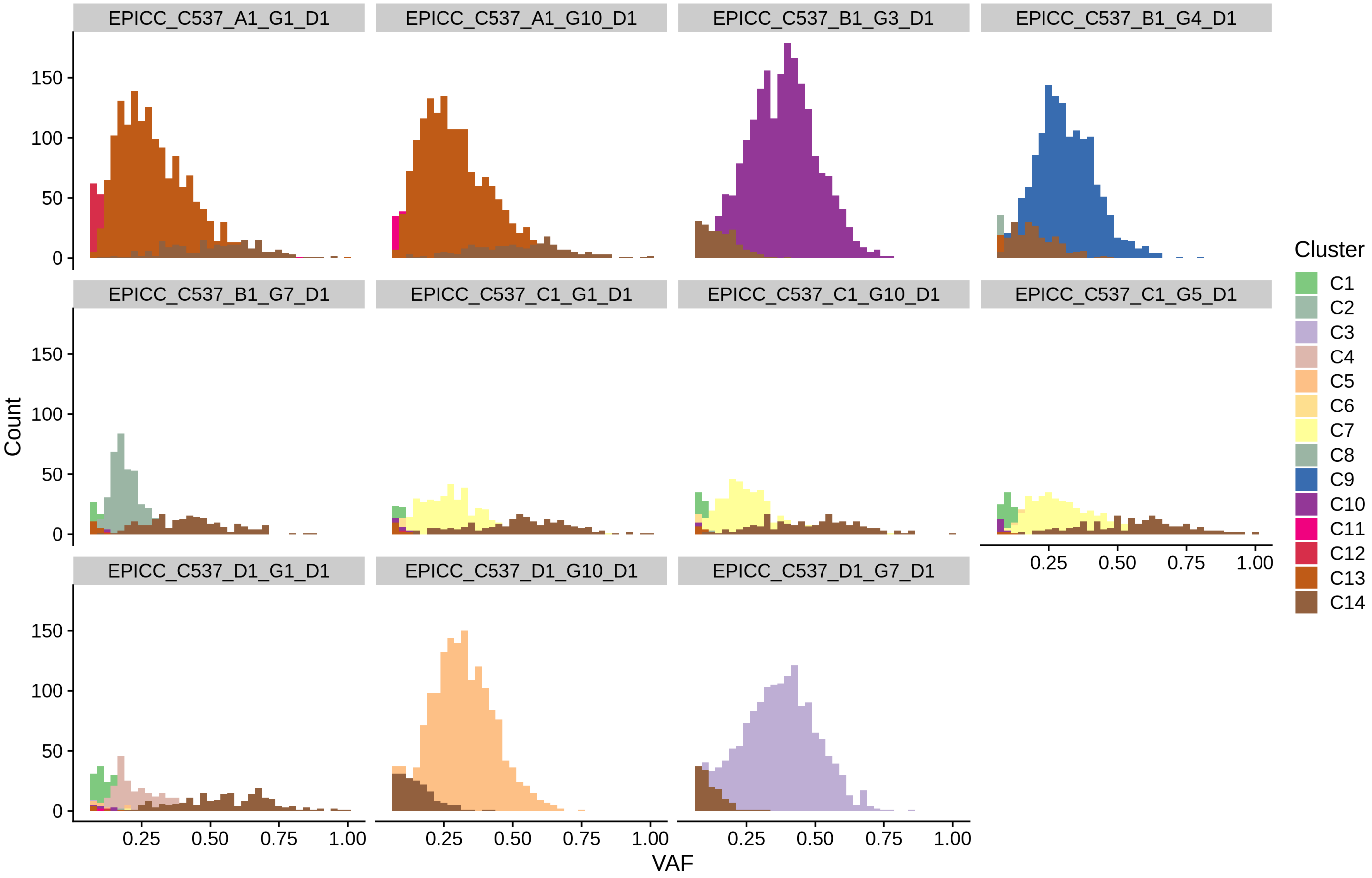

EPICC_C537_A1_G1_D1
EPICC_C537_A1_G10_D1
EPICC_C537_B1_G3_D1
EPICC_C537_B1_G4_D1
EPICC_C537_B1_G7_D1
EPICC_C537_C1_G1_D1
EPICC_C537_C1_G10_D1
EPICC_C537_C1_G5_D1
EPICC_C537_D1_G1_D1
EPICC_C537_D1_G10_D1
EPICC_C537_D1_G7_D1
Count
0
50
100
150
VAF
0.25
0.50
0.75
1.00
Cluster
C1
C2
C3
C4
C5
C6
C7
C8
C9
C10
C11
C12
C13
C14

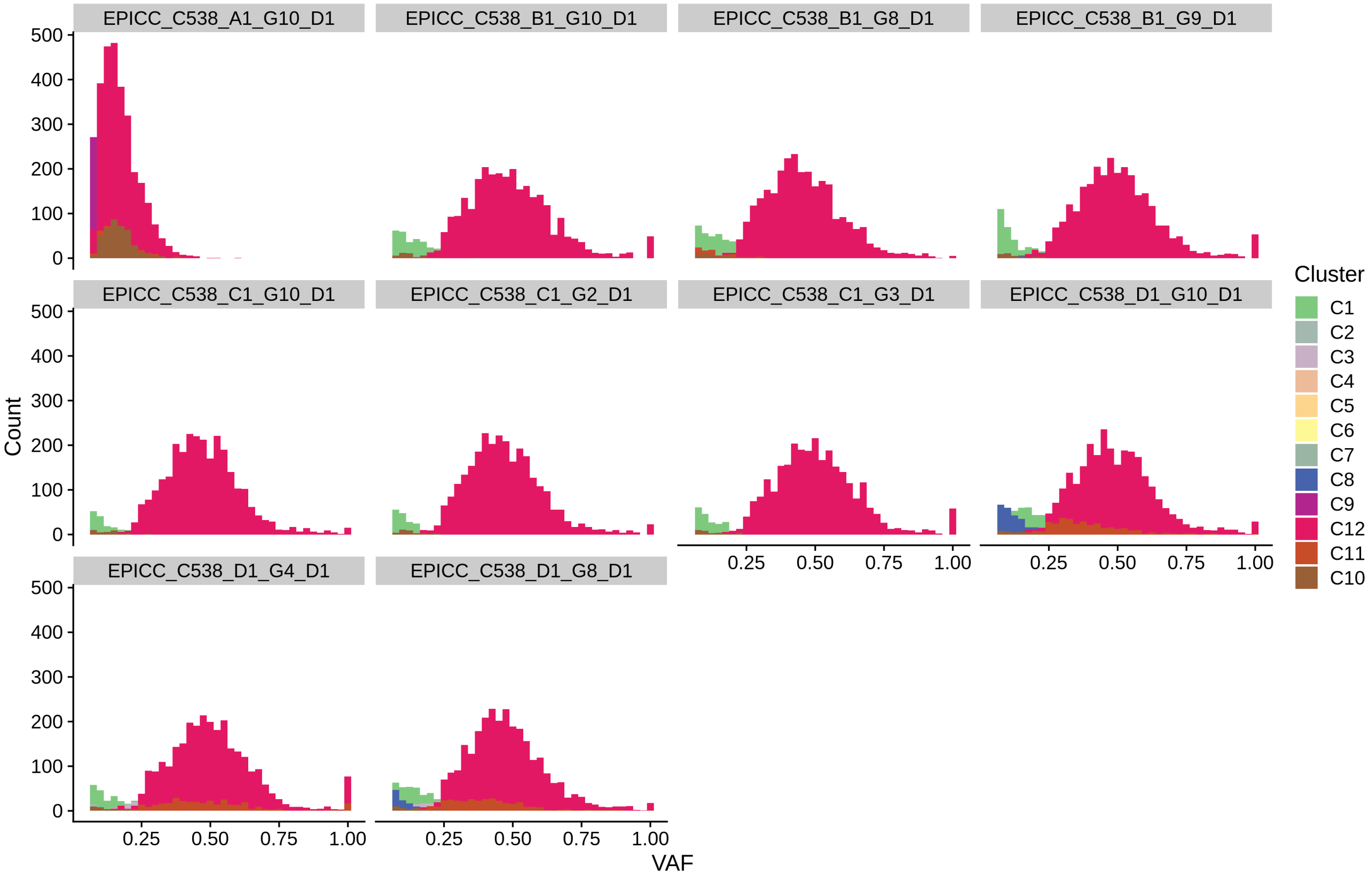
EPICC_C538_A1_G10_D1
EPICC_C538_B1_G10_D1
EPICC_C538_B1_G8_D1
EPICC_C538_B1_G9_D1
EPICC_C538_C1_G10_D1
EPICC_C538_C1_G2_D1
EPICC_C538_C1_G3_D1
EPICC_C538_D1_G10_D1
EPICC_C538_D1_G4_D1
EPICC_C538_D1_G8_D1
Count
0
100
200
300
400
500
VAF
0.25
0.50
0.75
1.00
Cluster
C1
C2
C3
C4
C5
C6
C7
C8
C9
C12
C11
C10

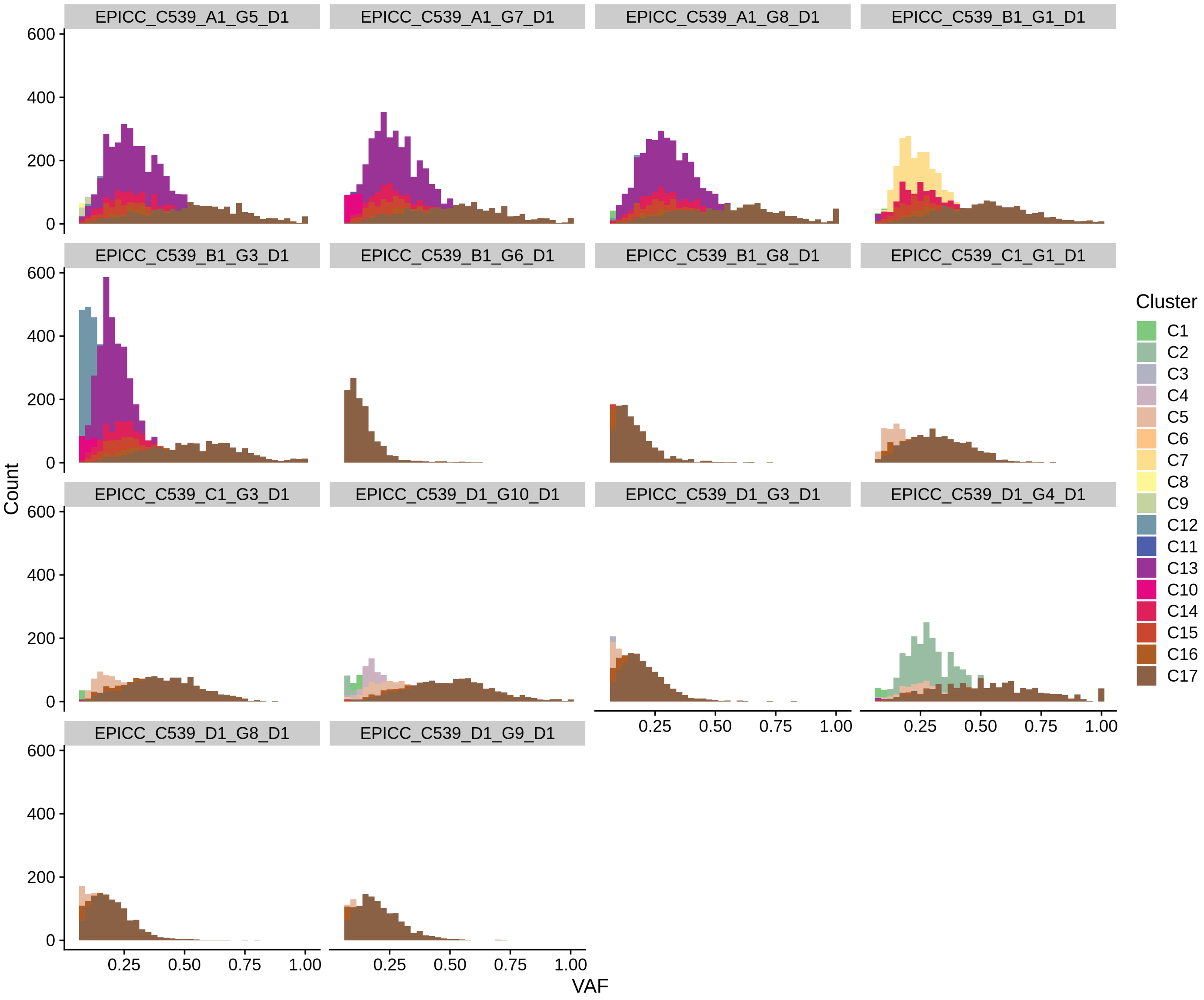

EPICC_C539_A1_G5_D1
EPICC_C539_A1_G7_D1
EPICC_C539_A1_G8_D1
EPICC_C539_B1_G1_D1
EPICC_C539_B1_G3_D1
EPICC_C539_B1_G6_D1
EPICC_C539_B1_G8_D1
EPICC_C539_C1_G1_D1
EPICC_C539_C1_G3_D1
EPICC_C539_D1_G10_D1
EPICC_C539_D1_G3_D1
EPICC_C539_D1_G4_D1
EPICC_C539_D1_G8_D1
EPICC_C539_D1_G9_D1
Count
0
200
400
600
0.25
0.50
0.75
1.00
VAF
Cluster
C1
C2
C3
C4
C5
C6
C7
C8
C9
C12
C11
C13
C10
C14
C15
C16
C17

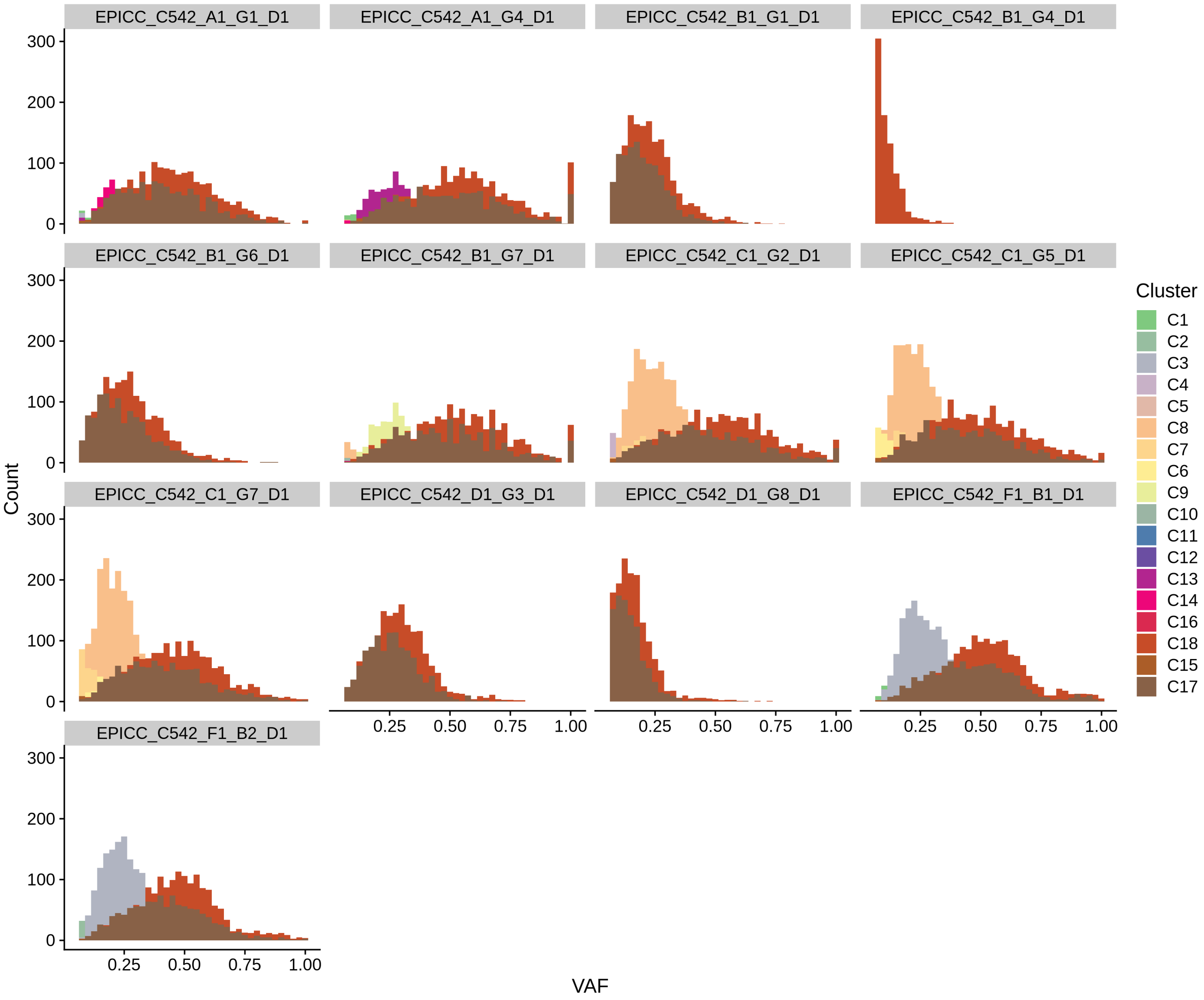

EPICC_C542_A1_G1_D1
EPICC_C542_A1_G4_D1
EPICC_C542_B1_G1_D1
EPICC_C542_B1_G4_D1
EPICC_C542_B1_G6_D1
EPICC_C542_B1_G7_D1
EPICC_C542_C1_G2_D1
EPICC_C542_C1_G5_D1
EPICC_C542_C1_G7_D1
EPICC_C542_D1_G3_D1
EPICC_C542_D1_G8_D1
EPICC_C542_F1_B1_D1
EPICC_C542_F1_B2_D1
Count
0
100
200
300
0.25
0.50
0.75
1.00
VAF
Cluster
C1
C2
C3
C4
C5
C8
C7
C6
C9
C10
C11
C12
C13
C14
C16
C18
C15
C17

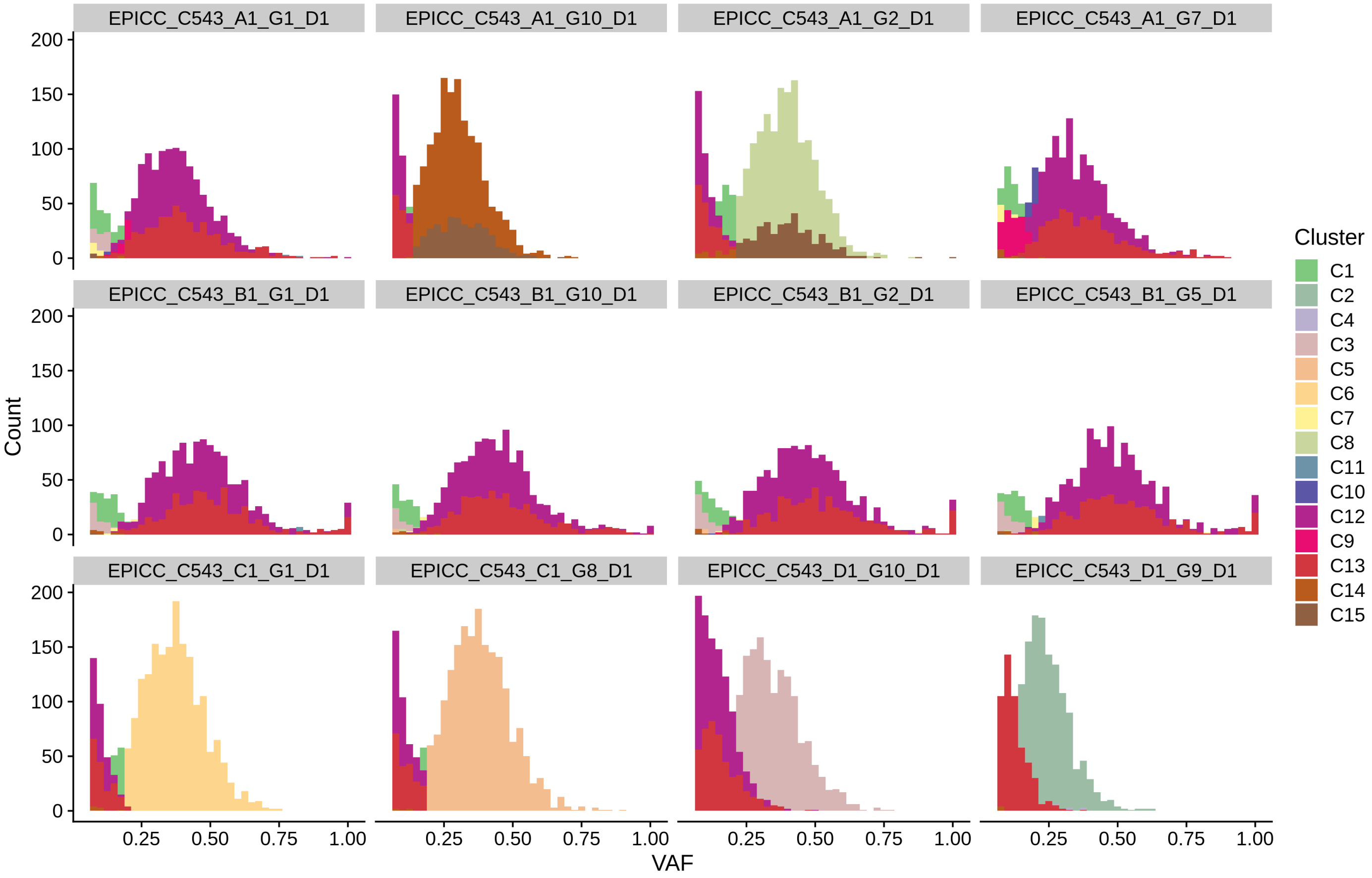
EPICC_C543_A1_G1_D1
EPICC_C543_A1_G10_D1
EPICC_C543_A1_G2_D1
EPICC_C543_A1_G7_D1
EPICC_C543_B1_G1_D1
EPICC_C543_B1_G10_D1
EPICC_C543_B1_G2_D1
EPICC_C543_B1_G5_D1
EPICC_C543_C1_G1_D1
EPICC_C543_C1_G8_D1
EPICC_C543_D1_G10_D1
EPICC_C543_D1_G9_D1
Count
0
50
100
150
200
VAF
0.25
0.50
0.75
1.00
Cluster
C1
C2
C4
C3
C5
C6
C7
C8
C11
C10
C12
C9
C13
C14
C15

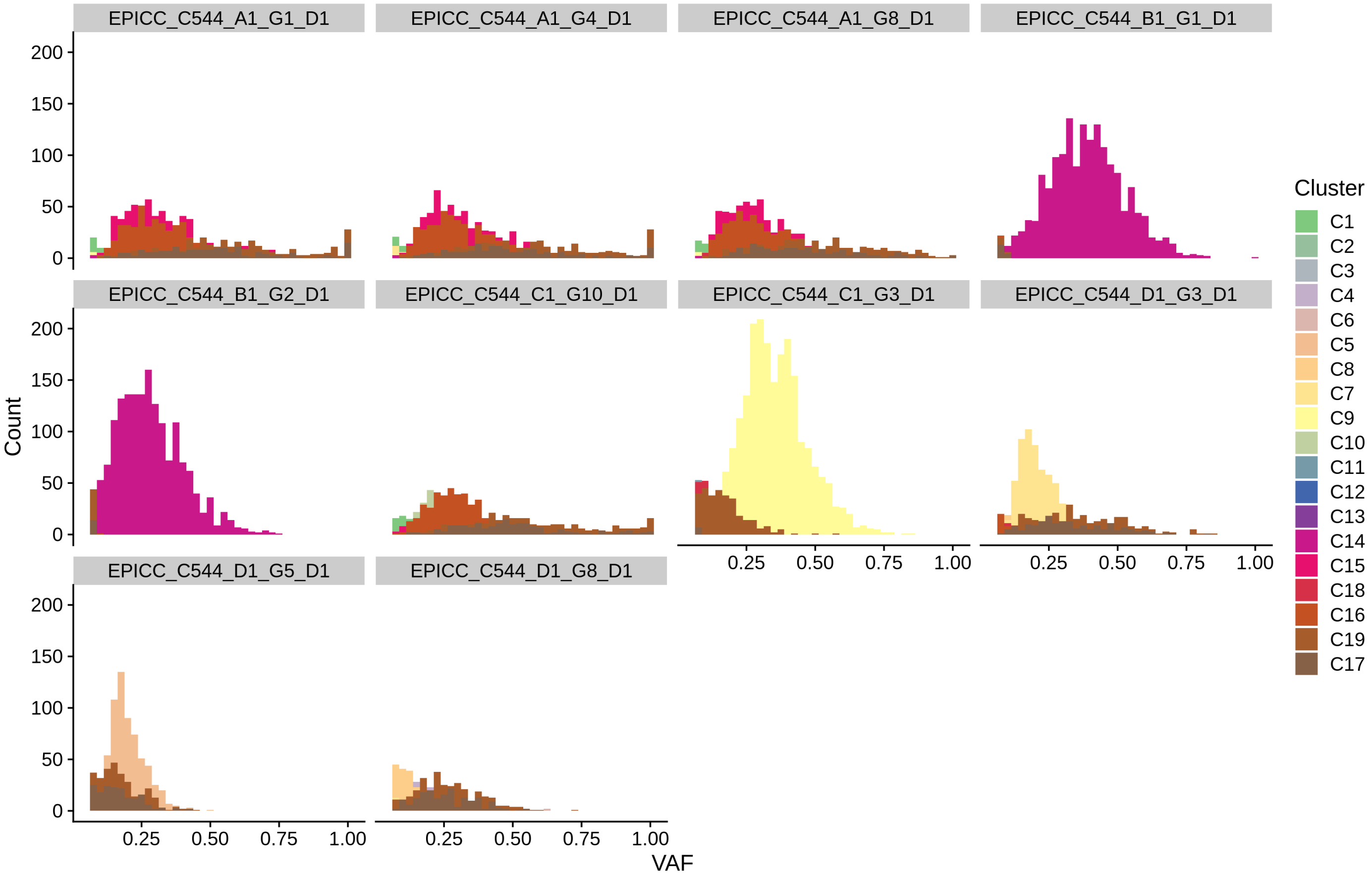
EPICC_C544_A1_G1_D1
EPICC_C544_A1_G4_D1
EPICC_C544_A1_G8_D1
EPICC_C544_B1_G1_D1
EPICC_C544_B1_G2_D1
EPICC_C544_C1_G10_D1
EPICC_C544_C1_G3_D1
EPICC_C544_D1_G3_D1
EPICC_C544_D1_G5_D1
EPICC_C544_D1_G8_D1
Count
VAF
0
50
100
150
200
0.25
0.50
0.75
1.00
Cluster
C1
C2
C3
C4
C6
C5
C8
C7
C9
C10
C11
C12
C13
C14
C15
C18
C16
C19
C17

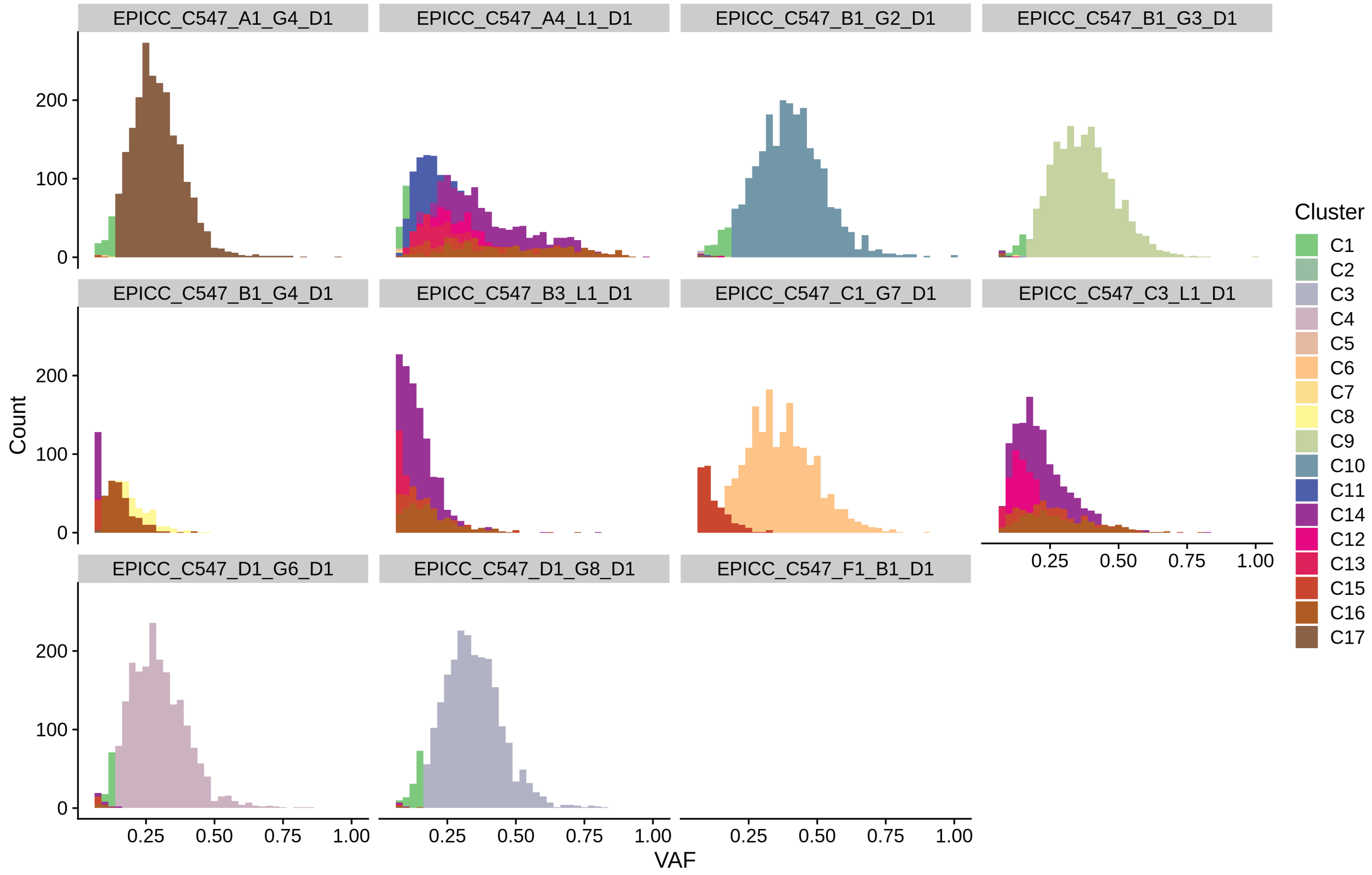
EPICC_C547_A1_G4_D1
EPICC_C547_A4_L1_D1
EPICC_C547_B1_G2_D1
EPICC_C547_B1_G3_D1
EPICC_C547_B1_G4_D1
EPICC_C547_B3_L1_D1
EPICC_C547_C1_G7_D1
EPICC_C547_C3_L1_D1
EPICC_C547_D1_G6_D1
EPICC_C547_D1_G8_D1
EPICC_C547_F1_B1_D1
Count
0
100
200
VAF
0.25
0.50
0.75
1.00
Cluster
C1
C2
C3
C4
C5
C6
C7
C8
C9
C10
C11
C14
C12
C13
C15
C16
C17

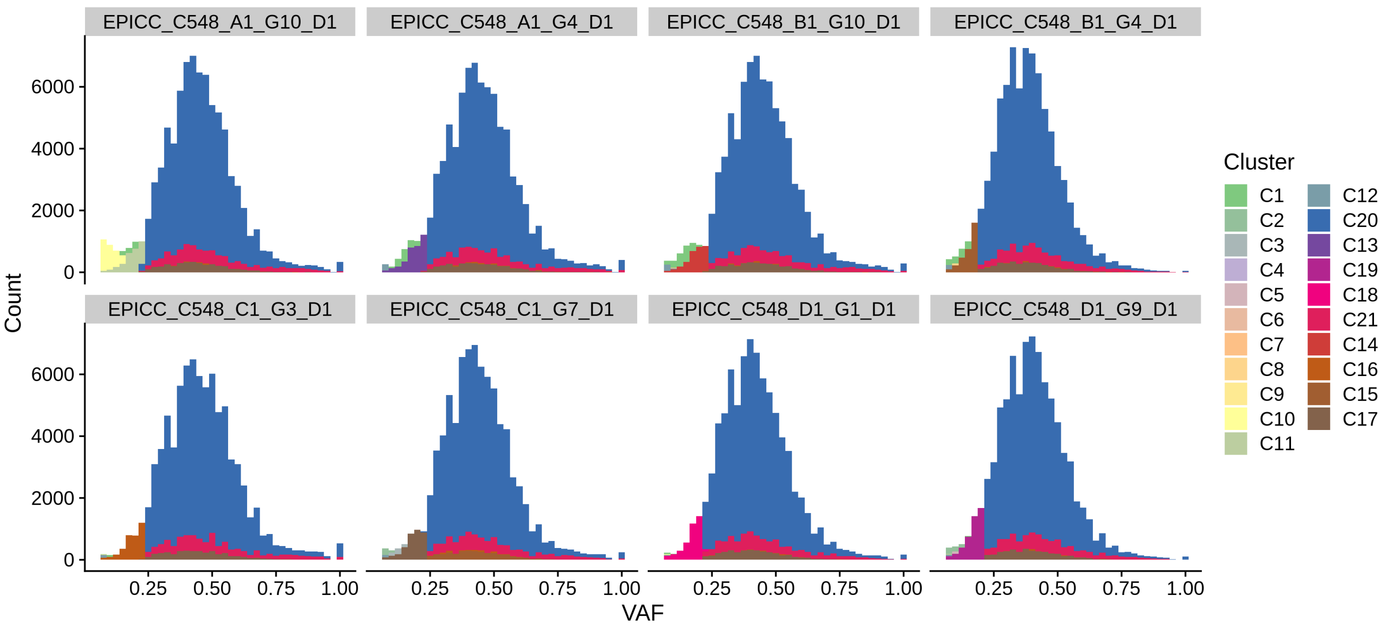
EPICC_C548_A1_G10_D1
EPICC_C548_A1_G4_D1
EPICC_C548_B1_G10_D1
EPICC_C548_B1_G4_D1
EPICC_C548_C1_G3_D1
EPICC_C548_C1_G7_D1
EPICC_C548_D1_G1_D1
EPICC_C548_D1_G9_D1
Count
VAF
0
2000
4000
6000
0.25
0.50
0.75
1.00
Cluster
C1
C2
C3
C4
C5
C6
C7
C8
C9
C10
C11
C12
C20
C13
C19
C18
C21
C14
C16
C15
C17

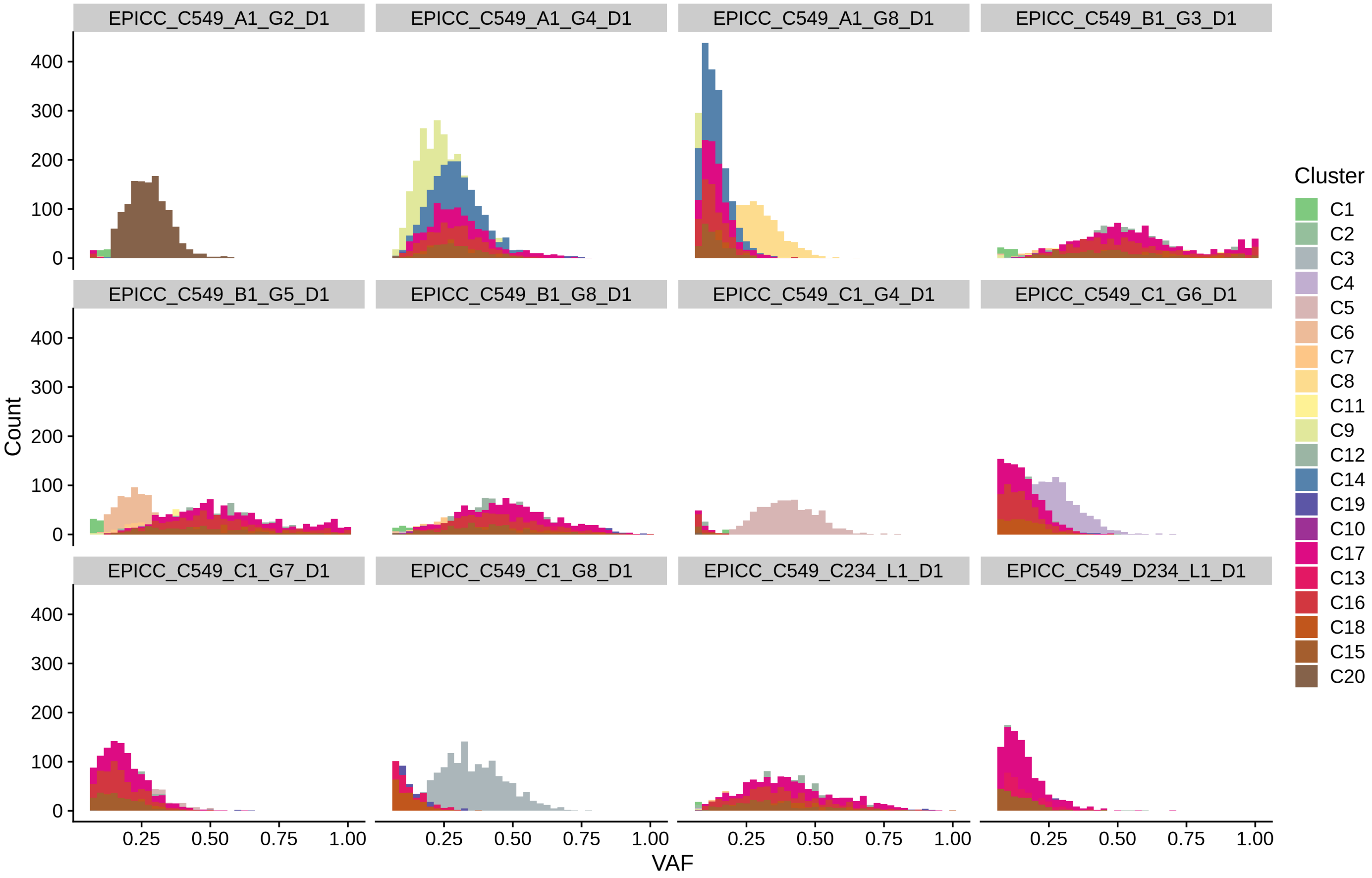

EPICC_C549_A1_G2_D1
EPICC_C549_A1_G4_D1
EPICC_C549_A1_G8_D1
EPICC_C549_B1_G3_D1
EPICC_C549_B1_G5_D1
EPICC_C549_B1_G8_D1
EPICC_C549_C1_G4_D1
EPICC_C549_C1_G6_D1
EPICC_C549_C1_G7_D1
EPICC_C549_C1_G8_D1
EPICC_C549_C234_L1_D1
EPICC_C549_D234_L1_D1
Count
0
100
200
300
400
VAF
0.25
0.50
0.75
1.00
Cluster
C1
C2
C3
C4
C5
C6
C7
C8
C11
C9
C12
C14
C19
C10
C17
C13
C16
C18
C15
C20

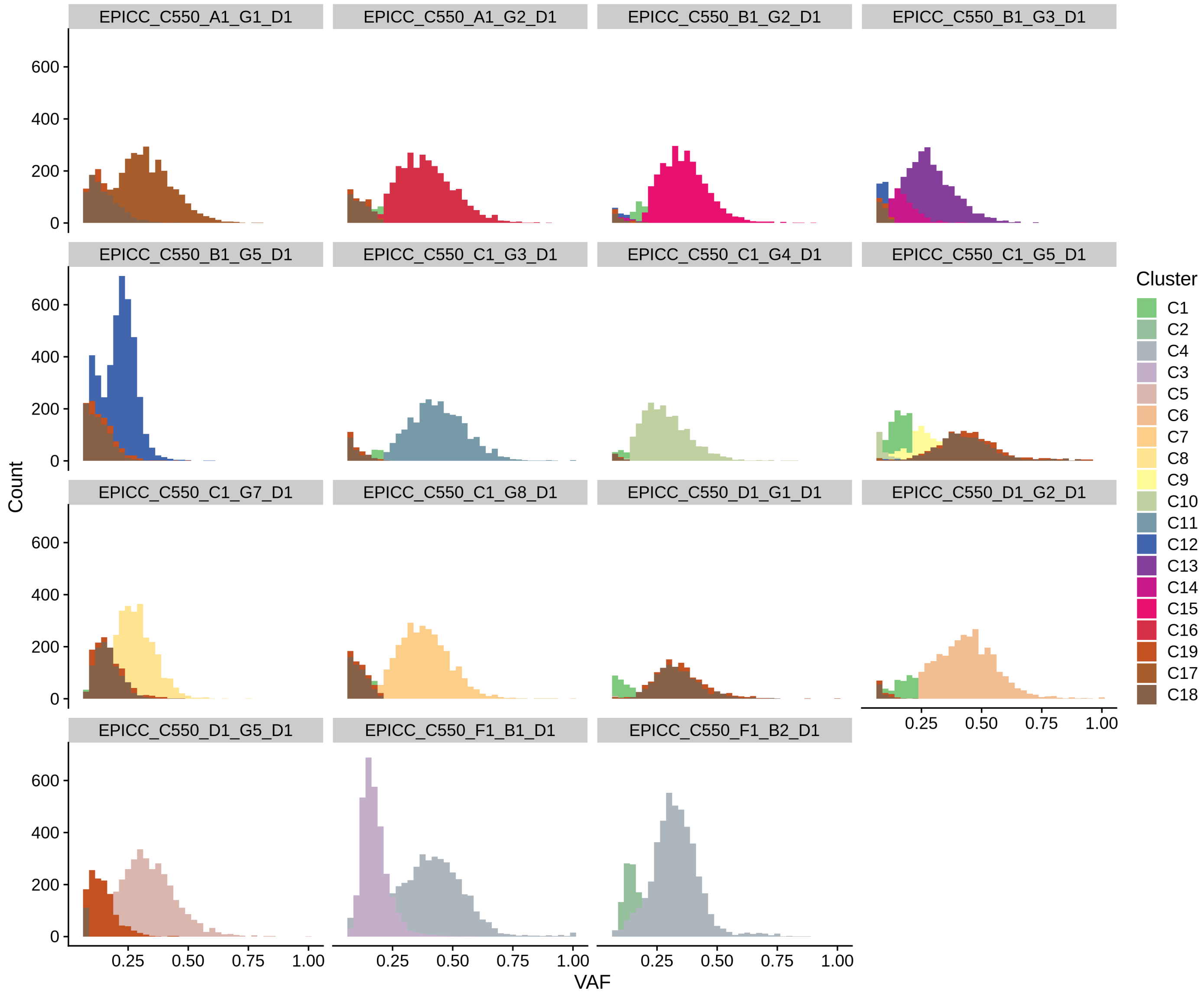
EPICC_C550_A1_G1_D1
EPICC_C550_A1_G2_D1
EPICC_C550_B1_G2_D1
EPICC_C550_B1_G3_D1
EPICC_C550_B1_G5_D1
EPICC_C550_C1_G3_D1
EPICC_C550_C1_G4_D1
EPICC_C550_C1_G5_D1
EPICC_C550_C1_G7_D1
EPICC_C550_C1_G8_D1
EPICC_C550_D1_G1_D1
EPICC_C550_D1_G2_D1
EPICC_C550_D1_G5_D1
EPICC_C550_F1_B1_D1
EPICC_C550_F1_B2_D1
Count
0
200
400
600
VAF
0.25
0.50
0.75
1.00
Cluster
C1
C2
C4
C3
C5
C6
C7
C8
C9
C10
C11
C12
C13
C14
C15
C16
C19
C17
C18

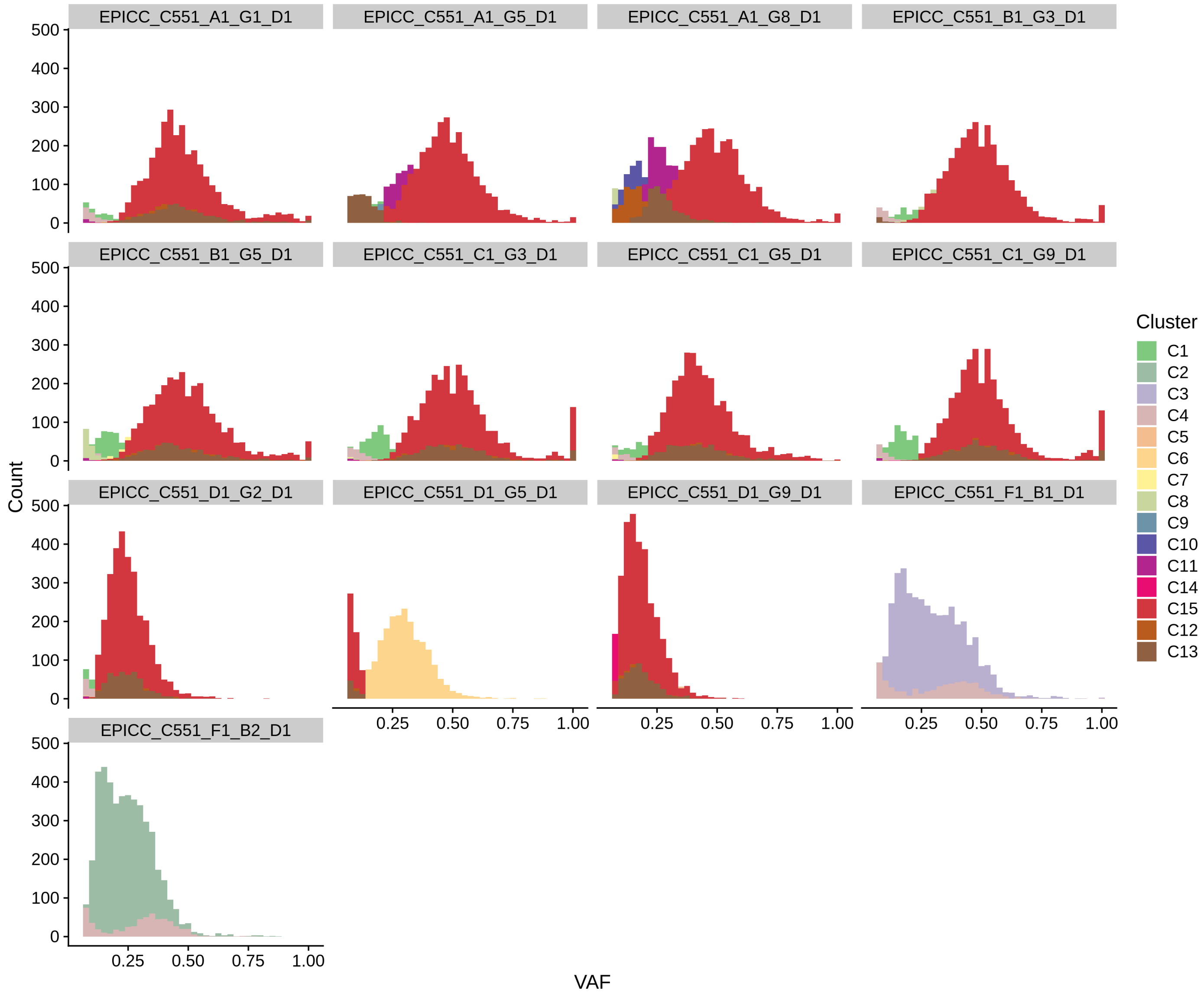
EPICC_C551_A1_G1_D1
EPICC_C551_A1_G5_D1
EPICC_C551_A1_G8_D1
EPICC_C551_B1_G3_D1
EPICC_C551_B1_G5_D1
EPICC_C551_C1_G3_D1
EPICC_C551_C1_G5_D1
EPICC_C551_C1_G9_D1
EPICC_C551_D1_G2_D1
EPICC_C551_D1_G5_D1
EPICC_C551_D1_G9_D1
EPICC_C551_F1_B1_D1
EPICC_C551_F1_B2_D1
Count
0
100
200
300
400
500
0.25
0.50
0.75
1.00
VAF
Cluster
C1
C2
C3
C4
C5
C6
C7
C8
C9
C10
C11
C14
C15
C12
C13

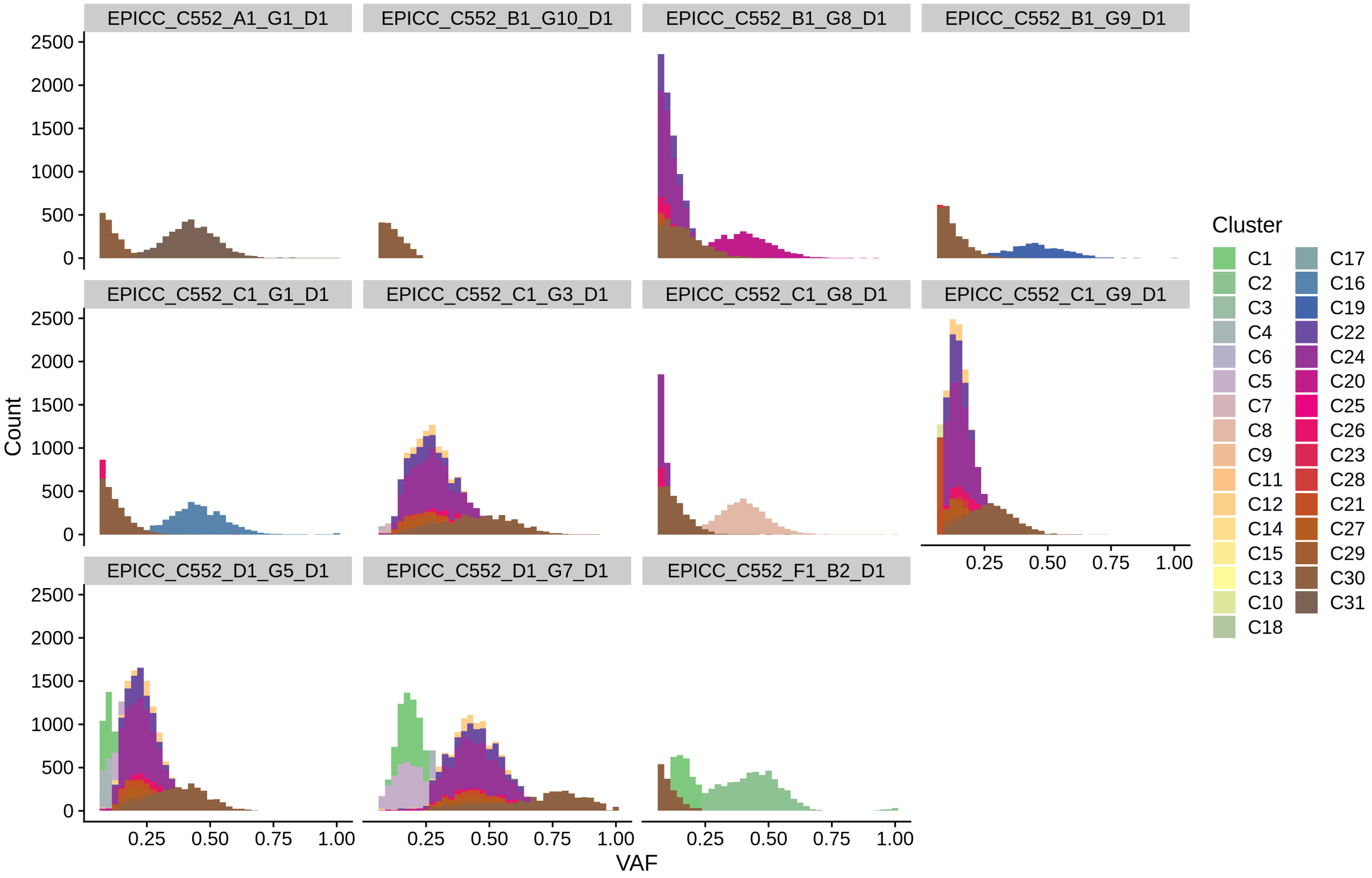

EPICC_C552_A1_G1_D1
EPICC_C552_B1_G10_D1
EPICC_C552_B1_G8_D1
EPICC_C552_B1_G9_D1
EPICC_C552_C1_G1_D1
EPICC_C552_C1_G3_D1
EPICC_C552_C1_G8_D1
EPICC_C552_C1_G9_D1
EPICC_C552_D1_G5_D1
EPICC_C552_D1_G7_D1
EPICC_C552_F1_B2_D1
Count
0
500
1000
1500
2000
2500
VAF
0.25
0.50
0.75
1.00
Cluster
C1
C2
C3
C4
C6
C5
C7
C8
C9
C11
C12
C14
C15
C13
C10
C18
C17
C16
C19
C22
C24
C20
C25
C26
C23
C28
C21
C27
C29
C30
C31

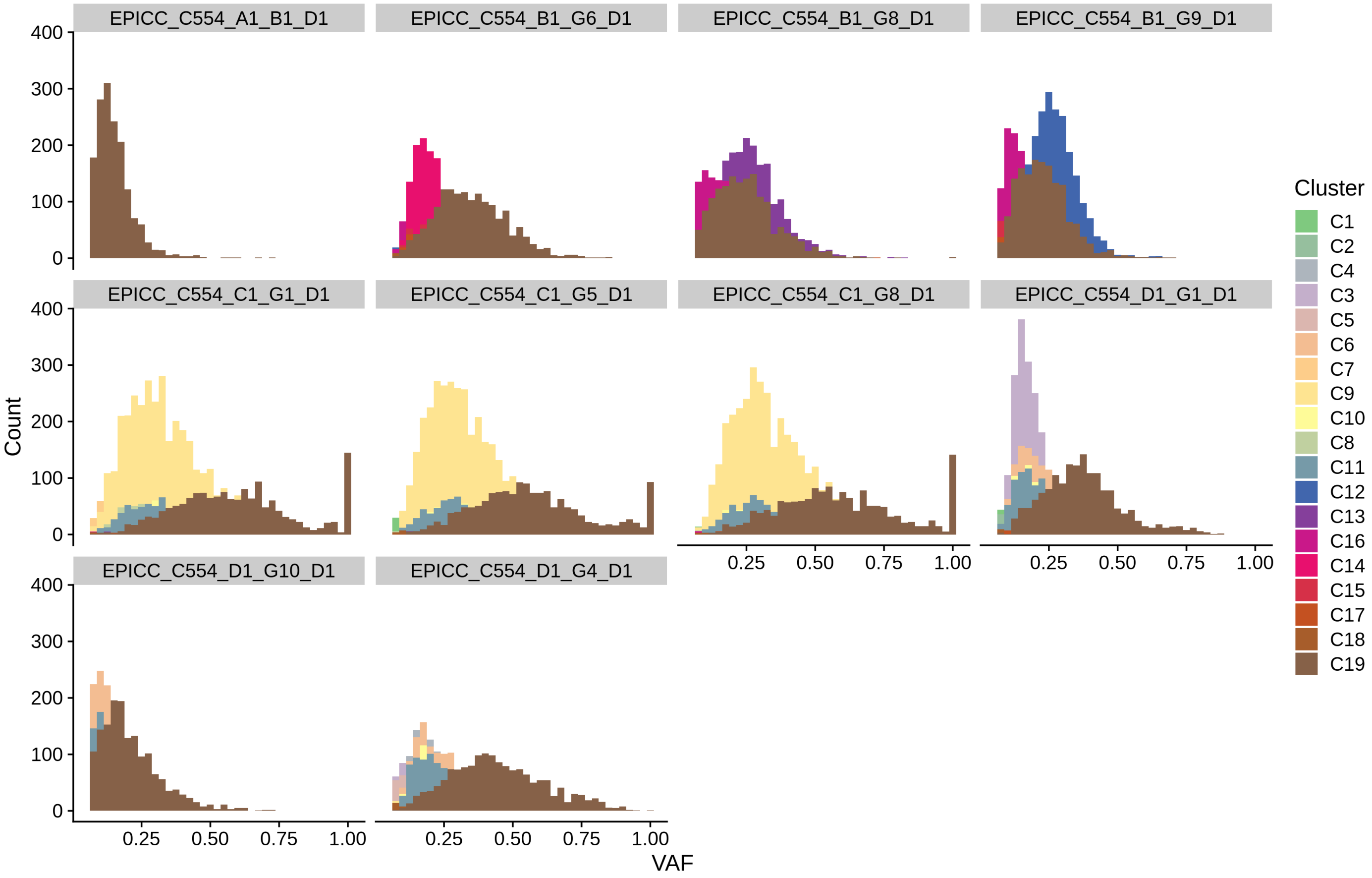

EPICC_C554_A1_B1_D1
EPICC_C554_B1_G6_D1
EPICC_C554_B1_G8_D1
EPICC_C554_B1_G9_D1
EPICC_C554_C1_G1_D1
EPICC_C554_C1_G5_D1
EPICC_C554_C1_G8_D1
EPICC_C554_D1_G1_D1
EPICC_C554_D1_G10_D1
EPICC_C554_D1_G4_D1
Count
0
100
200
300
400
VAF
0.25
0.50
0.75
1.00
Cluster
C1
C2
C4
C3
C5
C6
C7
C9
C10
C8
C11
C12
C13
C16
C14
C15
C17
C18
C19

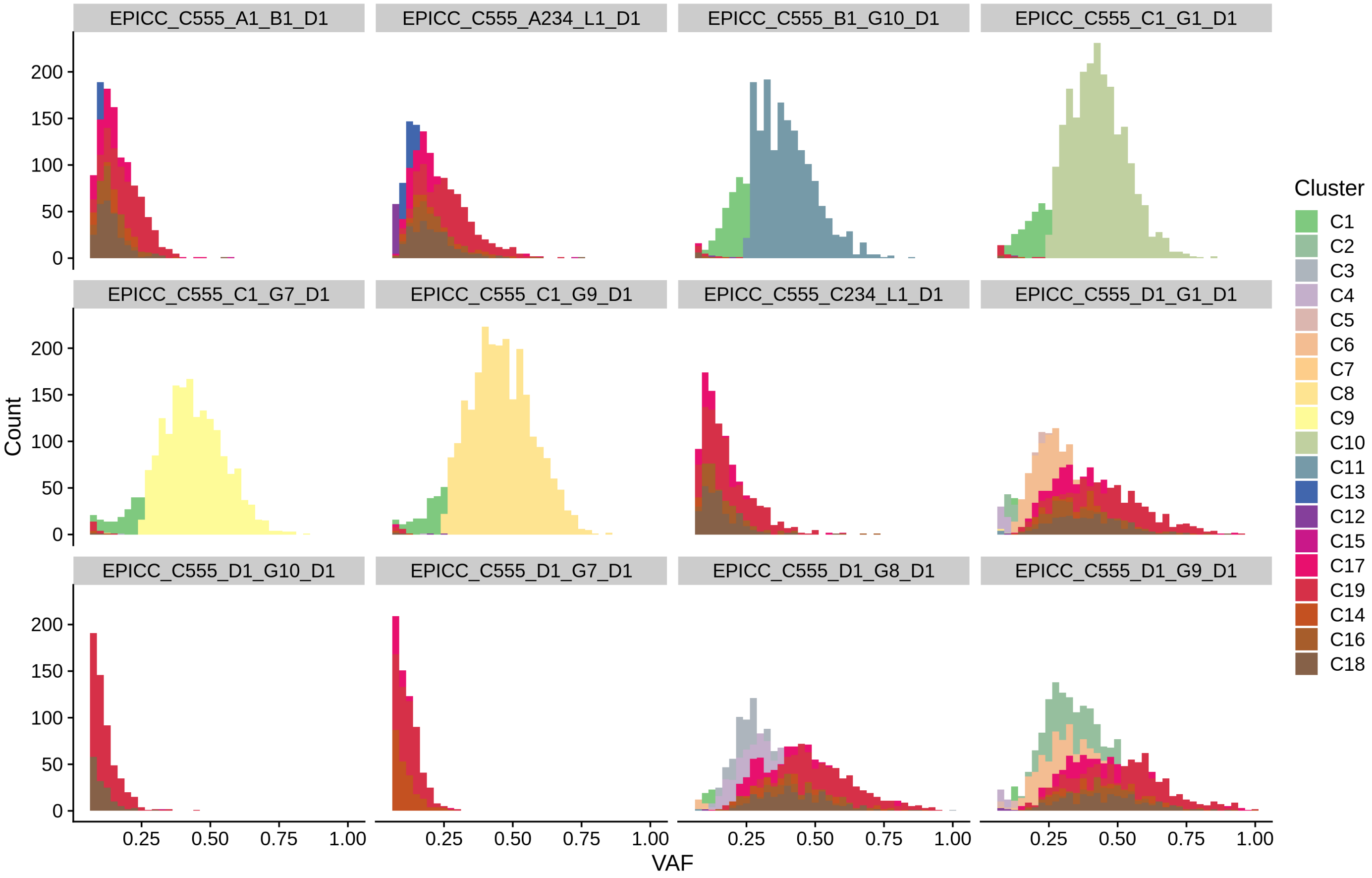

EPICC_C555_A1_B1_D1
EPICC_C555_A234_L1_D1
EPICC_C555_B1_G10_D1
EPICC_C555_C1_G1_D1
EPICC_C555_C1_G7_D1
EPICC_C555_C1_G9_D1
EPICC_C555_C234_L1_D1
EPICC_C555_D1_G1_D1
EPICC_C555_D1_G10_D1
EPICC_C555_D1_G7_D1
EPICC_C555_D1_G8_D1
EPICC_C555_D1_G9_D1
Count
0
50
100
150
200
0.25
0.50
0.75
1.00
VAF
Cluster
C1
C2
C3
C4
C5
C6
C7
C8
C9
C10
C11
C13
C12
C15
C17
C19
C14
C16
C18

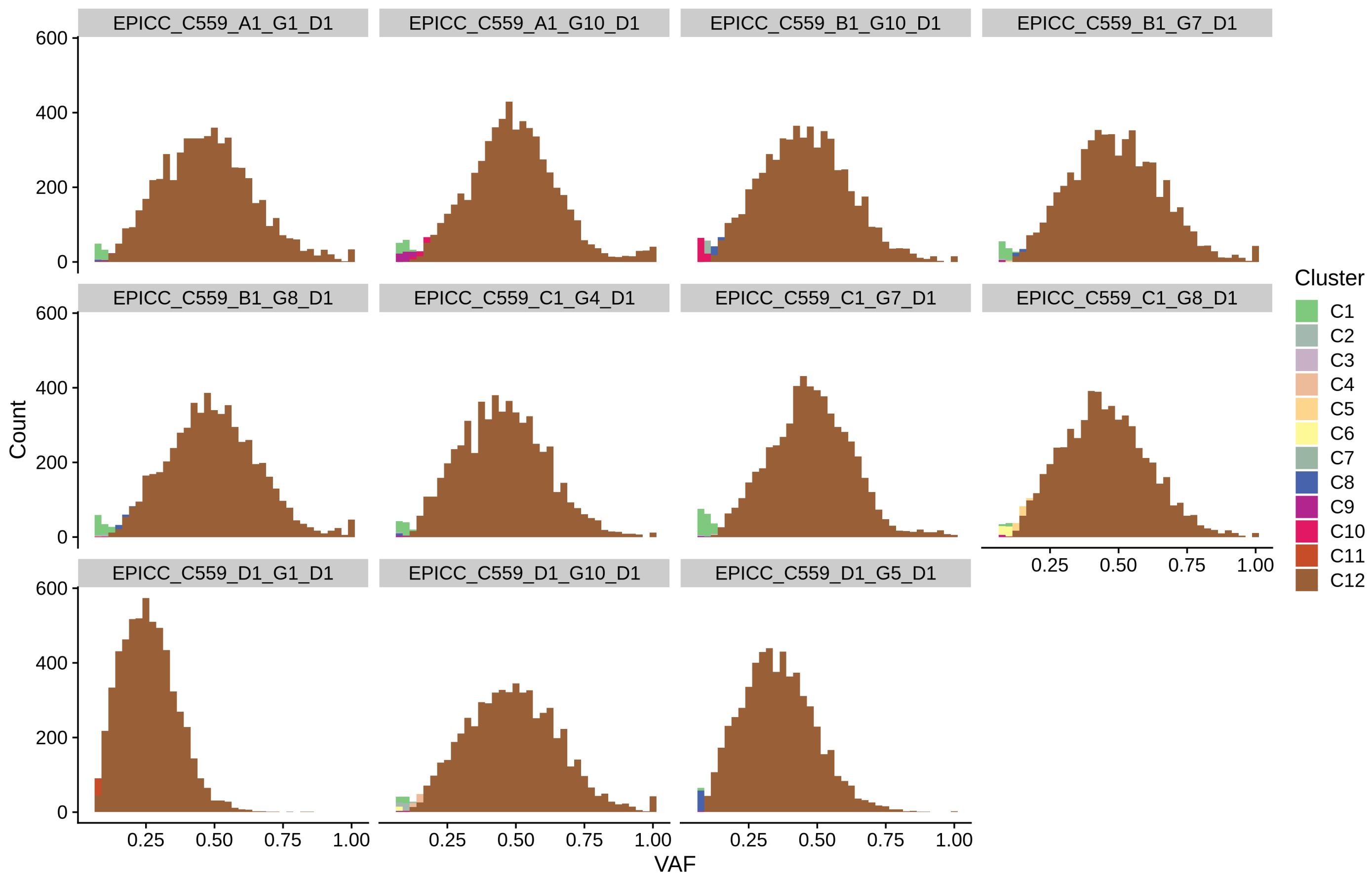
EPICC_C559_A1_G1_D1
EPICC_C559_A1_G10_D1
EPICC_C559_B1_G10_D1
EPICC_C559_B1_G7_D1
EPICC_C559_B1_G8_D1
EPICC_C559_C1_G4_D1
EPICC_C559_C1_G7_D1
EPICC_C559_C1_G8_D1
EPICC_C559_D1_G1_D1
EPICC_C559_D1_G10_D1
EPICC_C559_D1_G5_D1
Count
0
200
400
600
0.25
0.50
0.75
1.00
VAF
Cluster
C1
C2
C3
C4
C5
C6
C7
C8
C9
C10
C11
C12

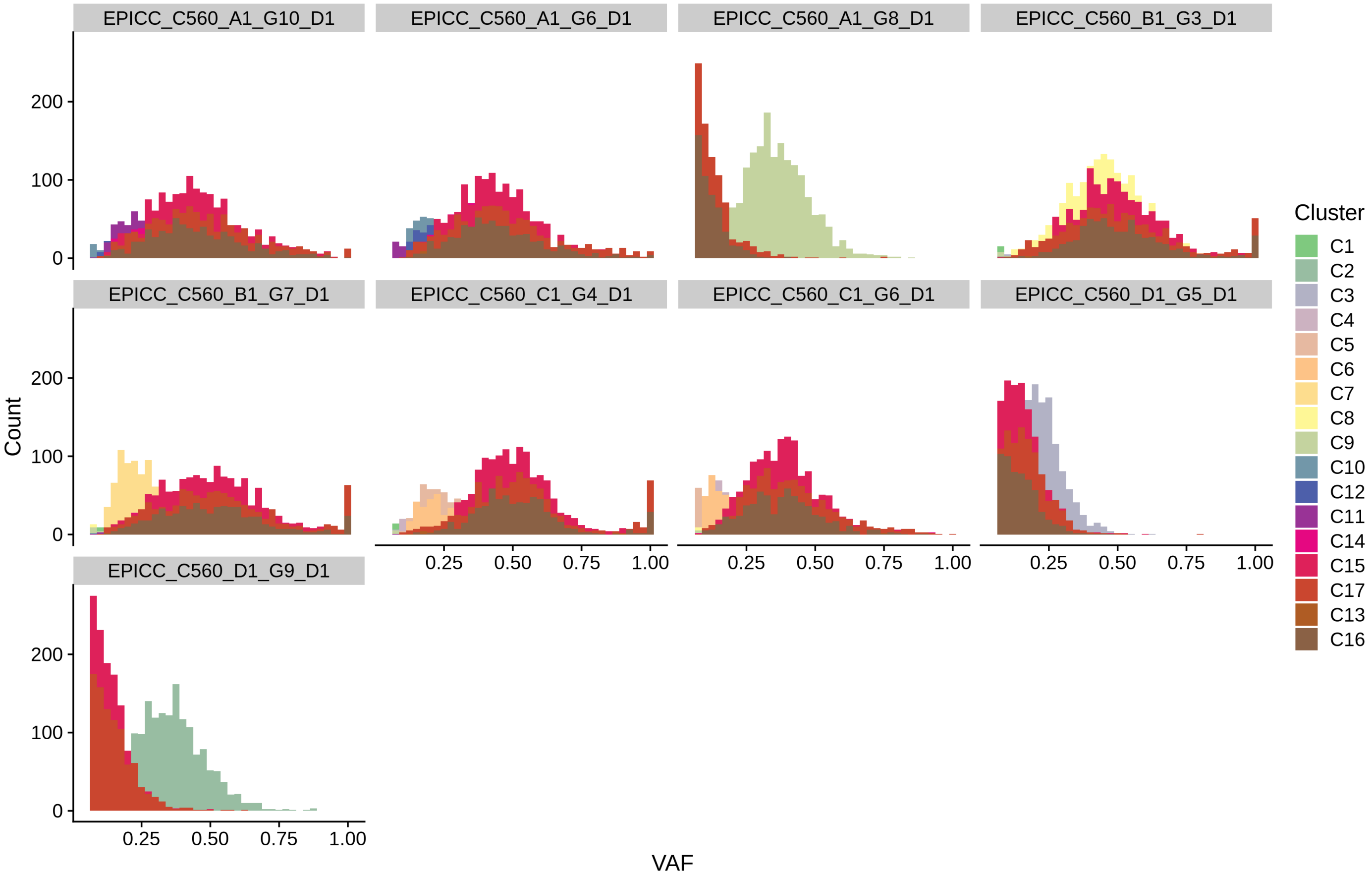

EPICC_C560_A1_G10_D1
EPICC_C560_A1_G6_D1
EPICC_C560_A1_G8_D1
EPICC_C560_B1_G3_D1
EPICC_C560_B1_G7_D1
EPICC_C560_C1_G4_D1
EPICC_C560_C1_G6_D1
EPICC_C560_D1_G5_D1
EPICC_C560_D1_G9_D1
Count
0
100
200
0.25
0.50
0.75
1.00
VAF
Cluster
C1
C2
C3
C4
C5
C6
C7
C8
C9
C10
C12
C11
C14
C15
C17
C13
C16

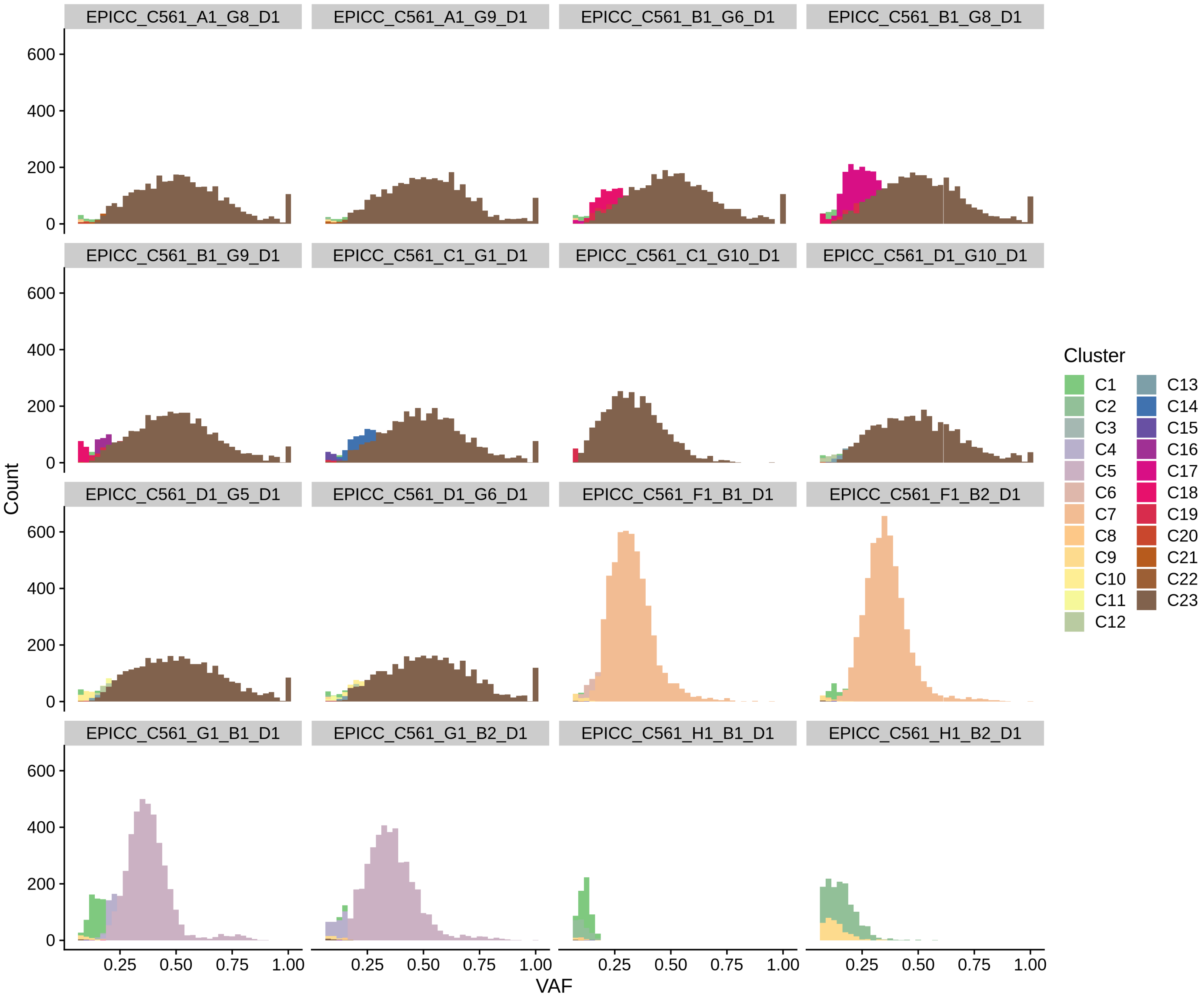
EPICC_C561_A1_G8_D1
EPICC_C561_A1_G9_D1
EPICC_C561_B1_G6_D1
EPICC_C561_B1_G8_D1
EPICC_C561_B1_G9_D1
EPICC_C561_C1_G1_D1
EPICC_C561_C1_G10_D1
EPICC_C561_D1_G10_D1
EPICC_C561_D1_G5_D1
EPICC_C561_D1_G6_D1
EPICC_C561_F1_B1_D1
EPICC_C561_F1_B2_D1
EPICC_C561_G1_B1_D1
EPICC_C561_G1_B2_D1
EPICC_C561_H1_B1_D1
EPICC_C561_H1_B2_D1
Count
0
200
400
600
0.25
0.50
0.75
1.00
VAF
Cluster
C1
C2
C3
C4
C5
C6
C7
C8
C9
C10
C11
C12
C13
C14
C15
C16
C17
C18
C19
C20
C21
C22
C23

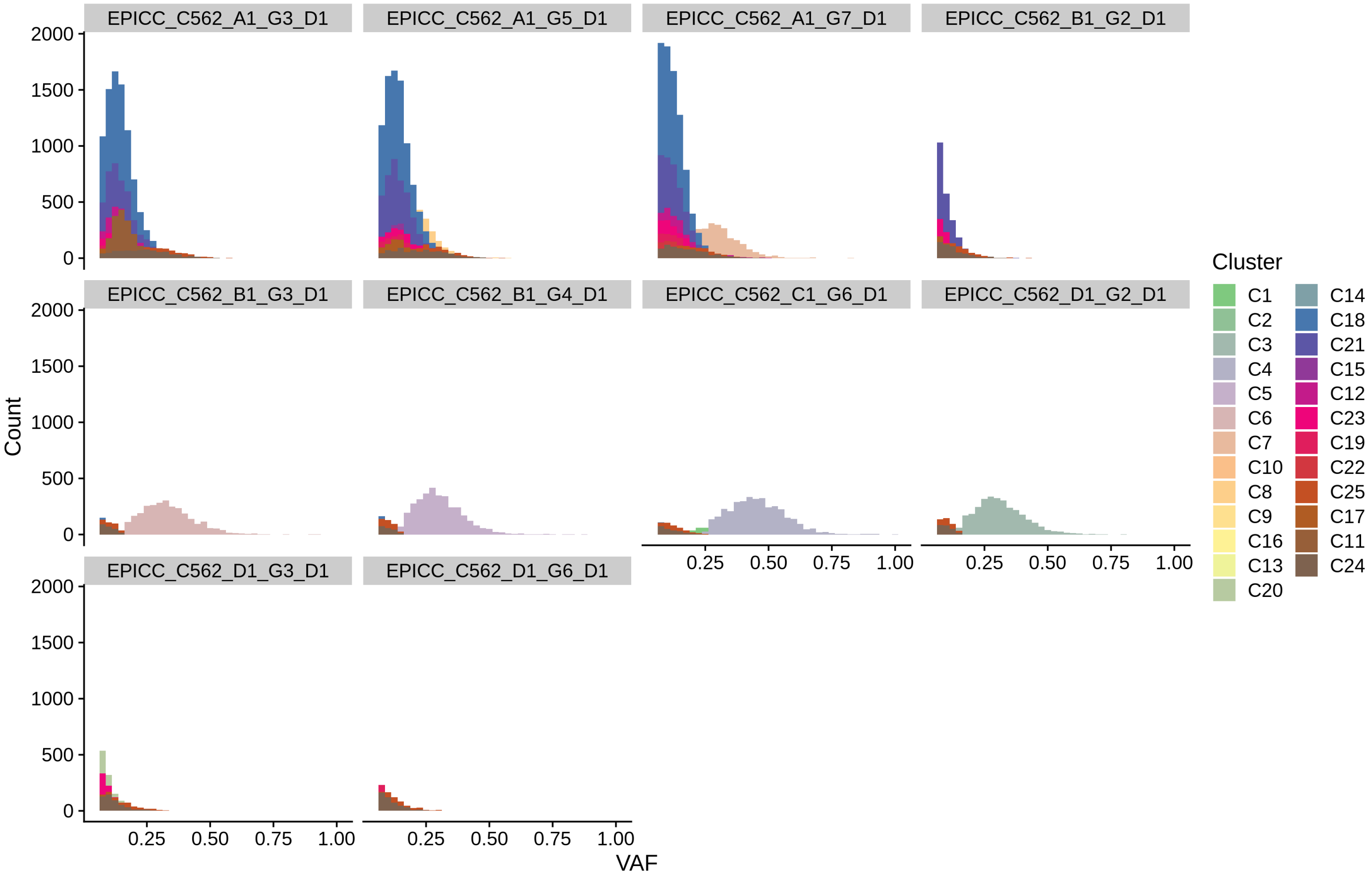
EPICC_C562_A1_G3_D1
EPICC_C562_A1_G5_D1
EPICC_C562_A1_G7_D1
EPICC_C562_B1_G2_D1
EPICC_C562_B1_G3_D1
EPICC_C562_B1_G4_D1
EPICC_C562_C1_G6_D1
EPICC_C562_D1_G2_D1
EPICC_C562_D1_G3_D1
EPICC_C562_D1_G6_D1
Count
0
500
1000
1500
2000
VAF
0.25
0.50
0.75
1.00
Cluster
C1
C2
C3
C4
C5
C6
C7
C10
C8
C9
C16
C13
C20
C14
C18
C21
C15
C12
C23
C19
C22
C25
C17
C11
C24
