## Supplementary Figures for "Assessment of the evolutionary consequence of putative driver mutations in colorectal cancer with spatial multiomic data": figS10.pdf

### EPICC inferences summary (WGS & LP)

May 4, 2021

### 1 Summary of model selection

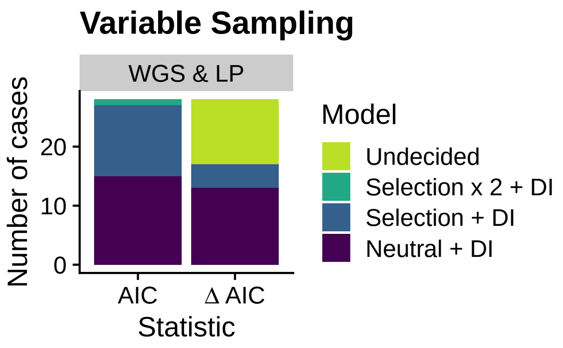

Figure 1: Number of selected models.

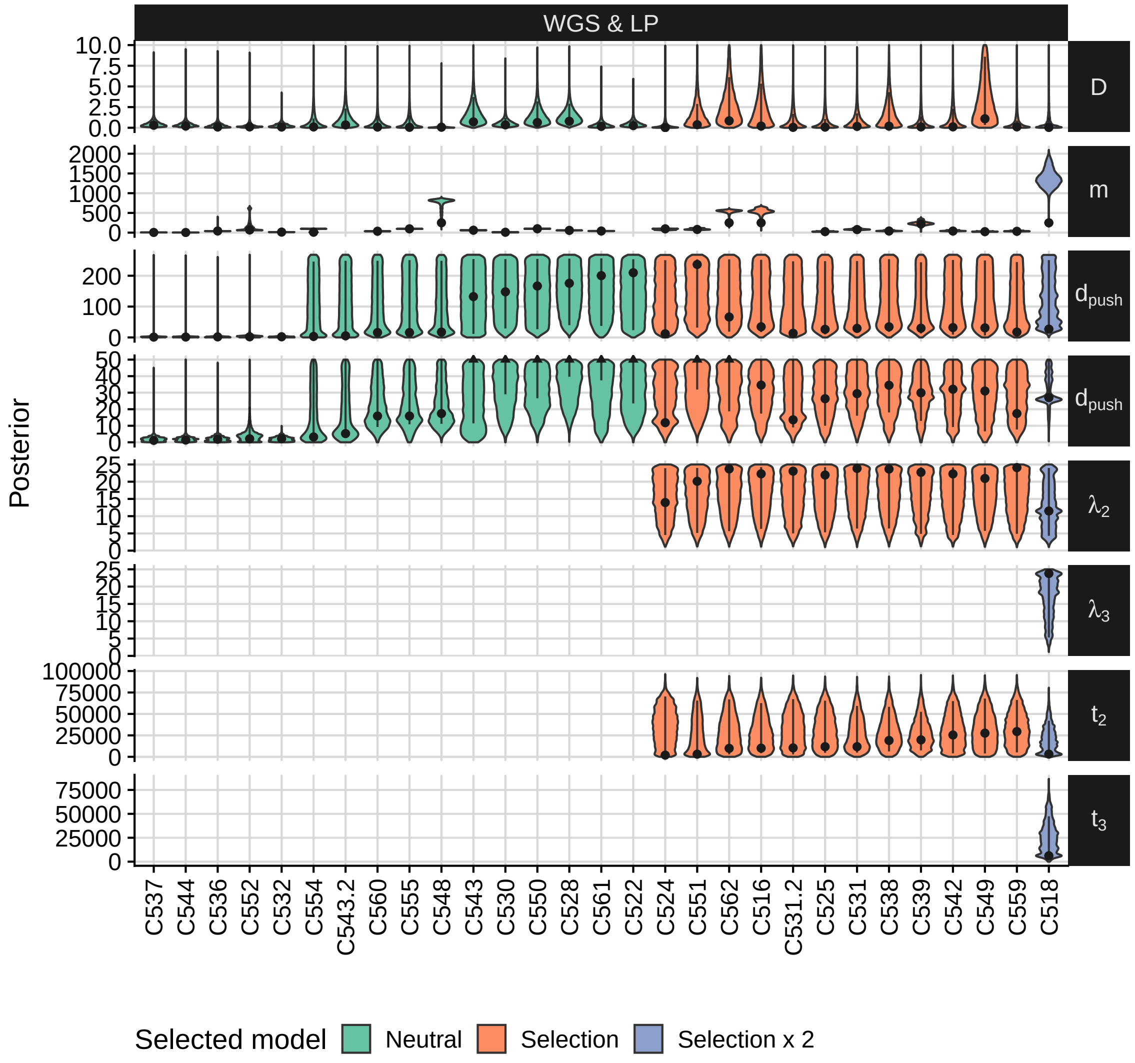

Figure 2: Posterior distribution.

Table 1: Model selection Variable Sampling - WGS + LP.

| | Case | Best Model | ΔAIC-N | ΔAIC-S | ΔAIC-Sx2 | p | Comment |
|---|---|---|---|---|---|---|---|
| 1 | C516 | **Selection** | 2.41 | 0.00 | 7.57 | 0.25 | Selection in B (Unknown driver?) |
| 2 | C518 | **Selection x 2** | 3.49 | 1.31 | 0 | 0.03 | Selection in A & B (PTEN p.C136R) |
| 5 | C524 | **Selection** | 2.1 | 0.00 | 4.79 | 0.05 | Selection in B (PIK3CA p.C378R) |
| 6 | C525 | **Selection** | **9.01** | 0.00 | 5.91 | 0.03 | Selection in C (PIK3CA p.Q546P) |
| 8 | C528 | Neutral | 0 | 2.60 | - | 0.00 | |
| 9 | C530 | Neutral | 0 | 5.05 | - | 0.01 | |
| 10 | C531 | **Selection** | 3.74 | 0.00 | 6.51 | 0.03 | Selection in D (PIK3CA p.Q546K) |
| 11 | C532 | Neutral | 0 | 10.30 | 20.6 | 0.13 | |
| 12 | C536 | Neutral | 0 | 15.90 | - | 0.59 | |
| 13 | C537 | Neutral | 0 | 6.19 | - | 0.01 | |
| 14 | C538 | **Selection** | **14.6** | 0.00 | 6.29 | 0.07 | Selection in D (RNF43 p.Q153*) |
| 15 | C539 | **Selection** | **28.6** | 0.00 | 3.05 | 0.03 | Selection in A (KRAS p.G12C) |
| 16 | C542 | **Selection** | **18.4** | 0.00 | 1.56 | 0.00 | Selection in A,B & D? (chr1p loss?) |
| 17 | C543 | Neutral | 0 | 0.88 | - | 0.00 | |
| 18 | C544 | Neutral | 0 | 9.77 | - | 0.22 | |
| 20 | C548 | Neutral | 0 | 7.28 | - | 0.19 | |
| 21 | C549 | **Selection** | 3.16 | 0.00 | 7.06 | 0.03 | Selection in A(chr1p loss?) |
| 22 | C550 | Neutral | 0 | 8.58 | - | 0.03 | |
| 23 | C551 | **Selection** | 1.59 | 0.00 | 7.52 | 0.09 | Selection in A & B (Unknown driver?) |
| 24 | C552 | Neutral | 0 | 11.30 | - | 0.50 | |
| 25 | C554 | Neutral | 0 | 4.08 | - | 0.12 | |
| 26 | C555 | Neutral | 0 | 8.16 | 16.3 | 0.55 | |
| 27 | C559 | **Selection** | **5.05** | 0.00 | - | 0.17 | Selection in A & B (Unknown driver?) |
| 28 | C560 | Neutral | 0 | 10.50 | - | 0.07 | |
| 29 | C561 | Neutral | 0 | 5.09 | - | 0.19 | |
| 30 | C562 | Selection | 0.321 | 0.00 | - | 0.12 | |
| 31 | C531 (excl. D1-G7) | **Selection** | **8.9** | 0.00 | 5.02 | 0.00 | Selection in B (SMAD4 p.A118V) |
| 32 | C543 (excl. A1-G9) | Neutral | 0 | 4.17 | 11.6 | 0.01 | |

### 2 Model selection data

## 2.1 C516

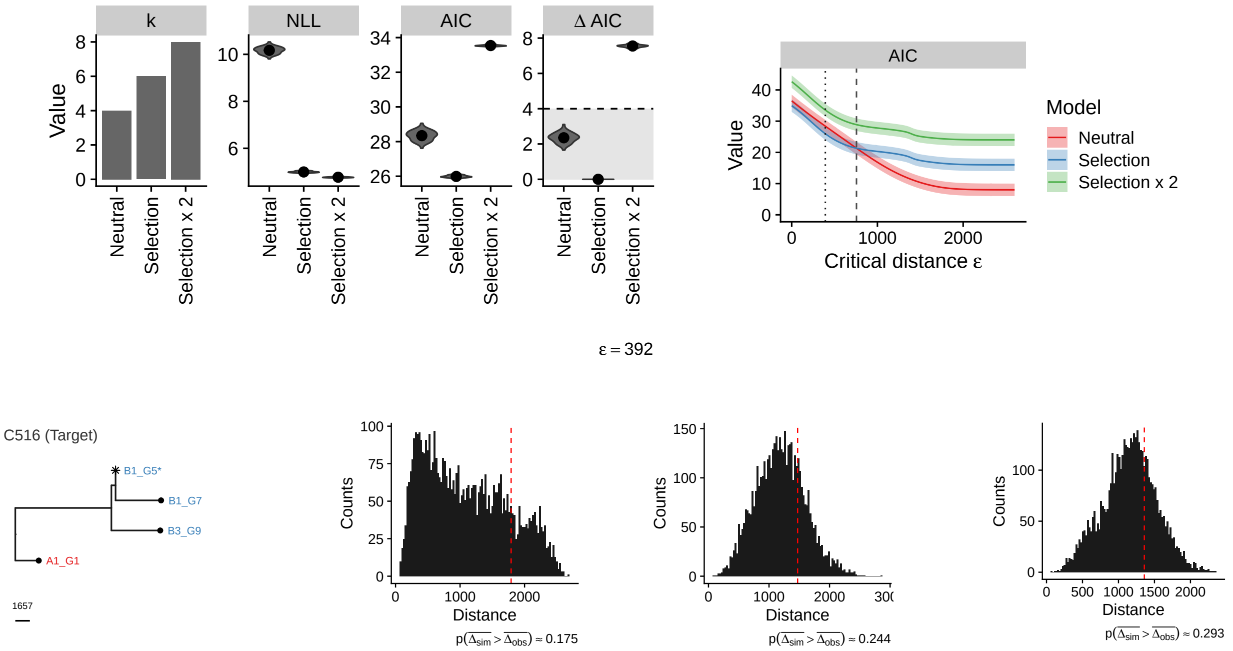

#### Neutral

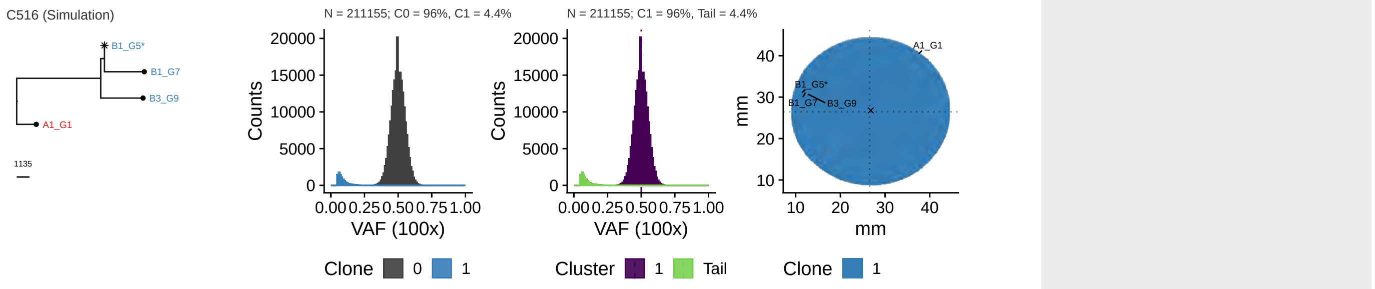

Parameters: $\lambda = 1$, $\mu = 0$, $d_{push} = 155$, $m = 0$, $t_s = 0$, $i_a = -1$

#### Selection

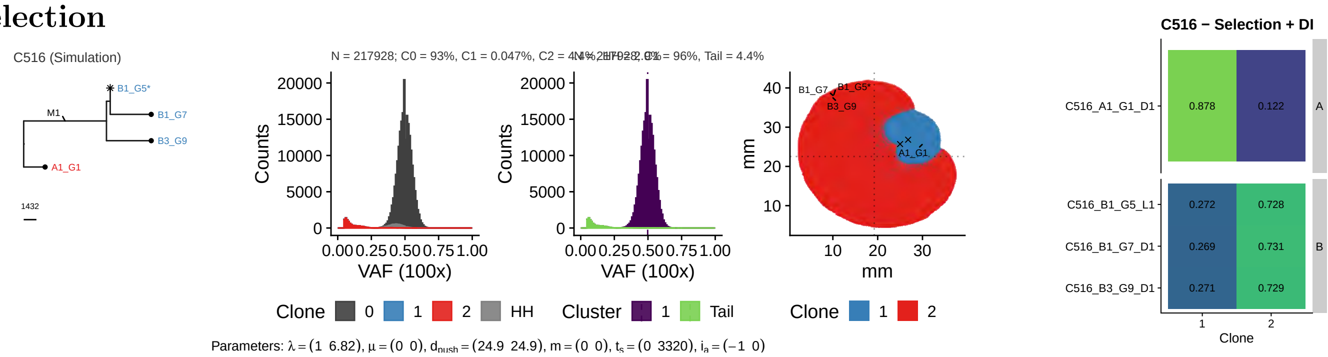

Parameters: $\lambda = (1\ \ 6.82)$, $\mu = (0\ \ 0)$, $d_{push} = (24.9\ \ 24.9)$, $m = (0\ \ 0)$, $t_s = (0\ \ 3320)$, $i_a = (-1\ \ 0)$

#### 2x Selection

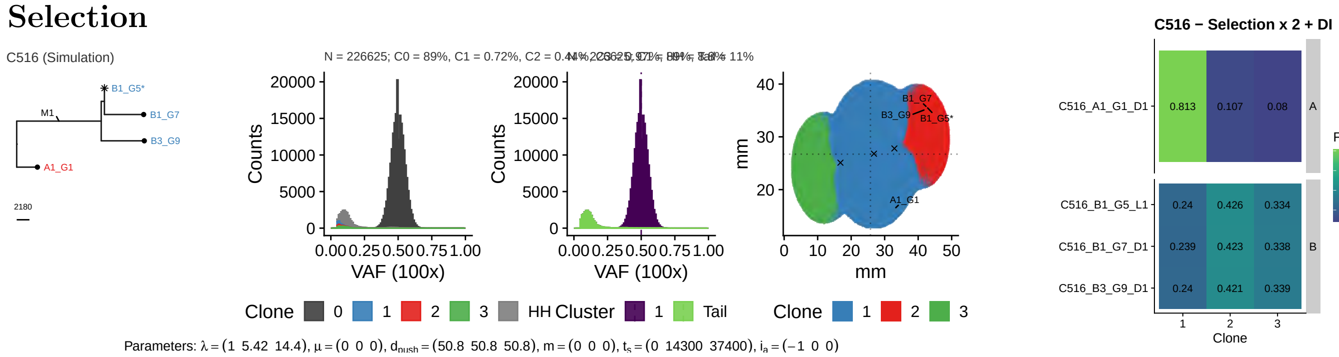

Parameters: $\lambda = (1\ \ 5.42\ \ 14.4)$, $\mu = (0\ \ 0\ \ 0)$, $d_{push} = (50.8\ \ 50.8\ \ 50.8)$, $m = (0\ \ 0\ \ 0)$, $t_s = (0\ \ 14300\ \ 37400)$, $i_a = (-1\ \ 0\ \ 0)$

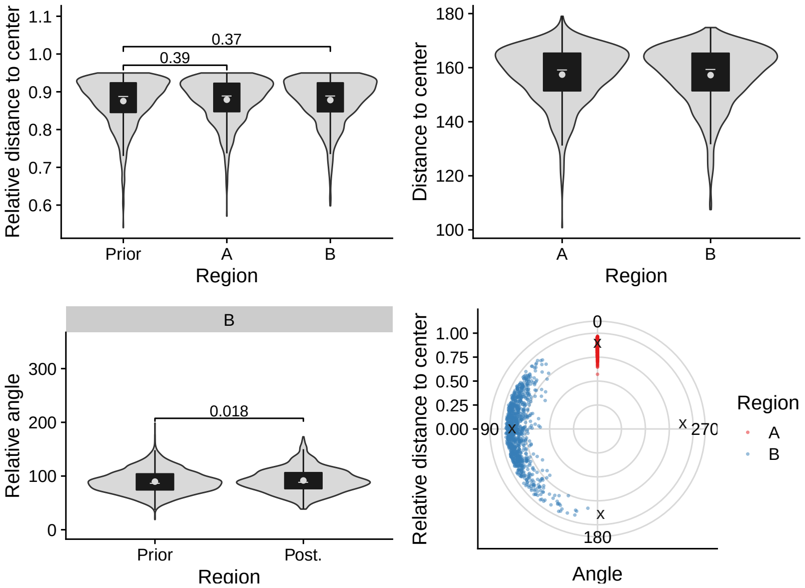
Relative distance to center
0.37
0.39
Prior
A
B
Region
Distance to center
Region
B
Relative angle
0.018
Prior
Post.
Region
Relative distance to center
0
90
270
180
Angle
Region
A
B

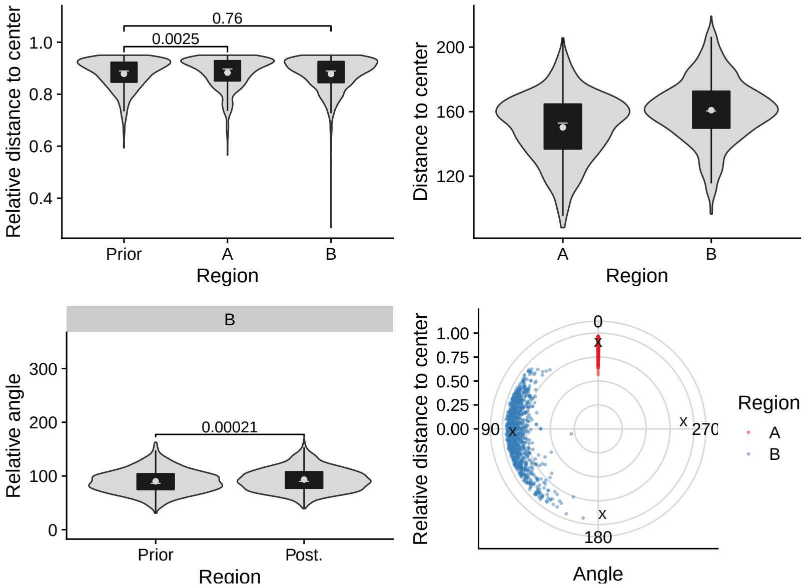
Relative distance to center
0.76
0.0025
Prior
A
B
Region
Distance to center
Region
B
Relative angle
0.00021
Prior
Post.
Region
Relative distance to center
0
90
270
180
Angle
Region
A
B

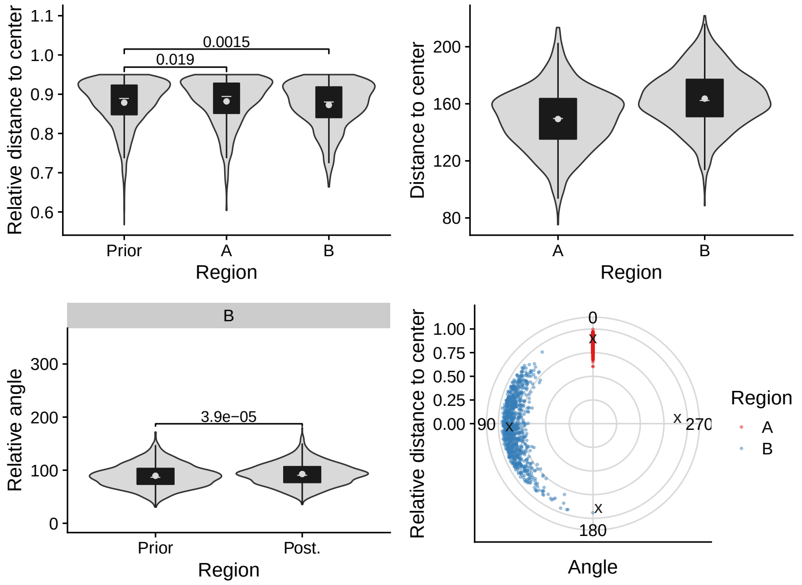
Relative distance to center
0.0015
0.019
Prior
A
B
Region
Distance to center
Region
B
Relative angle
3.9e−05
Prior
Post.
Region
Relative distance to center
0
90
270
180
Angle
Region
A
B

## 2.2 C518

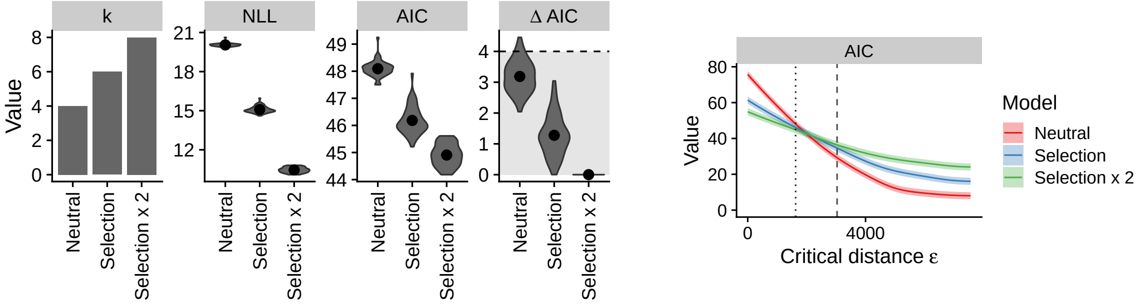

ε = 1630

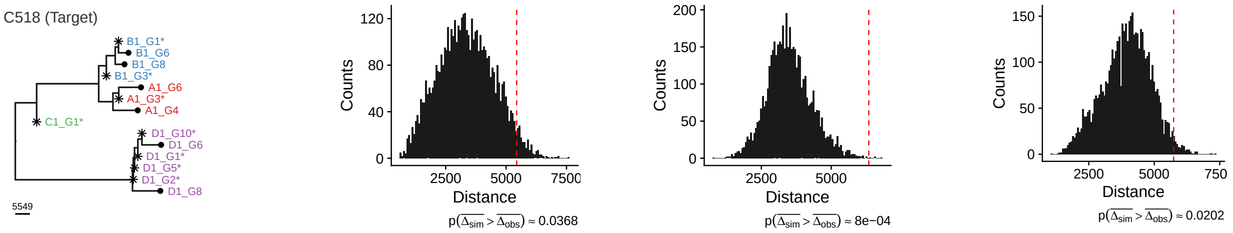

### Neutral

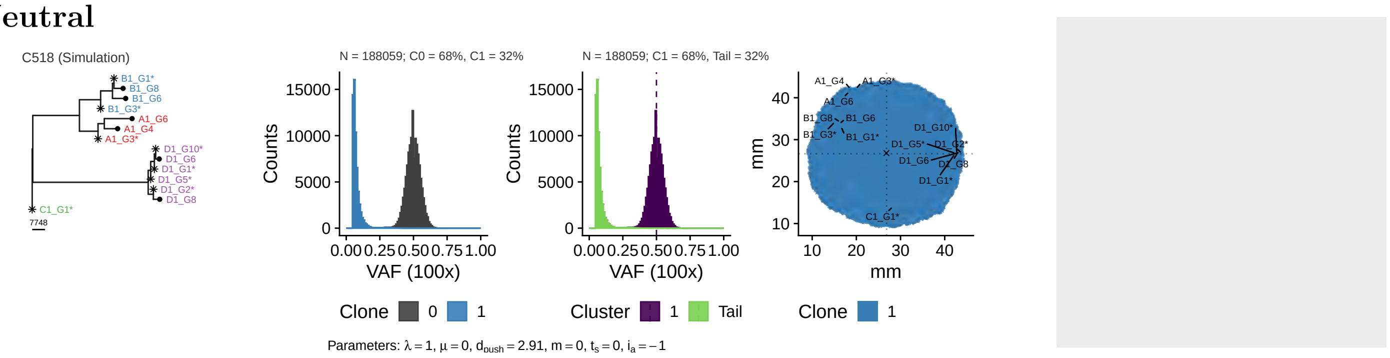

### Selection

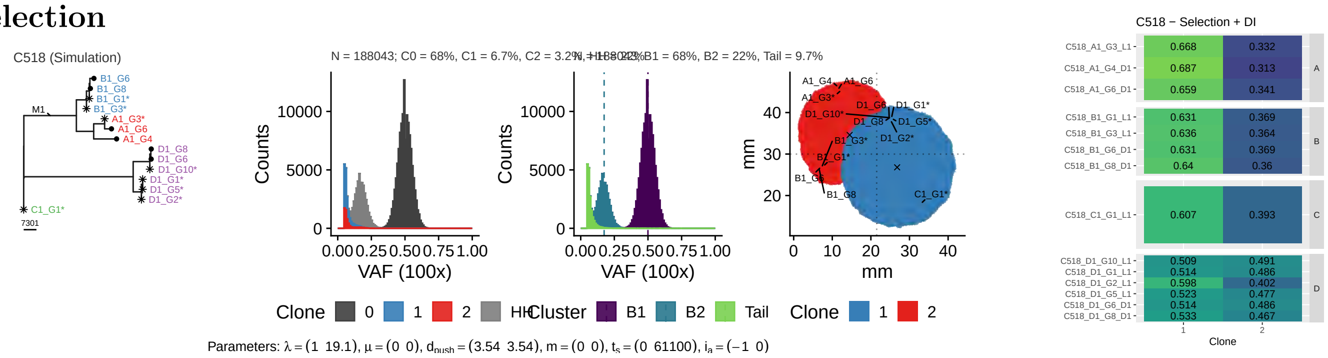

### 2x Selection

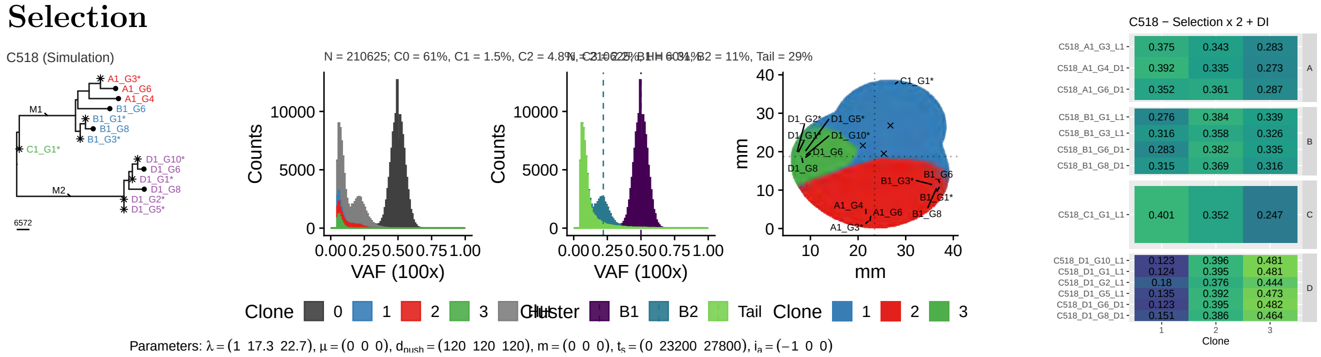

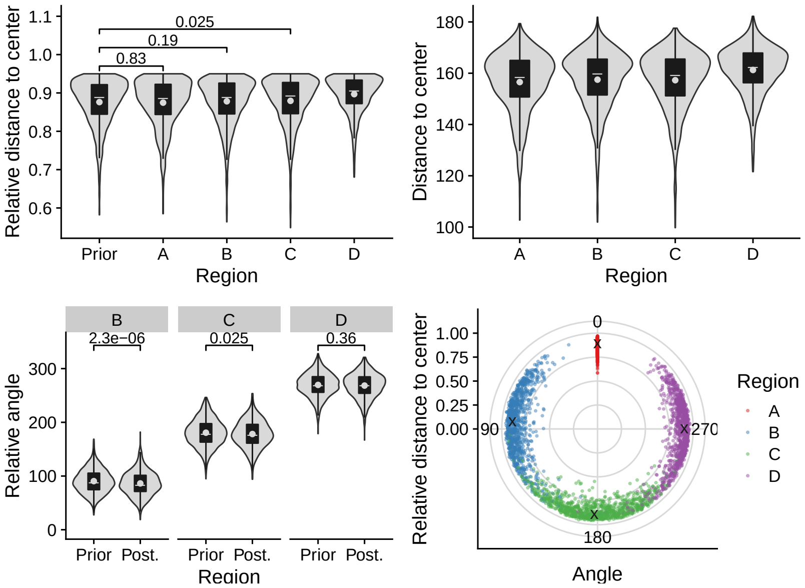
Relative distance to center
0.025
0.19
0.83
Prior
A
B
C
D
Region
Distance to center
Region
Relative angle
2.3e−06
0.025
0.36
Prior
Post.
Region
Relative distance to center
Angle
Region

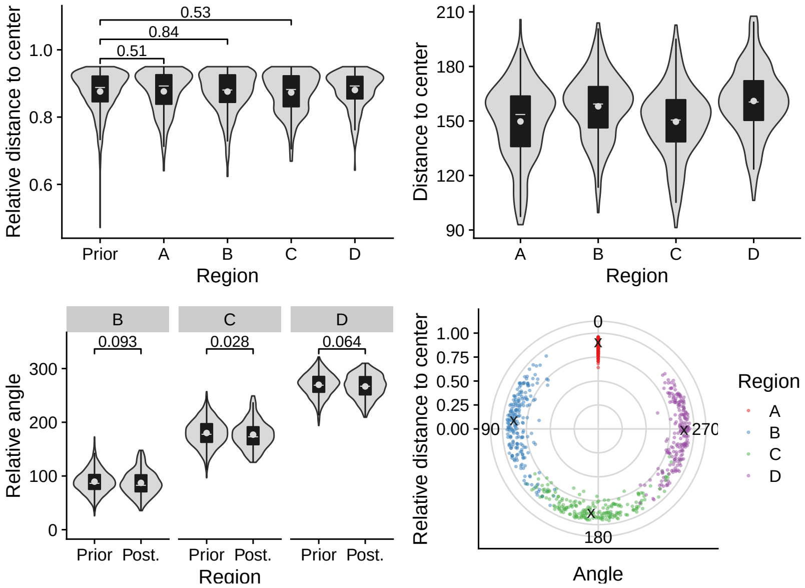
Relative distance to center
0.53
0.84
0.51
Prior
A
B
C
D
Region
Distance to center
Region
Relative angle
0.093
0.028
0.064
Prior
Post.
Region
Relative distance to center
Angle
Region

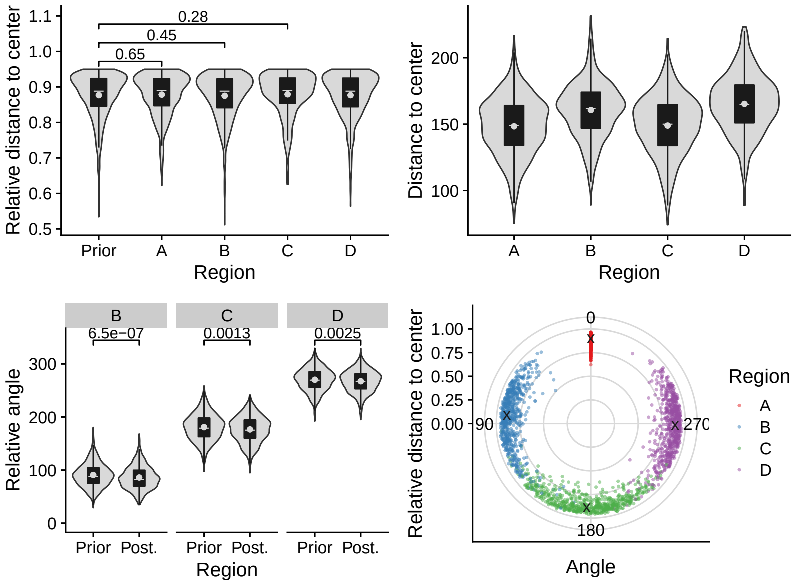
Relative distance to center
0.28
0.45
0.65
Prior
A
B
C
D
Region
Distance to center
Region
Relative angle
6.5e−07
0.0013
0.0025
Prior
Post.
Region
Relative distance to center
Angle
Region

## 2.3 C524

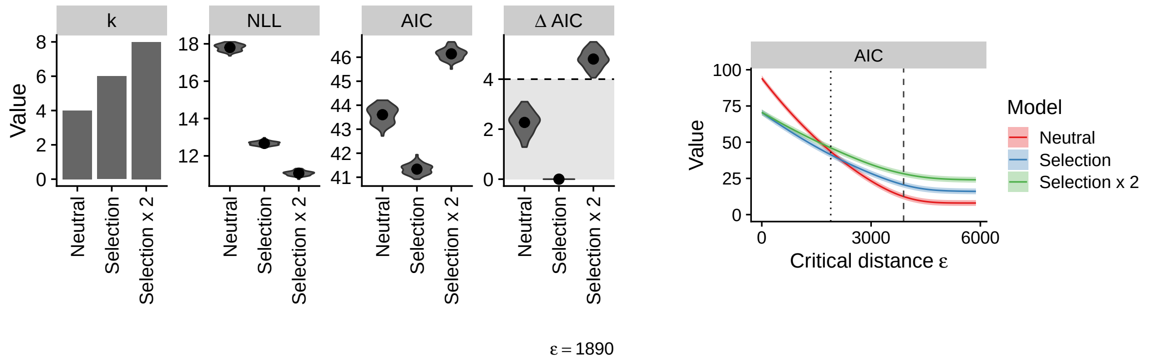

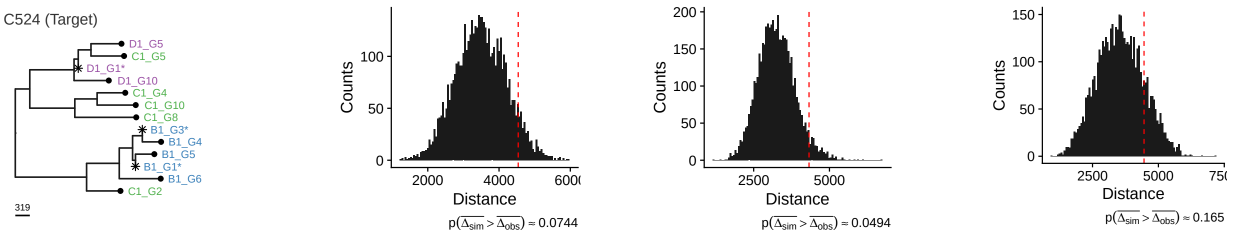

### Neutral

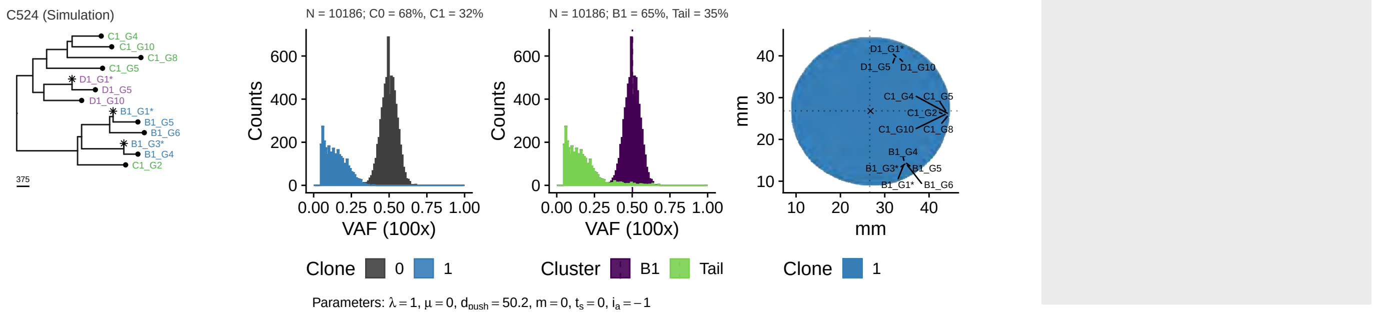

### Selection

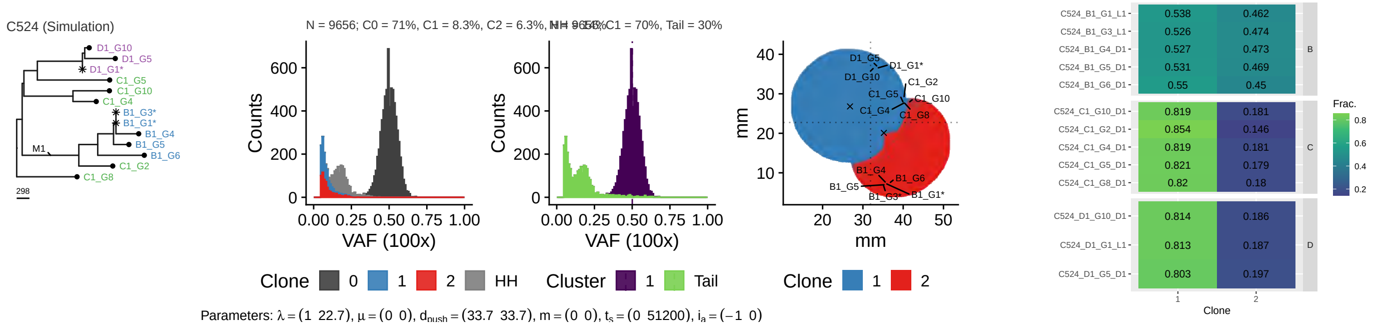

### 2x Selection

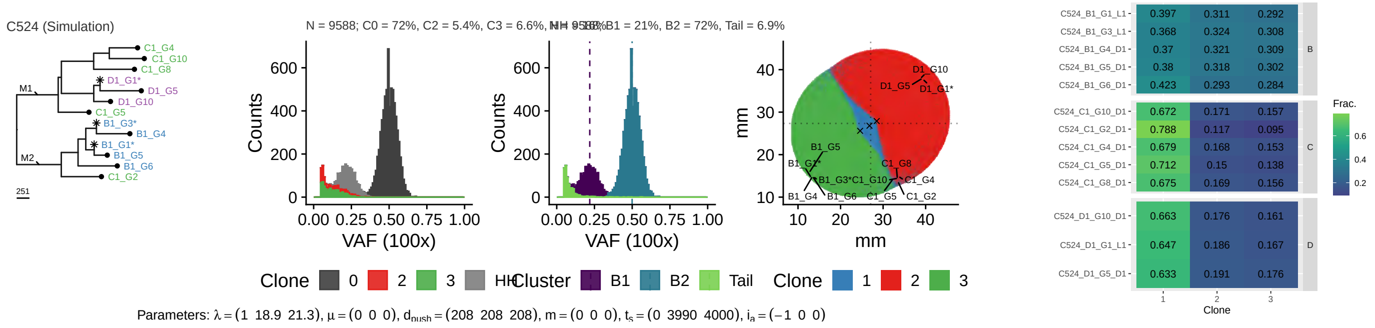

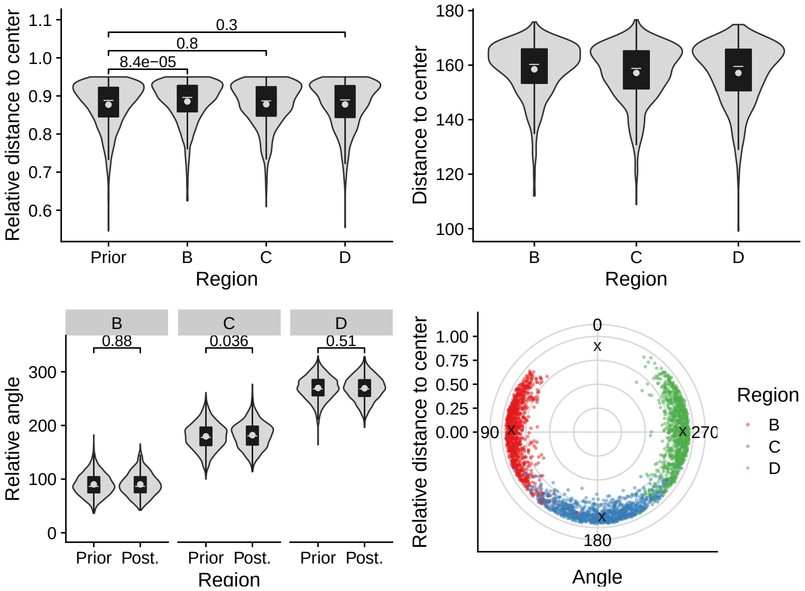

Relative distance to center
Region
Prior
B
C
D
8.4e−05
0.8
0.3
Distance to center
Relative angle
0.88
0.036
0.51
Prior Post.
Angle
Region

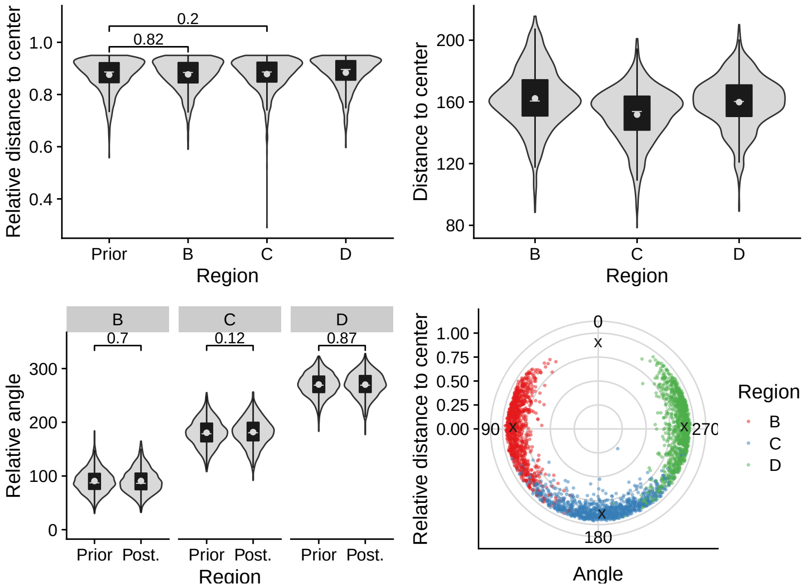

Relative distance to center
Region
Prior
B
C
D
0.82
0.2
Distance to center
Relative angle
0.7
0.12
0.87
Prior Post.
Angle
Region

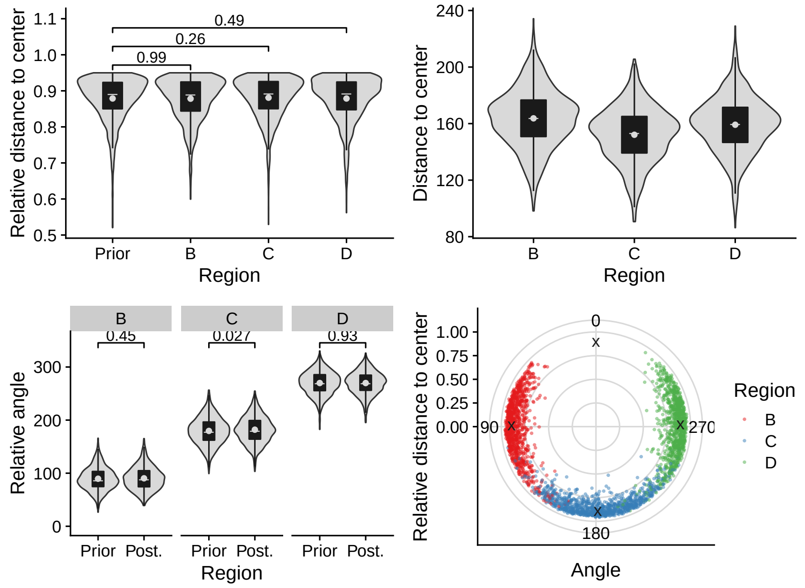

Relative distance to center
Region
Prior
B
C
D
0.99
0.26
0.49
Distance to center
Relative angle
0.45
0.027
0.93
Prior Post.
Angle
Region

## 2.4 C525

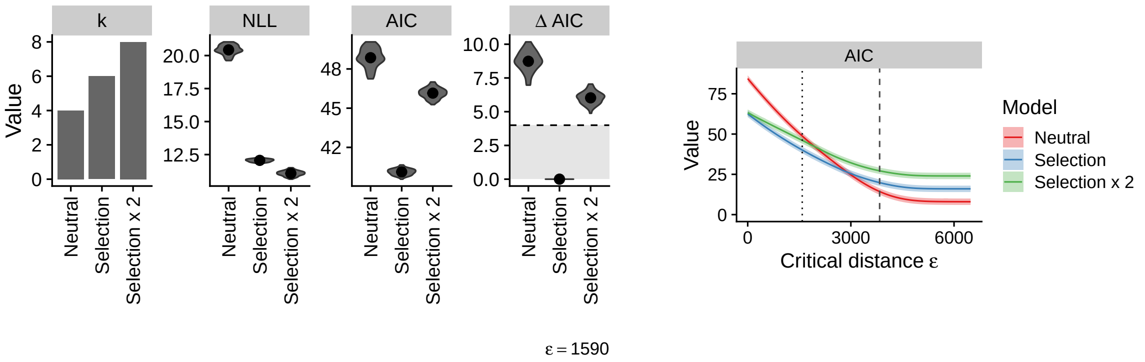

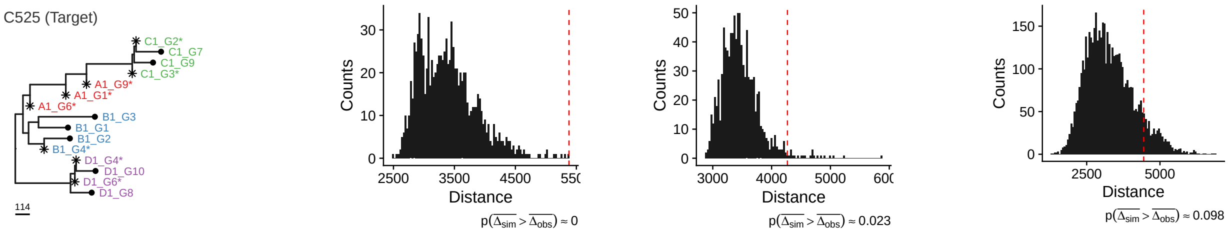

### Neutral

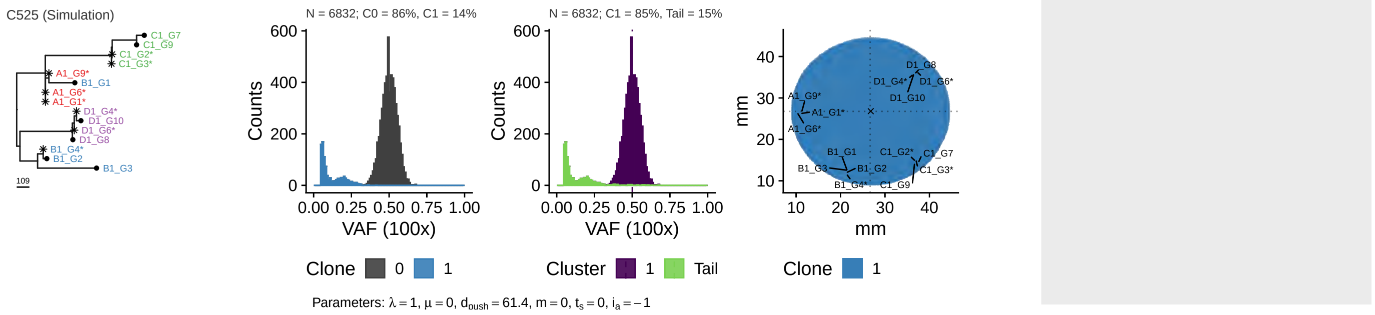

### Selection

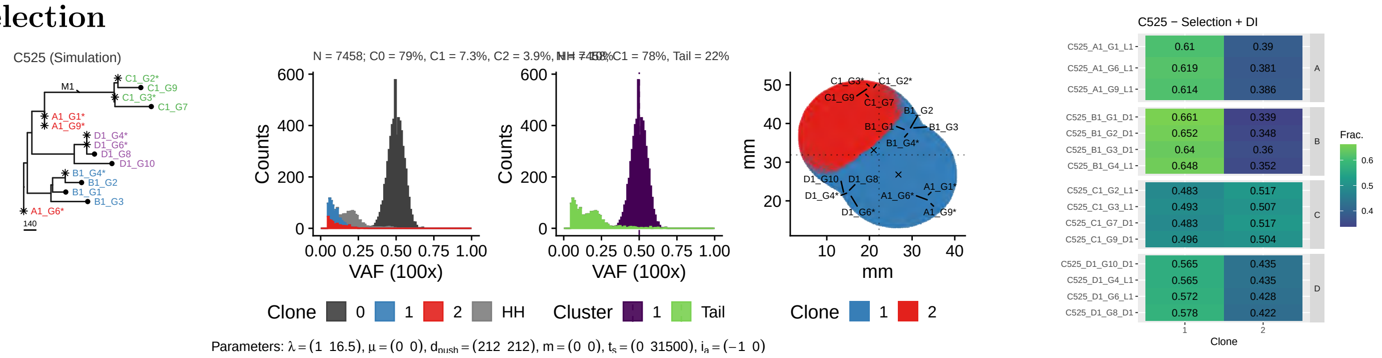

### 2x Selection

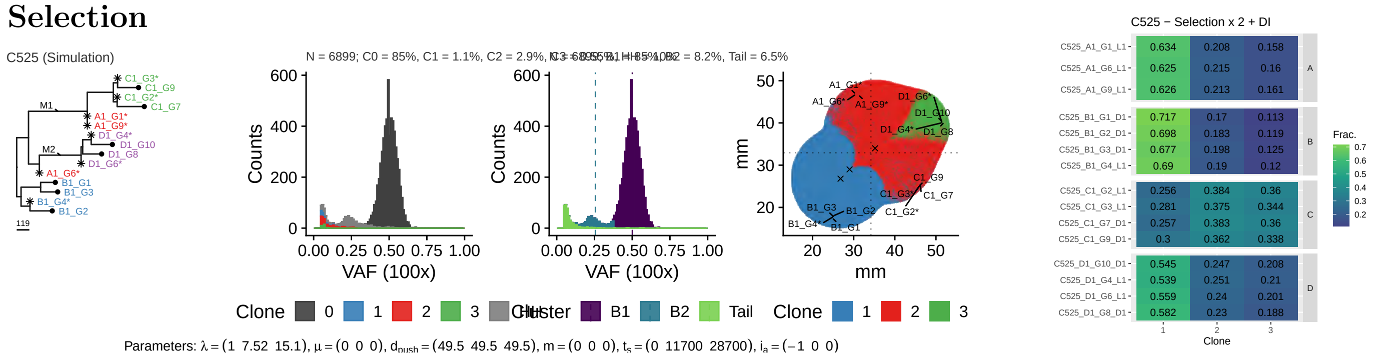

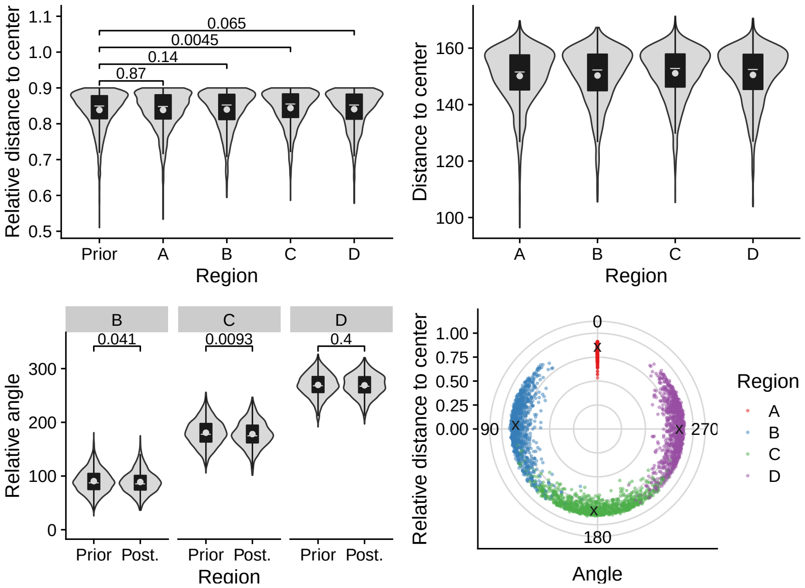
Relative distance to center
0.065
0.0045
0.14
0.87
Prior
A
B
C
D
Region
Distance to center
Relative angle
0.041
0.0093
0.4
Prior Post.
Relative distance to center
0
90
180
270
Angle
Region

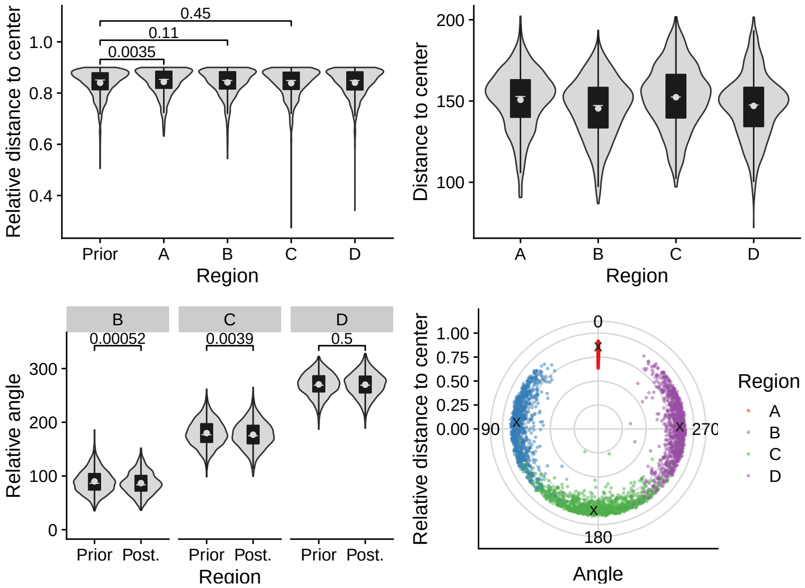
Relative distance to center
0.45
0.11
0.0035
Prior
A
B
C
D
Region
Distance to center
Relative angle
0.00052
0.0039
0.5
Prior Post.
Relative distance to center
0
90
180
270
Angle
Region

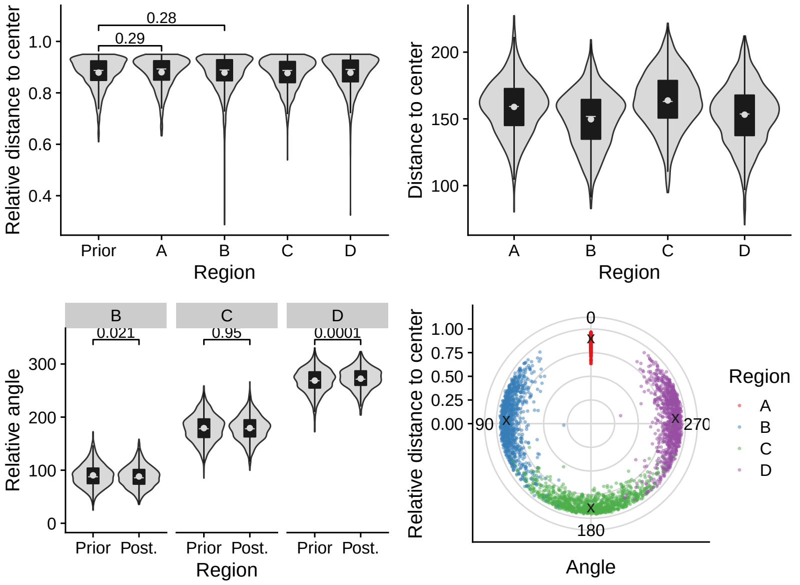
Relative distance to center
0.28
0.29
Prior
A
B
C
D
Region
Distance to center
Relative angle
0.021
0.95
0.0001
Prior Post.
Relative distance to center
0
90
180
270
Angle
Region

# 2.5 C528

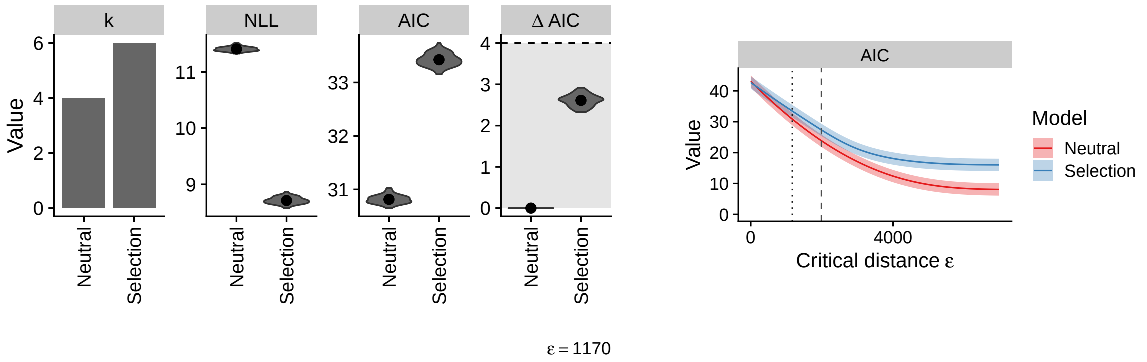

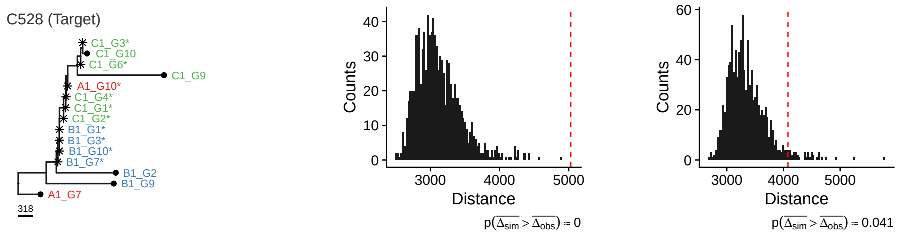

### Neutral

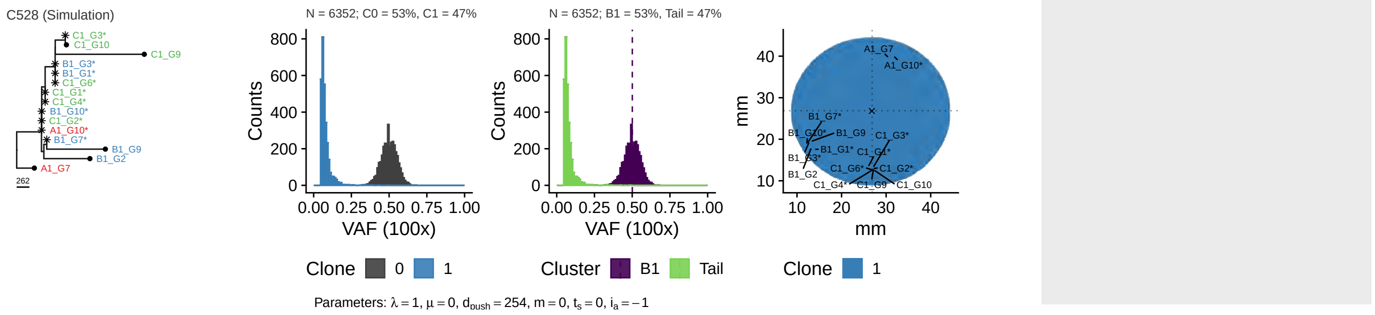

### Selection

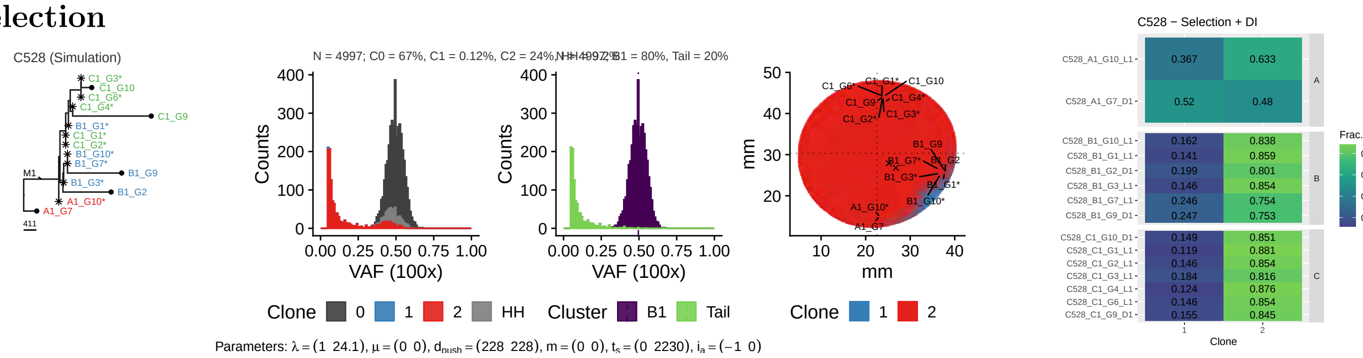

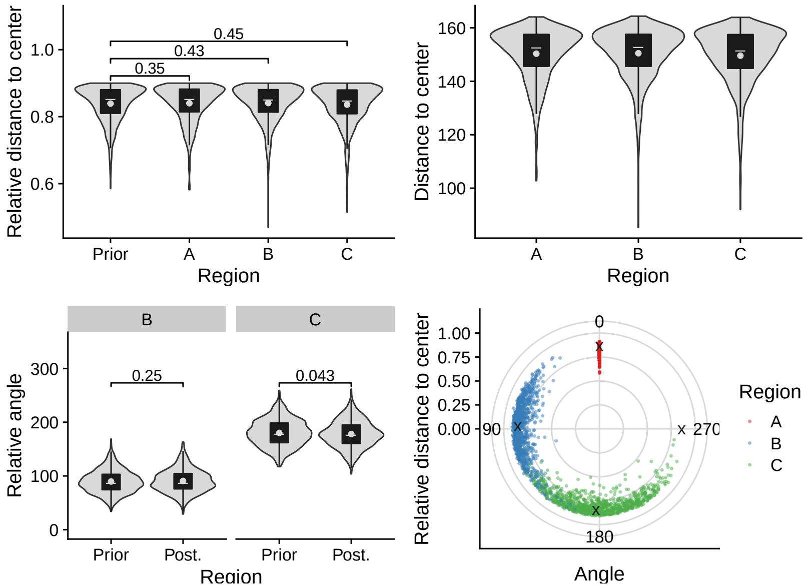
Relative distance to center
1.0
0.8
0.6
0.45
0.43
0.35
Prior
A
B
C
Region
Distance to center
160
140
120
100
A
B
C
Region
B
C
Relative angle
300
200
100
0
0.25
0.043
Prior
Post.
Prior
Post.
Region
Relative distance to center
1.00
0.75
0.50
0.25
0.00
0
90
270
180
Angle
Region
A
B
C

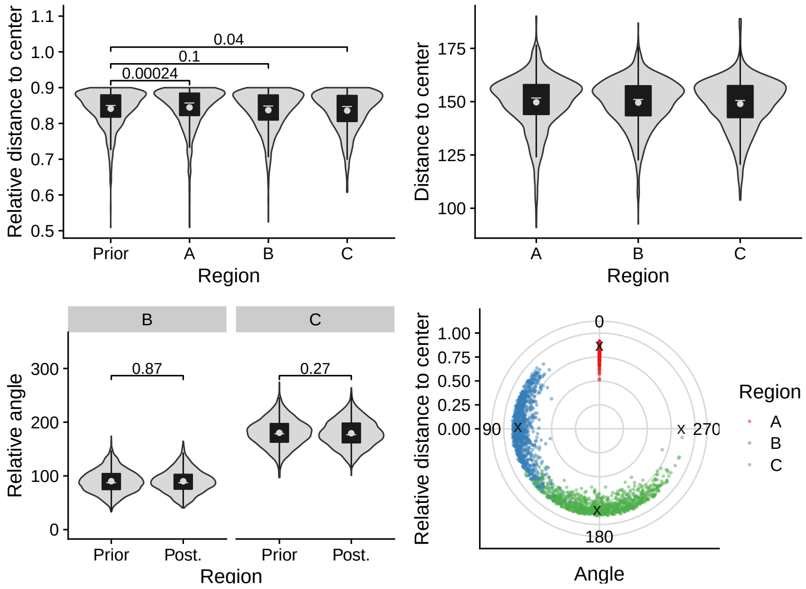
Relative distance to center
1.1
1.0
0.9
0.8
0.7
0.6
0.5
0.04
0.1
0.00024
Prior
A
B
C
Region
Distance to center
175
150
125
100
A
B
C
Region
B
C
Relative angle
300
200
100
0
0.87
0.27
Prior
Post.
Prior
Post.
Region
Relative distance to center
1.00
0.75
0.50
0.25
0.00
0
90
270
180
Angle
Region
A
B
C

## 2.6 C530

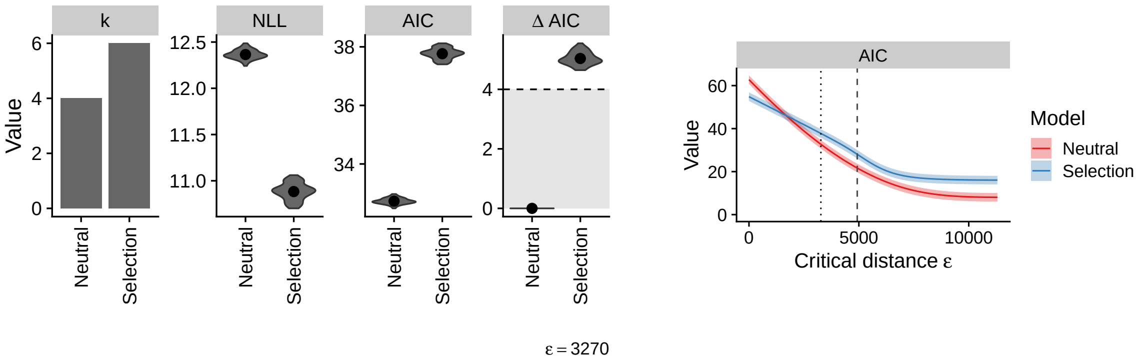

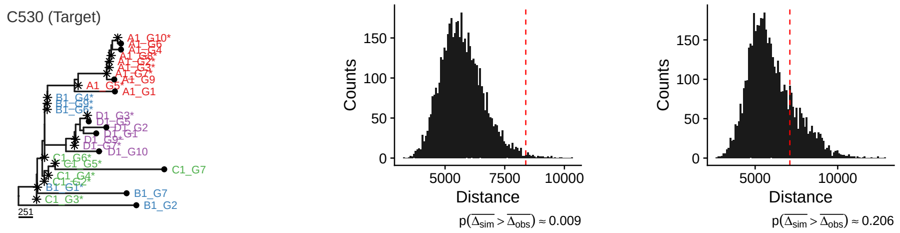

#### Neutral

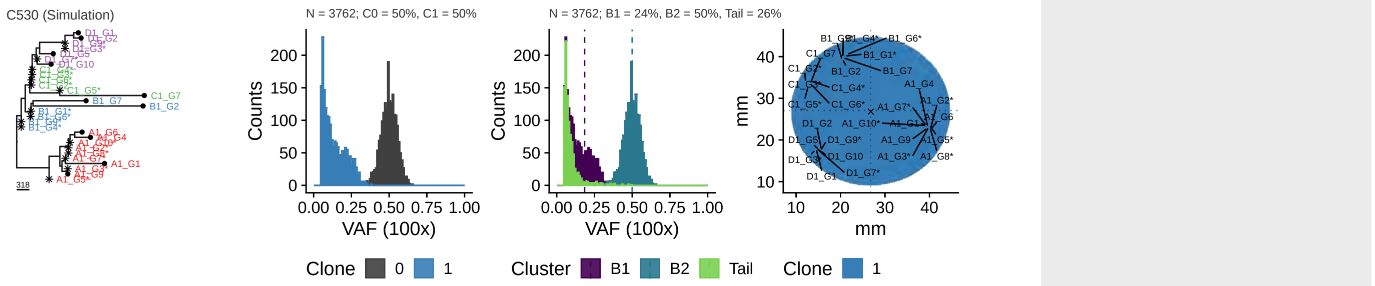

Parameters: $\lambda = 1$, $\mu = 0$, $d_{push} = 169$, $m = 0$, $t_s = 0$, $i_a = -1$

#### Selection

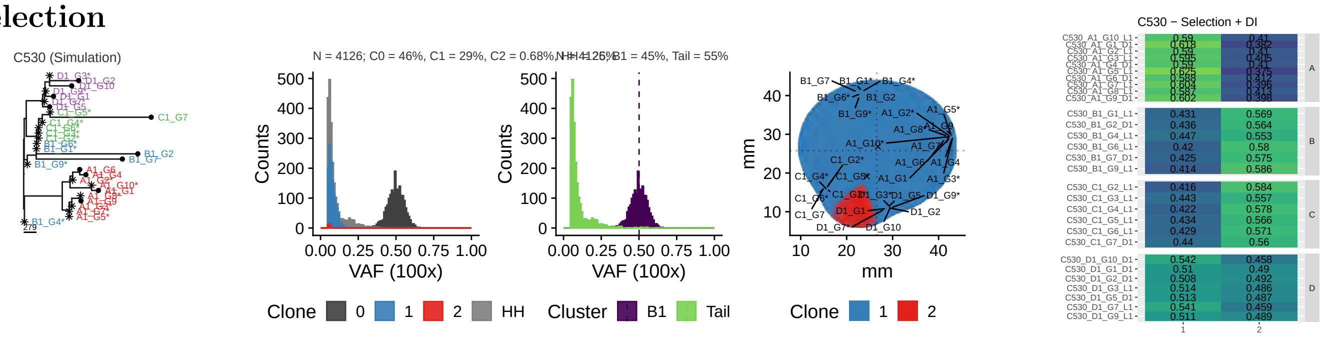

Parameters: $\lambda = (1\ \ 6.86)$, $\mu = (0\ \ 0)$, $d_{push} = (103\ \ 103)$, $m = (0\ \ 0)$, $t_s = (0\ \ 29300)$, $i_a = (-1\ \ 0)$

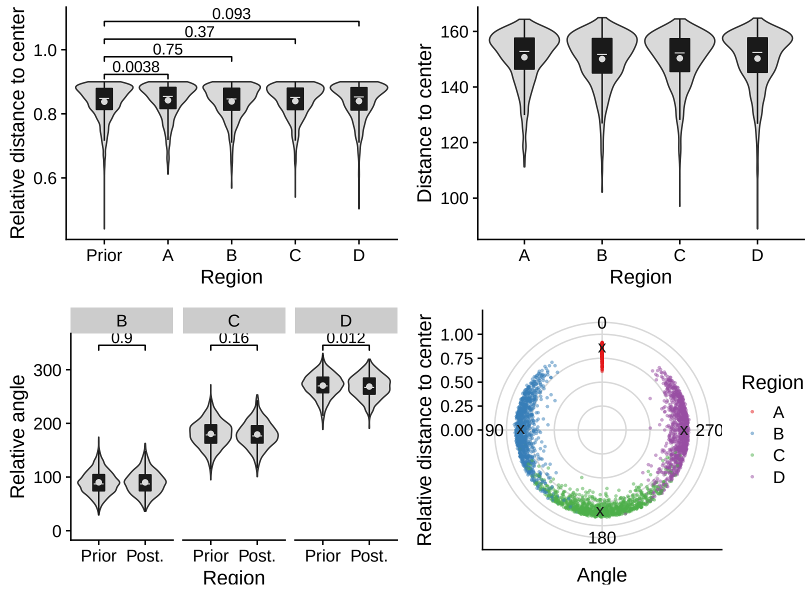
Relative distance to center
1.0
0.8
0.6
0.093
0.37
0.75
0.0038
Prior
A
B
C
D
Region
Distance to center
160
140
120
100
A
B
C
D
Region
B
C
D
0.9
0.16
0.012
Relative angle
300
200
100
0
Prior Post.
Prior Post.
Prior Post.
Region
Relative distance to center
1.00
0.75
0.50
0.25
0.00
0
90
270
180
Angle
Region
A
B
C
D

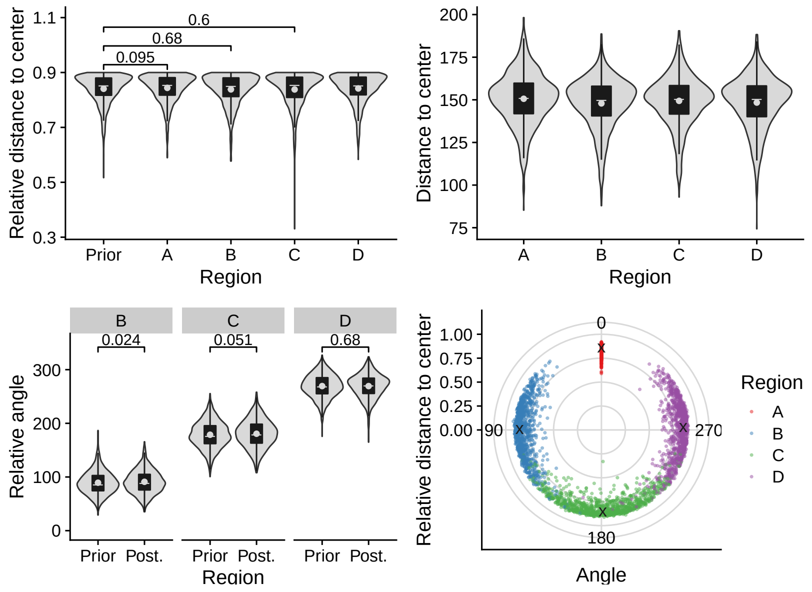
Relative distance to center
1.1
0.9
0.7
0.5
0.3
0.6
0.68
0.095
Prior
A
B
C
D
Region
Distance to center
200
175
150
125
100
75
A
B
C
D
Region
B
C
D
0.024
0.051
0.68
Relative angle
300
200
100
0
Prior Post.
Prior Post.
Prior Post.
Region
Relative distance to center
1.00
0.75
0.50
0.25
0.00
0
90
270
180
Angle
Region
A
B
C
D

## 2.7 C531

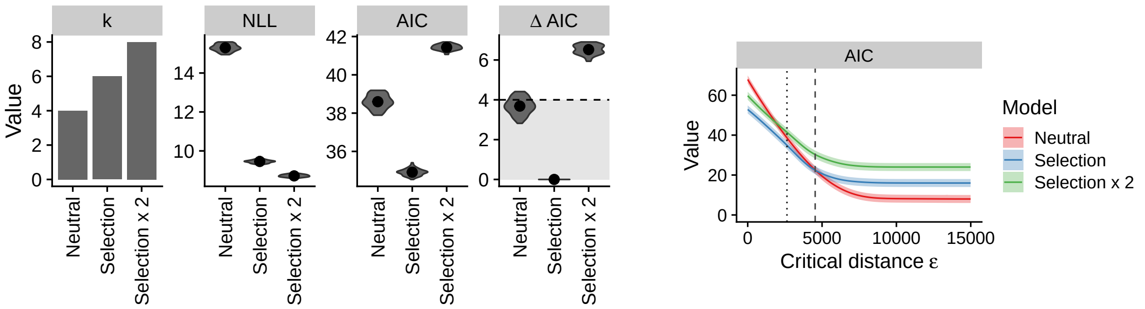

ε = 2640

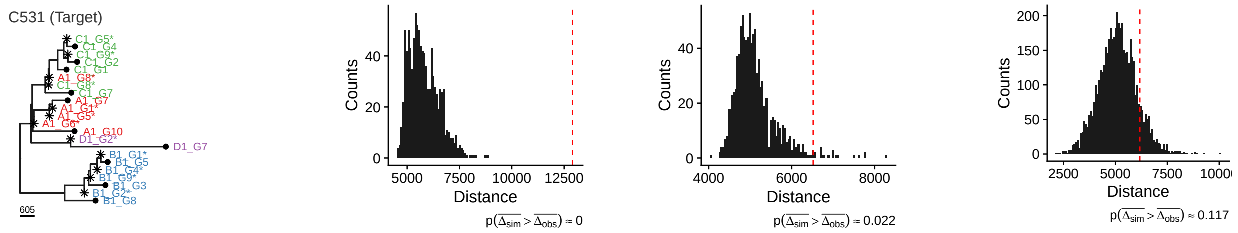

### Neutral

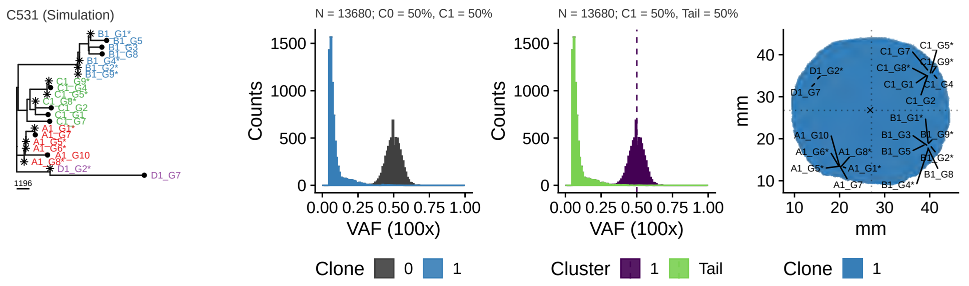

Parameters: $\lambda = 1$, $\mu = 0$, $d_{push} = 4.75$, $m = 0$, $t_s = 0$, $i_a = -1$

### Selection

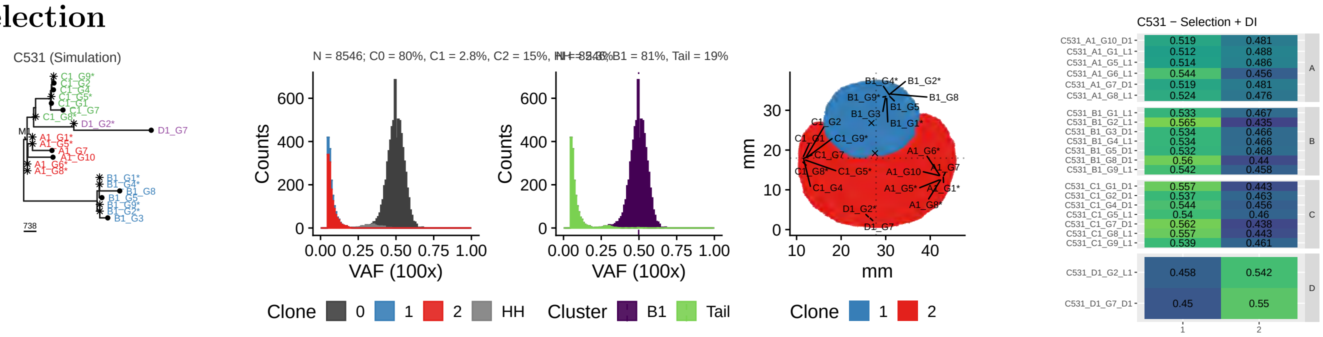

Parameters: $\lambda = (1\ \ 8.09)$, $\mu = (0\ \ 0)$, $d_{push} = (5.94\ \ 5.94)$, $m = (0\ \ 0)$, $t_s = (0\ \ 21500)$, $i_a = (-1\ \ 0)$

### 2x Selection

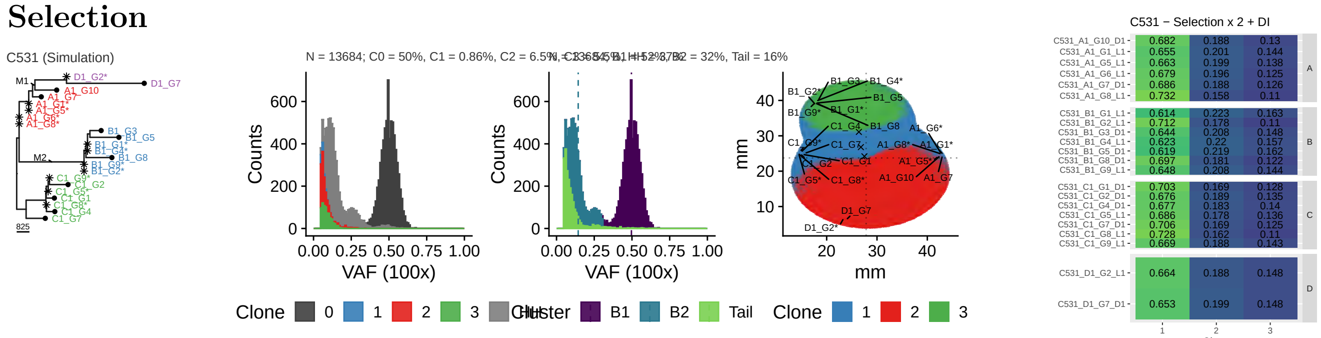

Parameters: $\lambda = (1\ \ 4.16\ \ 17)$, $\mu = (0\ \ 0\ \ 0)$, $d_{push} = (165\ \ 165\ \ 165)$, $m = (0\ \ 0\ \ 0)$, $t_s = (0\ \ 2140\ \ 21000)$, $i_a = (-1\ \ 0\ \ 0)$

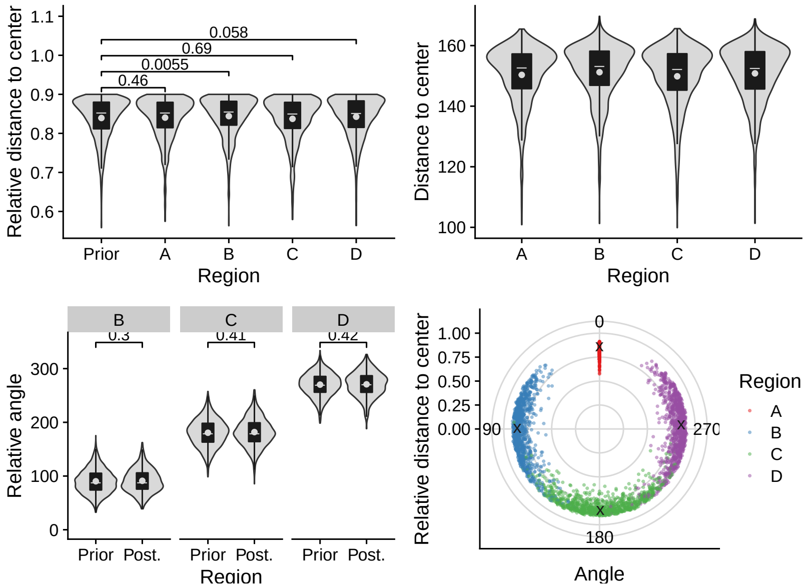
Relative distance to center
Region
Prior
A
B
C
D
0.058
0.69
0.0055
0.46
Distance to center
Relative angle
0.3
0.41
0.42
Prior Post.
Angle
0
90
180
270
A
B
C
D

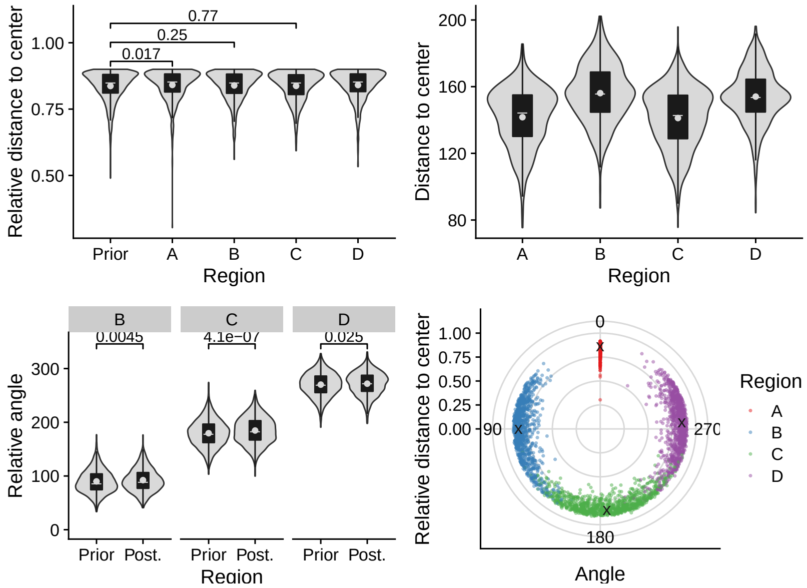
Relative distance to center
Region
Prior
A
B
C
D
0.77
0.25
0.017
Distance to center
Relative angle
0.0045
4.1e−07
0.025
Prior Post.
Angle
0
90
180
270
A
B
C
D

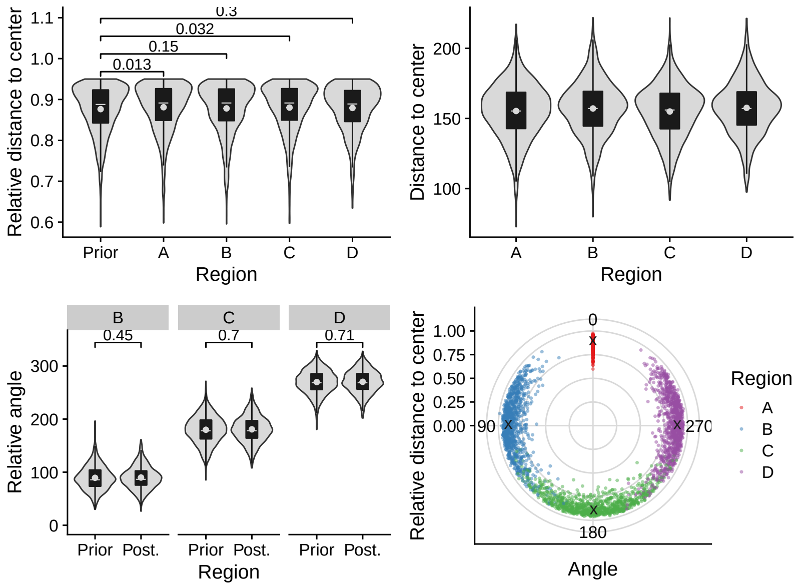
Relative distance to center
Region
Prior
A
B
C
D
0.3
0.032
0.15
0.013
Distance to center
Relative angle
0.45
0.7
0.71
Prior Post.
Angle
0
90
180
270
A
B
C
D

#### 2.8 C531 (excl. D1-G7)

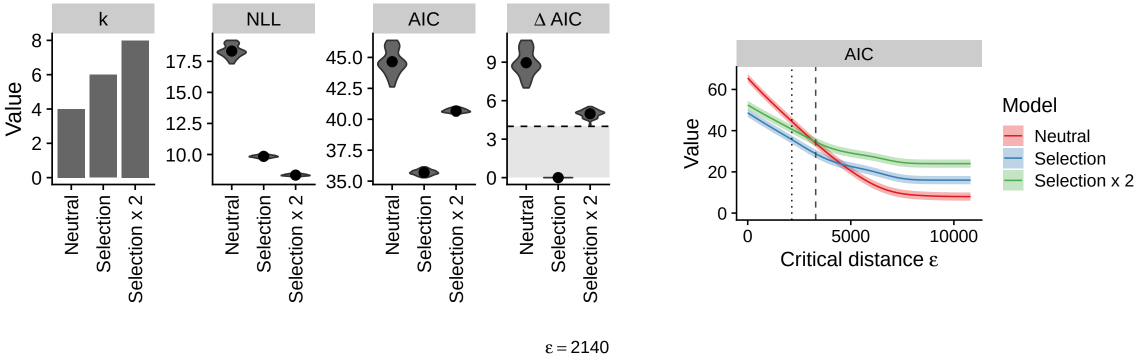

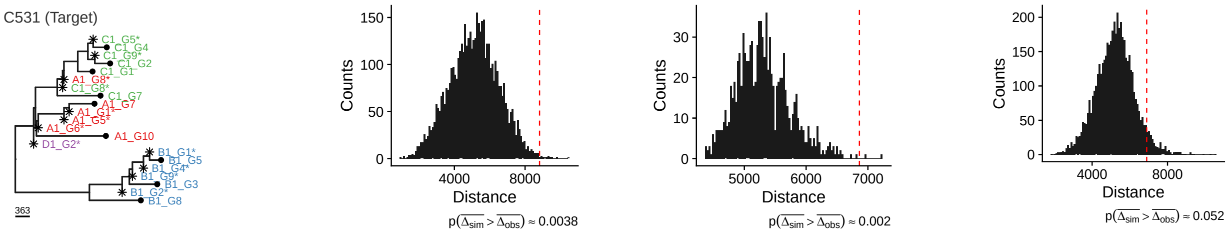

#### Neutral

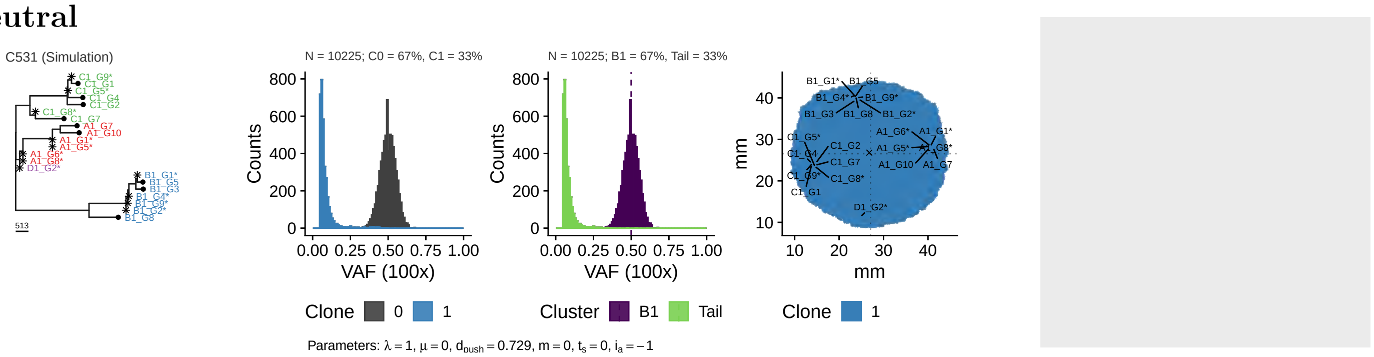

#### Selection

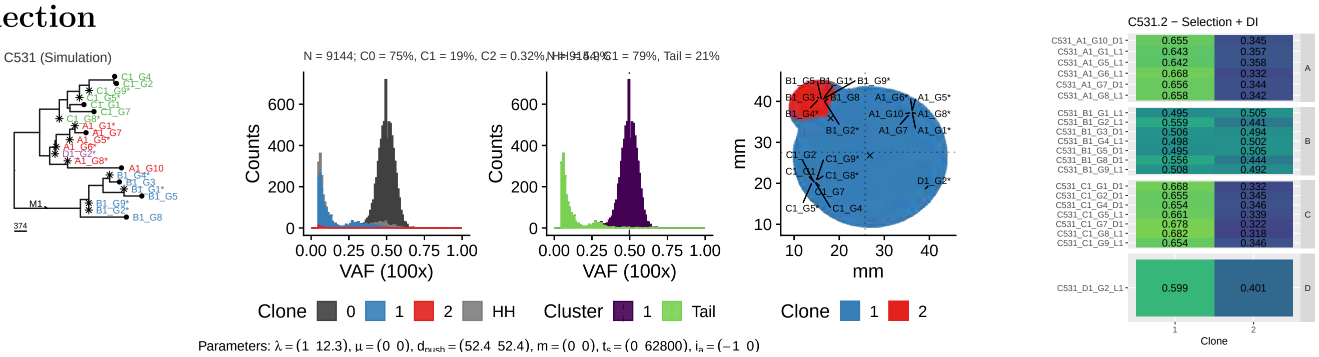

#### 2x Selection

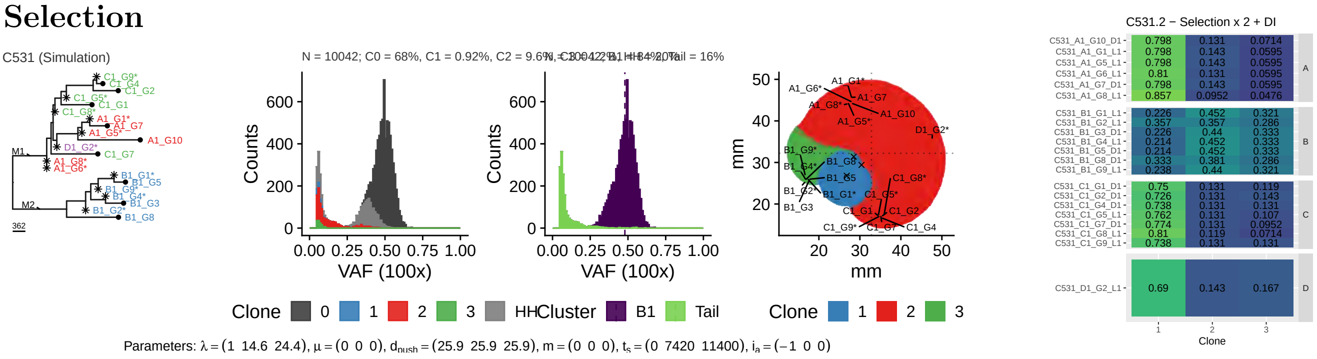

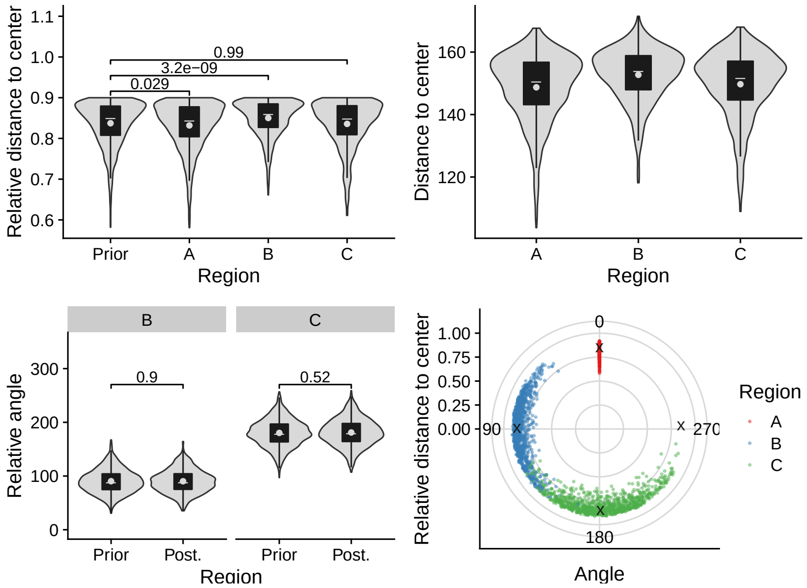
Relative distance to center
Region
Prior
A
B
C
0.99
3.2e−09
0.029
Distance to center
Relative angle
0.9
0.52
Prior
Post.
Angle
0
90
180
270
A
B
C

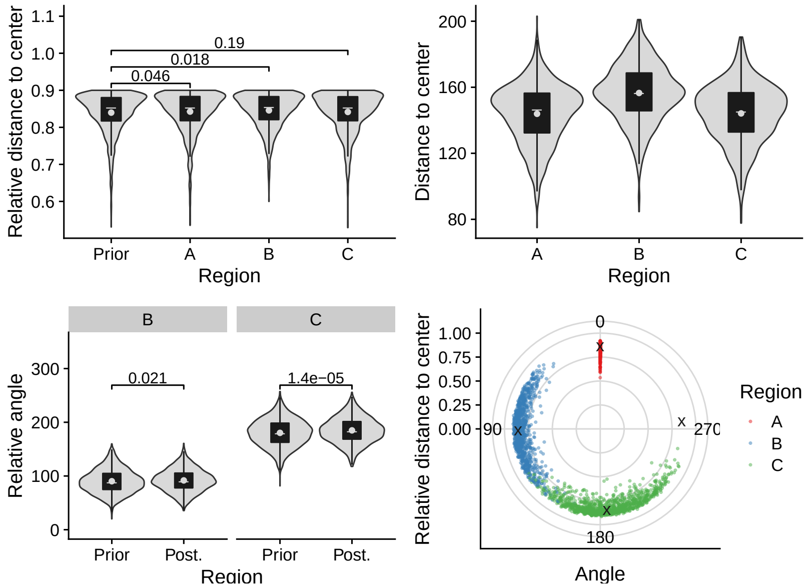
Relative distance to center
Region
Prior
A
B
C
0.19
0.018
0.046
Distance to center
Relative angle
0.021
1.4e−05
Prior
Post.
Angle
0
90
180
270
A
B
C

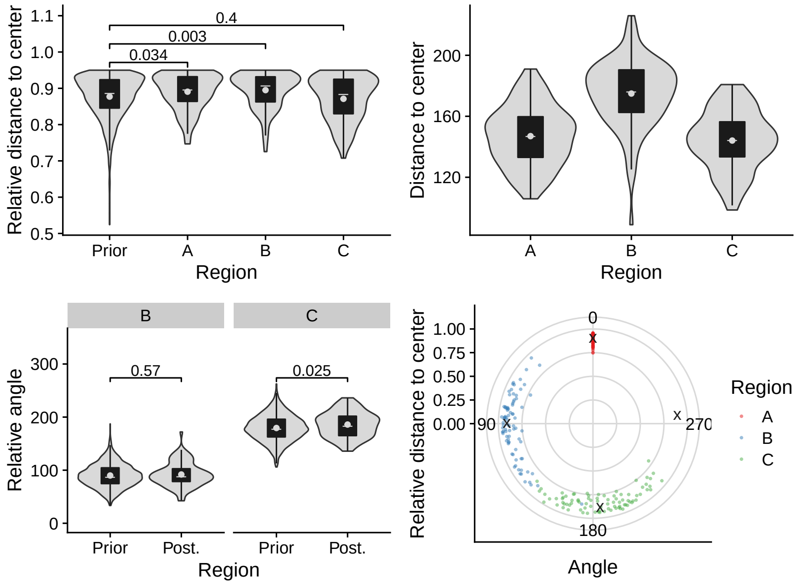
Relative distance to center
Region
Prior
A
B
C
0.4
0.003
0.034
Distance to center
Relative angle
0.57
0.025
Prior
Post.
Angle
0
90
180
270
A
B
C

## 2.9 C532

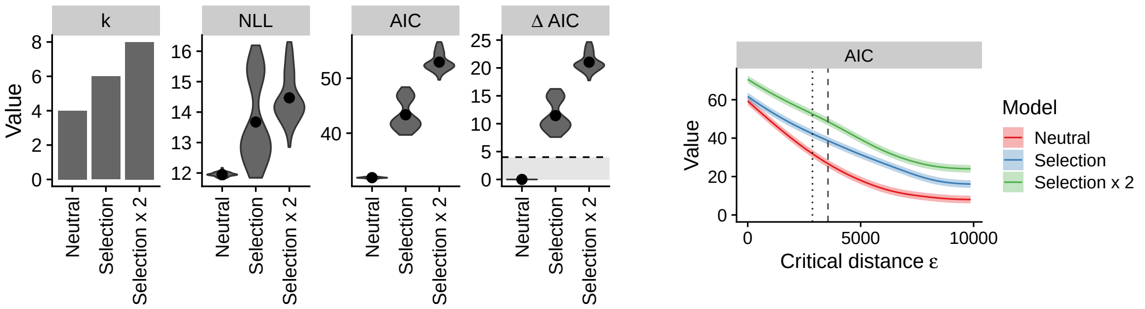

ε = 2870

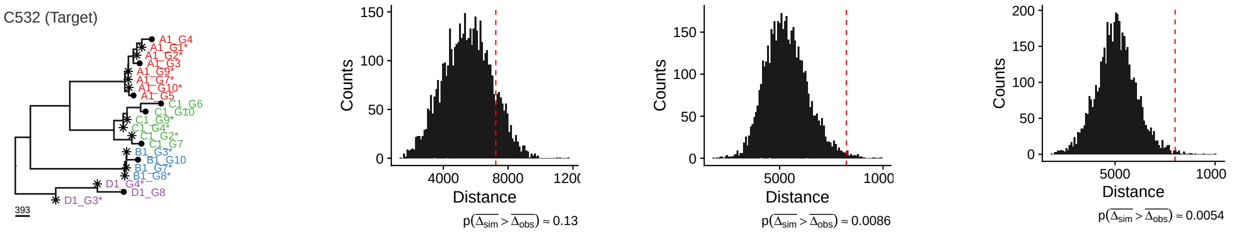

### Neutral

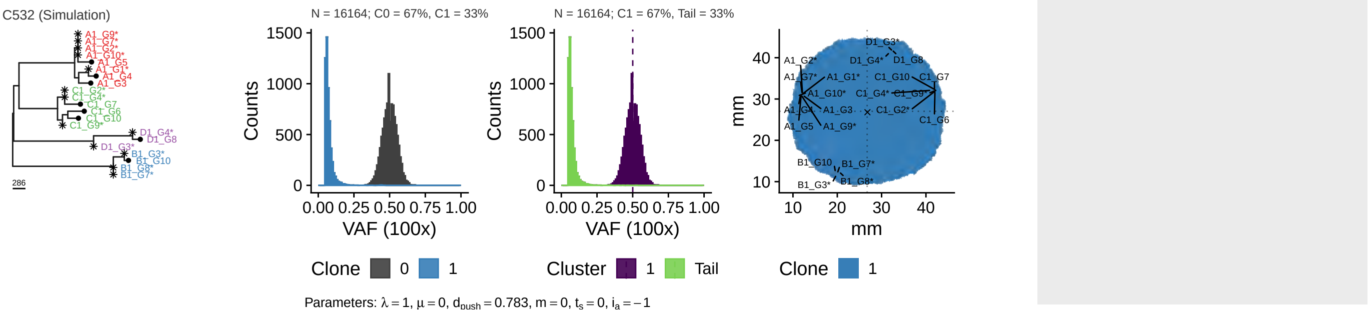

### Selection

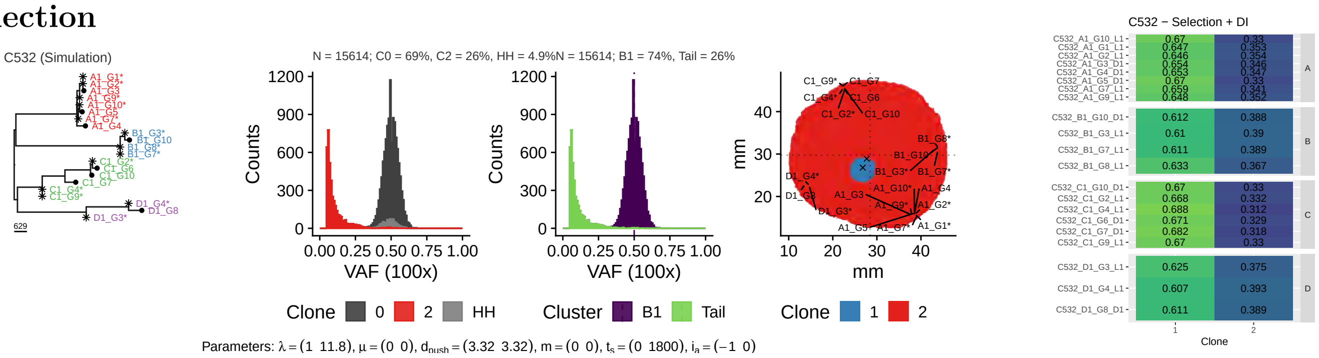

### 2x Selection

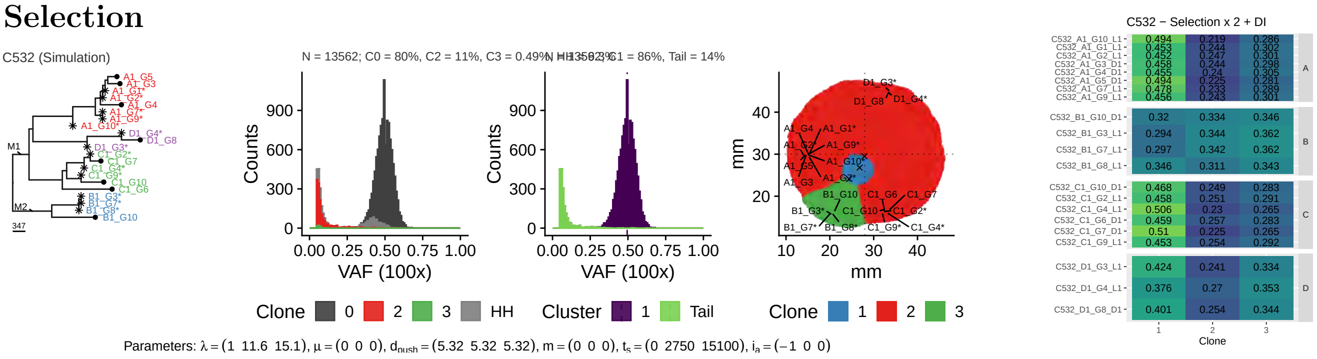

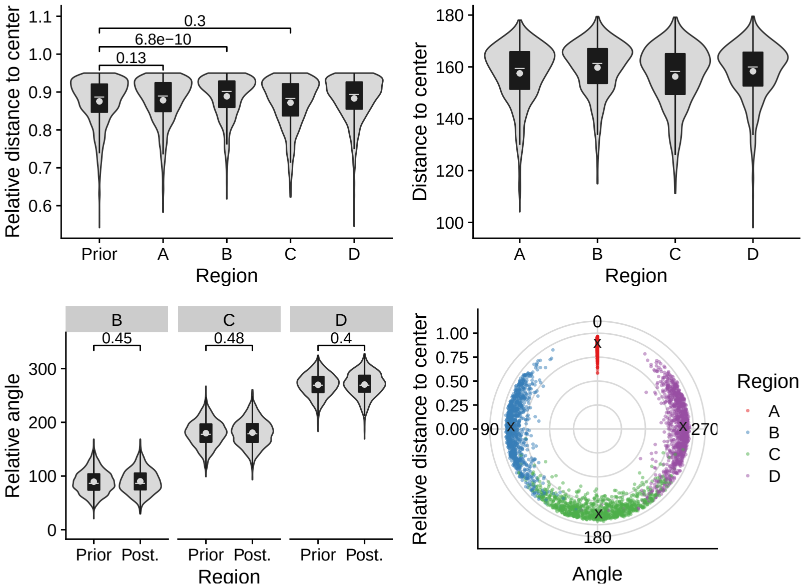
Relative distance to center
0.3
6.8e−10
0.13
Prior
A
B
C
D
Region
Distance to center
B
C
D
0.45
0.48
0.4
Relative angle
Prior Post.
Relative distance to center
0
90
180
270
Angle
Region
A
B
C
D

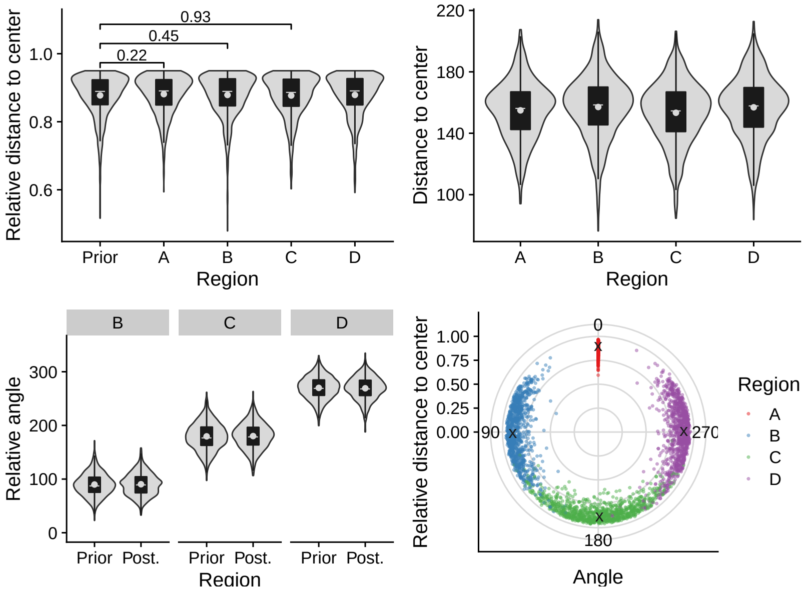
Relative distance to center
0.93
0.45
0.22
Prior
A
B
C
D
Region
Distance to center
B
C
D
Relative angle
Prior Post.
Relative distance to center
0
90
180
270
Angle
Region
A
B
C
D

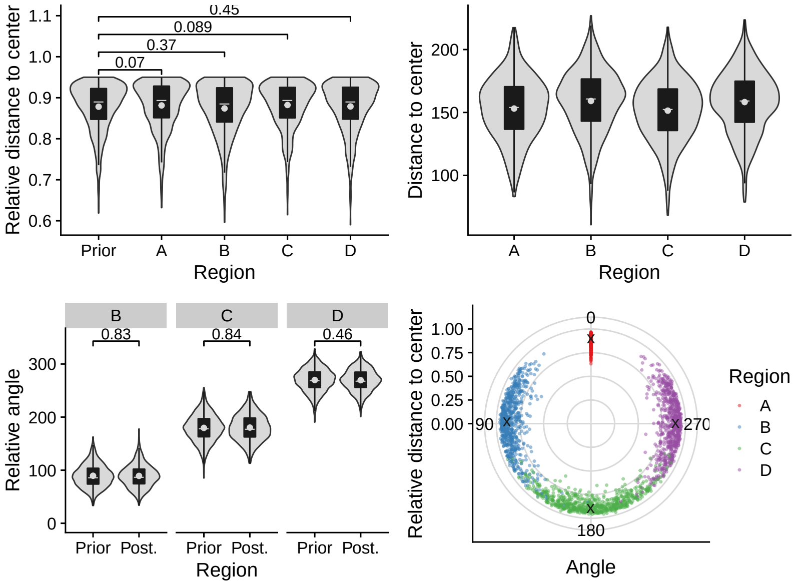
Relative distance to center
0.45
0.089
0.37
0.07
Prior
A
B
C
D
Region
Distance to center
B
C
D
0.83
0.84
0.46
Relative angle
Prior Post.
Relative distance to center
0
90
180
270
Angle
Region
A
B
C
D

## 2.10 C536

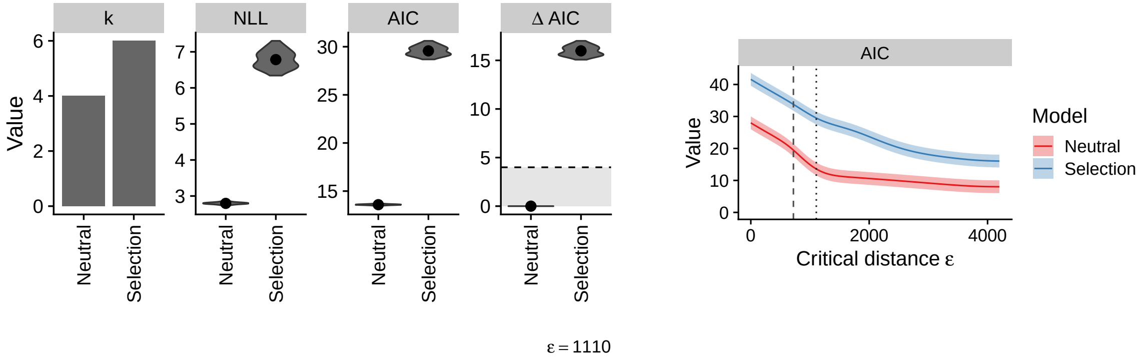

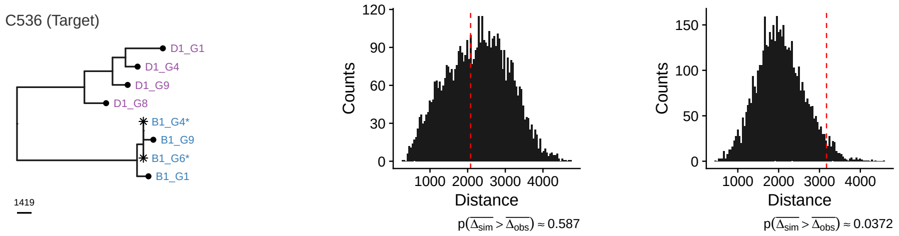

#### Neutral

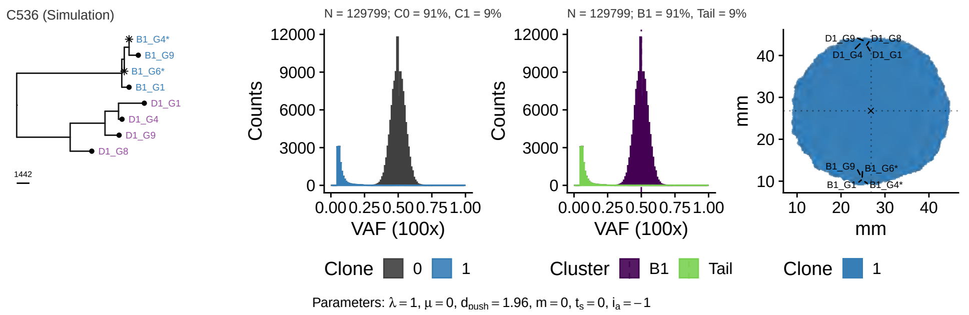

#### Selection

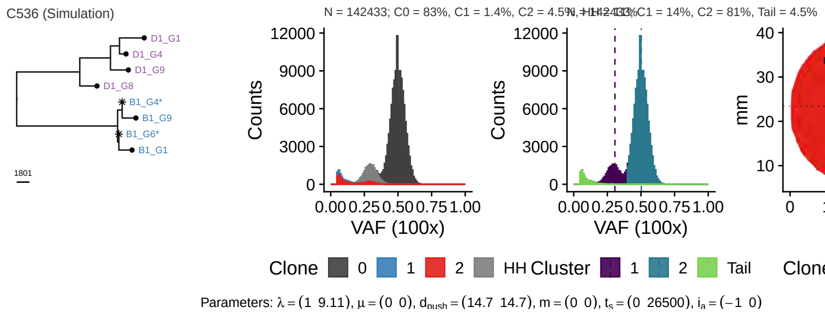

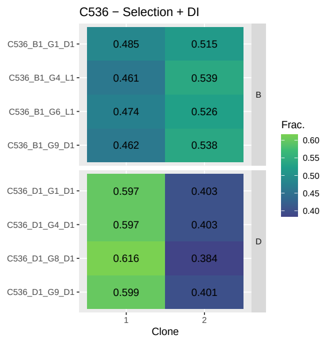

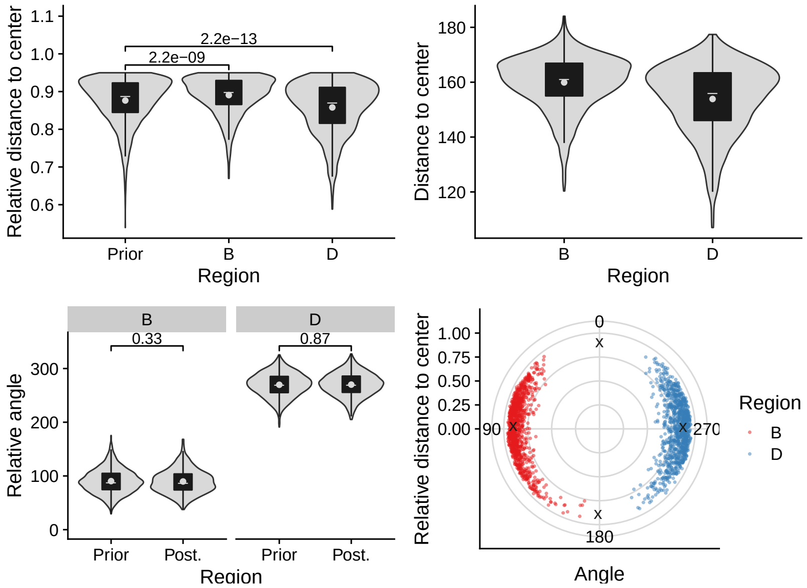
Relative distance to center
2.2e−13
2.2e−09
Prior
B
D
Region
Distance to center
Relative angle
0.33
0.87
Prior
Post.
Relative distance to center
Angle
Region
B
D

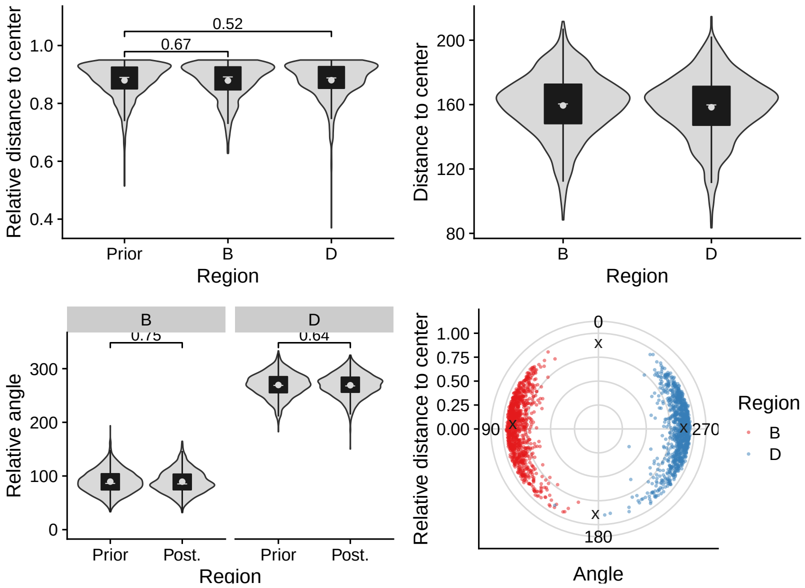
Relative distance to center
0.52
0.67
Prior
B
D
Region
Distance to center
Relative angle
0.75
0.64
Prior
Post.
Relative distance to center
Angle
Region
B
D

## 2.11 C537

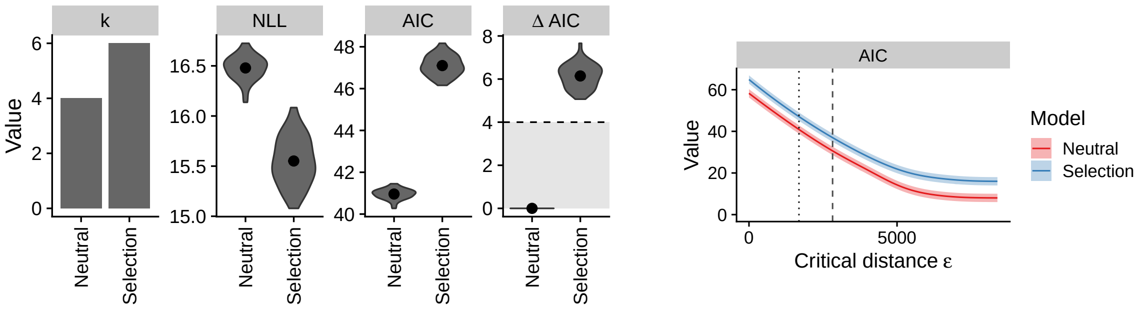

ε = 1690

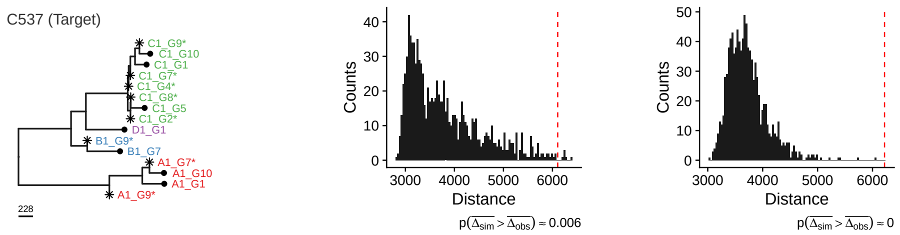

#### Neutral

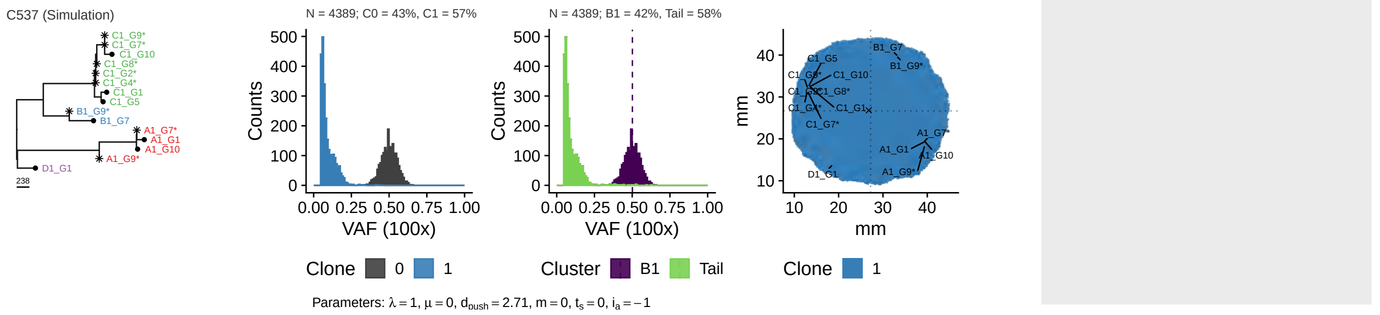

#### Selection

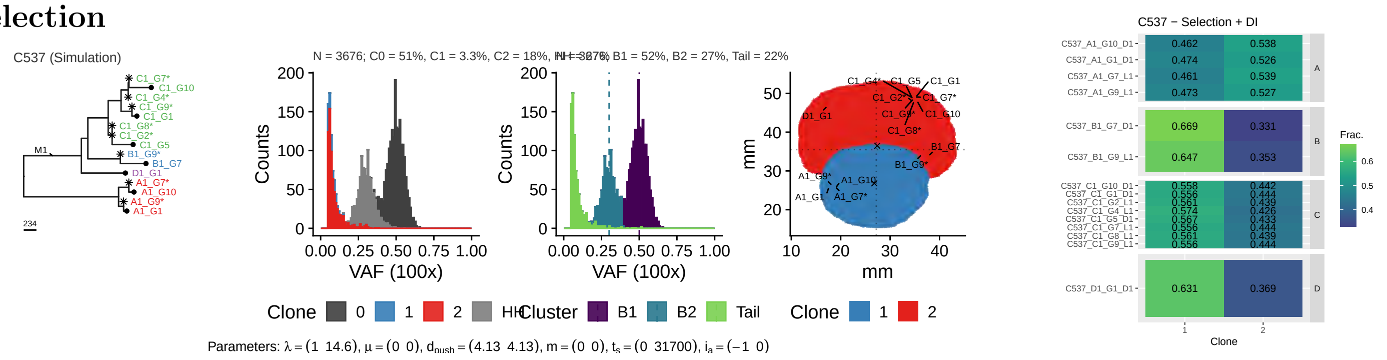

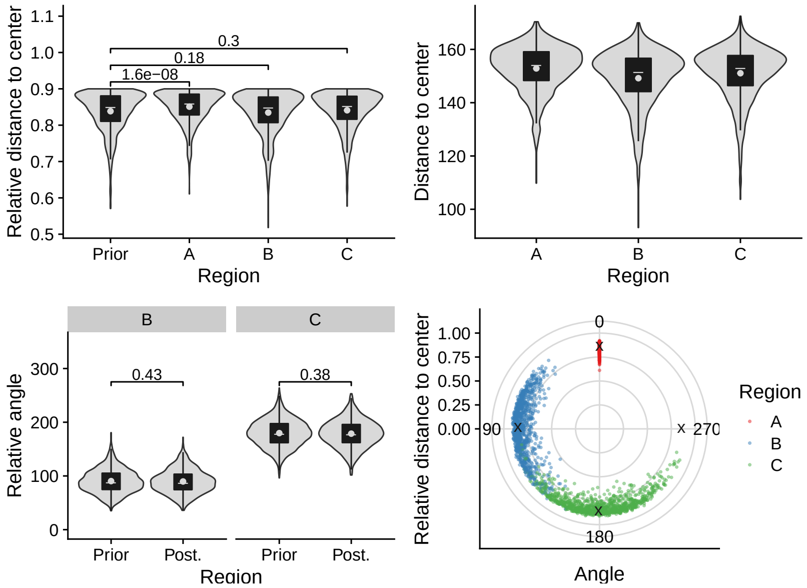

Relative distance to center
1.1
1.0
0.9
0.8
0.7
0.6
0.5
0.3
0.18
1.6e−08
Prior
A
B
C
Region
Distance to center
160
140
120
100
A
B
C
Region
B
C
Relative angle
300
200
100
0
0.43
0.38
Prior
Post.
Prior
Post.
Region
Relative distance to center
1.00
0.75
0.50
0.25
0.00
0
90
270
180
Angle
Region
A
B
C

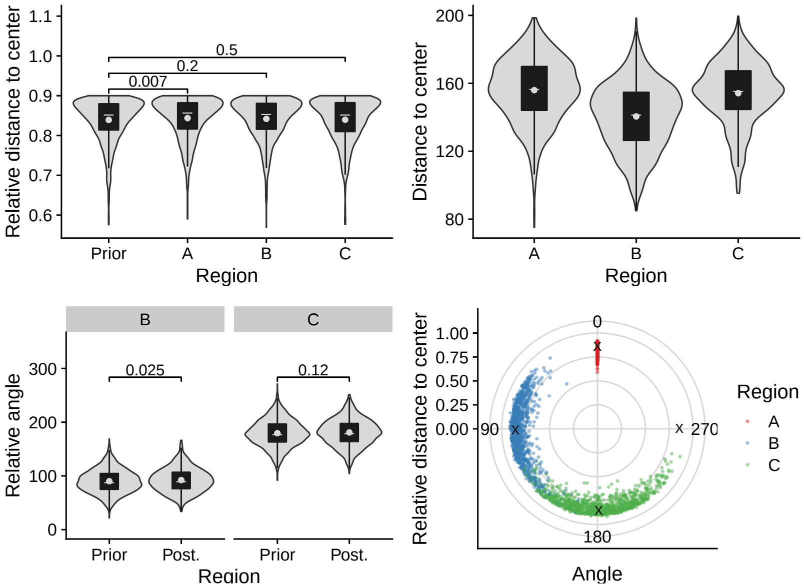

Relative distance to center
1.1
1.0
0.9
0.8
0.7
0.6
0.5
0.2
0.007
Prior
A
B
C
Region
Distance to center
200
160
120
80
A
B
C
Region
B
C
Relative angle
300
200
100
0
0.025
0.12
Prior
Post.
Prior
Post.
Region
Relative distance to center
1.00
0.75
0.50
0.25
0.00
0
90
270
180
Angle
Region
A
B
C

## 2.12 C538

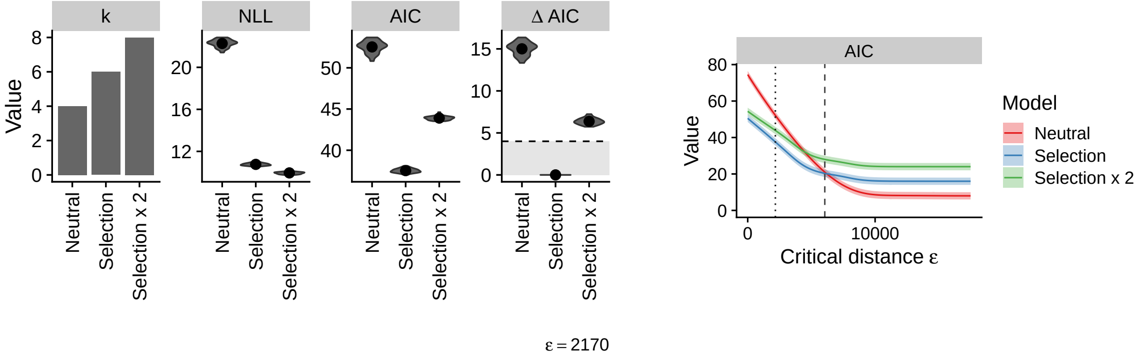

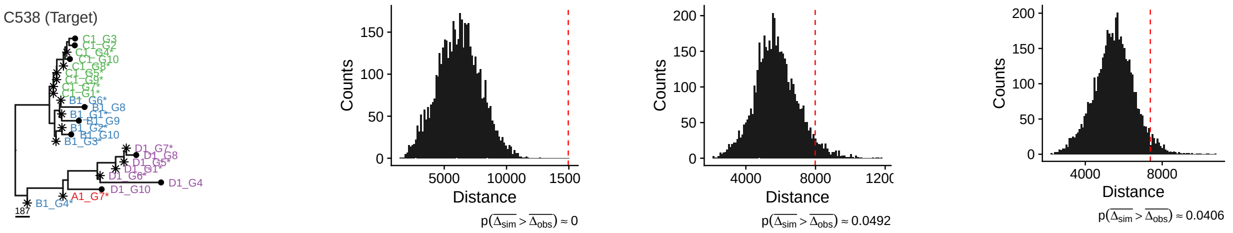

#### Neutral

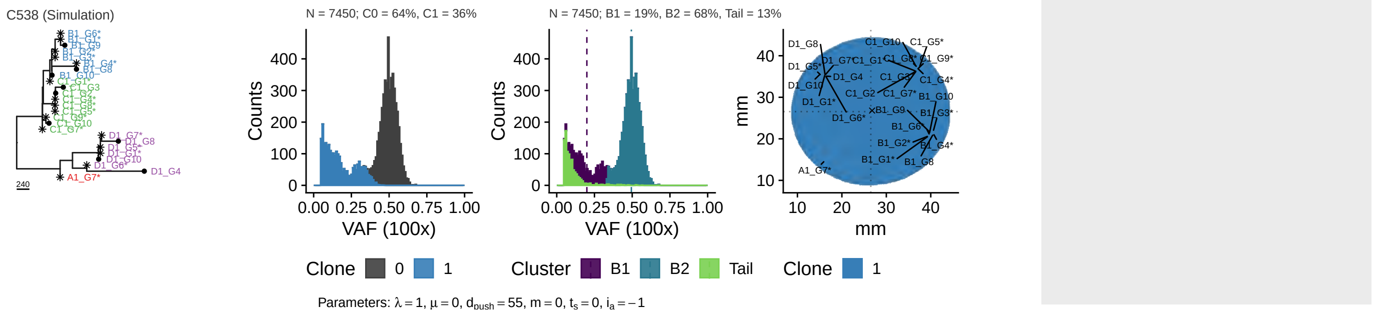

#### Selection

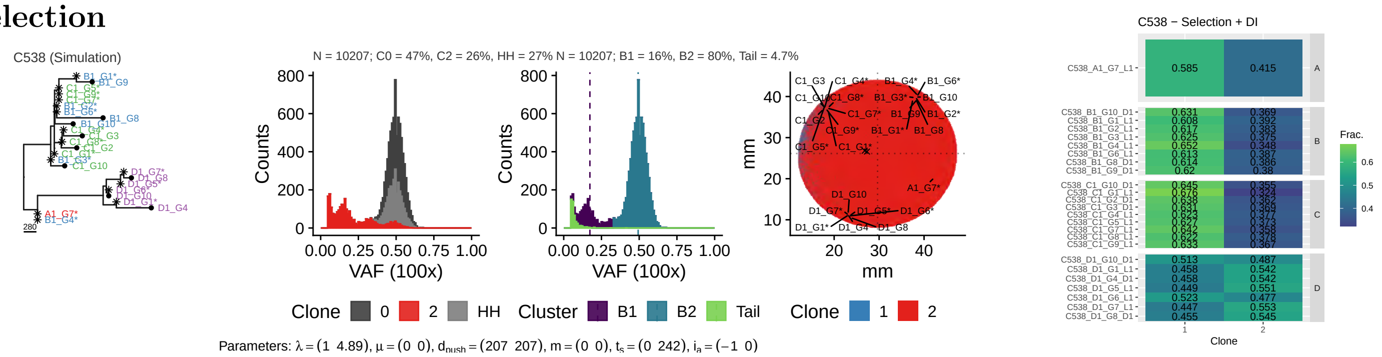

#### 2x Selection

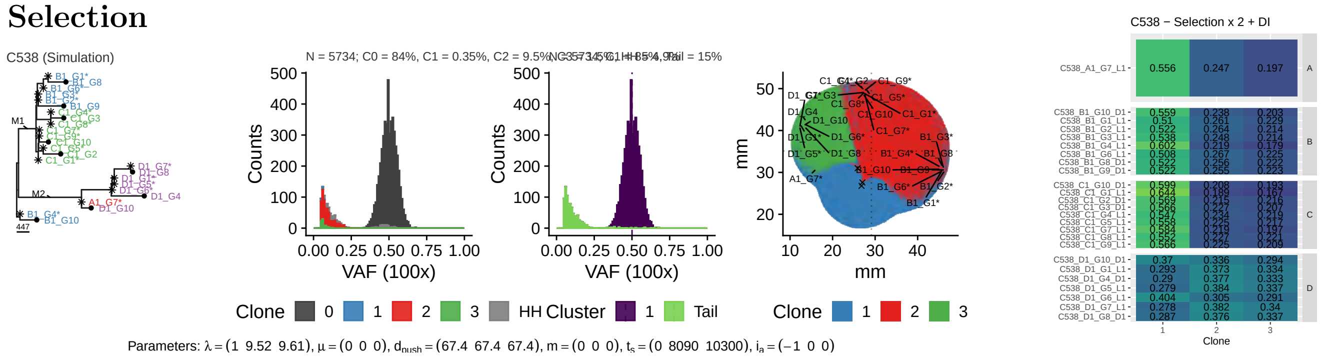

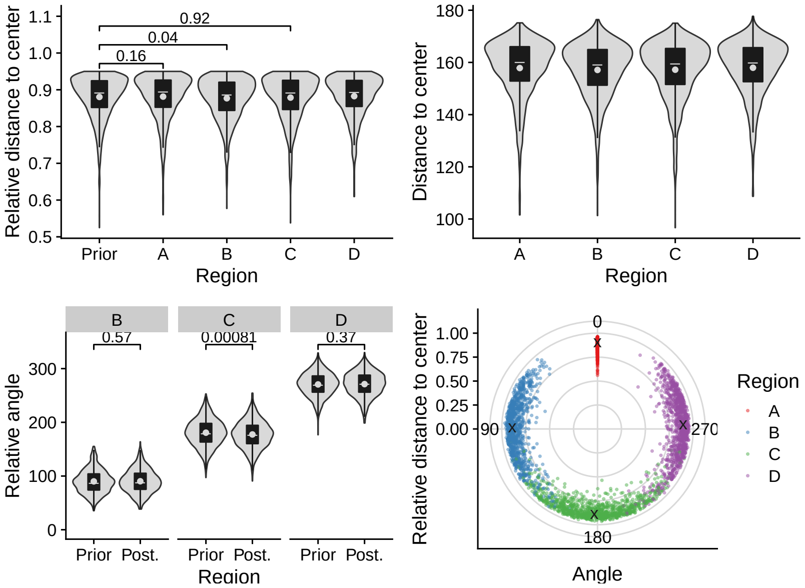
0.92
0.04
0.16
Relative distance to center
Prior A B C D
Region
Distance to center
A B C D
Region
B C D
0.57
0.00081
0.37
Relative angle
Prior Post. Prior Post. Prior Post.
Region
Relative distance to center
0
90
270
180
Angle
Region
A
B
C
D

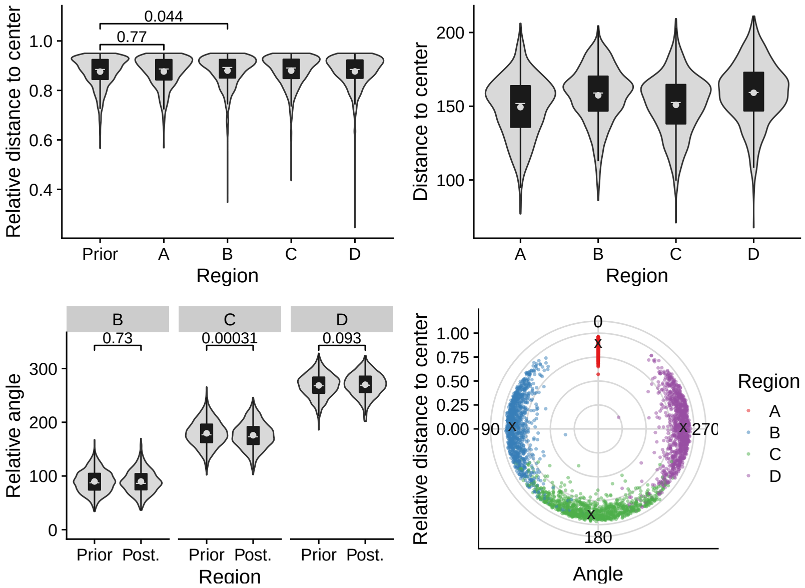
0.044
0.77
Relative distance to center
Prior A B C D
Region
Distance to center
A B C D
Region
B C D
0.73
0.00031
0.093
Relative angle
Prior Post. Prior Post. Prior Post.
Region
Relative distance to center
0
90
270
180
Angle
Region
A
B
C
D

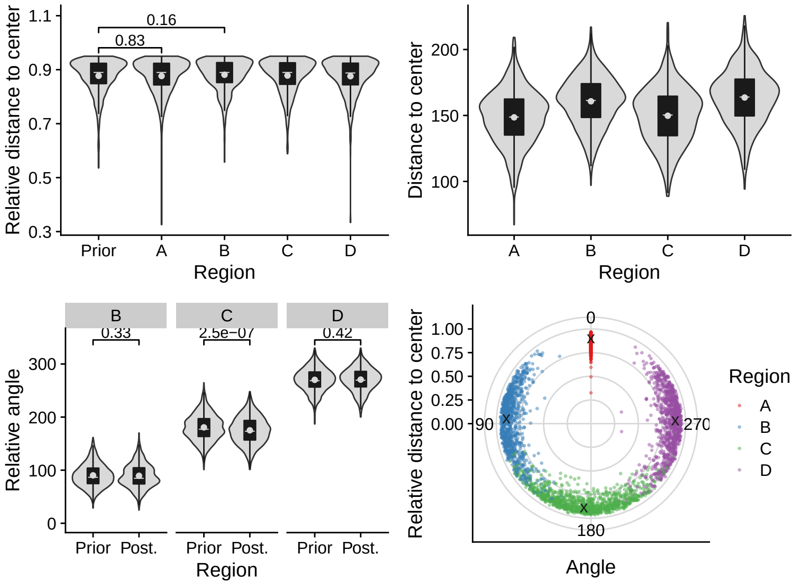
0.16
0.83
Relative distance to center
Prior A B C D
Region
Distance to center
A B C D
Region
B C D
0.33
2.5e−07
0.42
Relative angle
Prior Post. Prior Post. Prior Post.
Region
Relative distance to center
0
90
270
180
Angle
Region
A
B
C
D

## 2.13 C539

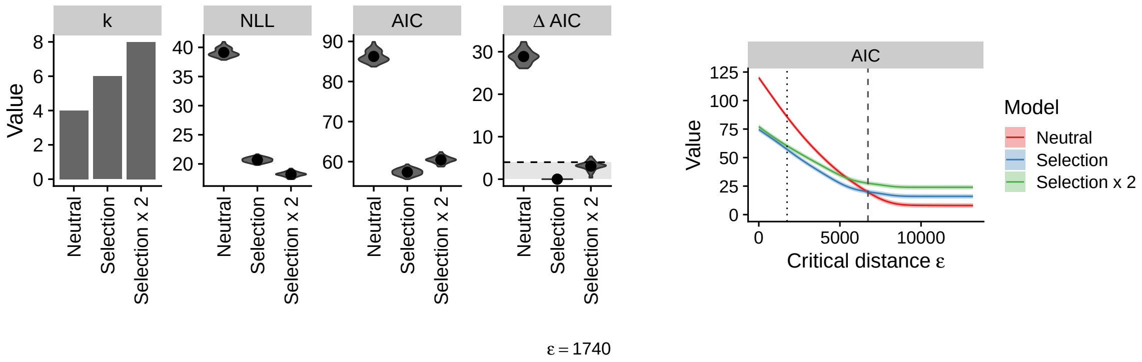

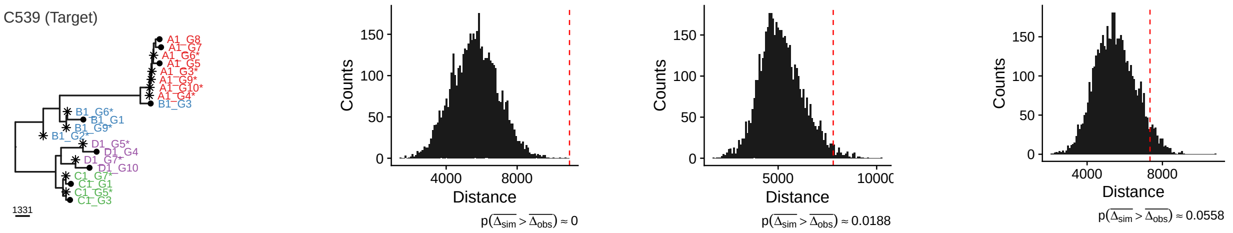

#### Neutral

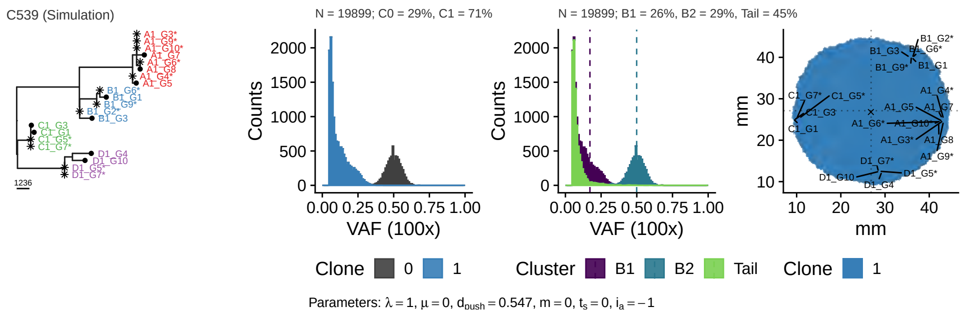

#### Selection

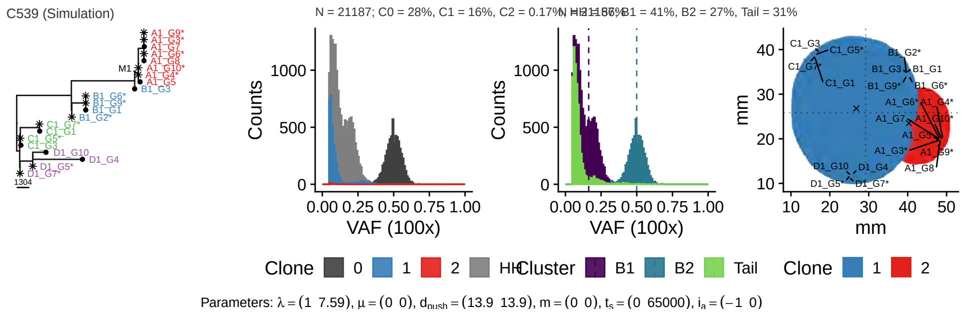

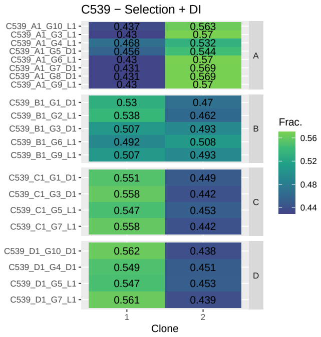

#### 2x Selection

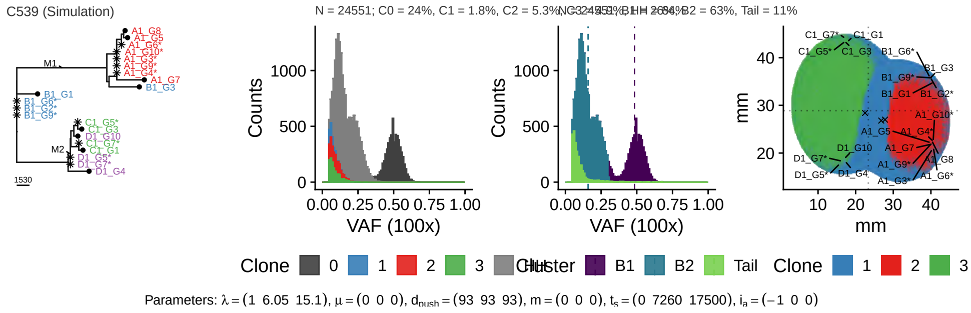

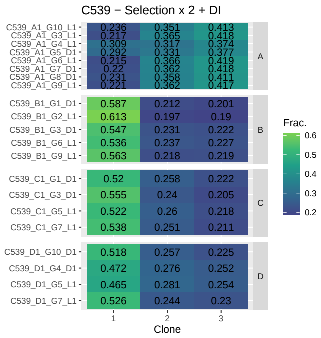

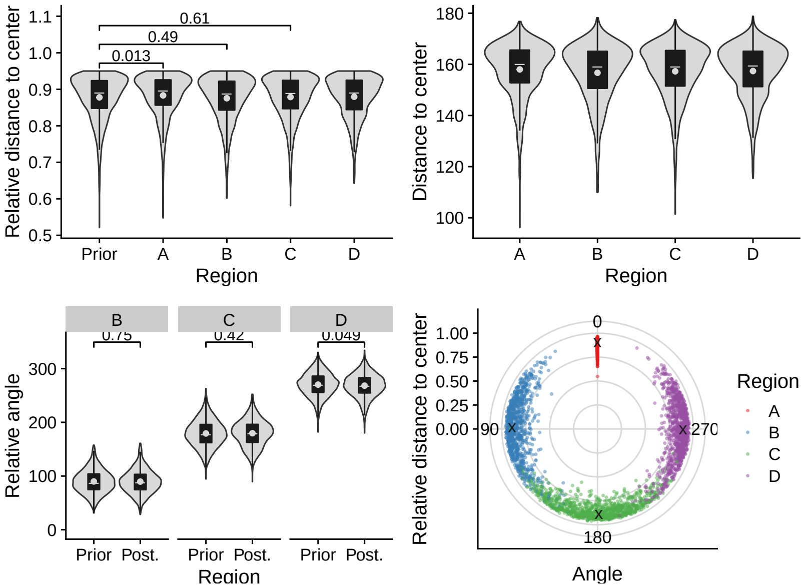
Relative distance to center
0.61
0.49
0.013
Prior A B C D
Region
Distance to center
A B C D
Region
B C D
0.75 0.42 0.049
Relative angle
Prior Post. Prior Post. Prior Post.
Region
Relative distance to center
0
90
270
180
Angle
Region
A
B
C
D

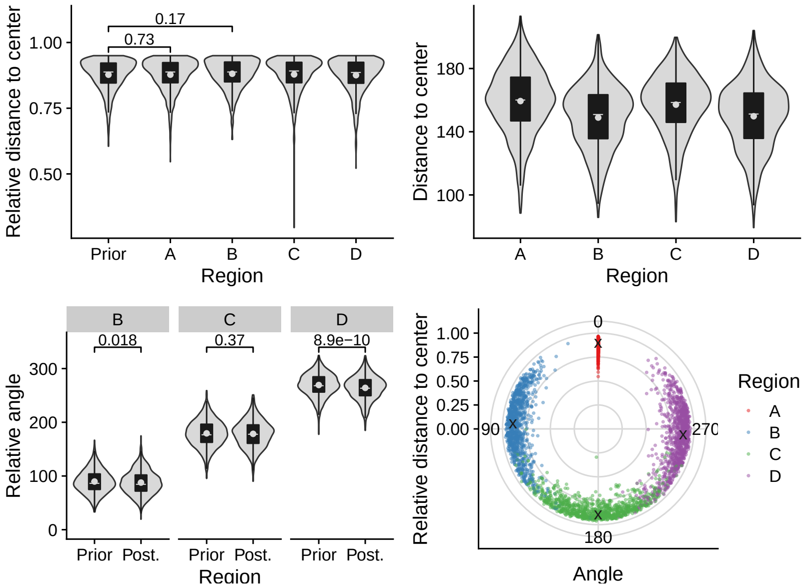
Relative distance to center
0.17
0.73
Prior A B C D
Region
Distance to center
A B C D
Region
B C D
0.018 0.37 8.9e−10
Relative angle
Prior Post. Prior Post. Prior Post.
Region
Relative distance to center
0
90
270
180
Angle
Region
A
B
C
D

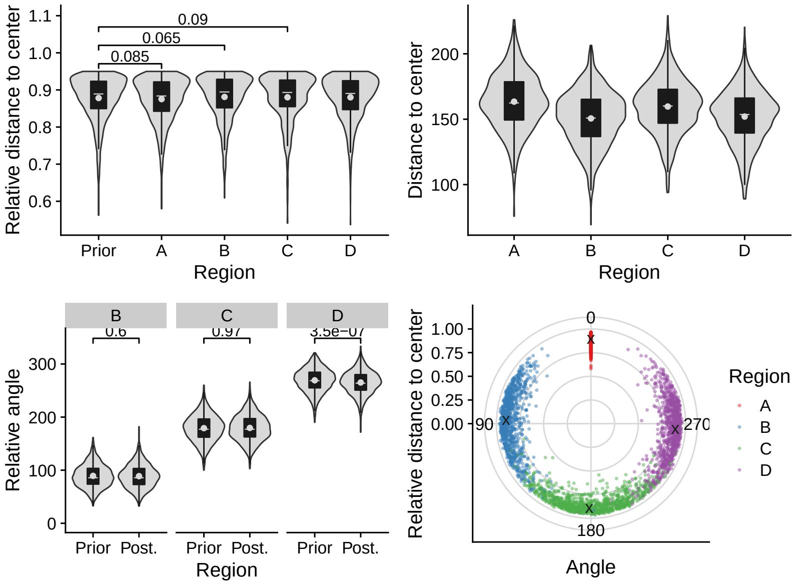
Relative distance to center
0.09
0.065
0.085
Prior A B C D
Region
Distance to center
A B C D
Region
B C D
0.6 0.97 3.5e−07
Relative angle
Prior Post. Prior Post. Prior Post.
Region
Relative distance to center
0
90
270
180
Angle
Region
A
B
C
D

## 2.14 C542

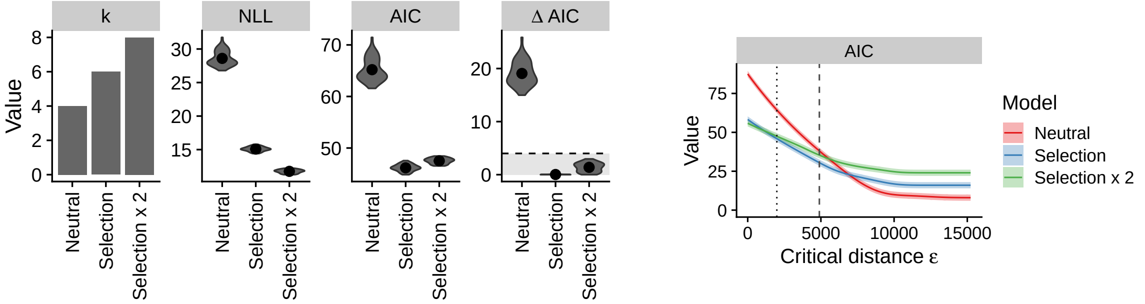

ε = 1990

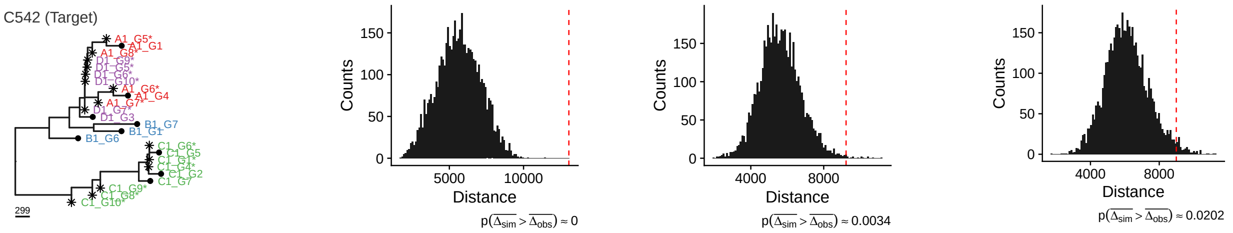

#### Neutral

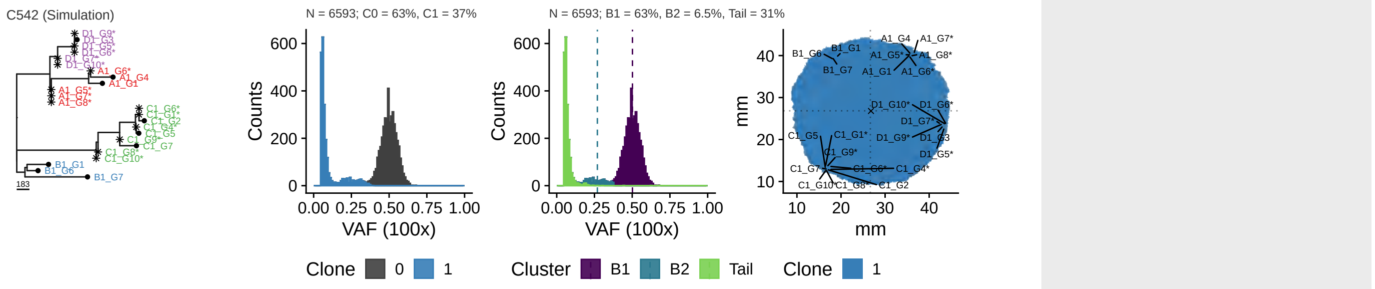

Parameters: $\lambda = 1$, $\mu = 0$, $d_{push} = 1.02$, $m = 0$, $t_s = 0$, $i_a = -1$

#### Selection

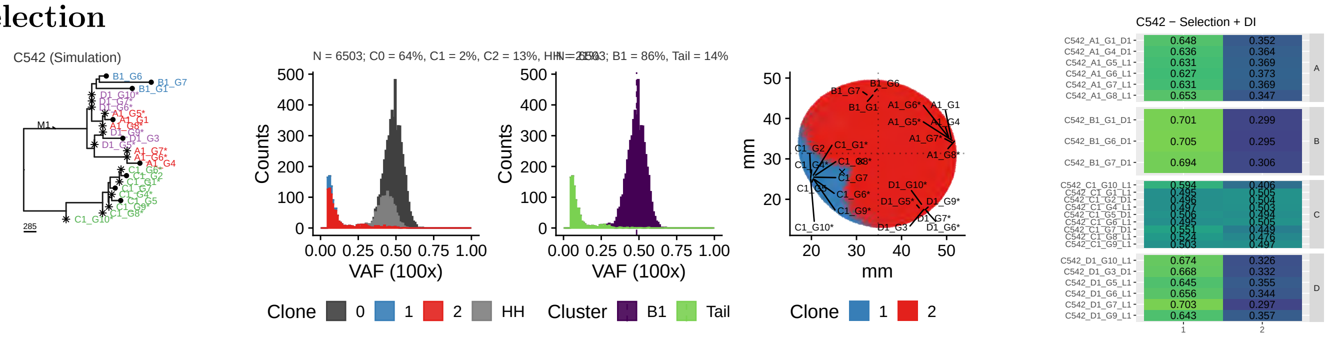

Parameters: $\lambda = (1\ \ 20.4)$, $\mu = (0\ \ 0)$, $d_{push} = (235\ \ 235)$, $m = (0\ \ 0)$, $t_s = (0\ \ 7650)$, $i_a = (-1\ \ 0)$

#### 2x Selection

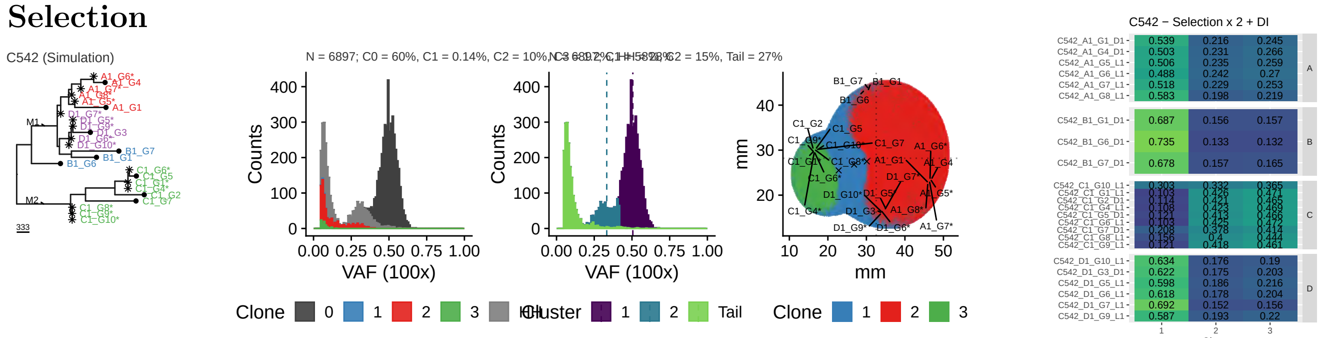

Parameters: $\lambda = (1\ \ 10.6\ \ 22.8)$, $\mu = (0\ \ 0\ \ 0)$, $d_{push} = (188\ \ 188\ \ 188)$, $m = (0\ \ 0\ \ 0)$, $t_s = (0\ \ 9000\ \ 17300)$, $i_a = (-1\ \ 0\ \ 0)$

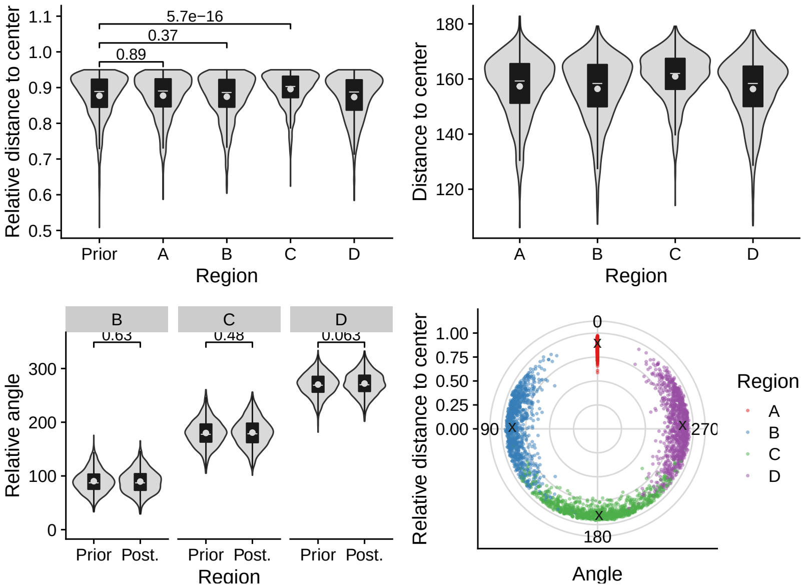
Relative distance to center
5.7e−16
0.37
0.89
Prior
A
B
C
D
Region
Distance to center
B
0.63
C
0.48
D
0.063
Relative angle
Prior Post.
0
90
180
270
Angle

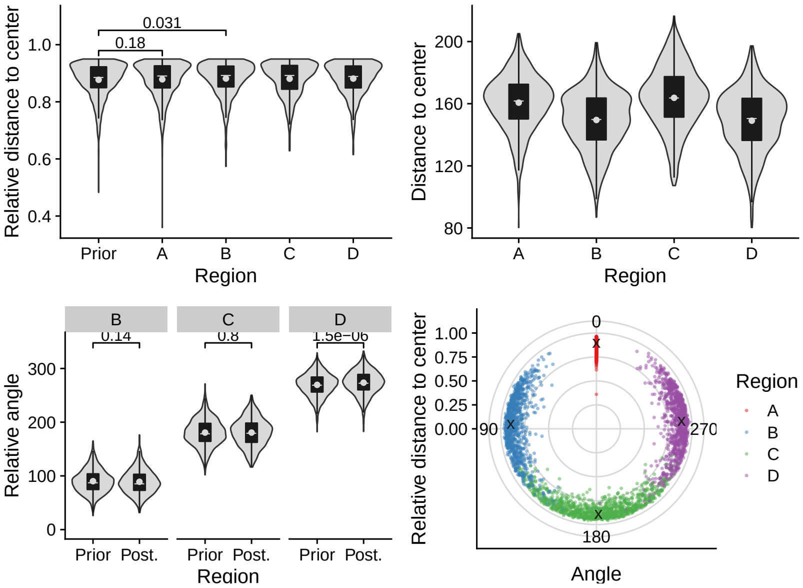
Relative distance to center
0.031
0.18
Prior
A
B
C
D
Region
Distance to center
B
0.14
C
0.8
D
1.5e−06
Relative angle
Prior Post.
0
90
180
270
Angle

## 2.15 C543

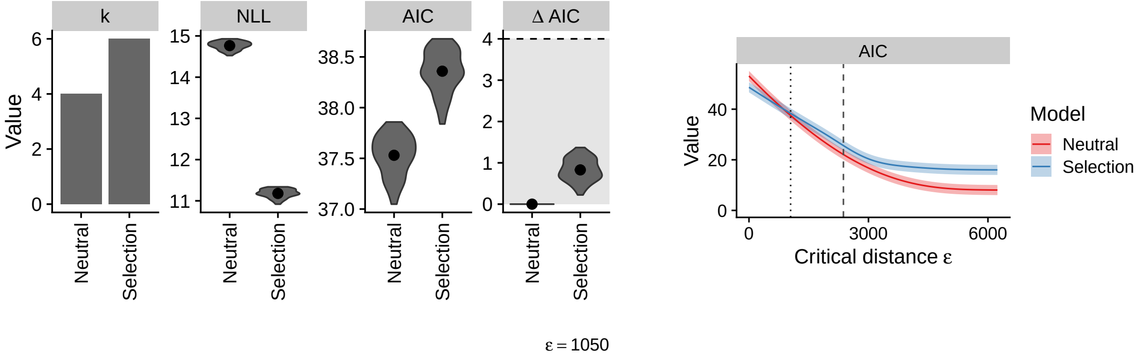

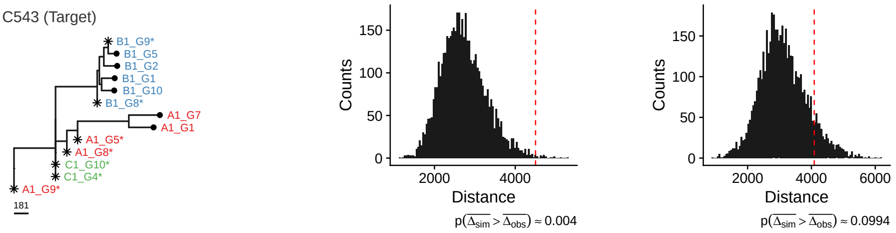

#### Neutral

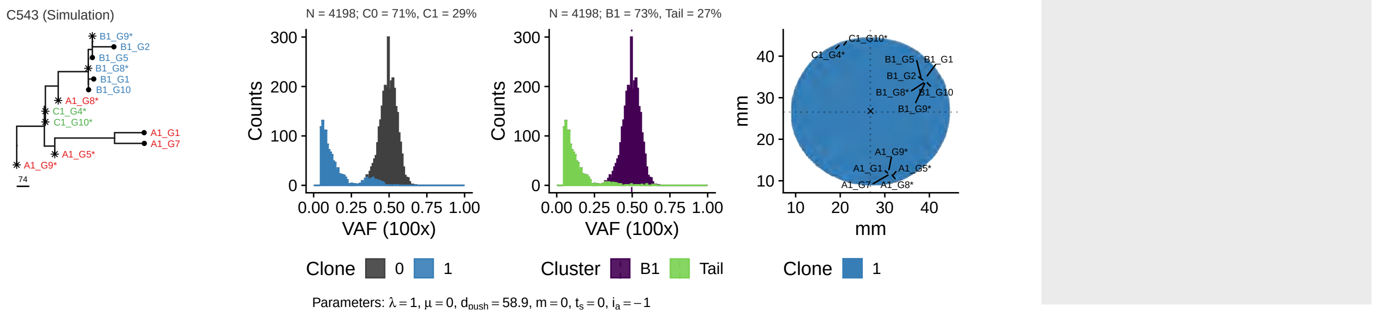

#### Selection

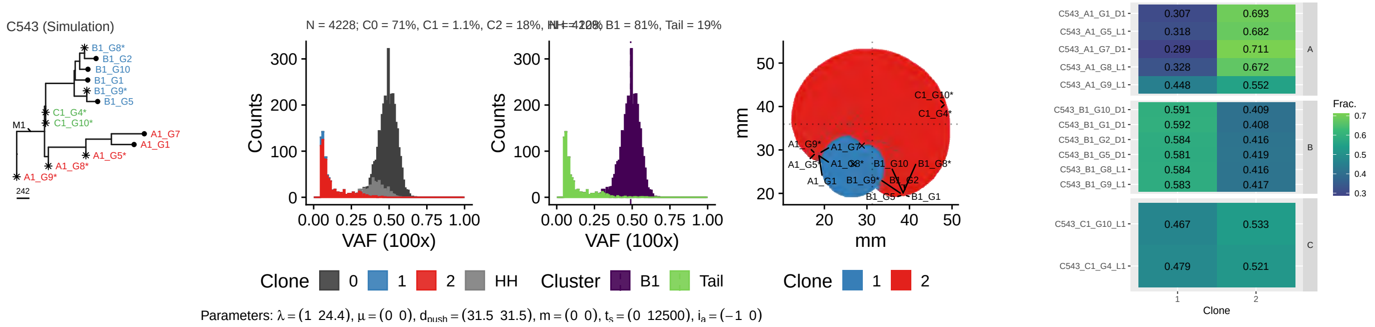

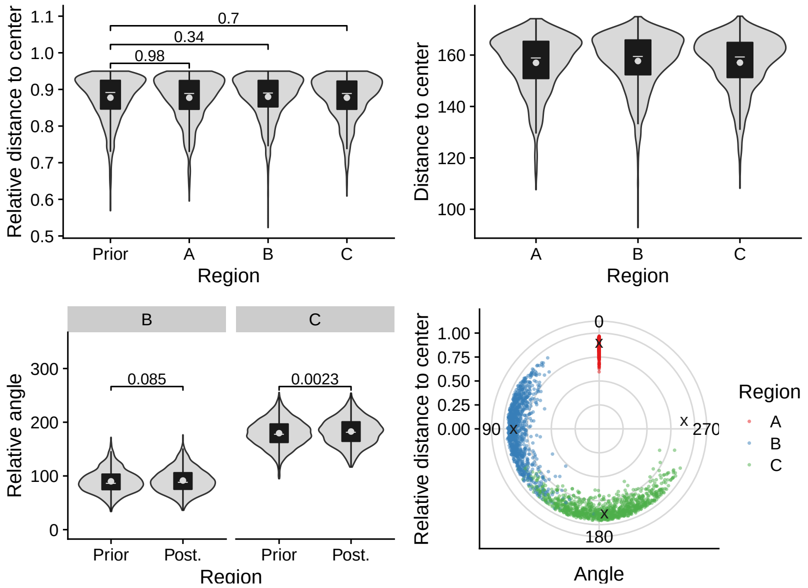
Relative distance to center
1.1
1.0
0.9
0.8
0.7
0.6
0.5
0.7
0.34
0.98
Prior
A
B
C
Region
Distance to center
160
140
120
100
A
B
C
Region
B
C
Relative angle
300
200
100
0
0.085
0.0023
Prior
Post.
Prior
Post.
Region
Relative distance to center
1.00
0.75
0.50
0.25
0.00
0
90
270
180
Angle
Region
A
B
C

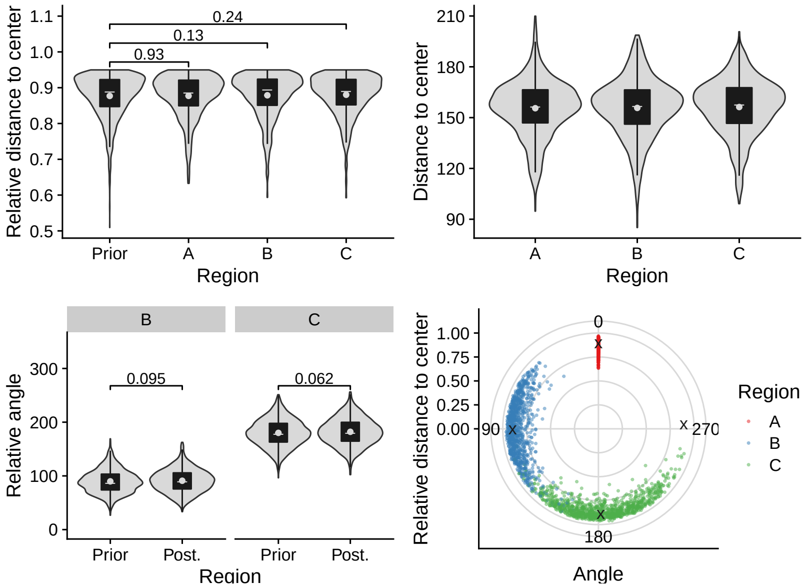
Relative distance to center
1.1
1.0
0.9
0.8
0.7
0.6
0.5
0.24
0.13
0.93
Prior
A
B
C
Region
Distance to center
210
180
150
120
90
A
B
C
Region
B
C
Relative angle
300
200
100
0
0.095
0.062
Prior
Post.
Prior
Post.
Region
Relative distance to center
1.00
0.75
0.50
0.25
0.00
0
90
270
180
Angle
Region
A
B
C

#### 2.16 C543 (excl. A1-G9)

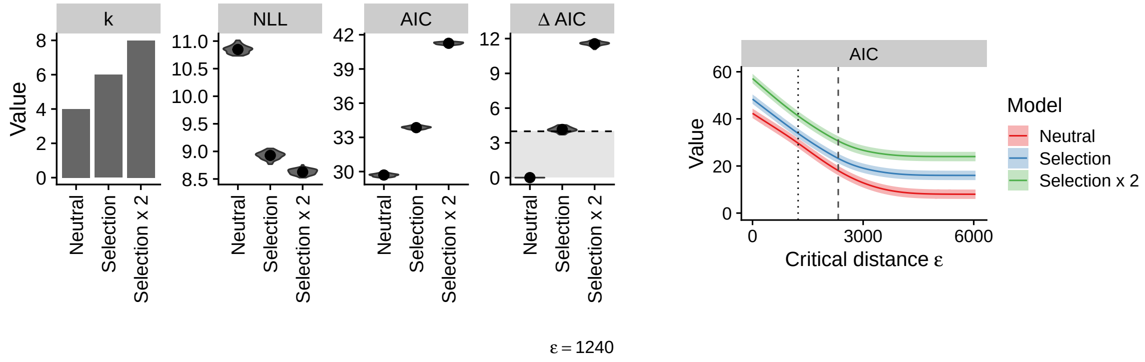

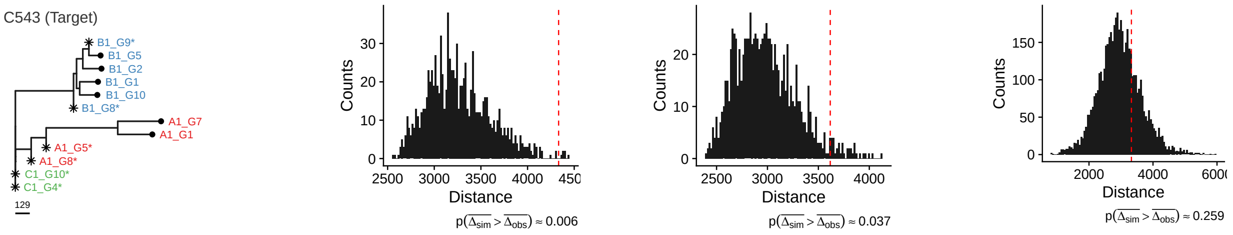

#### Neutral

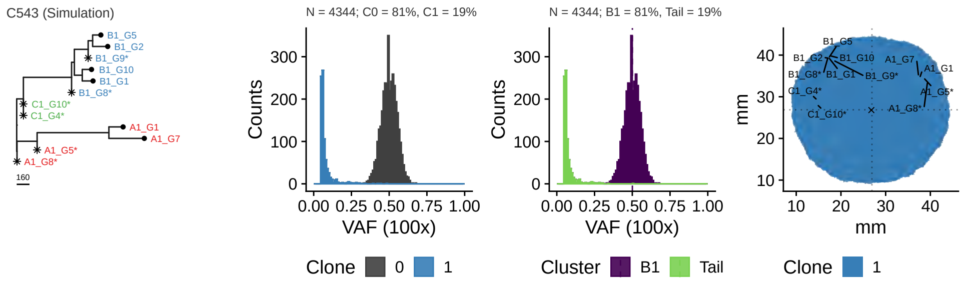

Parameters: $\lambda = 1$, $\mu = 0$, $d_{push} = 3.39$, $m = 0$, $t_s = 0$, $i_a = -1$

#### Selection

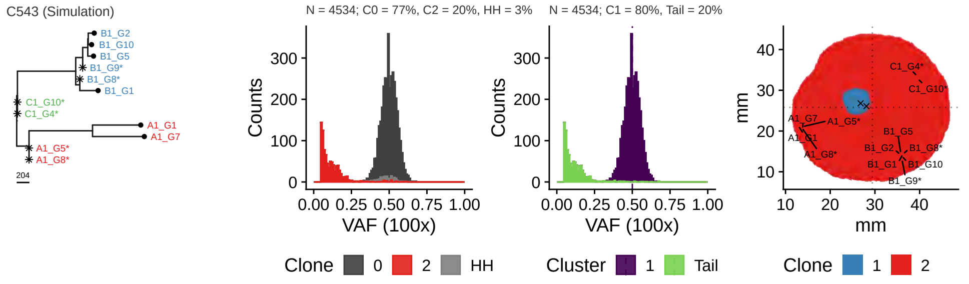

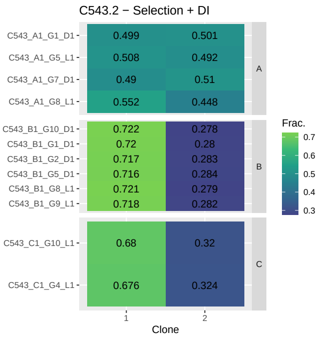

Parameters: $\lambda = (1\ \ 6.77)$, $\mu = (0\ \ 0)$, $d_{push} = (5.13\ \ 5.13)$, $m = (0\ \ 0)$, $t_s = (0\ \ 1290)$, $i_a = (-1\ \ 0)$

#### 2x Selection

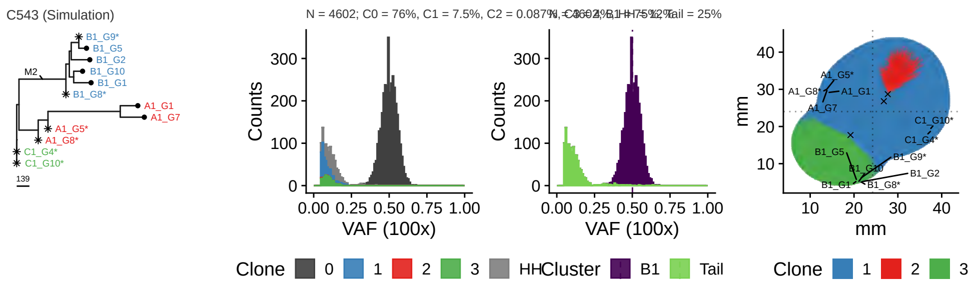

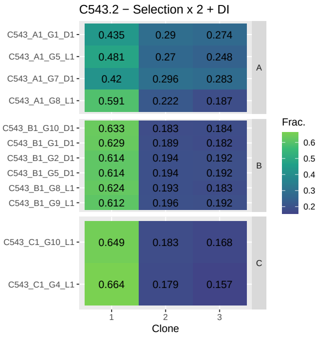

Parameters: $\lambda = (1\ \ 7.39\ \ 20.4)$, $\mu = (0\ \ 0\ \ 0)$, $d_{push} = (189\ \ 189\ \ 189)$, $m = (0\ \ 0\ \ 0)$, $t_s = (0\ \ 22300\ \ 44900)$, $i_a = (-1\ \ 0\ \ 0)$

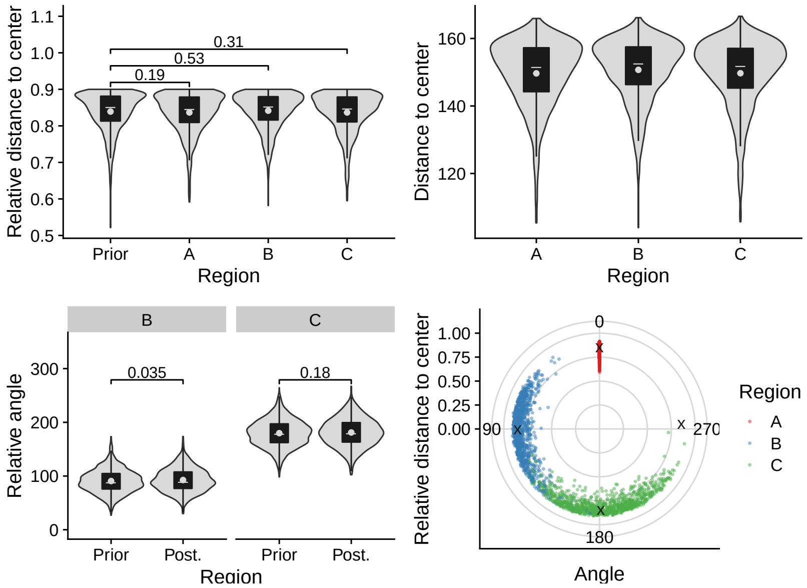
Relative distance to center
0.31
0.53
0.19
Prior
A
B
C
Region
Distance to center
Relative angle
0.035
0.18
Post.
Angle
Region
A
B
C

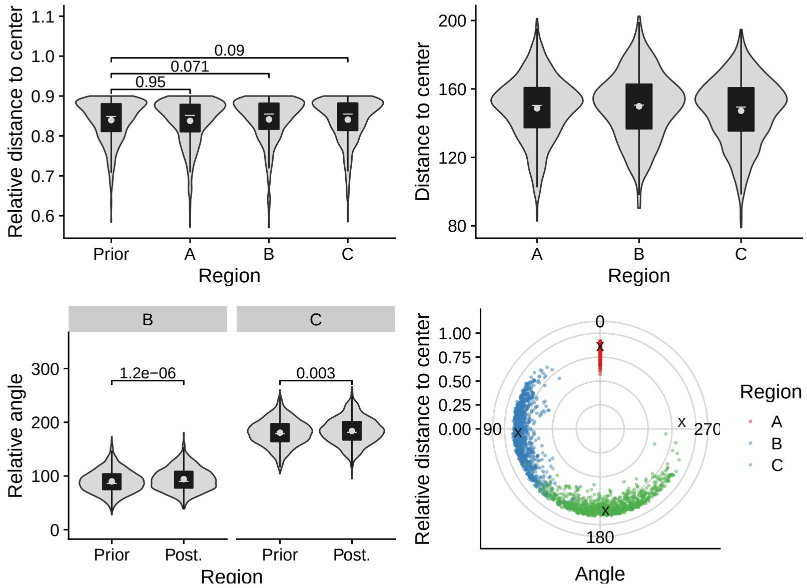
Relative distance to center
0.09
0.071
0.95
Prior
A
B
C
Region
Distance to center
Relative angle
1.2e−06
0.003
Post.
Angle
Region
A
B
C

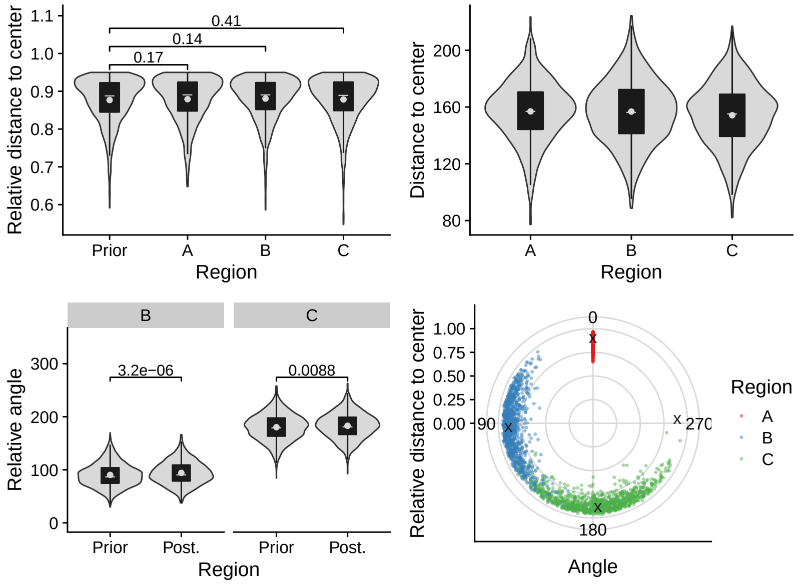
Relative distance to center
0.41
0.14
0.17
Prior
A
B
C
Region
Distance to center
Relative angle
3.2e−06
0.0088
Post.
Angle
Region
A
B
C

# 2.17 C544

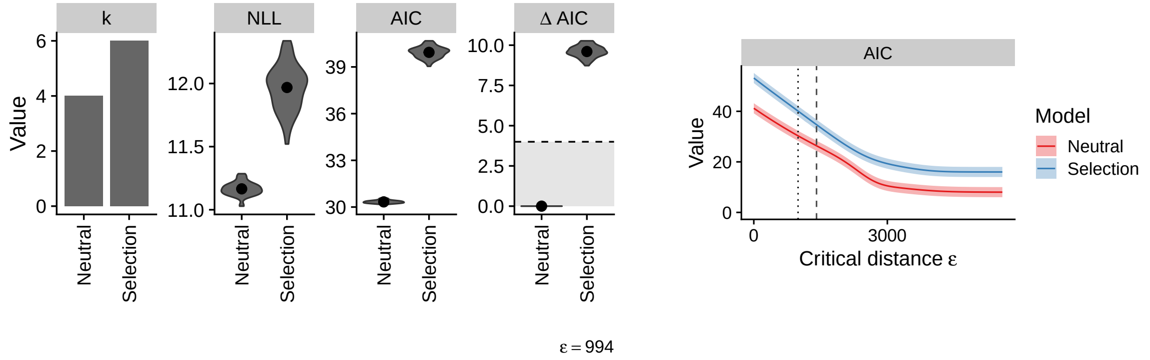

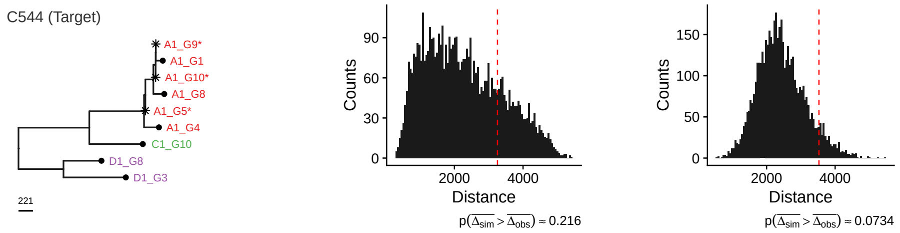

#### Neutral

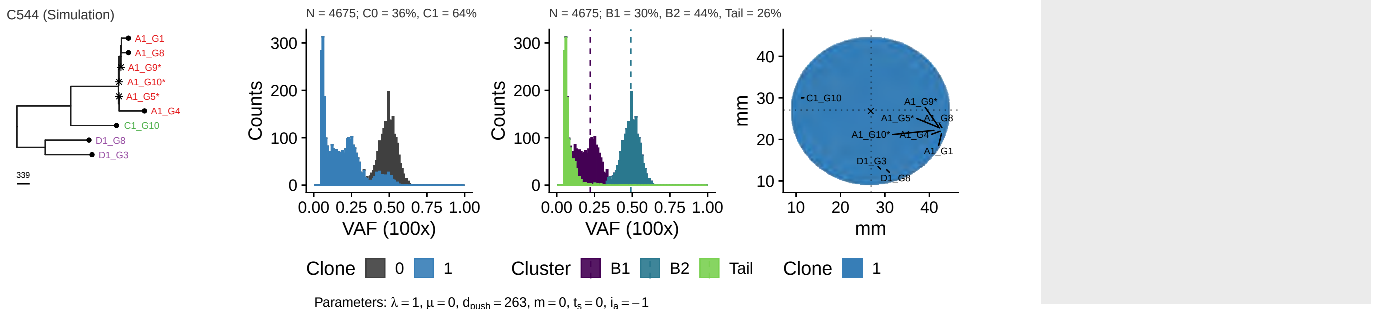

#### Selection

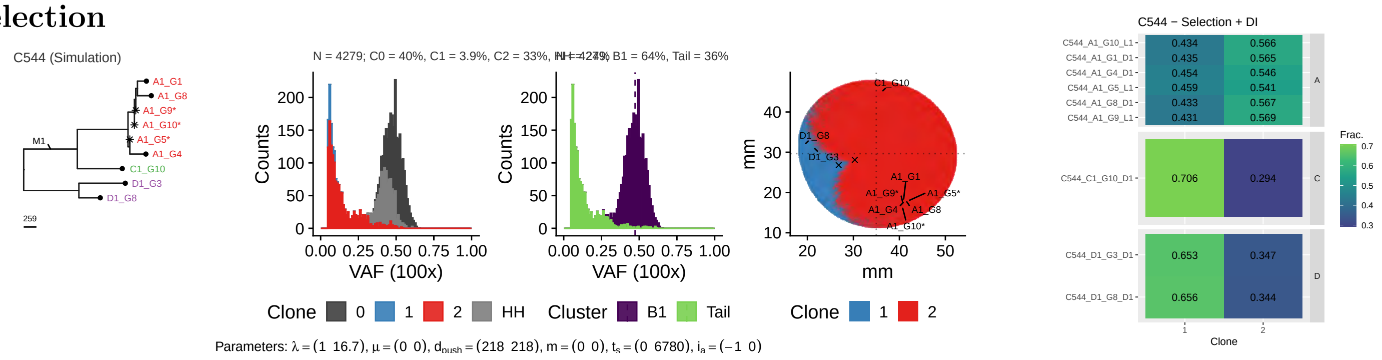

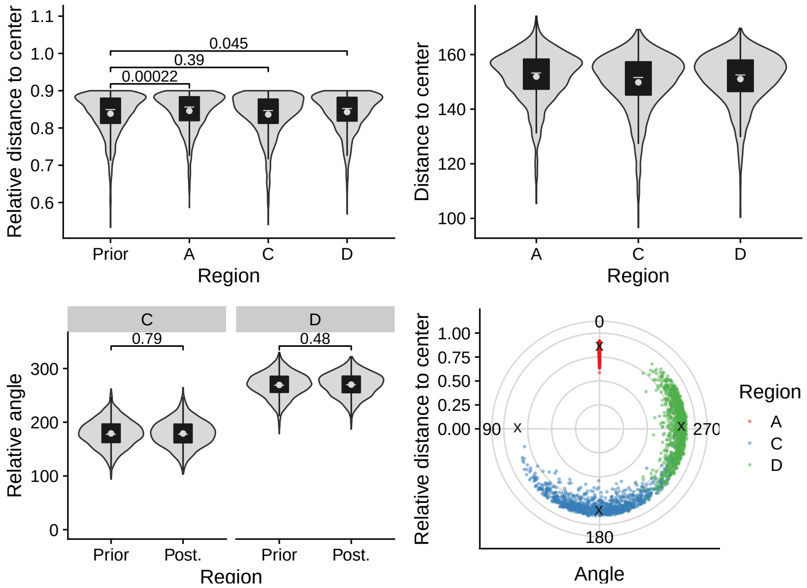
Relative distance to center
0.045
0.39
0.00022
Prior
A
C
D
Region
Distance to center
Region
C
D
0.79
0.48
Relative angle
Prior
Post.
Region
Relative distance to center
Angle
Region
A
C
D

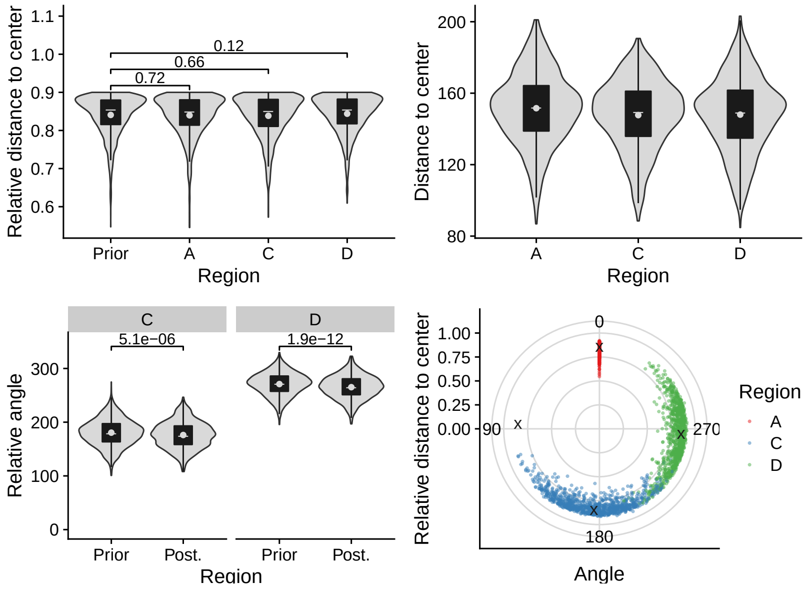
Relative distance to center
0.12
0.66
0.72
Prior
A
C
D
Region
Distance to center
Region
C
D
5.1e−06
1.9e−12
Relative angle
Prior
Post.
Region
Relative distance to center
Angle
Region
A
C
D

## 2.18 C548

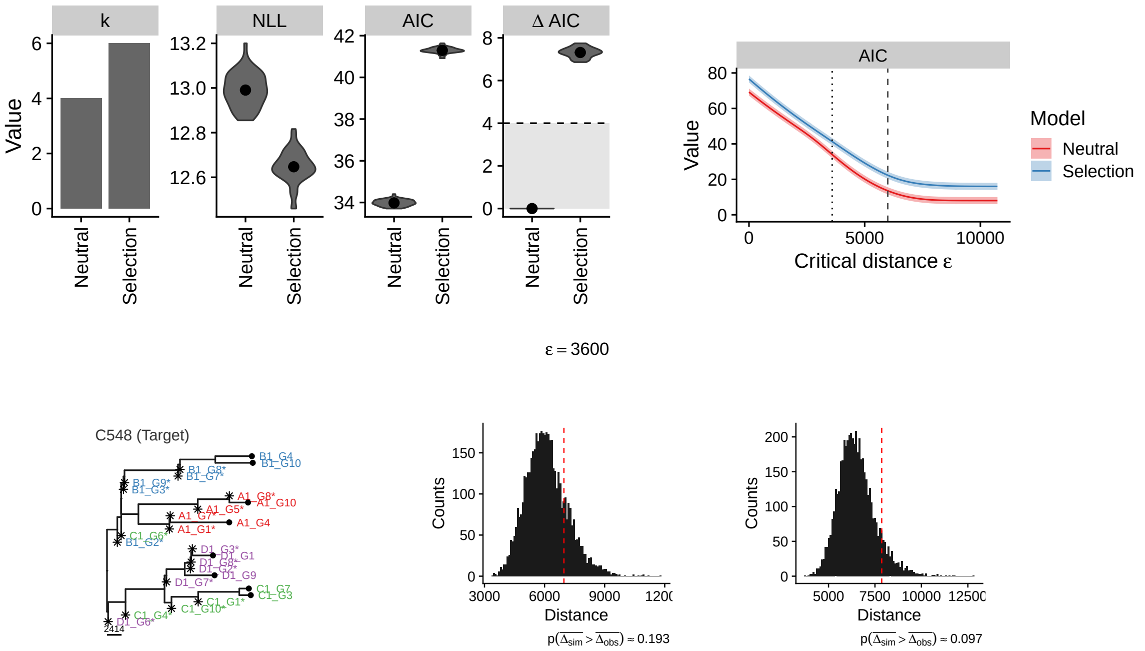

#### Neutral

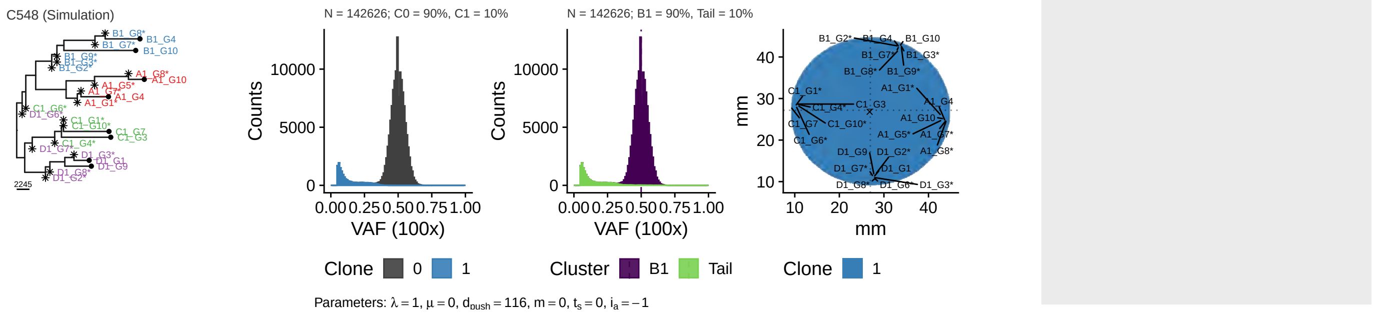

#### Selection

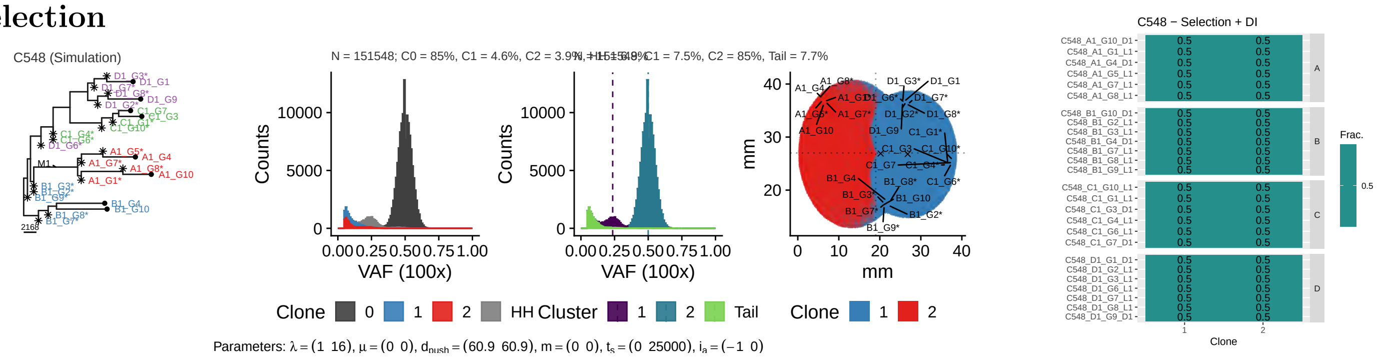

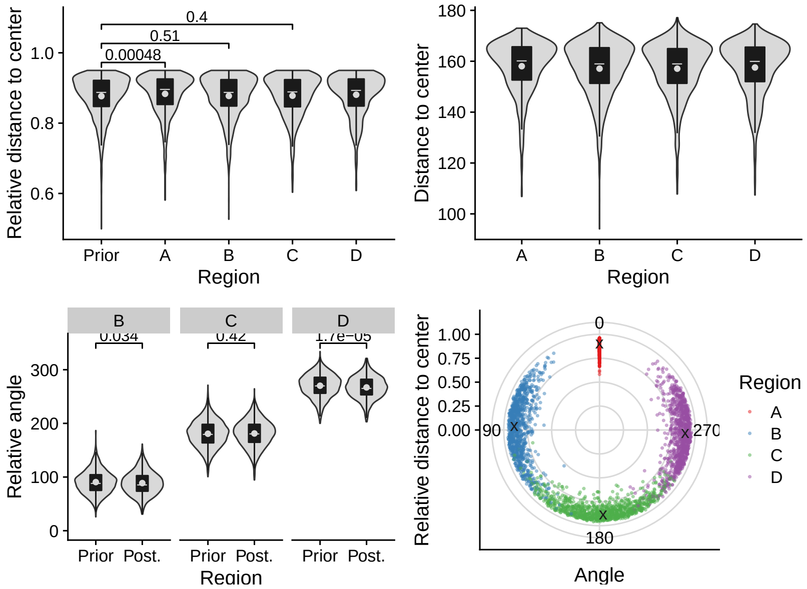
Relative distance to center
0.4
0.51
0.00048
1.0
0.8
0.6
Prior
A
B
C
D
Region
Distance to center
180
160
140
120
100
A
B
C
D
Region
B
C
D
0.034
0.42
1.7e−05
Relative angle
300
200
100
0
Prior Post.
Prior Post.
Prior Post.
Region
Relative distance to center
1.00
0.75
0.50
0.25
0.00
0
90
270
180
Angle
Region
A
B
C
D

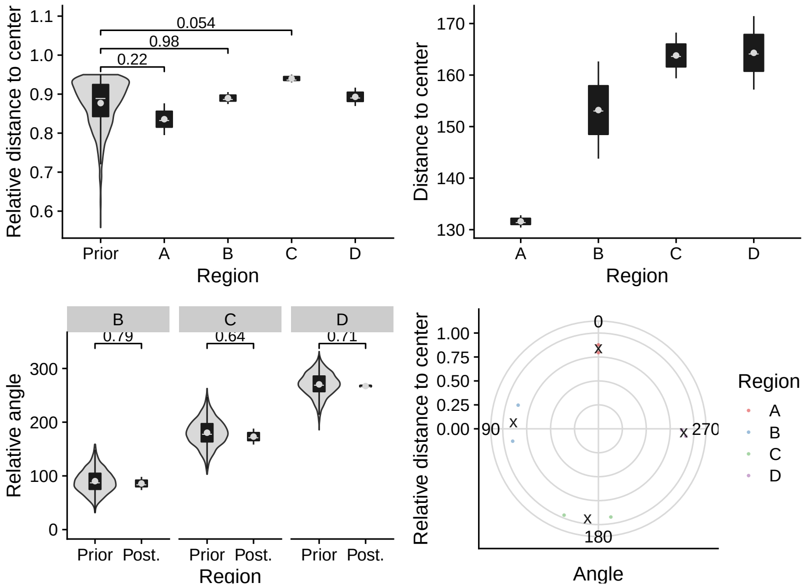
Relative distance to center
0.054
0.98
0.22
1.1
1.0
0.9
0.8
0.7
0.6
Prior
A
B
C
D
Region
Distance to center
170
160
150
140
130
A
B
C
D
Region
B
C
D
0.79
0.64
0.71
Relative angle
300
200
100
0
Prior Post.
Prior Post.
Prior Post.
Region
Relative distance to center
1.00
0.75
0.50
0.25
0.00
0
90
270
180
Angle
Region
A
B
C
D

## 2.19 C549

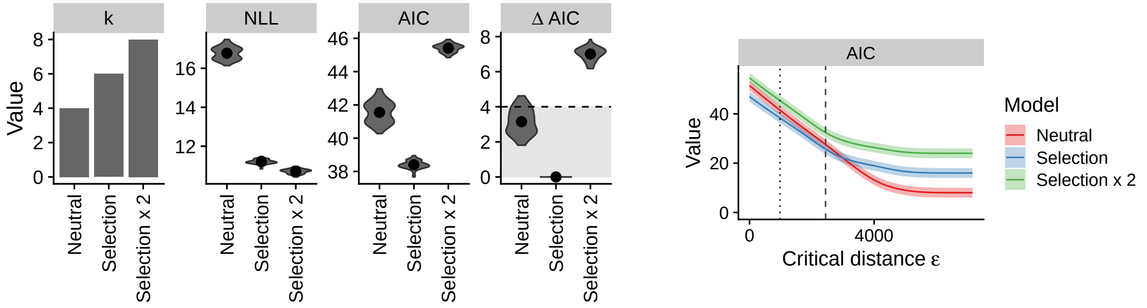

ε = 976

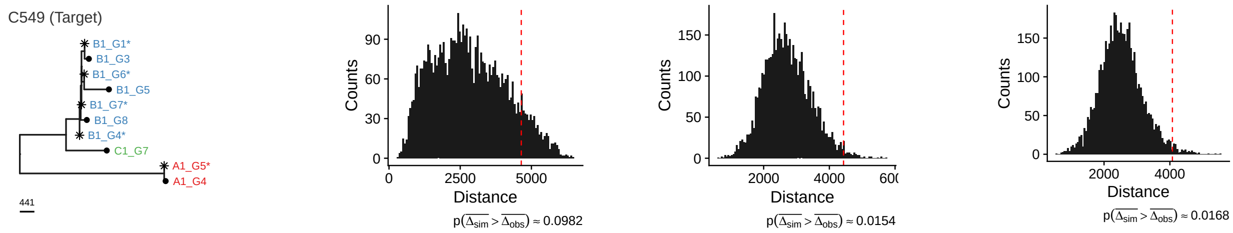

#### Neutral

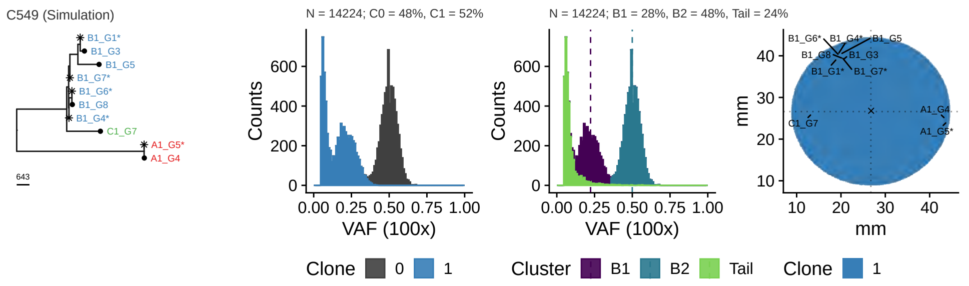

Parameters: $\lambda = 1$, $\mu = 0$, $d_{push} = 217$, $m = 0$, $t_s = 0$, $i_a = -1$

#### Selection

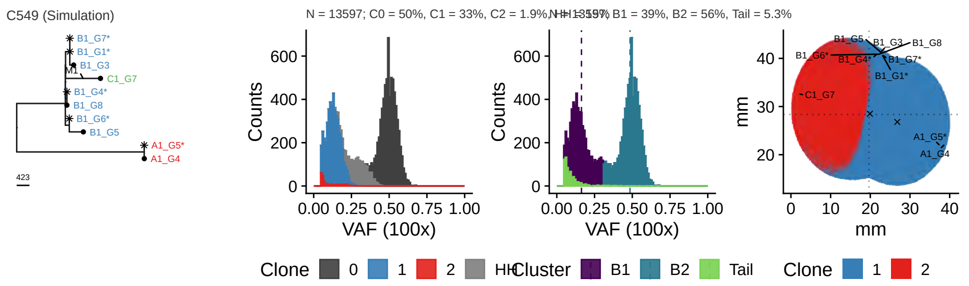

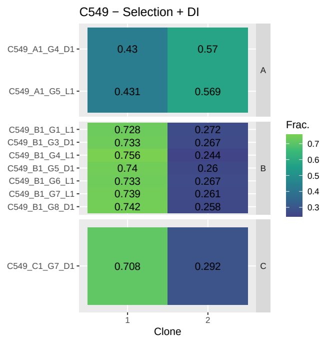

Parameters: $\lambda = (1\ \ 17.9)$, $\mu = (0\ \ 0)$, $d_{push} = (266\ \ 266)$, $m = (0\ \ 0)$, $t_s = (0\ \ 25500)$, $i_a = (-1\ \ 0)$

#### 2x Selection

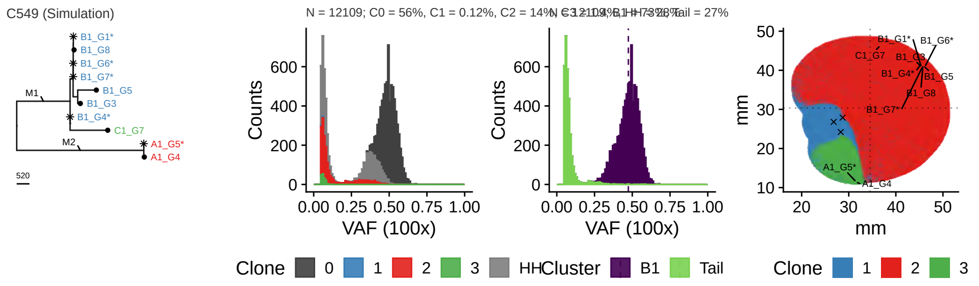

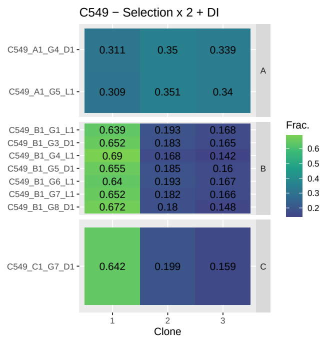

Parameters: $\lambda = (1\ \ 20\ \ 20.6)$, $\mu = (0\ \ 0\ \ 0)$, $d_{push} = (49.2\ \ 49.2\ \ 49.2)$, $m = (0\ \ 0\ \ 0)$, $t_s = (0\ \ 7620\ \ 7780)$, $i_a = (-1\ \ 0\ \ 0)$

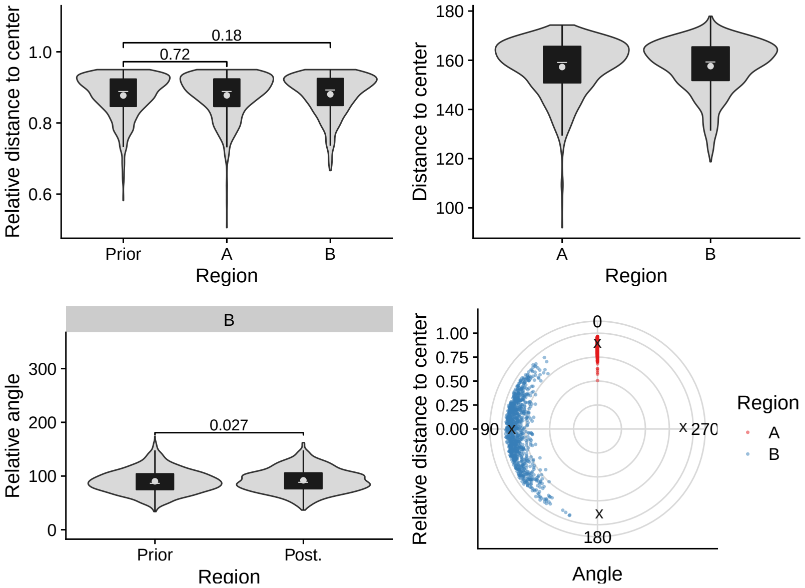
Relative distance to center
1.0
0.8
0.6
0.18
0.72
Prior
A
B
Region
Distance to center
180
160
140
120
100
A
B
Region
B
Relative angle
300
200
100
0
0.027
Prior
Post.
Region
Relative distance to center
1.00
0.75
0.50
0.25
0.00
0
90
270
180
Angle
Region
A
B

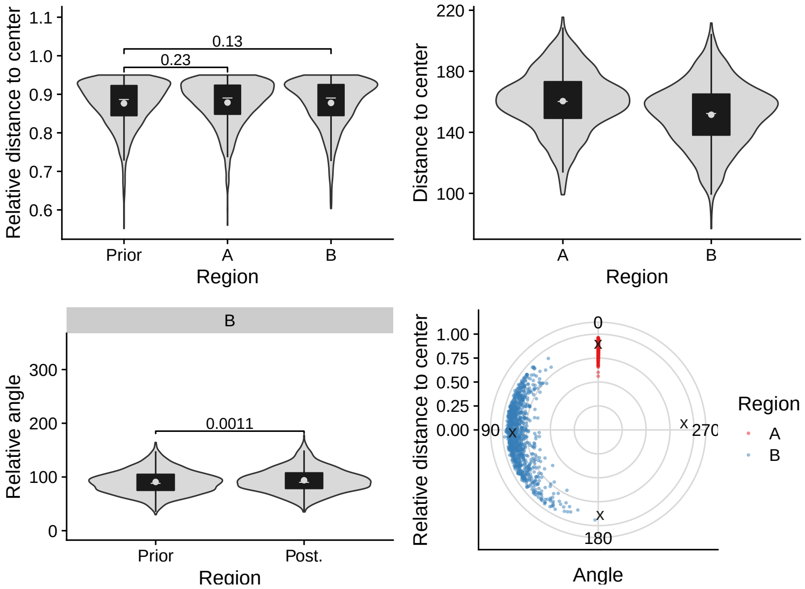
Relative distance to center
1.1
1.0
0.9
0.8
0.7
0.6
0.13
0.23
Prior
A
B
Region
Distance to center
220
180
140
100
A
B
Region
B
Relative angle
300
200
100
0
0.0011
Prior
Post.
Region
Relative distance to center
1.00
0.75
0.50
0.25
0.00
0
90
270
180
Angle
Region
A
B

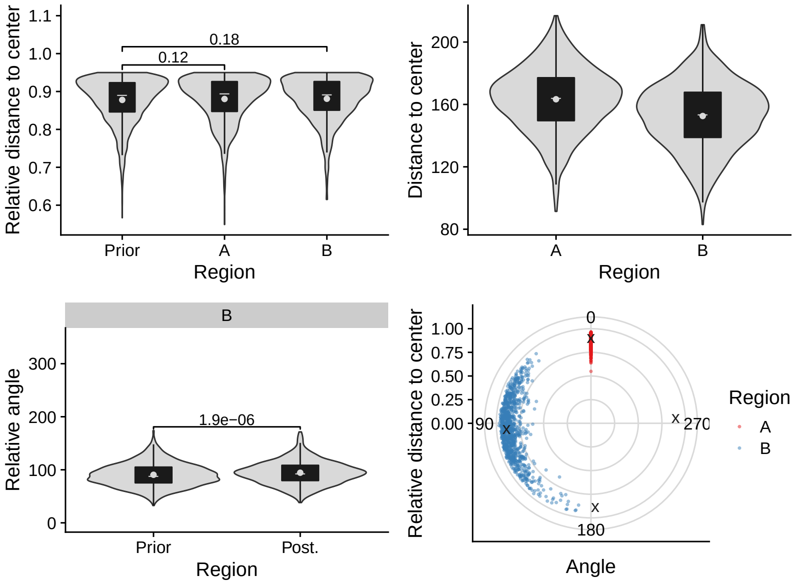
Relative distance to center
1.1
1.0
0.9
0.8
0.7
0.6
0.18
0.12
Prior
A
B
Region
Distance to center
200
160
120
80
A
B
Region
B
Relative angle
300
200
100
0
1.9e−06
Prior
Post.
Region
Relative distance to center
1.00
0.75
0.50
0.25
0.00
0
90
270
180
Angle
Region
A
B

## 2.20 C550

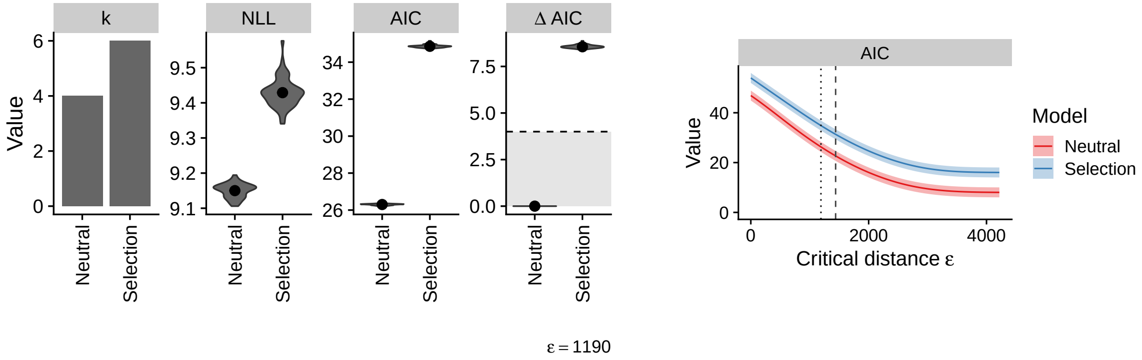

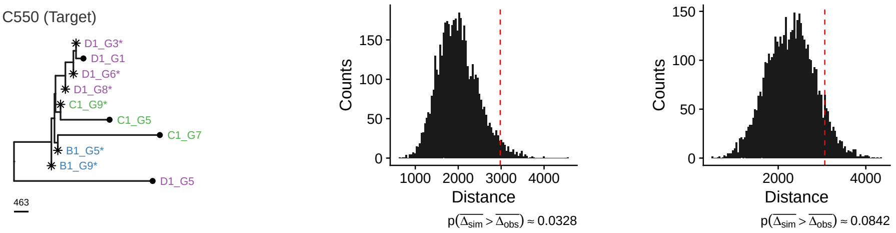

### Neutral

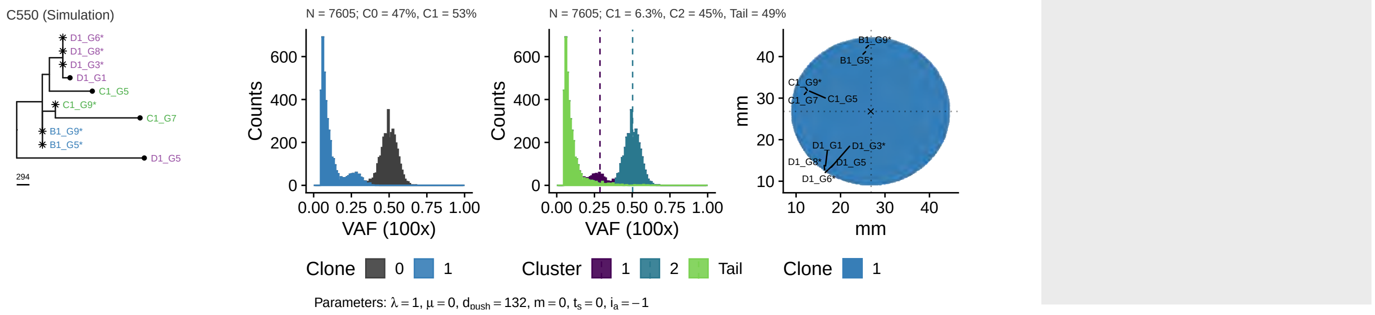

### Selection

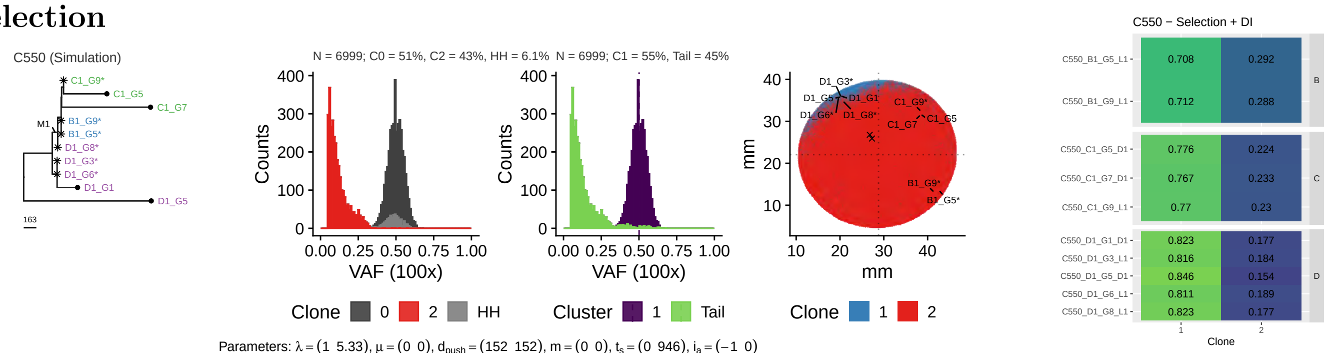

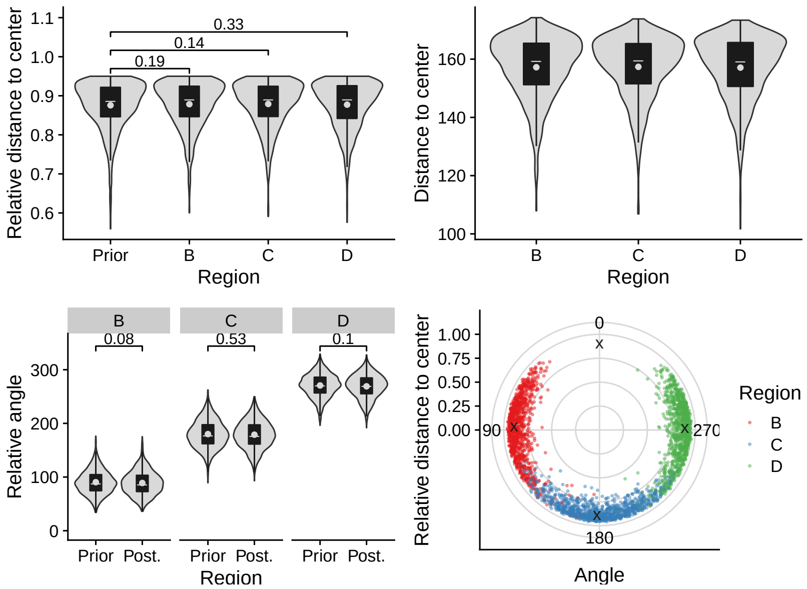
Relative distance to center
0.33
0.14
0.19
Prior B C D
Region
Distance to center
B C D
Region
B C D
0.08 0.53 0.1
Relative angle
Prior Post. Prior Post. Prior Post.
Region
Relative distance to center
0
90
270
180
Angle
Region
B
C
D

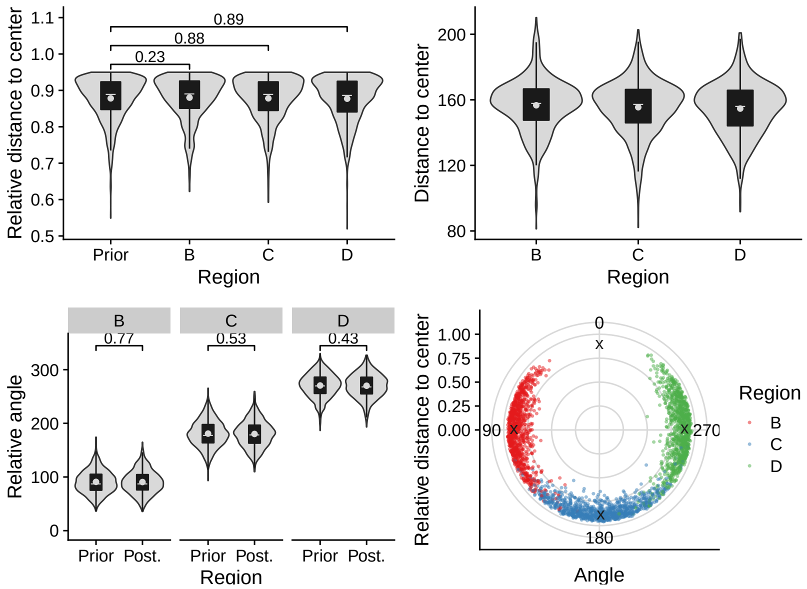
Relative distance to center
0.89
0.88
0.23
Prior B C D
Region
Distance to center
B C D
Region
B C D
0.77 0.53 0.43
Relative angle
Prior Post. Prior Post. Prior Post.
Region
Relative distance to center
0
90
270
180
Angle
Region
B
C
D

## 2.21 C551

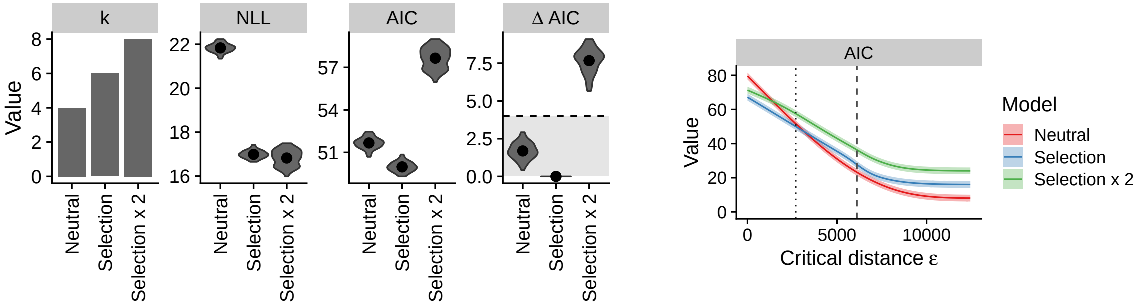

$\varepsilon = 2700$

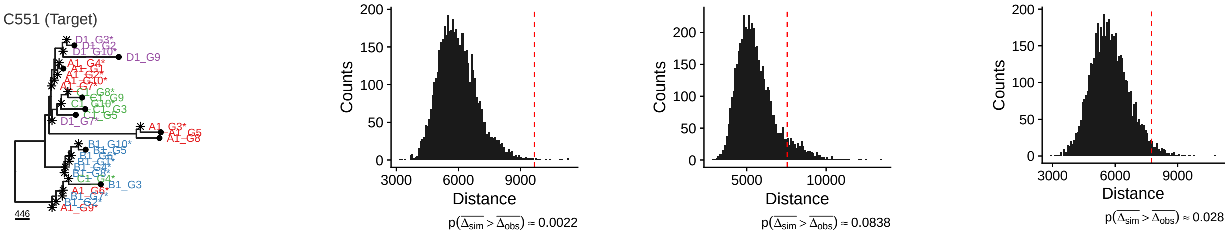

#### Neutral

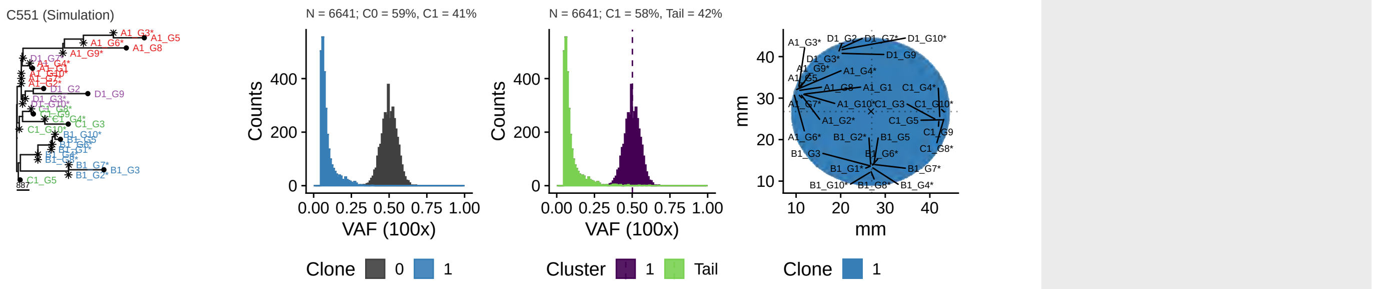

Parameters: $\lambda = 1$, $\mu = 0$, $d_{push} = 239$, $m = 0$, $t_s = 0$, $i_a = -1$

#### Selection

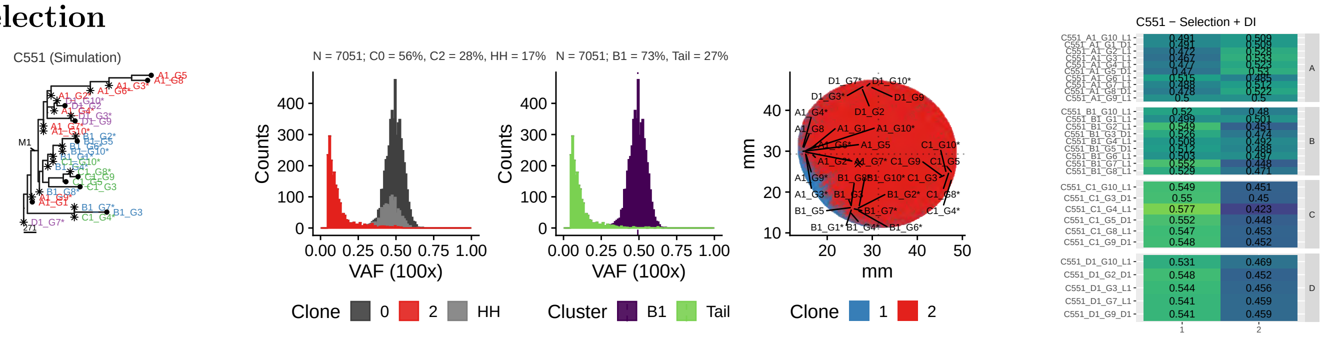

Parameters: $\lambda = (1\ \ 13.1)$, $\mu = (0\ \ 0)$, $d_{push} = (92.9\ \ 92.9)$, $m = (0\ \ 0)$, $t_s = (0\ \ 3390)$, $i_a = (-1\ \ 0)$

#### 2x Selection

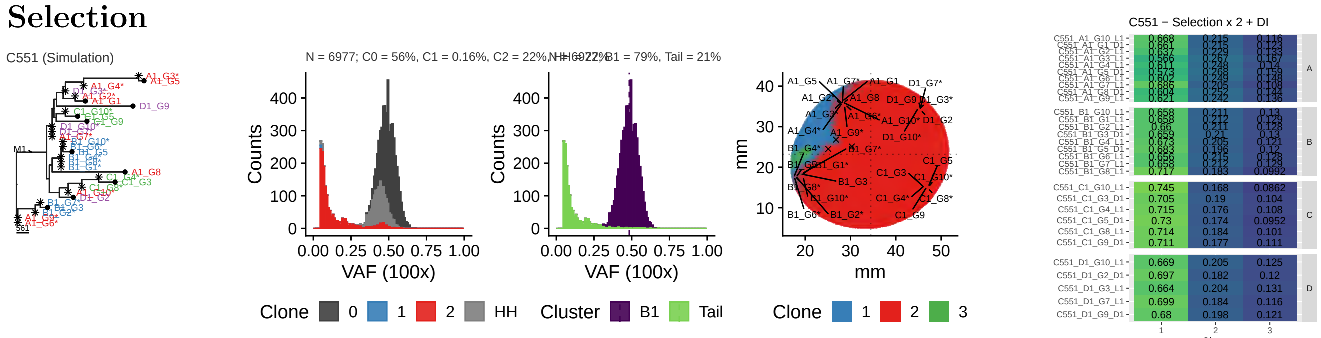

Parameters: $\lambda = (1\ \ 13.7\ \ 11.5)$, $\mu = (0\ \ 0\ \ 0)$, $d_{push} = (238\ \ 238\ \ 238)$, $m = (0\ \ 0\ \ 0)$, $t_s = (0\ \ 5030\ \ 5950)$, $i_a = (-1\ \ 0\ \ 0)$

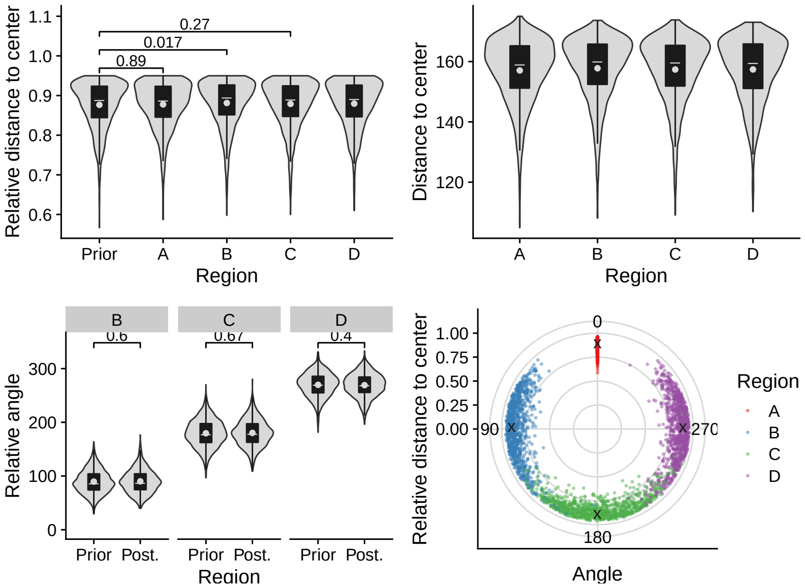
Relative distance to center
0.27
0.017
0.89
Prior A B C D
Region
Distance to center
A B C D
Region
B C D
0.6 0.67 0.4
Relative angle
Prior Post. Prior Post. Prior Post.
Region
Relative distance to center
0 90 180 270
Angle
Region
A
B
C
D

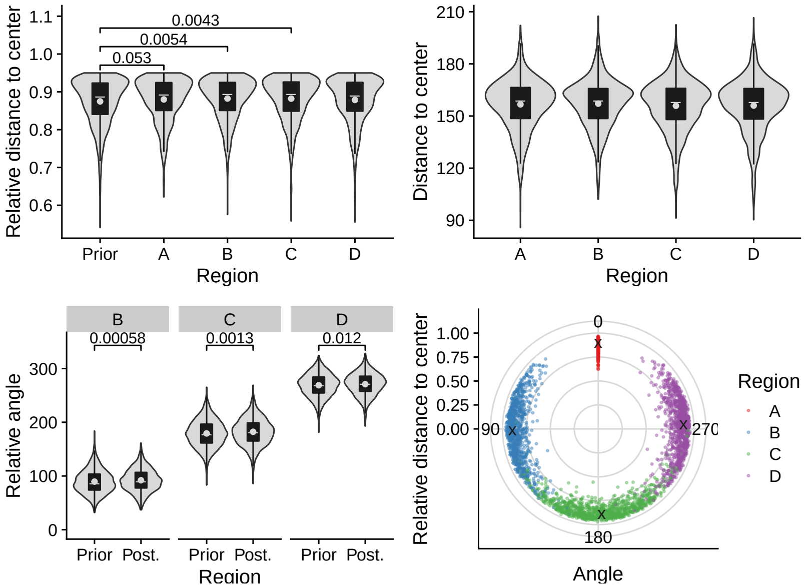
Relative distance to center
0.0043
0.0054
0.053
Prior A B C D
Region
Distance to center
A B C D
Region
B C D
0.00058 0.0013 0.012
Relative angle
Prior Post. Prior Post. Prior Post.
Region
Relative distance to center
0 90 180 270
Angle
Region
A
B
C
D

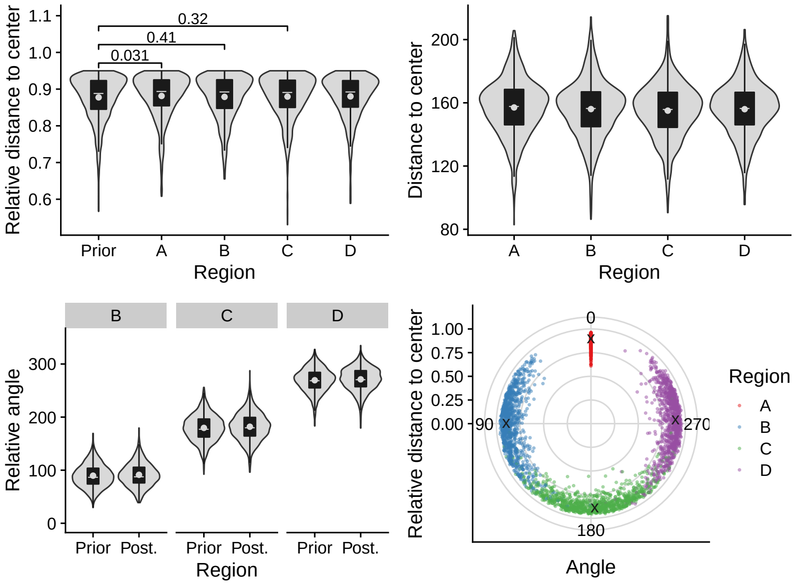
Relative distance to center
0.32
0.41
0.031
Prior A B C D
Region
Distance to center
A B C D
Region
B C D
Relative angle
Prior Post. Prior Post. Prior Post.
Region
Relative distance to center
0 90 180 270
Angle
Region
A
B
C
D

## 2.22 C552

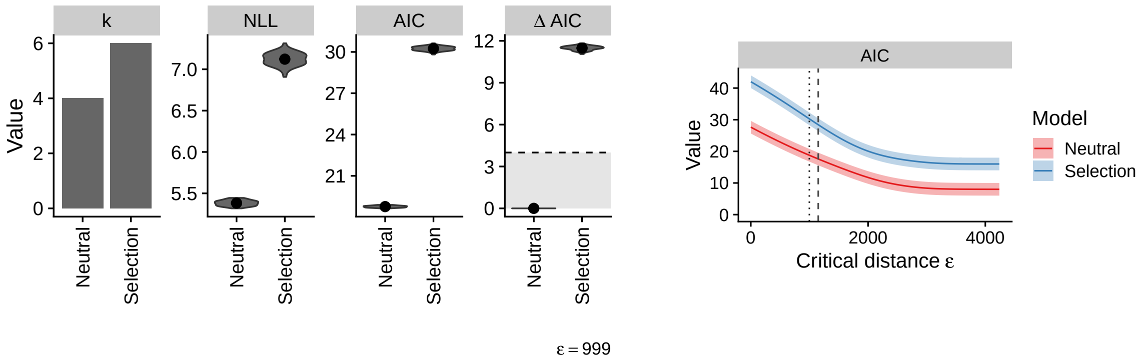

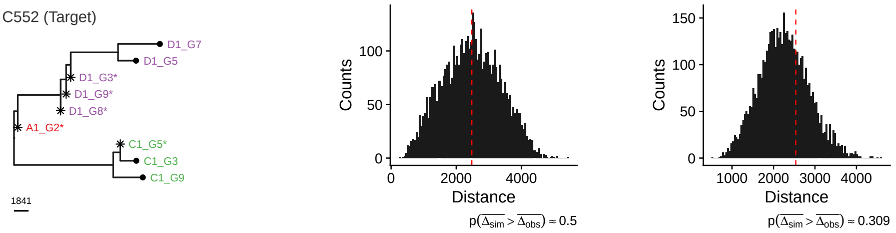

#### Neutral

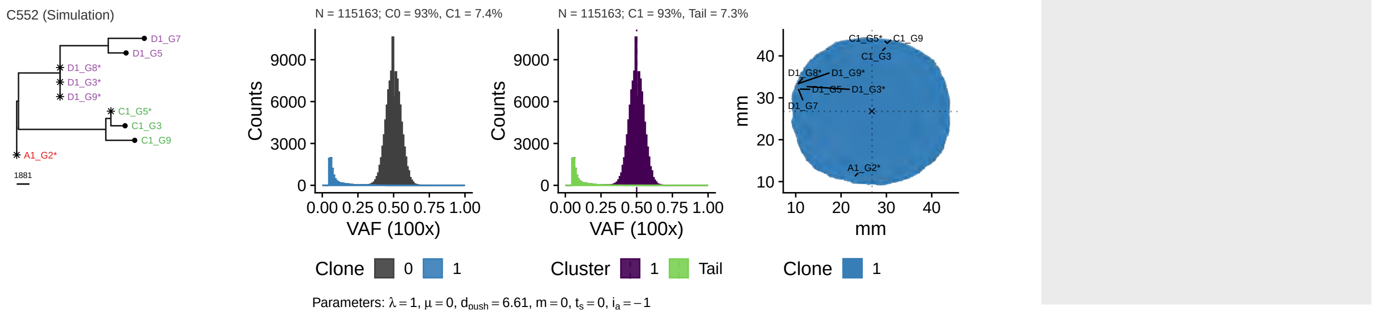

#### Selection

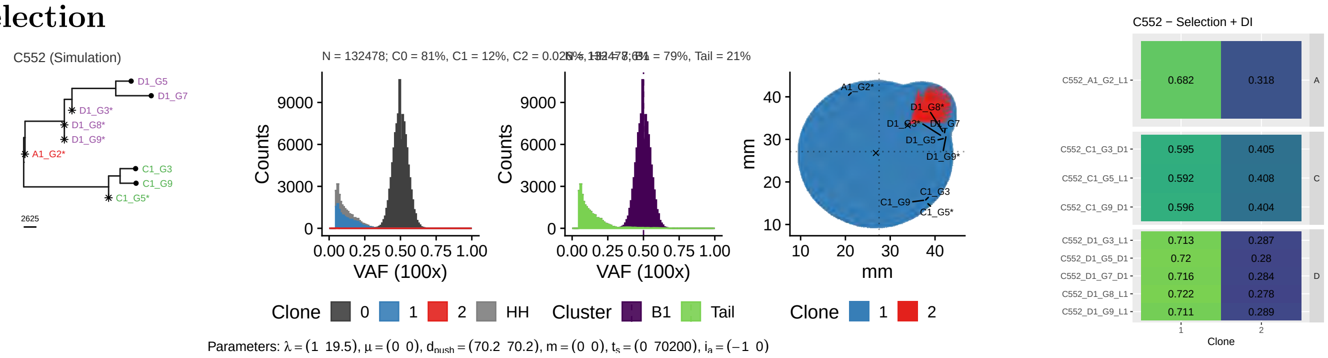

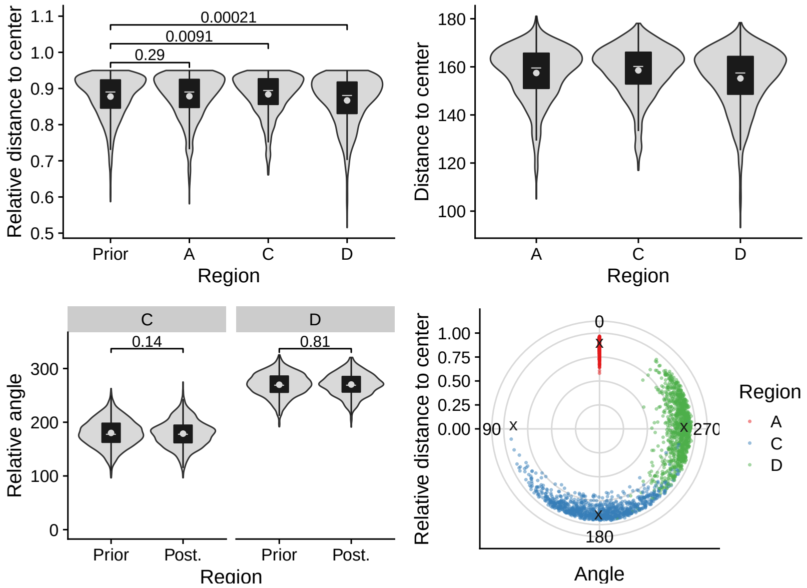
Relative distance to center
0.00021
0.0091
0.29
Prior
A
C
D
Region
Distance to center
A
C
D
Region
C
D
0.14
0.81
Relative angle
Prior
Post.
Prior
Post.
Region
Relative distance to center
0
90
270
180
Angle
Region
A
C
D

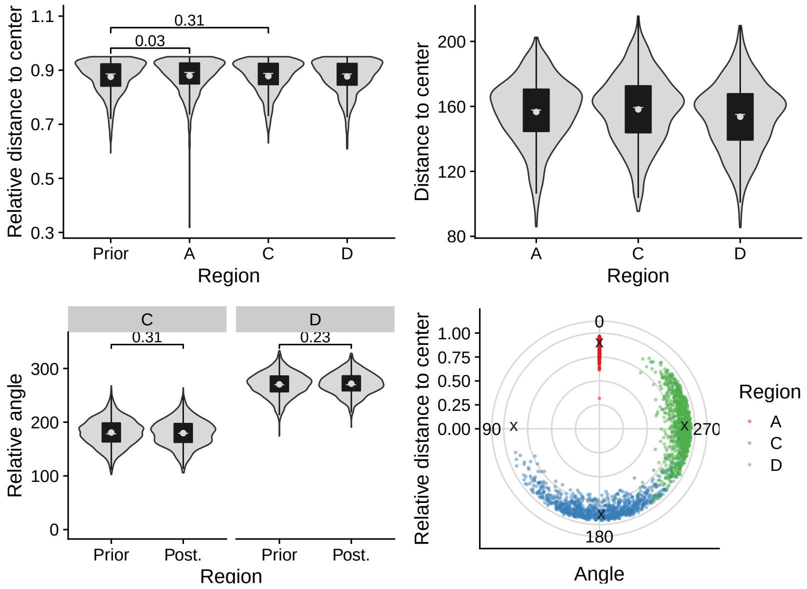
Relative distance to center
0.31
0.03
Prior
A
C
D
Region
Distance to center
A
C
D
Region
C
D
0.31
0.23
Relative angle
Prior
Post.
Prior
Post.
Region
Relative distance to center
0
90
270
180
Angle
Region
A
C
D

## 2.23 C554

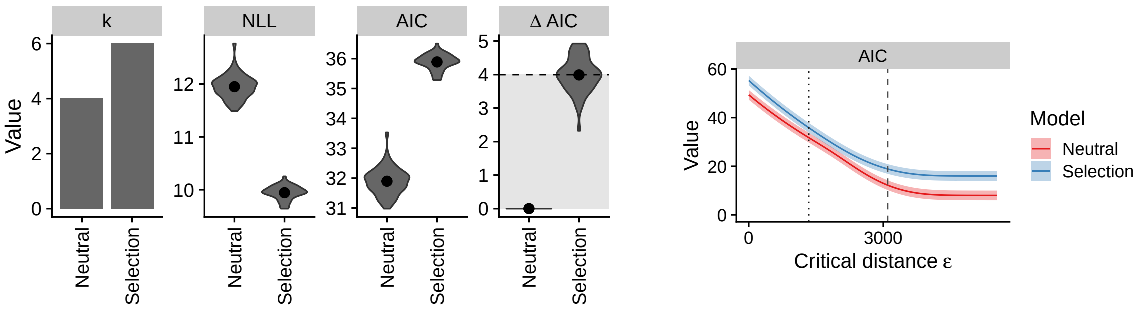

ε = 1340

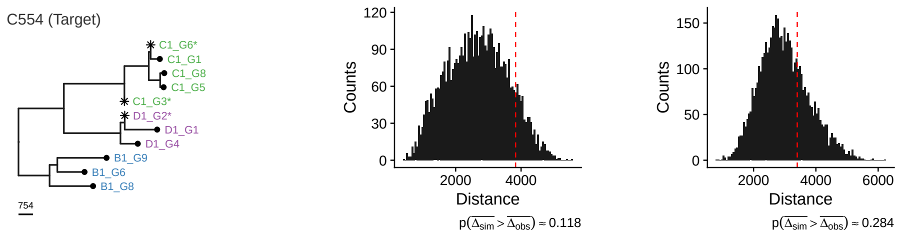

##### Neutral

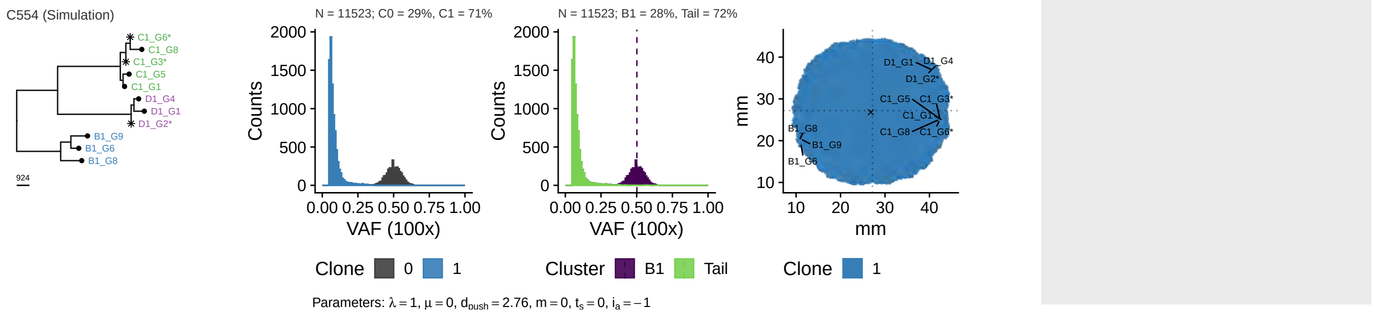

##### Selection

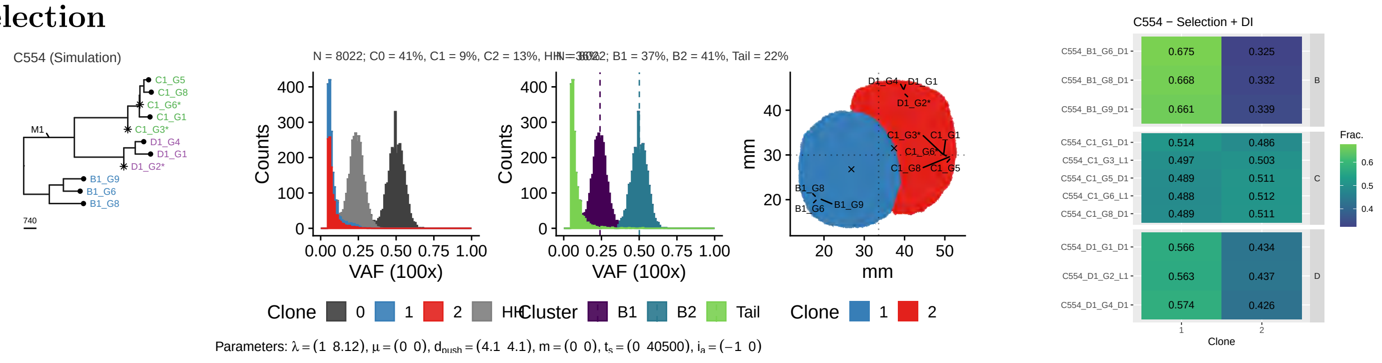

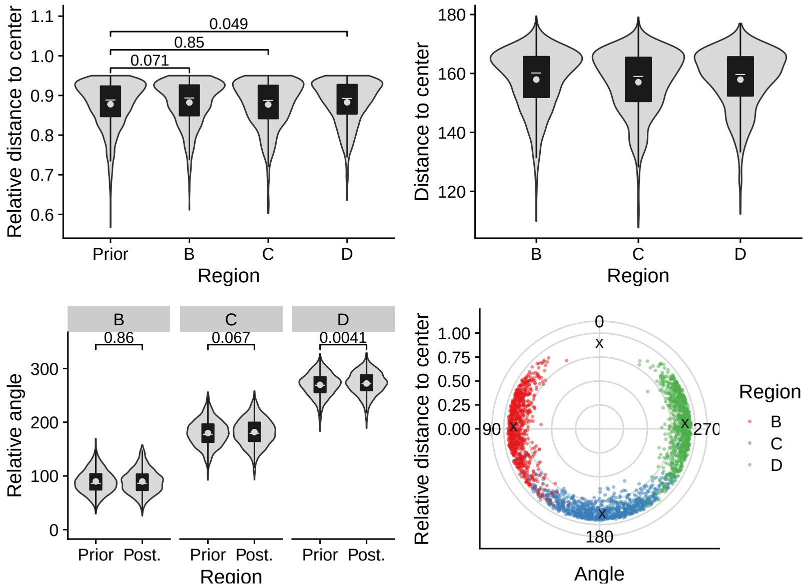
0.049
0.85
0.071
Relative distance to center
1.1
1.0
0.9
0.8
0.7
0.6
Prior
B
C
D
Region
Distance to center
180
160
140
120
B
C
D
Region
B
C
D
0.86
0.067
0.0041
Relative angle
300
200
100
0
Prior Post.
Prior Post.
Prior Post.
Region
Relative distance to center
1.00
0.75
0.50
0.25
0.00
0
90
270
180
Angle
Region
B
C
D

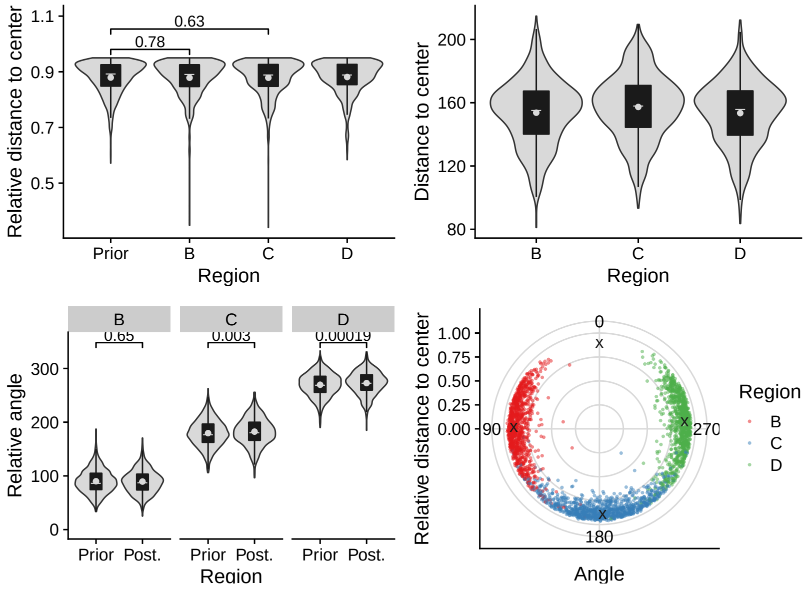
0.63
0.78
Relative distance to center
1.1
0.9
0.7
0.5
Prior
B
C
D
Region
Distance to center
200
160
120
80
B
C
D
Region
B
C
D
0.65
0.003
0.00019
Relative angle
300
200
100
0
Prior Post.
Prior Post.
Prior Post.
Region
Relative distance to center
1.00
0.75
0.50
0.25
0.00
0
90
270
180
Angle
Region
B
C
D

## 2.24 C555

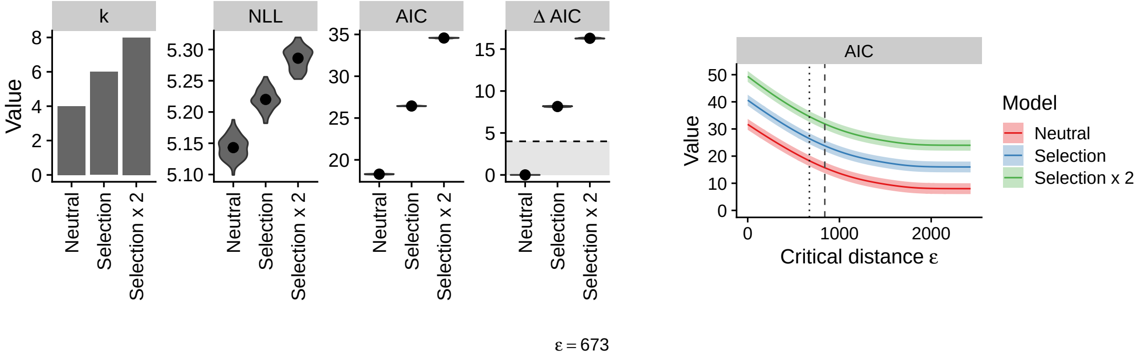

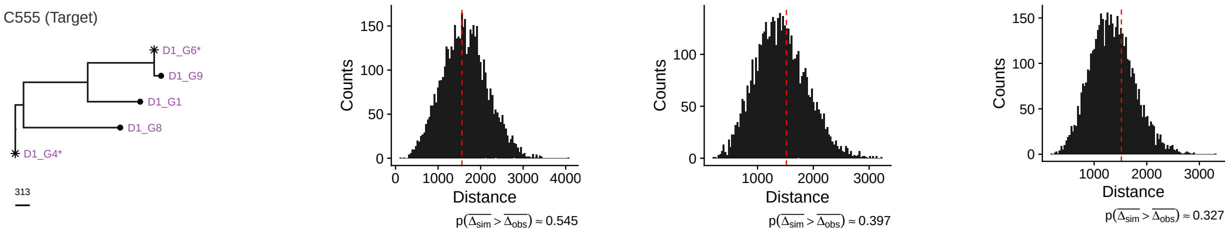

#### Neutral

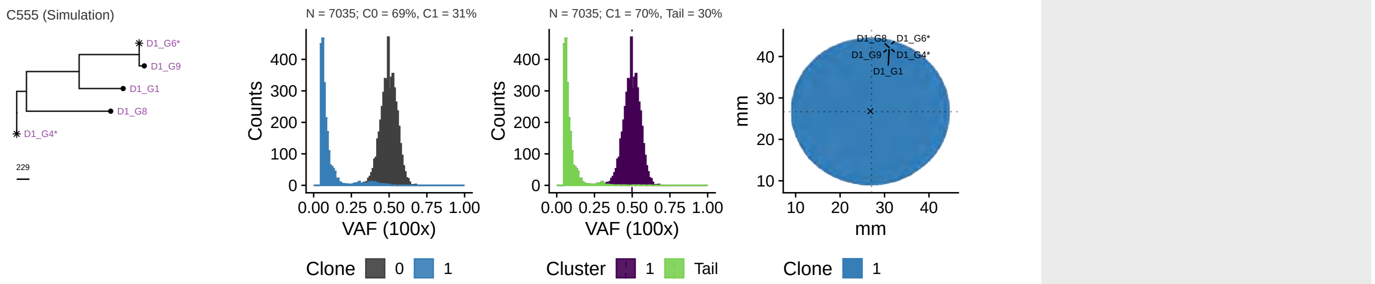

Parameters: $\lambda = 1$, $\mu = 0$, $d_{push} = 242$, $m = 0$, $t_s = 0$, $i_a = -1$

#### Selection

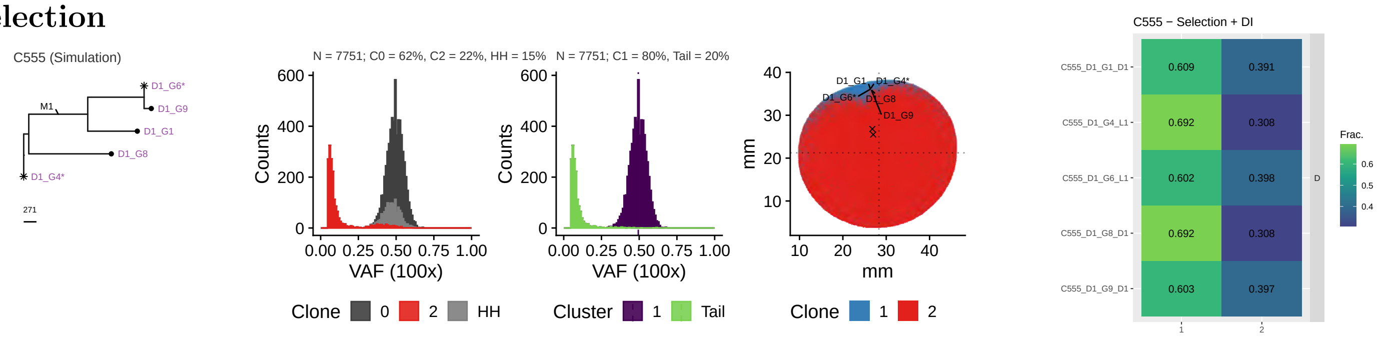

Parameters: $\lambda = (1 \;\; 21.8)$, $\mu = (0 \;\; 0)$, $d_{push} = (94.7 \;\; 94.7)$, $m = (0 \;\; 0)$, $t_s = (0 \;\; 3630)$, $i_a = (-1 \;\; 0)$

#### 2x Selection

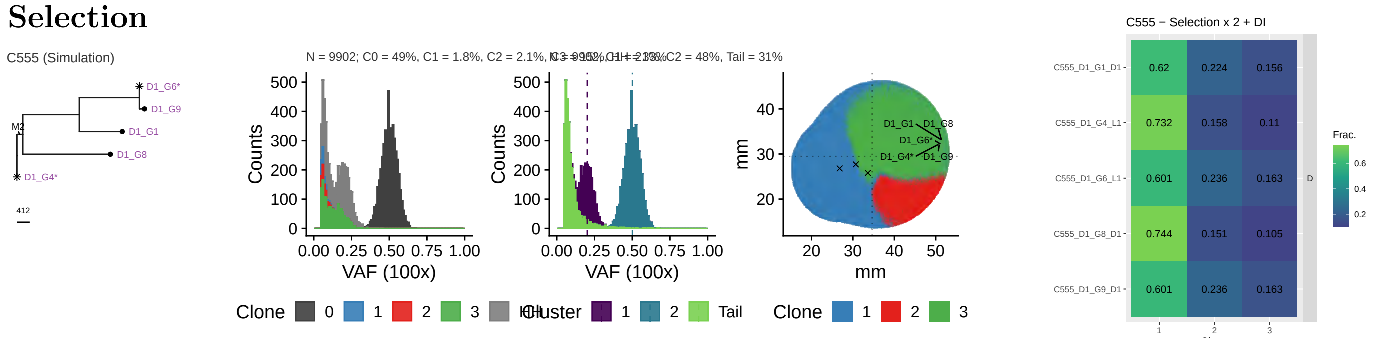

Parameters: $\lambda = (1 \;\; 8.77 \;\; 15.1)$, $\mu = (0 \;\; 0 \;\; 0)$, $d_{push} = (170 \;\; 170 \;\; 170)$, $m = (0 \;\; 0 \;\; 0)$, $t_s = (0 \;\; 15300 \;\; 18800)$, $i_a = (-1 \;\; 0 \;\; 0)$

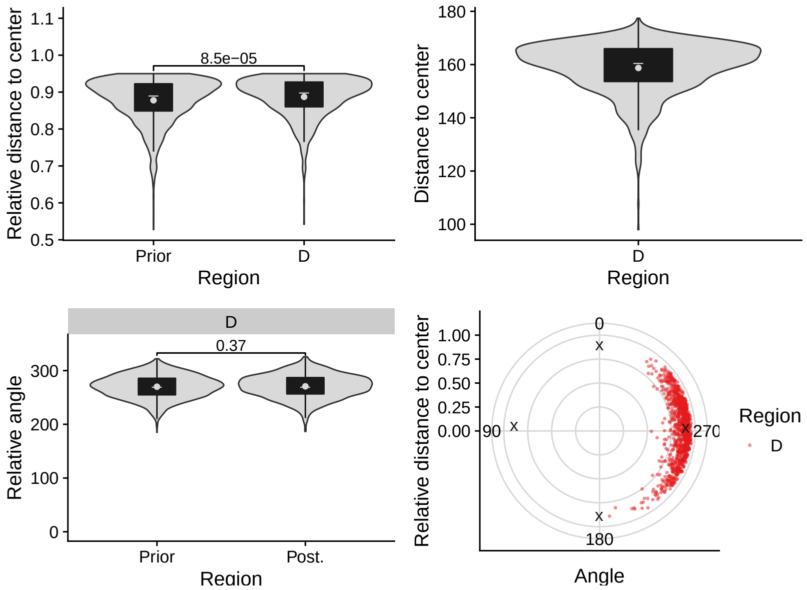
Relative distance to center
8.5e−05
Prior
D
Region
Distance to center
D
Region
D
0.37
Relative angle
Prior
Post.
Region
Relative distance to center
0
90
270
180
Angle
Region
D

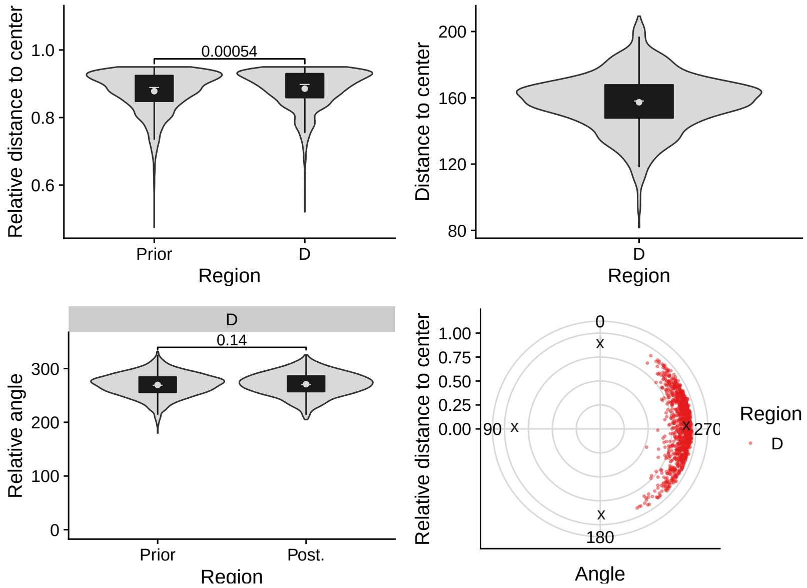
Relative distance to center
0.00054
Prior
D
Region
Distance to center
D
Region
D
0.14
Relative angle
Prior
Post.
Region
Relative distance to center
0
90
270
180
Angle
Region
D

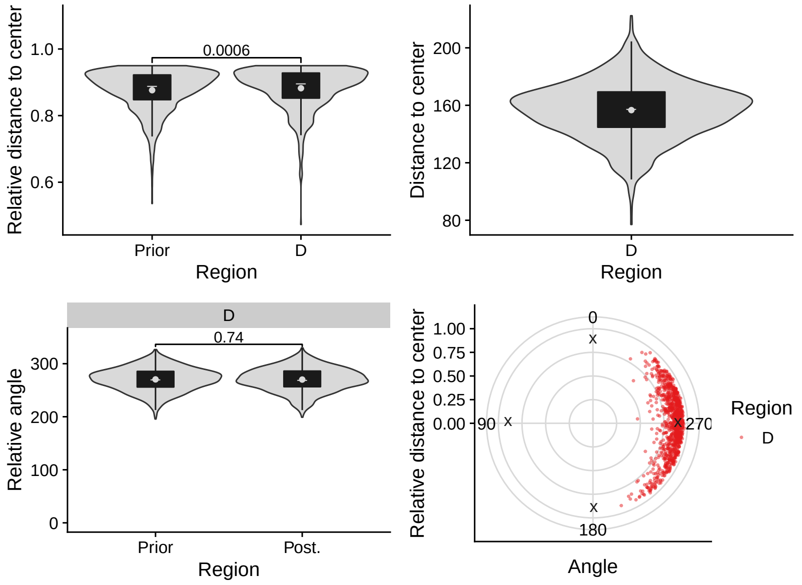
Relative distance to center
0.0006
Prior
D
Region
Distance to center
D
Region
D
0.74
Relative angle
Prior
Post.
Region
Relative distance to center
0
90
270
180
Angle
Region
D

## 2.25 C559

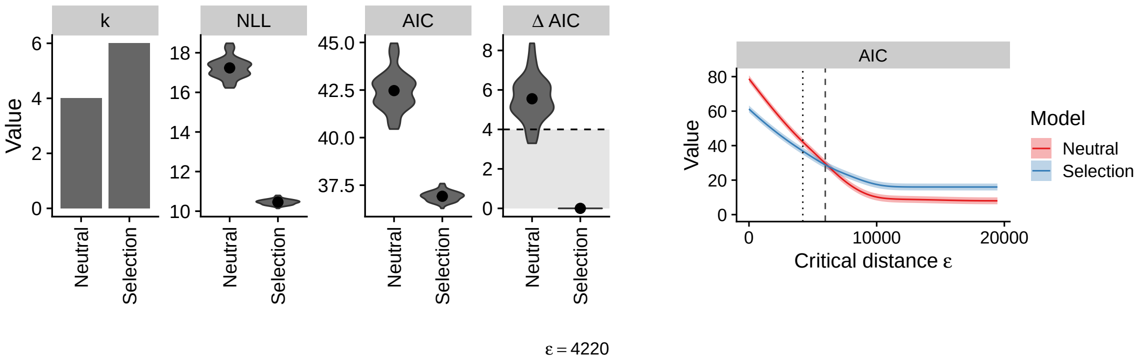

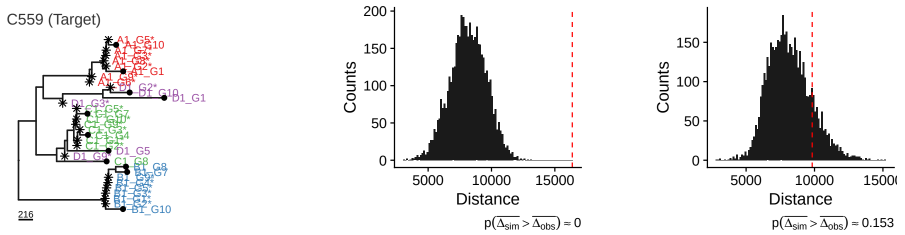

#### Neutral

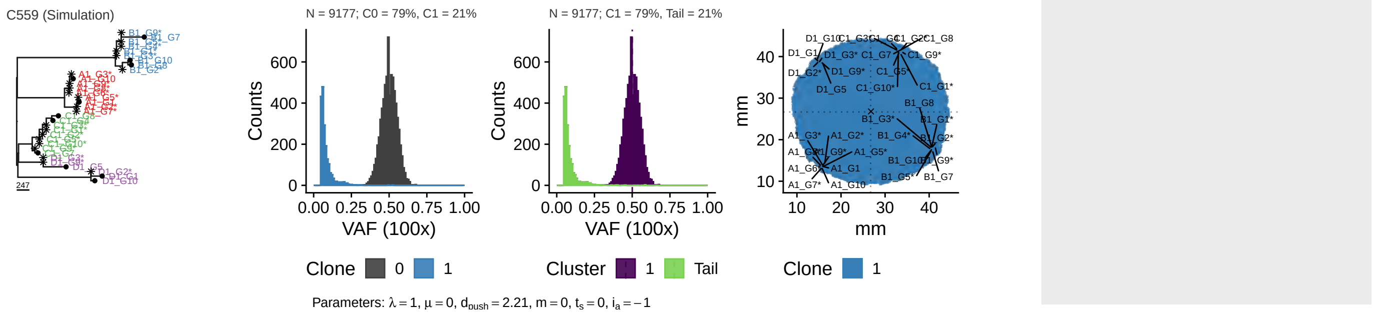

#### Selection

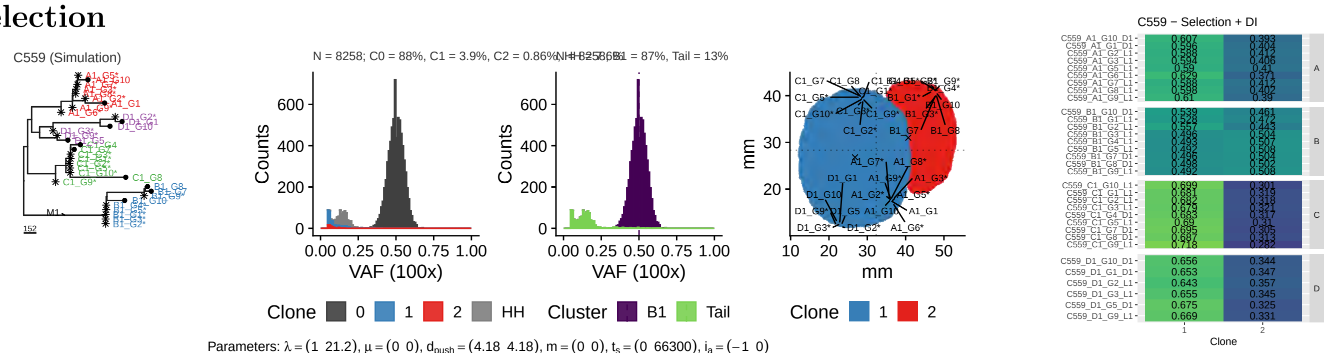

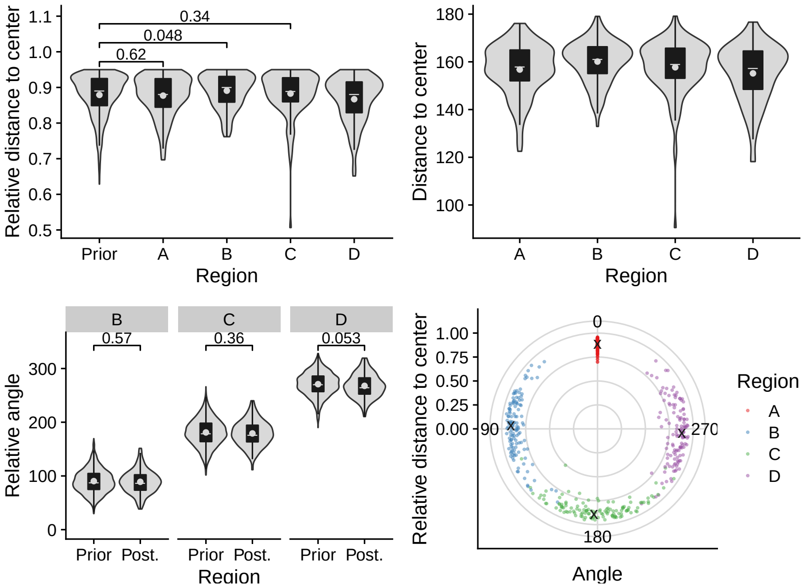
Relative distance to center
1.1
1.0
0.9
0.8
0.7
0.6
0.5
0.34
0.048
0.62
Prior
A
B
C
D
Region
Distance to center
180
160
140
120
100
A
B
C
D
Region
B
C
D
0.57
0.36
0.053
Relative angle
300
200
100
0
Prior Post.
Prior Post.
Prior Post.
Region
Relative distance to center
1.00
0.75
0.50
0.25
0.00
0
90
270
180
Angle
Region
A
B
C
D

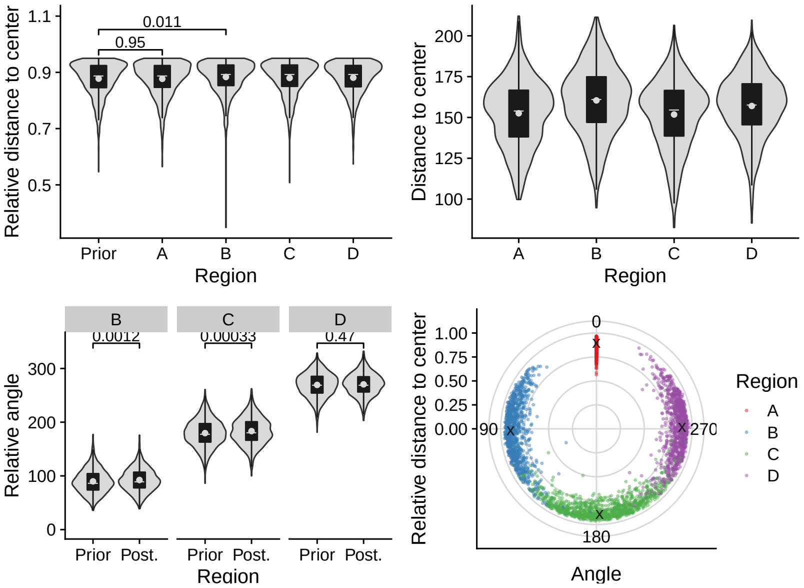
Relative distance to center
1.1
0.9
0.7
0.5
0.011
0.95
Prior
A
B
C
D
Region
Distance to center
200
175
150
125
100
A
B
C
D
Region
B
C
D
0.0012
0.00033
0.47
Relative angle
300
200
100
0
Prior Post.
Prior Post.
Prior Post.
Region
Relative distance to center
1.00
0.75
0.50
0.25
0.00
0
90
270
180
Angle
Region
A
B
C
D

## 2.26 C560

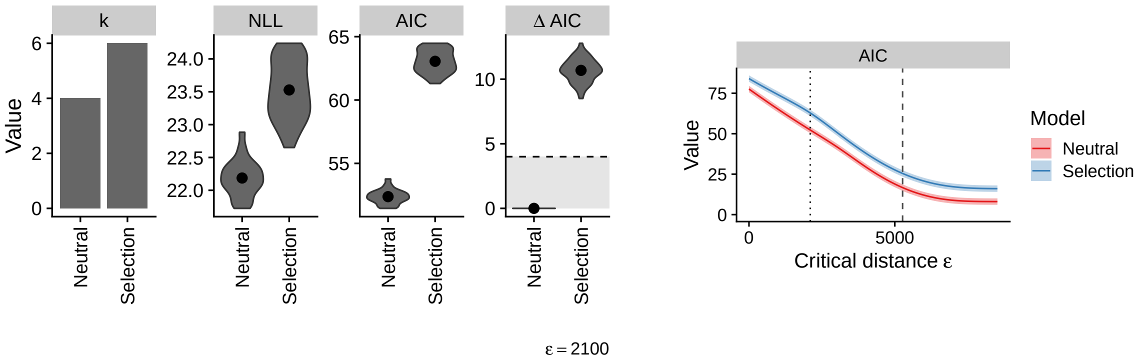

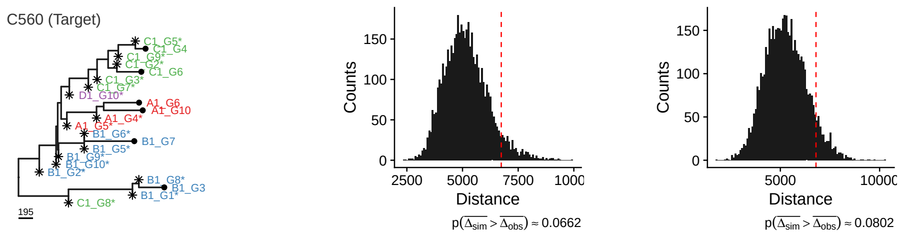

#### Neutral

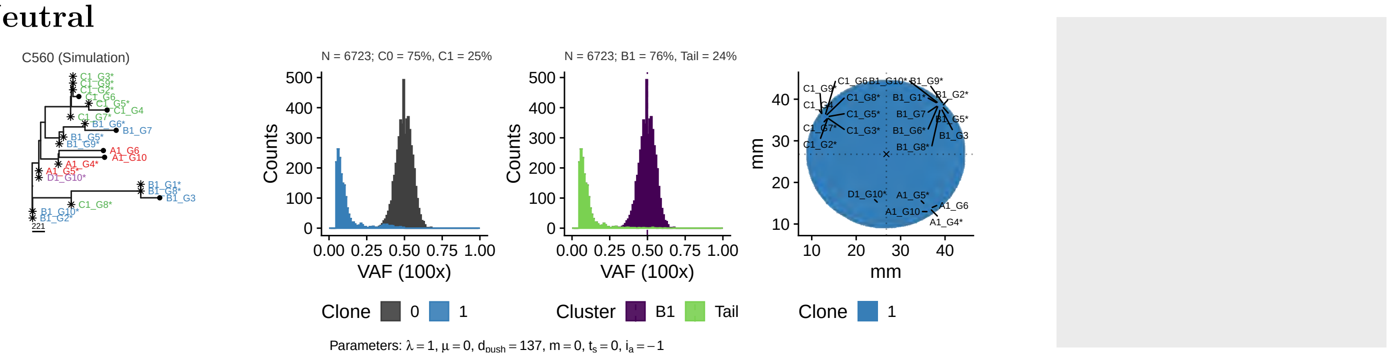

#### Selection

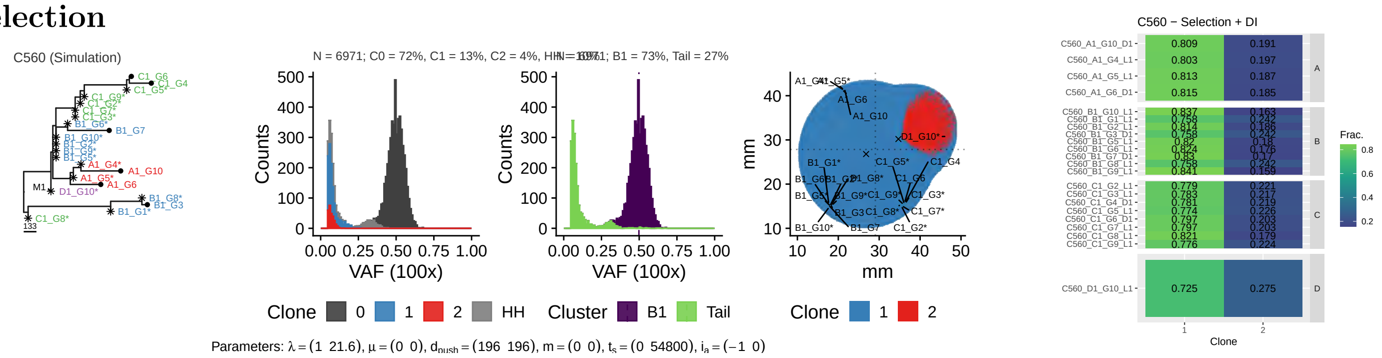

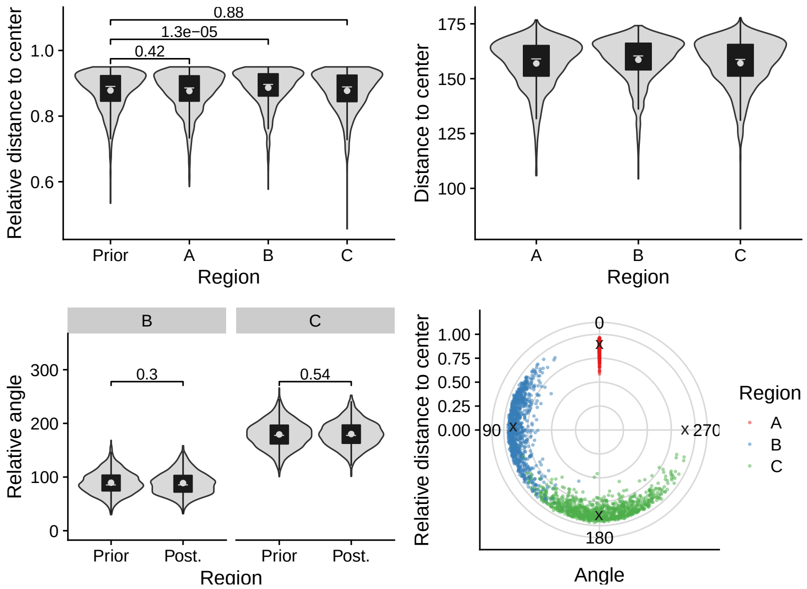
0.88
1.3e−05
0.42
Relative distance to center
Prior
A
B
C
Region
Distance to center
Relative angle
0.3
0.54
Post.
Angle
Relative distance to center
0
90
180
270
Region
A
B
C

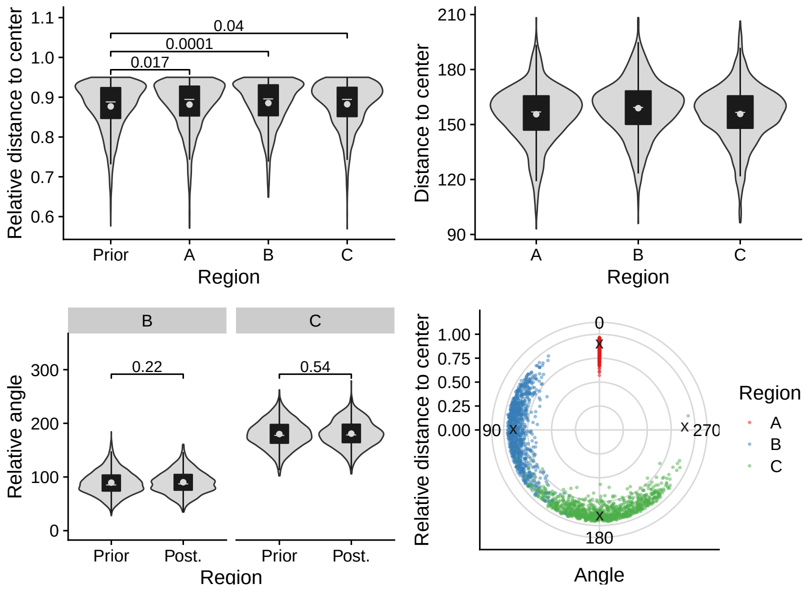
0.04
0.0001
0.017
Relative distance to center
Prior
A
B
C
Region
Distance to center
Relative angle
0.22
0.54
Post.
Angle
Relative distance to center
0
90
180
270
Region
A
B
C

## 2.27 C561

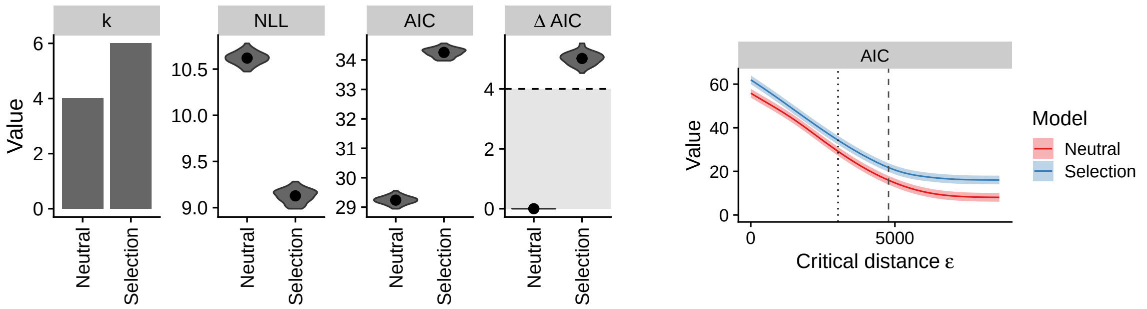

ε = 3030

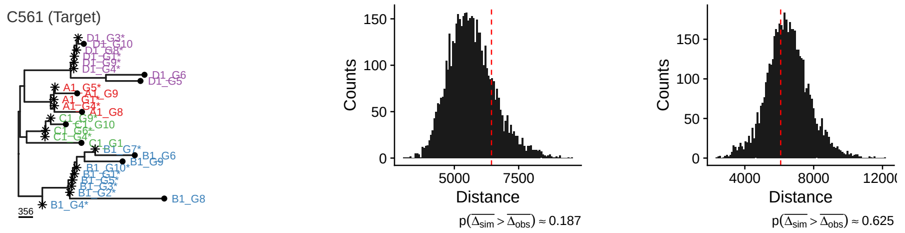

#### Neutral

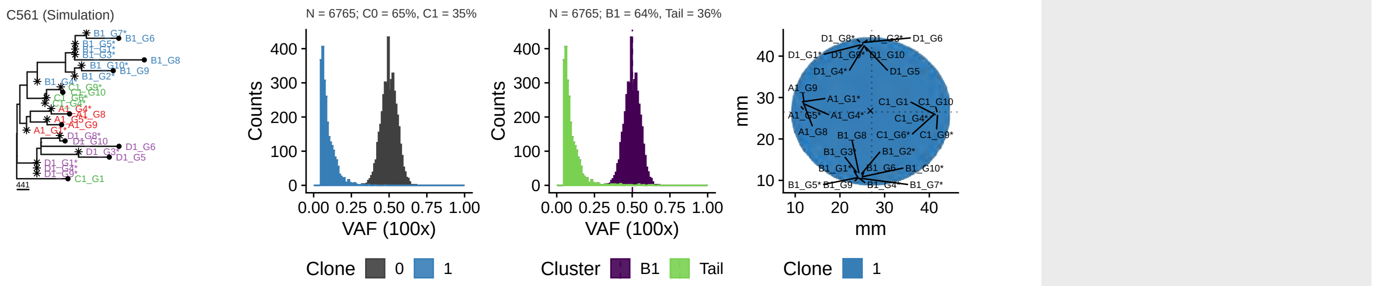

Parameters: $\lambda = 1$, $\mu = 0$, $d_{push} = 47.7$, $m = 0$, $t_s = 0$, $i_a = -1$

#### Selection

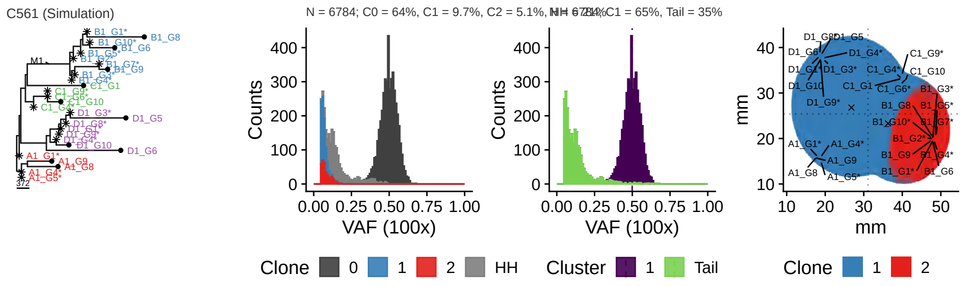

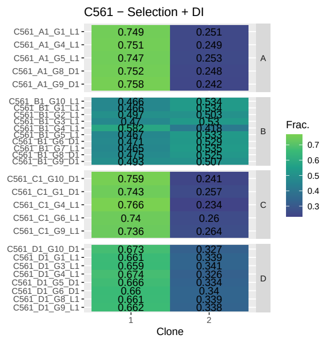

Parameters: $\lambda = (1\ \ 20.6)$, $\mu = (0\ \ 0)$, $d_{push} = (206\ \ 206)$, $m = (0\ \ 0)$, $t_s = (0\ \ 41300)$, $i_a = (-1\ \ 0)$

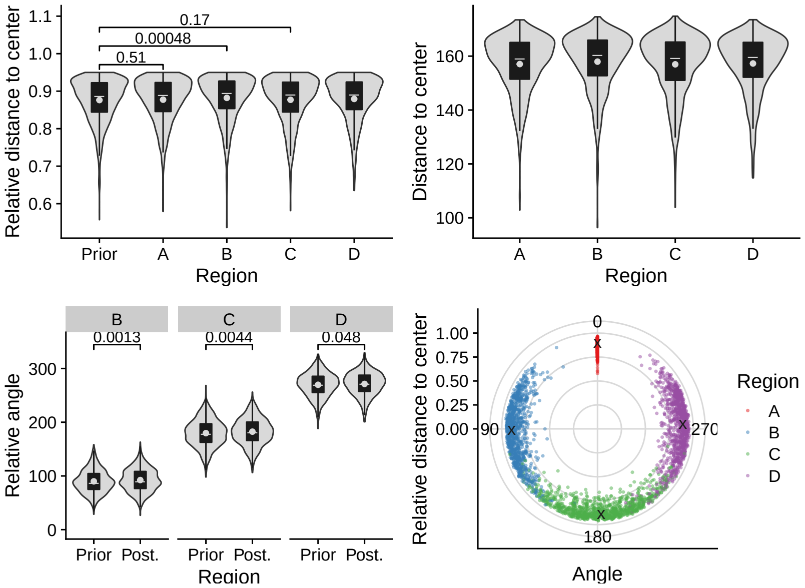
0.17
0.00048
0.51
Relative distance to center
Prior
A
B
C
D
Region
Distance to center
B
0.0013
C
0.0044
D
0.048
Relative angle
Prior
Post.
0
90
180
270
Angle
Region

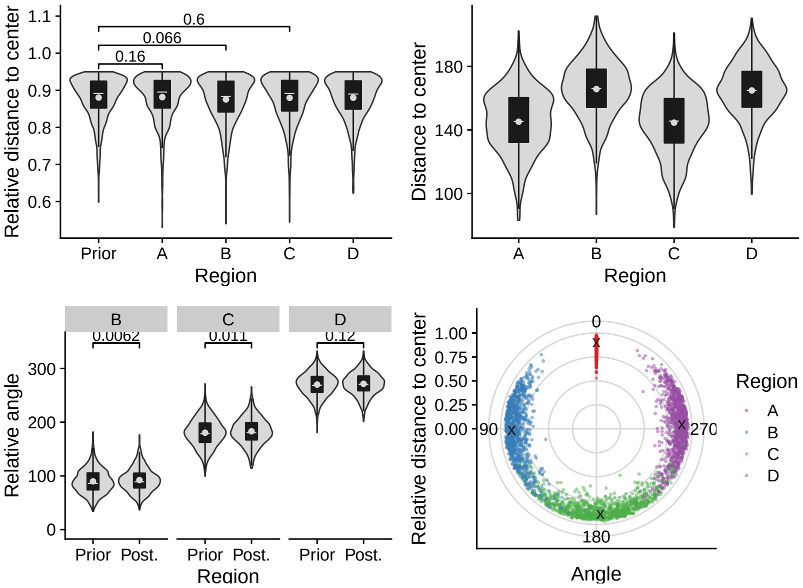
0.6
0.066
0.16
Relative distance to center
Prior
A
B
C
D
Region
Distance to center
B
0.0062
C
0.011
D
0.12
Relative angle
Prior
Post.
0
90
180
270
Angle
Region

## 2.28 C562

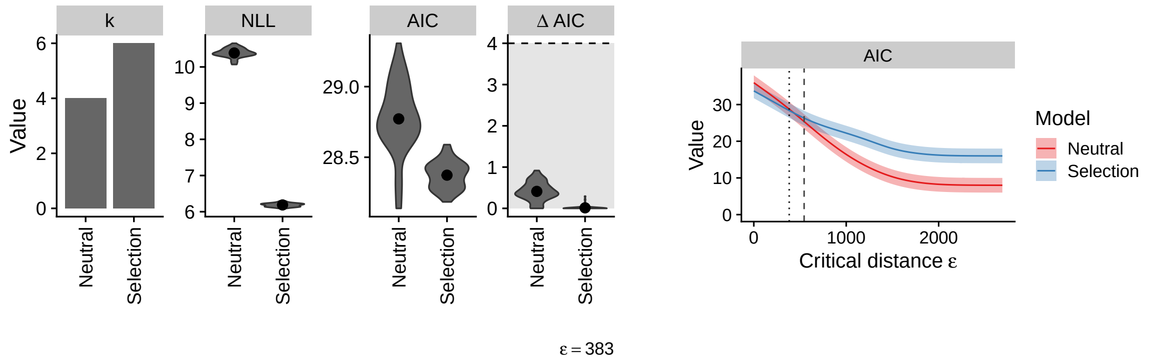

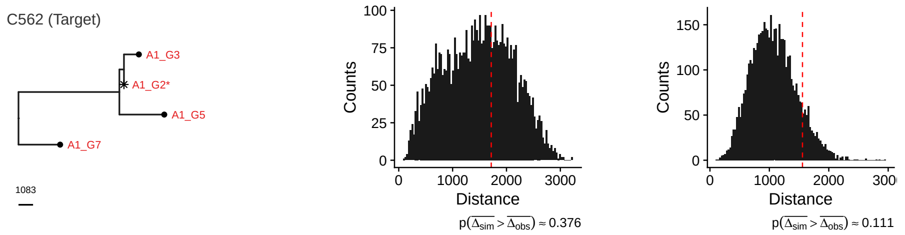

### Neutral

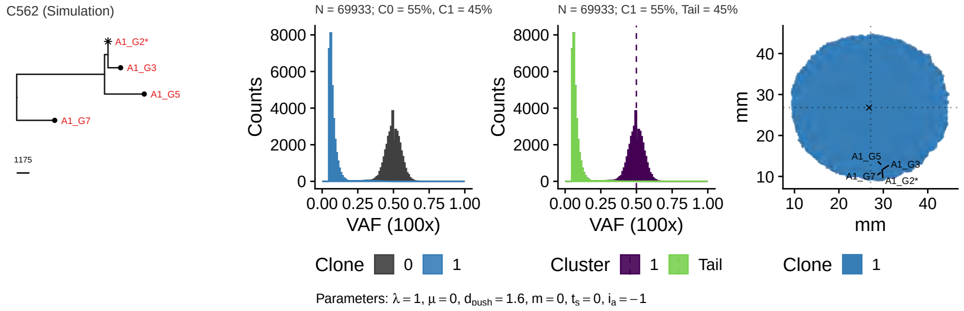

### Selection

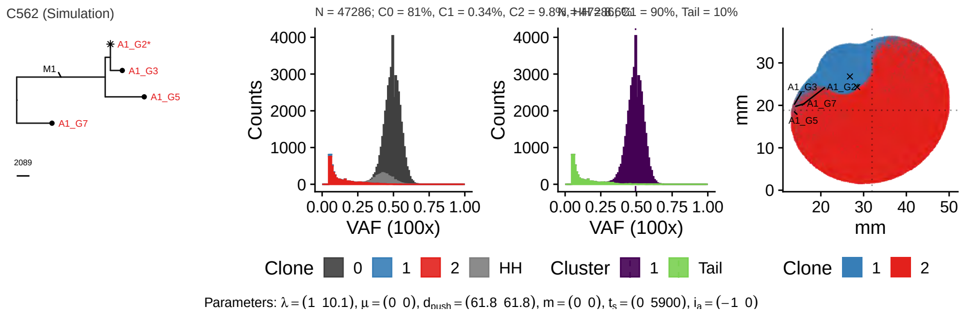

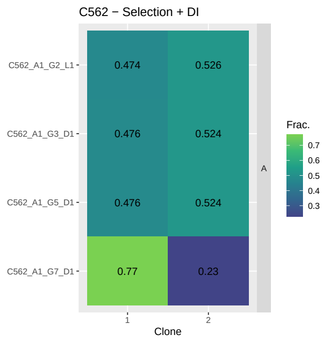
