## Supplementary Figures for "Assessment of the evolutionary consequence of putative driver mutations in colorectal cancer with spatial multiomic data": figS11.pdf

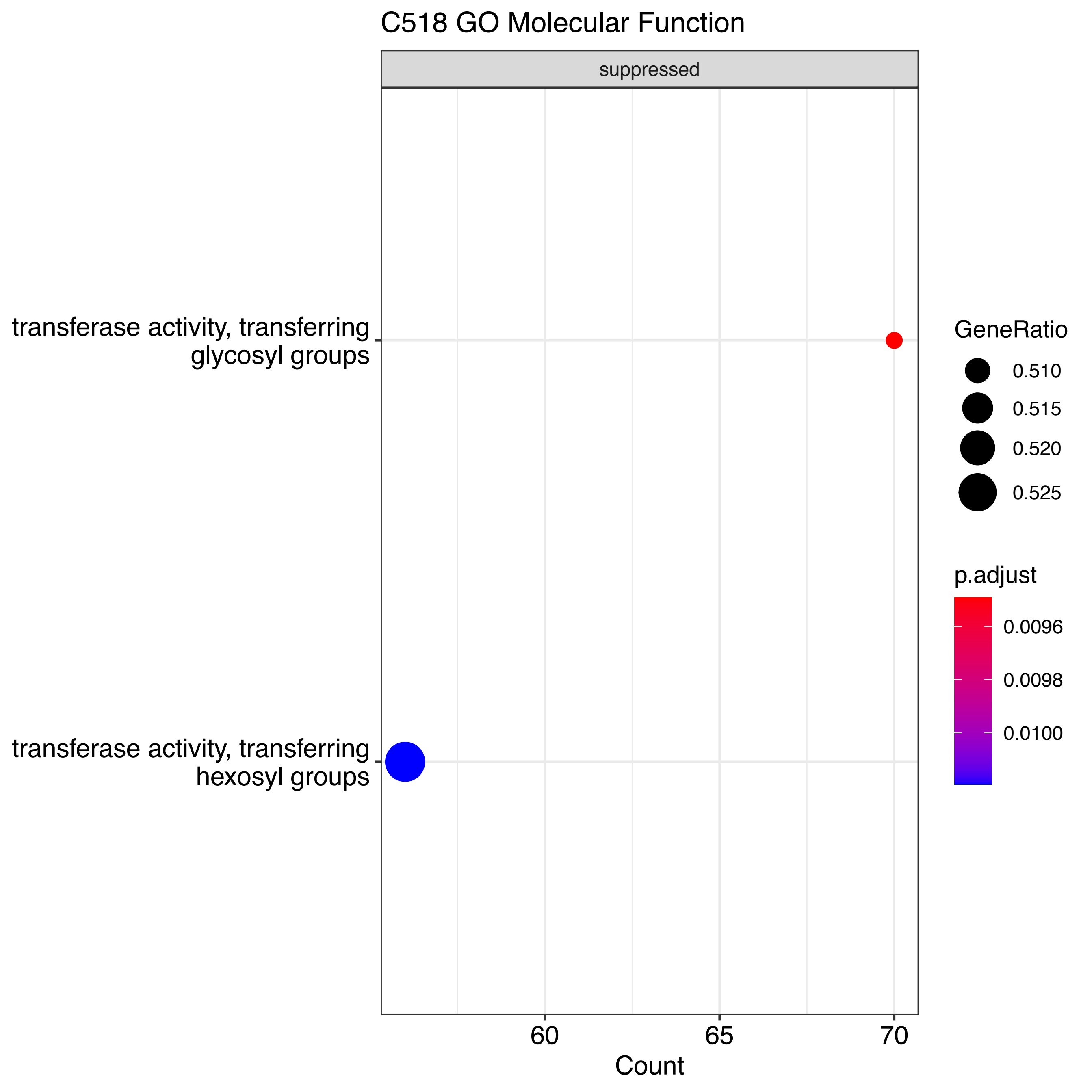
C518 GO Molecular Function
suppressed
transferase activity, transferring glycosyl groups
transferase activity, transferring hexosyl groups
60
65
70
Count
GeneRatio
0.510
0.515
0.520
0.525
p.adjust
0.0096
0.0098
0.0100

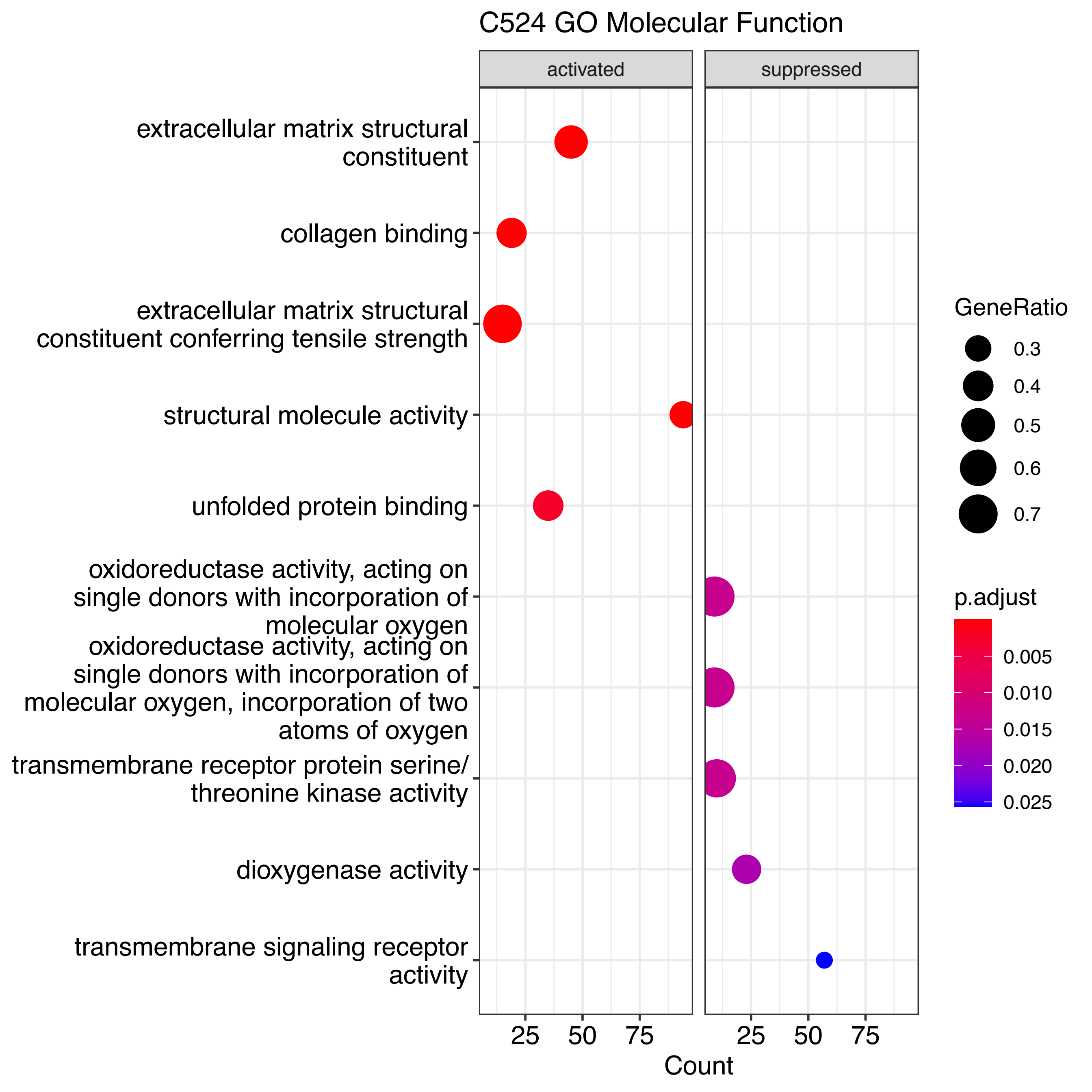

C524 GO Molecular Function
activated
suppressed
extracellular matrix structural constituent
collagen binding
extracellular matrix structural constituent conferring tensile strength
structural molecule activity
unfolded protein binding
oxidoreductase activity, acting on single donors with incorporation of molecular oxygen
oxidoreductase activity, acting on single donors with incorporation of molecular oxygen, incorporation of two atoms of oxygen
transmembrane receptor protein serine/threonine kinase activity
dioxygenase activity
transmembrane signaling receptor activity
25 50 75
25 50 75
Count
GeneRatio
0.3
0.4
0.5
0.6
0.7
p.adjust
0.005
0.010
0.015
0.020
0.025

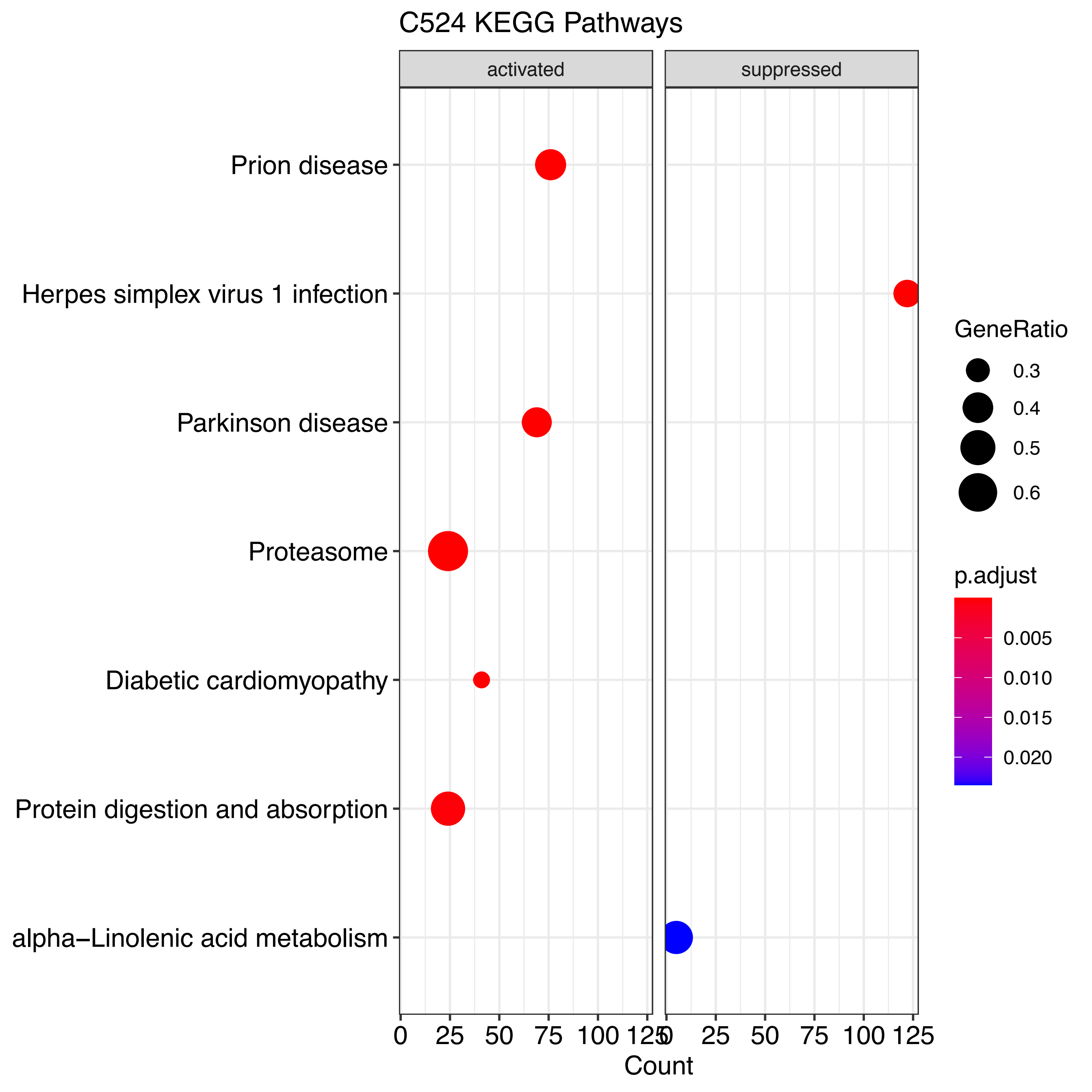
C524 KEGG Pathways
activated
suppressed
Prion disease
Herpes simplex virus 1 infection
Parkinson disease
Proteasome
Diabetic cardiomyopathy
Protein digestion and absorption
alpha−Linolenic acid metabolism
0
25
50
75
100
125
Count
GeneRatio
0.3
0.4
0.5
0.6
p.adjust
0.005
0.010
0.015
0.020

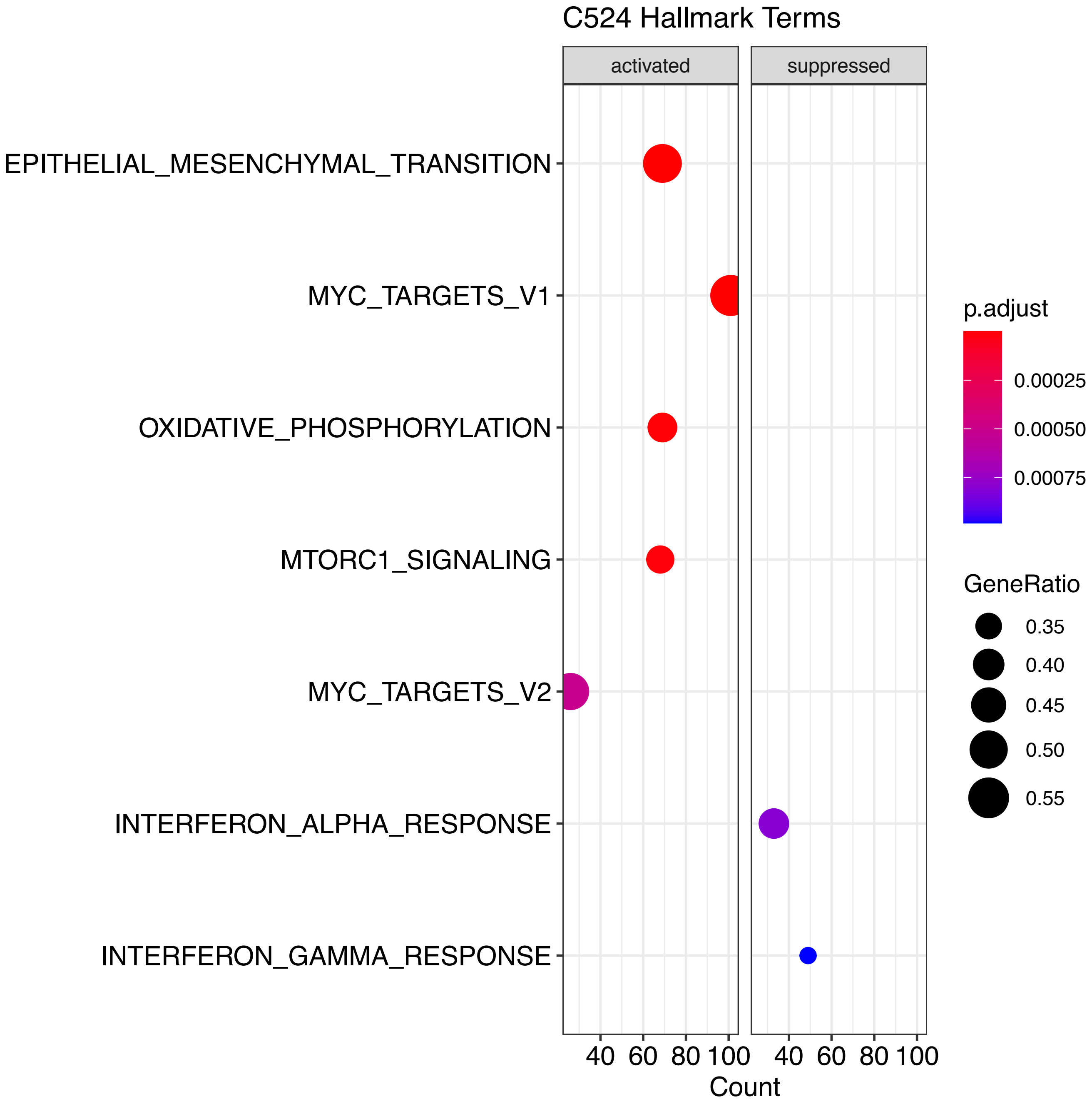

C524 Hallmark Terms
activated
suppressed
EPITHELIAL_MESENCHYMAL_TRANSITION
MYC_TARGETS_V1
OXIDATIVE_PHOSPHORYLATION
MTORC1_SIGNALING
MYC_TARGETS_V2
INTERFERON_ALPHA_RESPONSE
INTERFERON_GAMMA_RESPONSE
40 60 80 100
40 60 80 100
Count
p.adjust
0.00025
0.00050
0.00075
GeneRatio
0.35
0.40
0.45
0.50
0.55

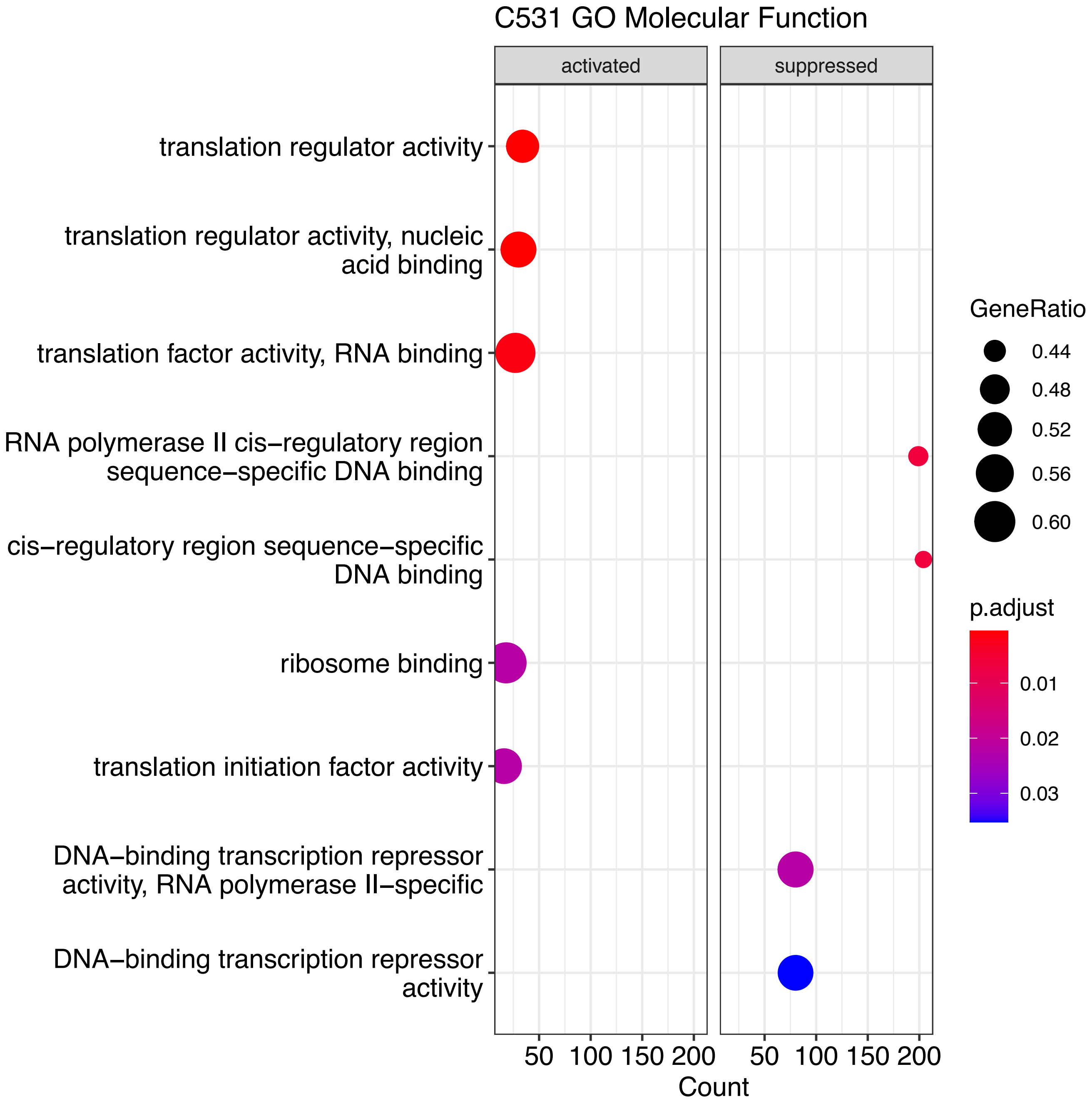
C531 GO Molecular Function
activated
suppressed
translation regulator activity
translation regulator activity, nucleic acid binding
translation factor activity, RNA binding
RNA polymerase II cis−regulatory region sequence−specific DNA binding
cis−regulatory region sequence−specific DNA binding
ribosome binding
translation initiation factor activity
DNA−binding transcription repressor activity, RNA polymerase II−specific
DNA−binding transcription repressor activity
50 100 150 200
50 100 150 200
Count
GeneRatio
0.44
0.48
0.52
0.56
0.60
p.adjust
0.01
0.02
0.03

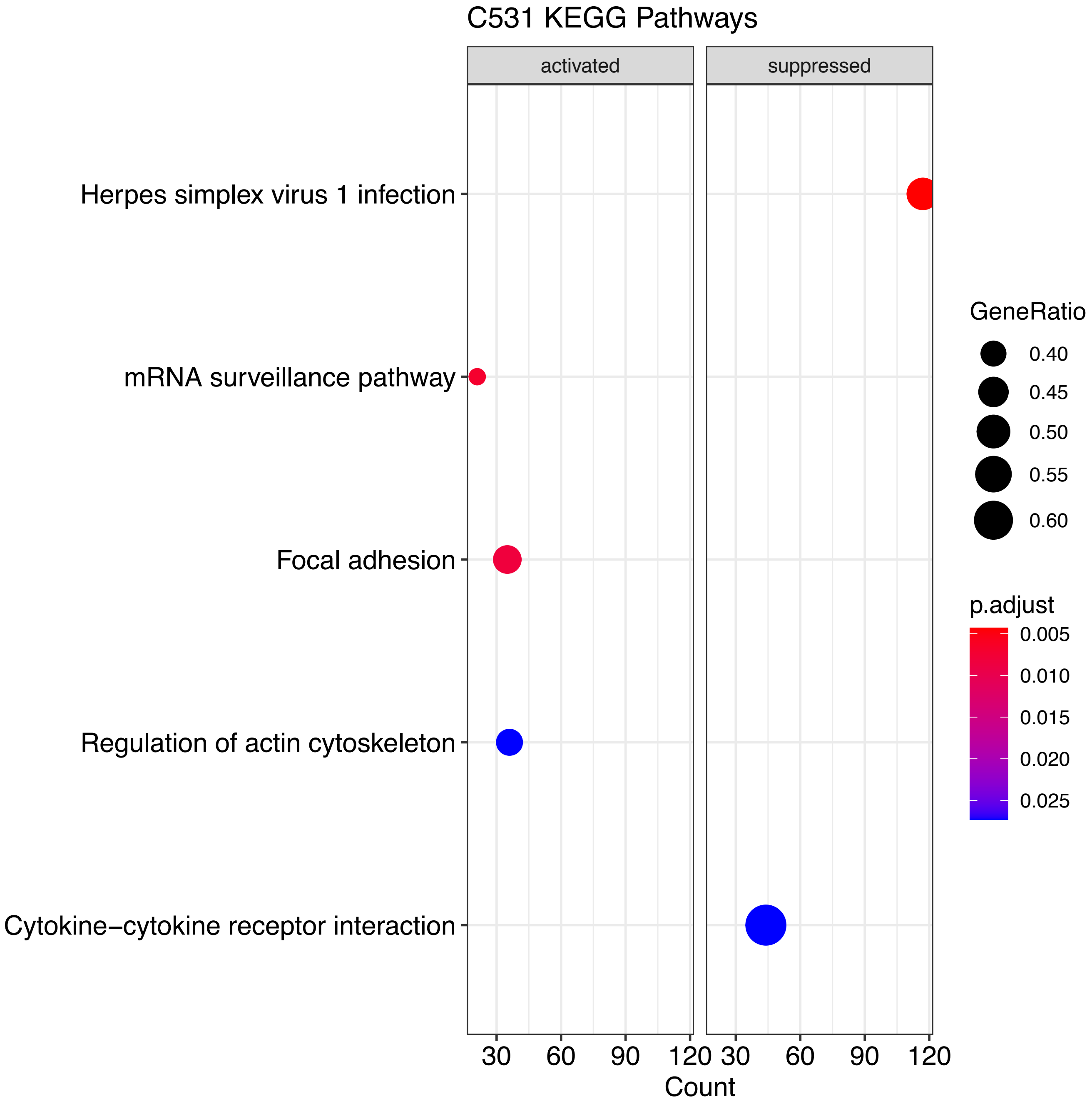

C531 KEGG Pathways
activated
suppressed
Herpes simplex virus 1 infection
mRNA surveillance pathway
Focal adhesion
Regulation of actin cytoskeleton
Cytokine−cytokine receptor interaction
30
60
90
120
30
60
90
120
Count
GeneRatio
0.40
0.45
0.50
0.55
0.60
p.adjust
0.005
0.010
0.015
0.020
0.025

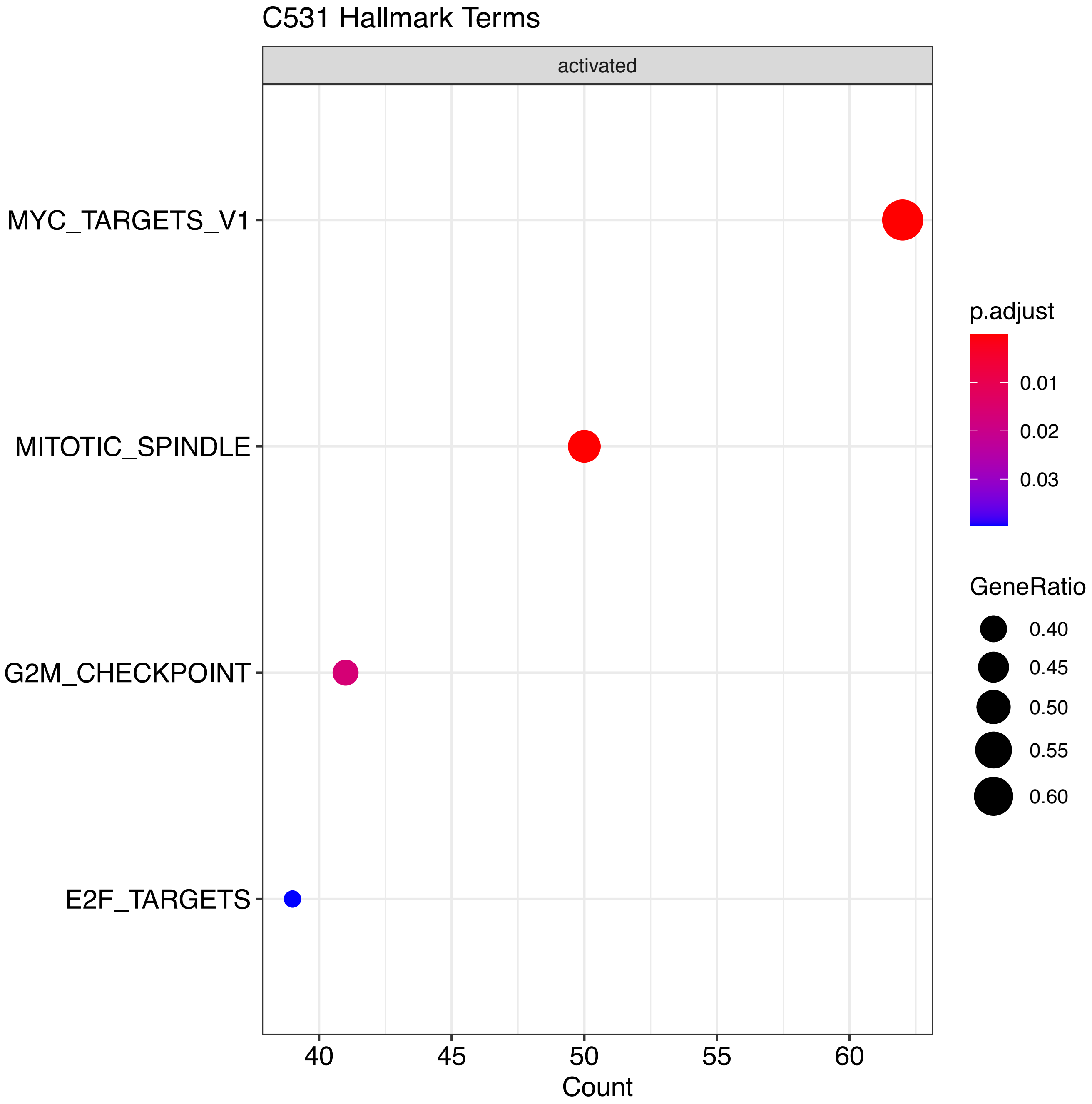
C531 Hallmark Terms
activated
MYC_TARGETS_V1
MITOTIC_SPINDLE
G2M_CHECKPOINT
E2F_TARGETS
40
45
50
55
60
Count
p.adjust
0.01
0.02
0.03
GeneRatio
0.40
0.45
0.50
0.55
0.60

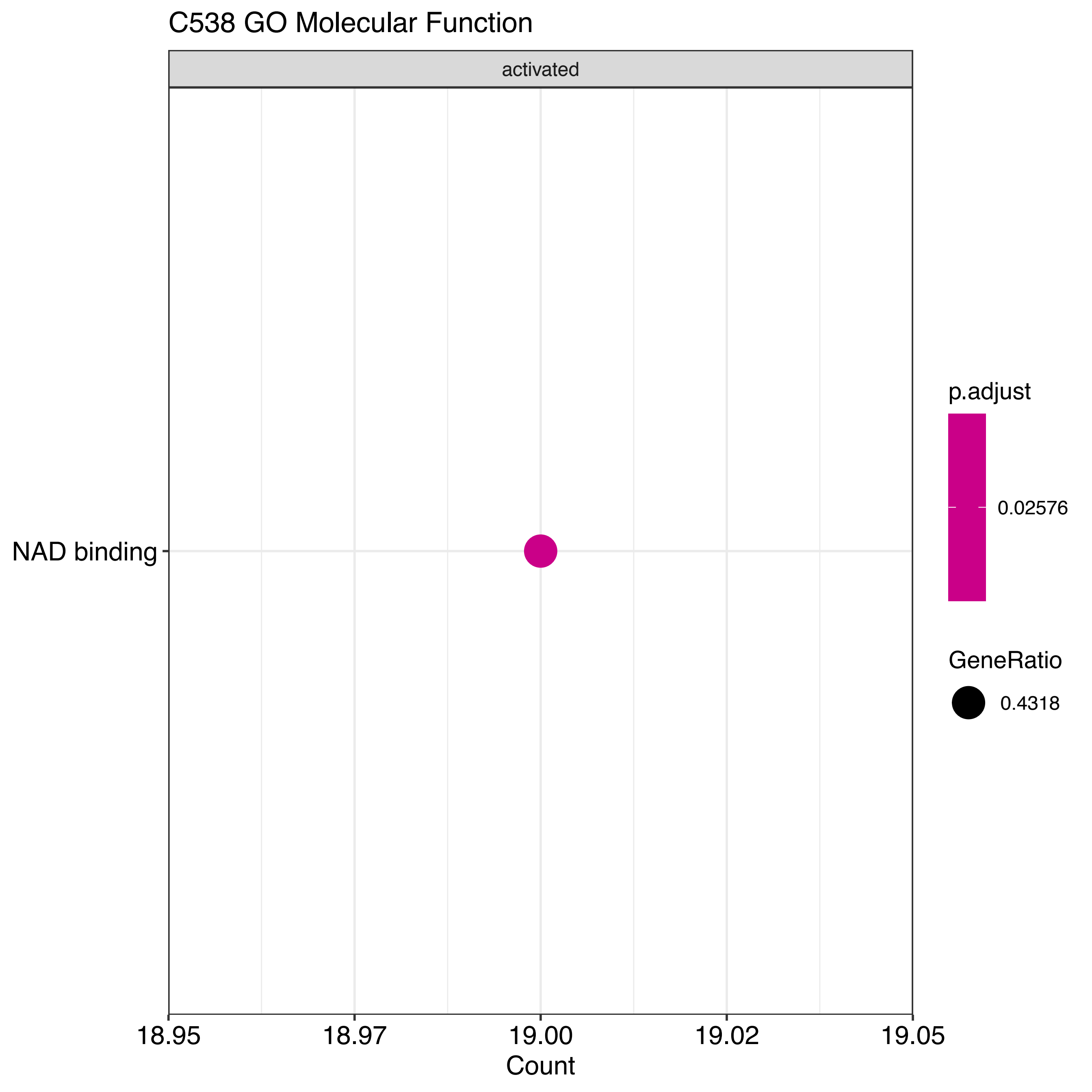

C538 GO Molecular Function
activated
NAD binding
18.95
18.97
19.00
19.02
19.05
Count
p.adjust
0.02576
GeneRatio
0.4318

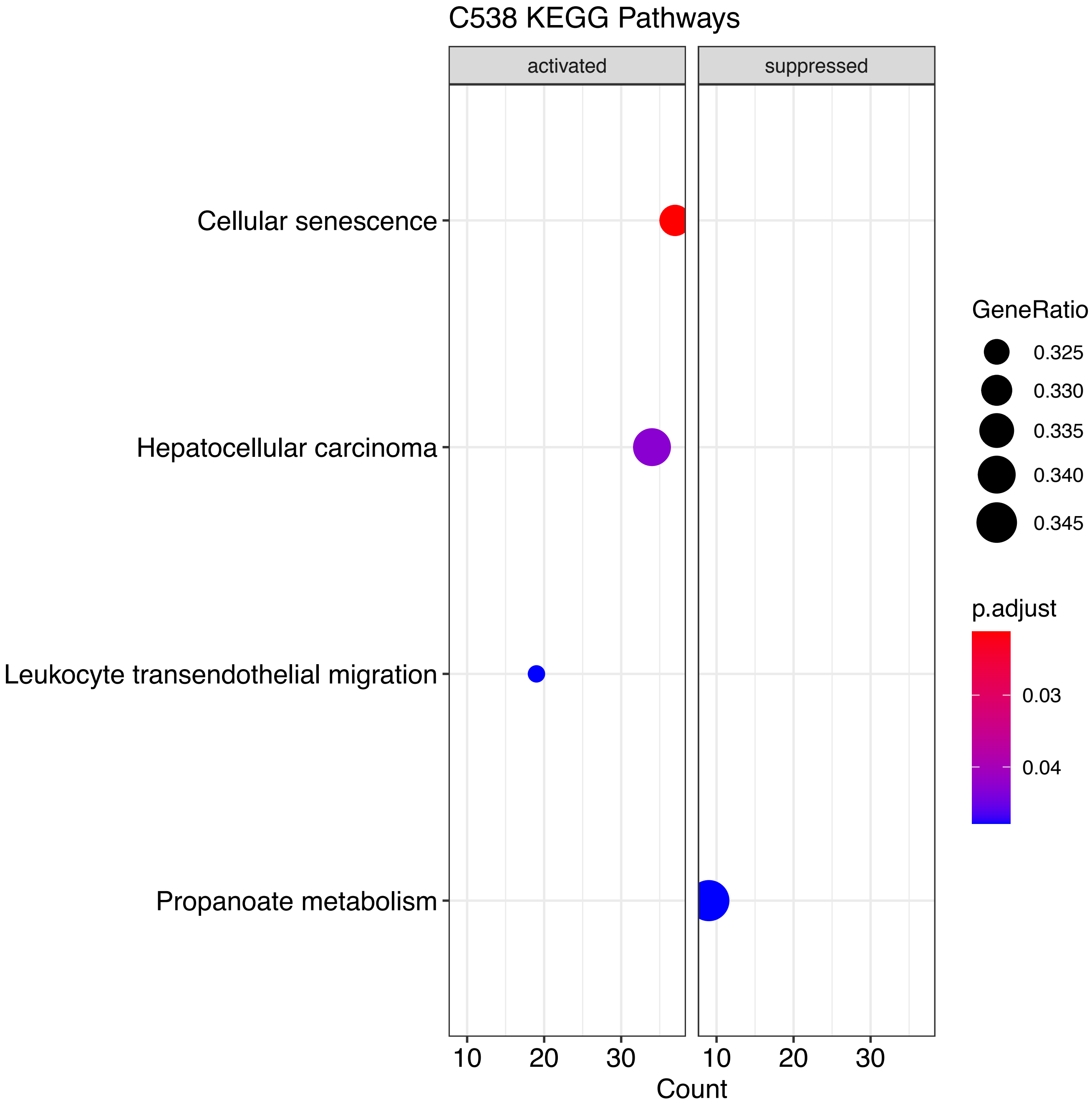
C538 KEGG Pathways
activated
suppressed
Cellular senescence
Hepatocellular carcinoma
Leukocyte transendothelial migration
Propanoate metabolism
10
20
30
10
20
30
Count
GeneRatio
0.325
0.330
0.335
0.340
0.345
p.adjust
0.03
0.04

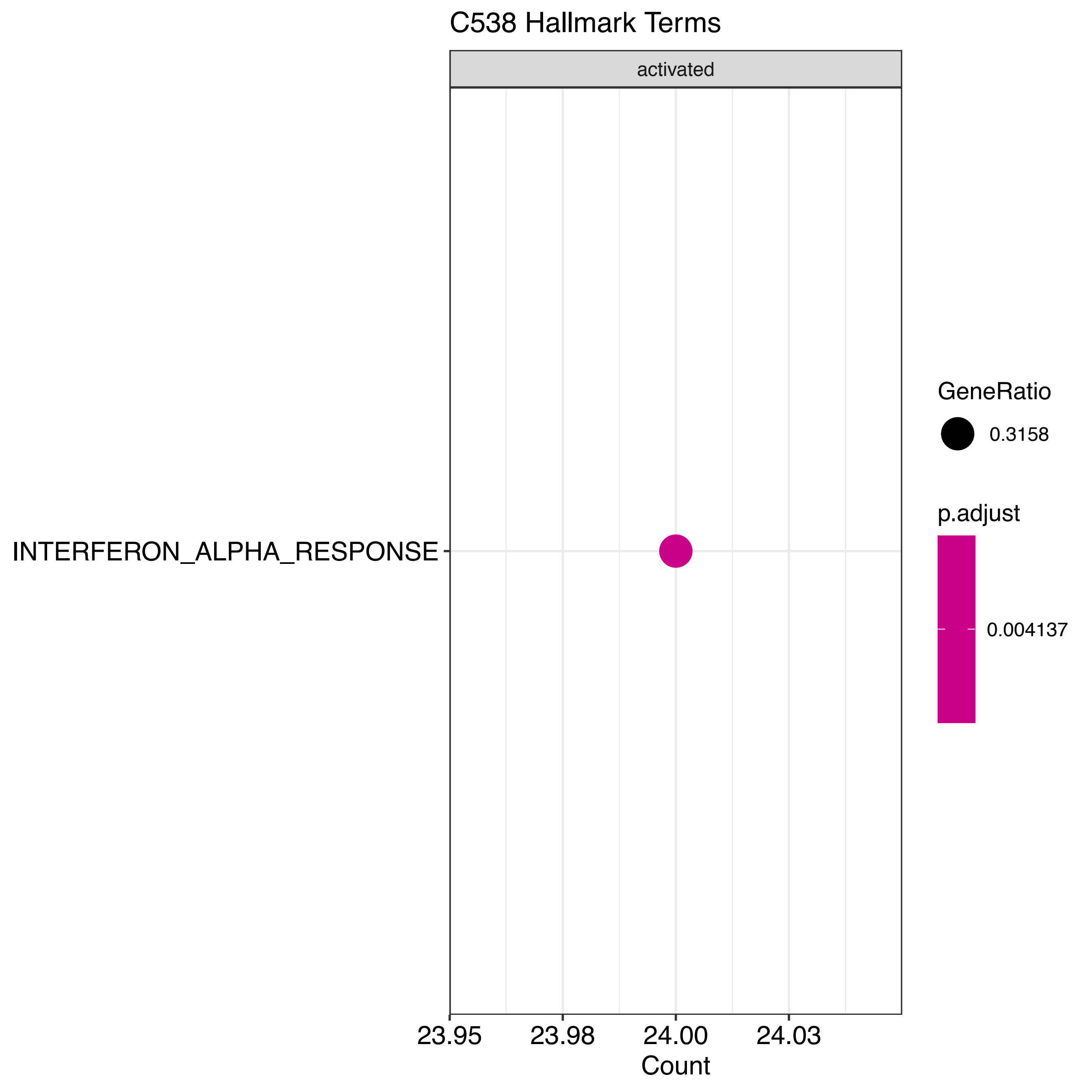

C538 Hallmark Terms
activated
INTERFERON_ALPHA_RESPONSE
23.95
23.98
24.00
24.03
Count
GeneRatio
0.3158
p.adjust
0.004137

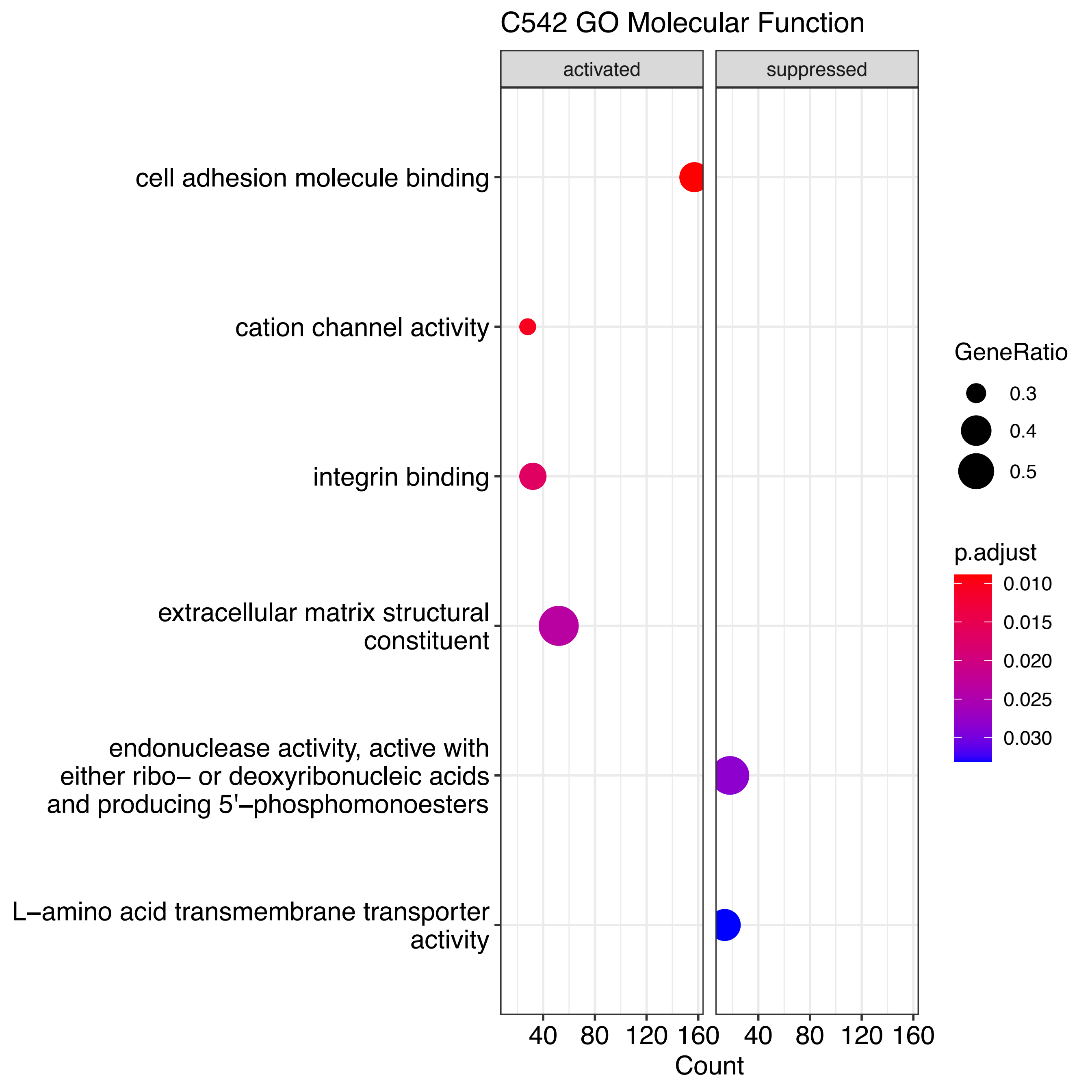

C542 GO Molecular Function
activated
suppressed
cell adhesion molecule binding
cation channel activity
integrin binding
extracellular matrix structural constituent
endonuclease activity, active with either ribo− or deoxyribonucleic acids and producing 5'−phosphomonoesters
L−amino acid transmembrane transporter activity
40
80
120
160
40
80
120
160
Count
GeneRatio
0.3
0.4
0.5
p.adjust
0.010
0.015
0.020
0.025
0.030

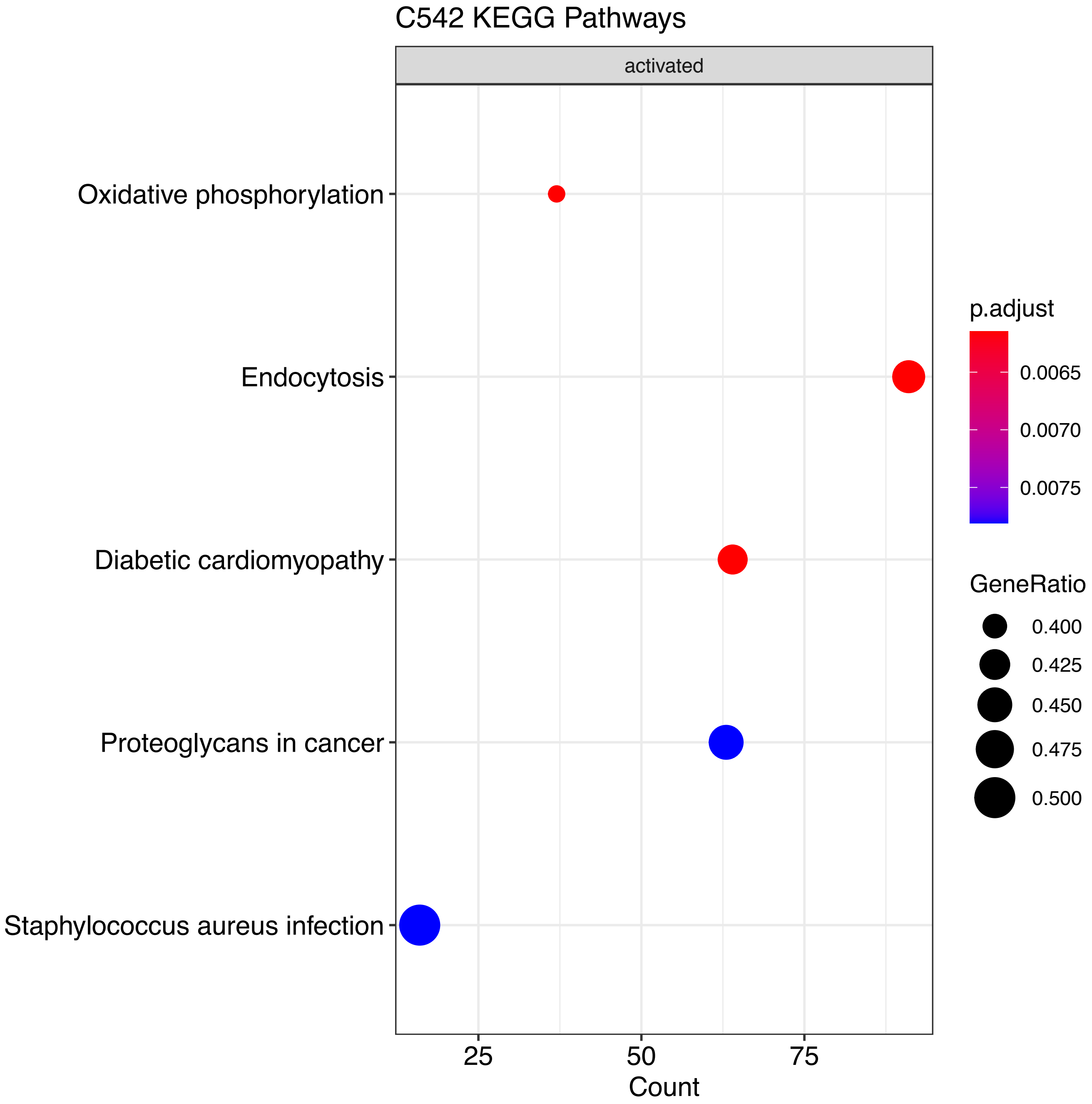

C542 KEGG Pathways
activated
Oxidative phosphorylation
Endocytosis
Diabetic cardiomyopathy
Proteoglycans in cancer
Staphylococcus aureus infection
25
50
75
Count
p.adjust
0.0065
0.0070
0.0075
GeneRatio
0.400
0.425
0.450
0.475
0.500

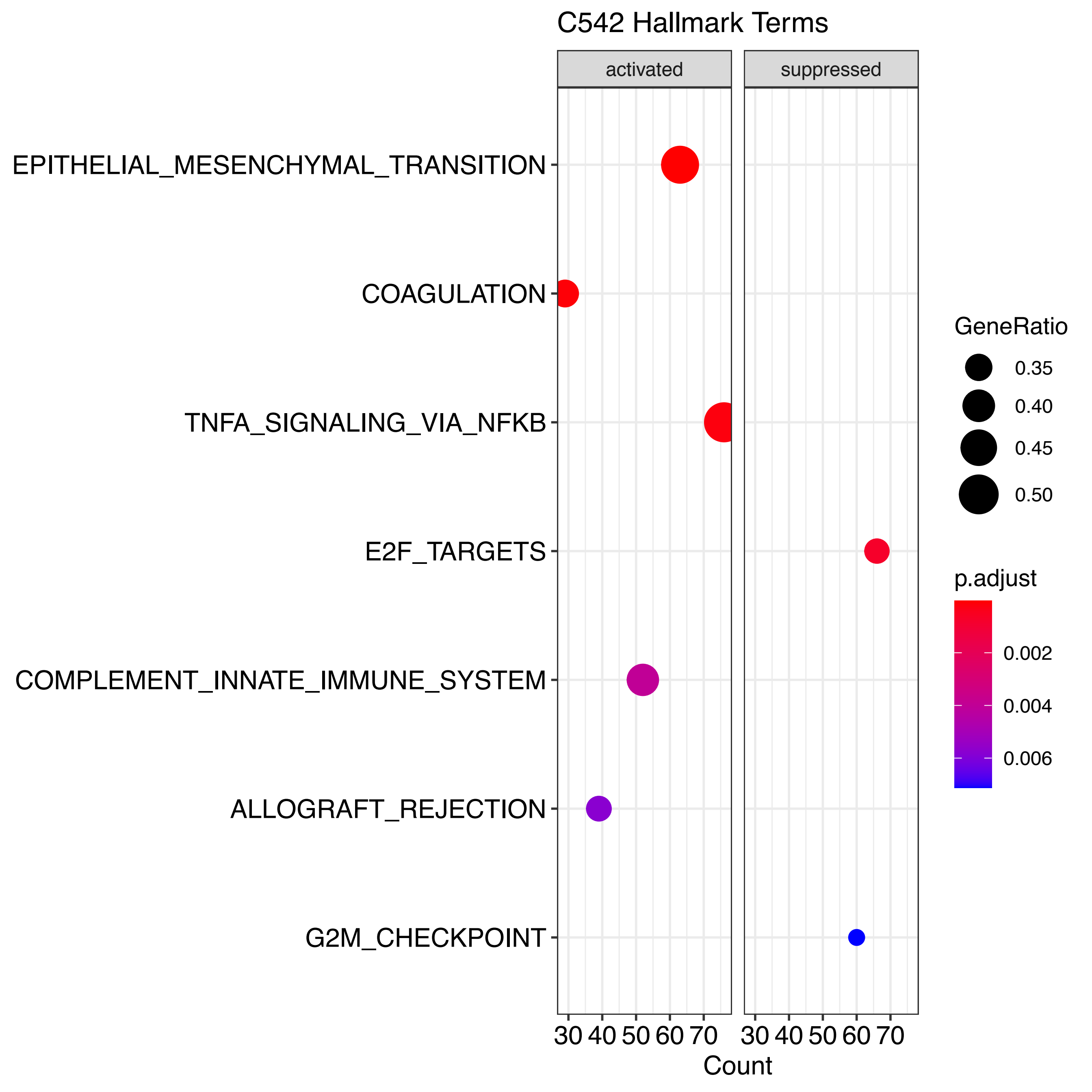
C542 Hallmark Terms
activated
suppressed
EPITHELIAL_MESENCHYMAL_TRANSITION
COAGULATION
TNFA_SIGNALING_VIA_NFKB
E2F_TARGETS
COMPLEMENT_INNATE_IMMUNE_SYSTEM
ALLOGRAFT_REJECTION
G2M_CHECKPOINT
30 40 50 60 70
30 40 50 60 70
Count
GeneRatio
0.35
0.40
0.45
0.50
p.adjust
0.002
0.004
0.006

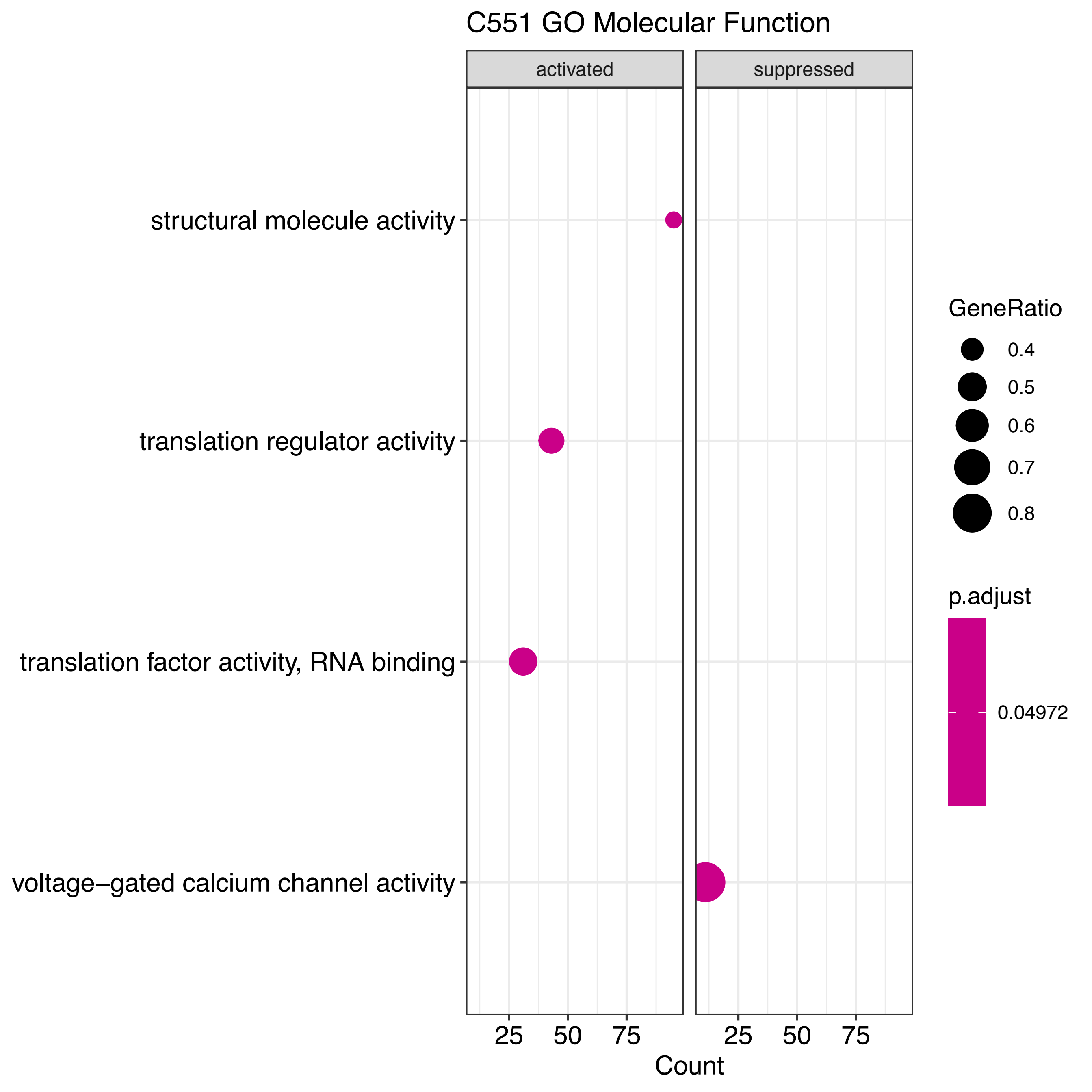
C551 GO Molecular Function
activated
suppressed
structural molecule activity
translation regulator activity
translation factor activity, RNA binding
voltage−gated calcium channel activity
25
50
75
25
50
75
Count
GeneRatio
0.4
0.5
0.6
0.7
0.8
p.adjust
0.04972

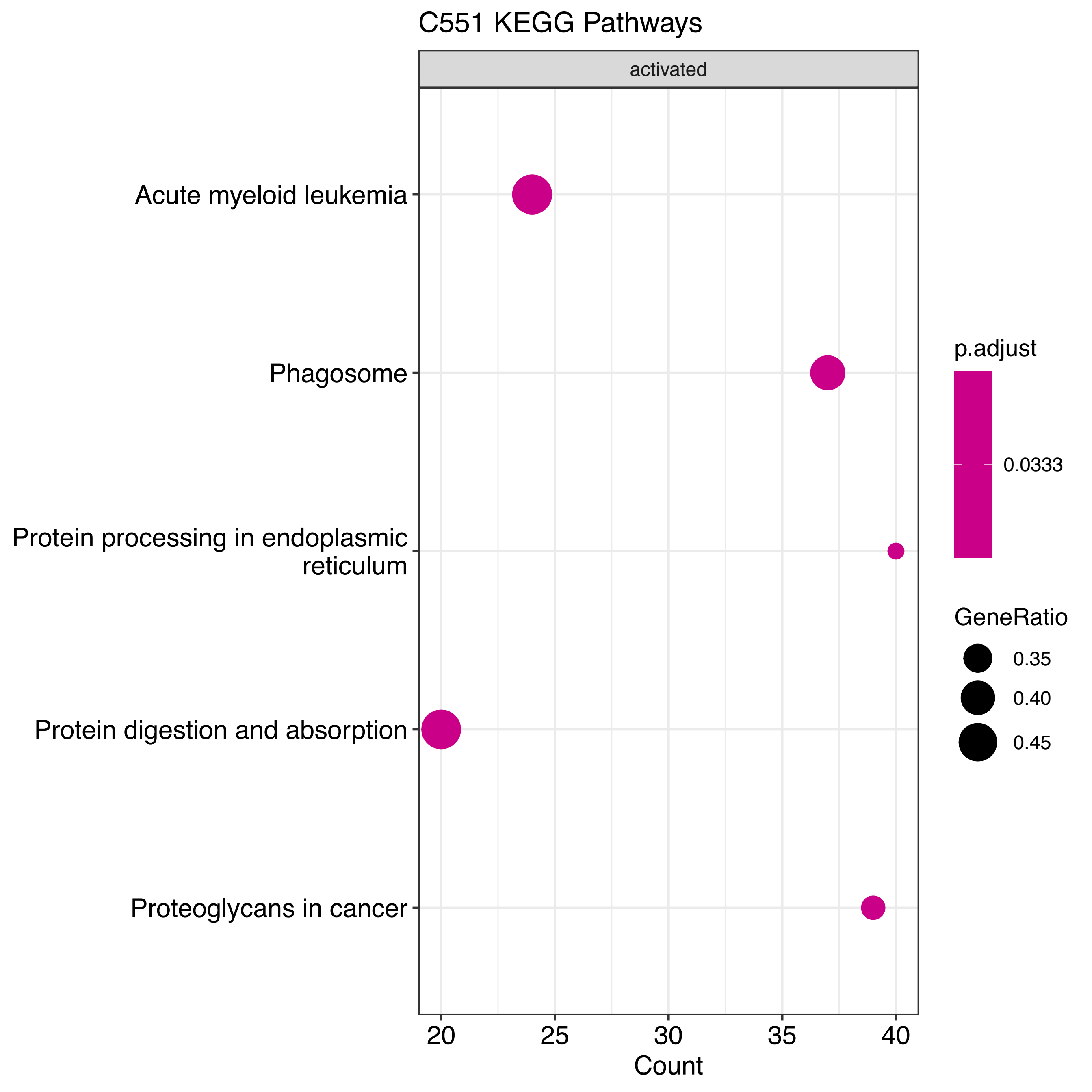

C551 KEGG Pathways
activated
Acute myeloid leukemia
Phagosome
Protein processing in endoplasmic reticulum
Protein digestion and absorption
Proteoglycans in cancer
20
25
30
35
40
Count
p.adjust
0.0333
GeneRatio
0.35
0.40
0.45

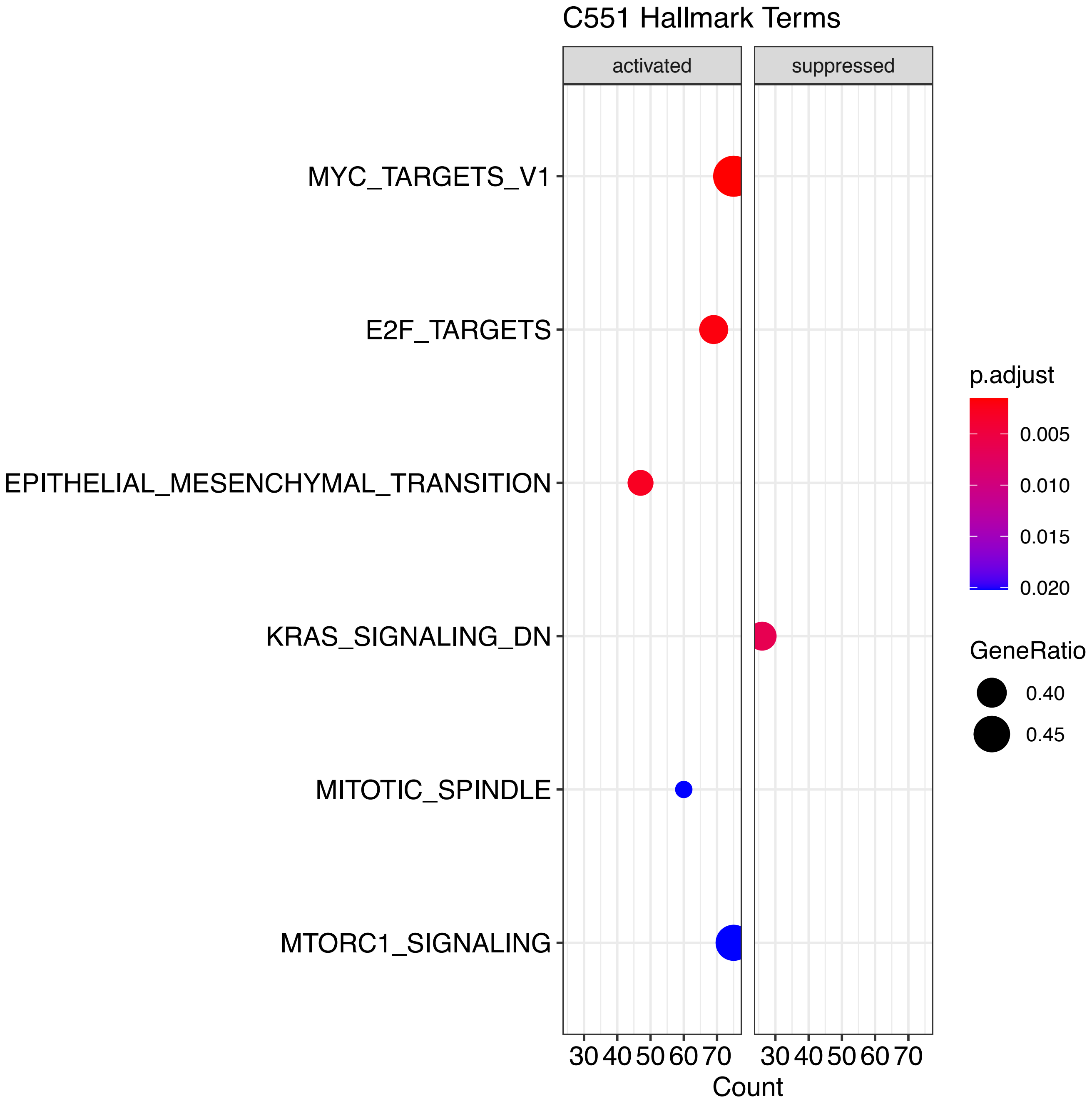
C551 Hallmark Terms
activated
suppressed
MYC_TARGETS_V1
E2F_TARGETS
EPITHELIAL_MESENCHYMAL_TRANSITION
KRAS_SIGNALING_DN
MITOTIC_SPINDLE
MTORC1_SIGNALING
30 40 50 60 70
30 40 50 60 70
Count
p.adjust
0.005
0.010
0.015
0.020
GeneRatio
0.40
0.45

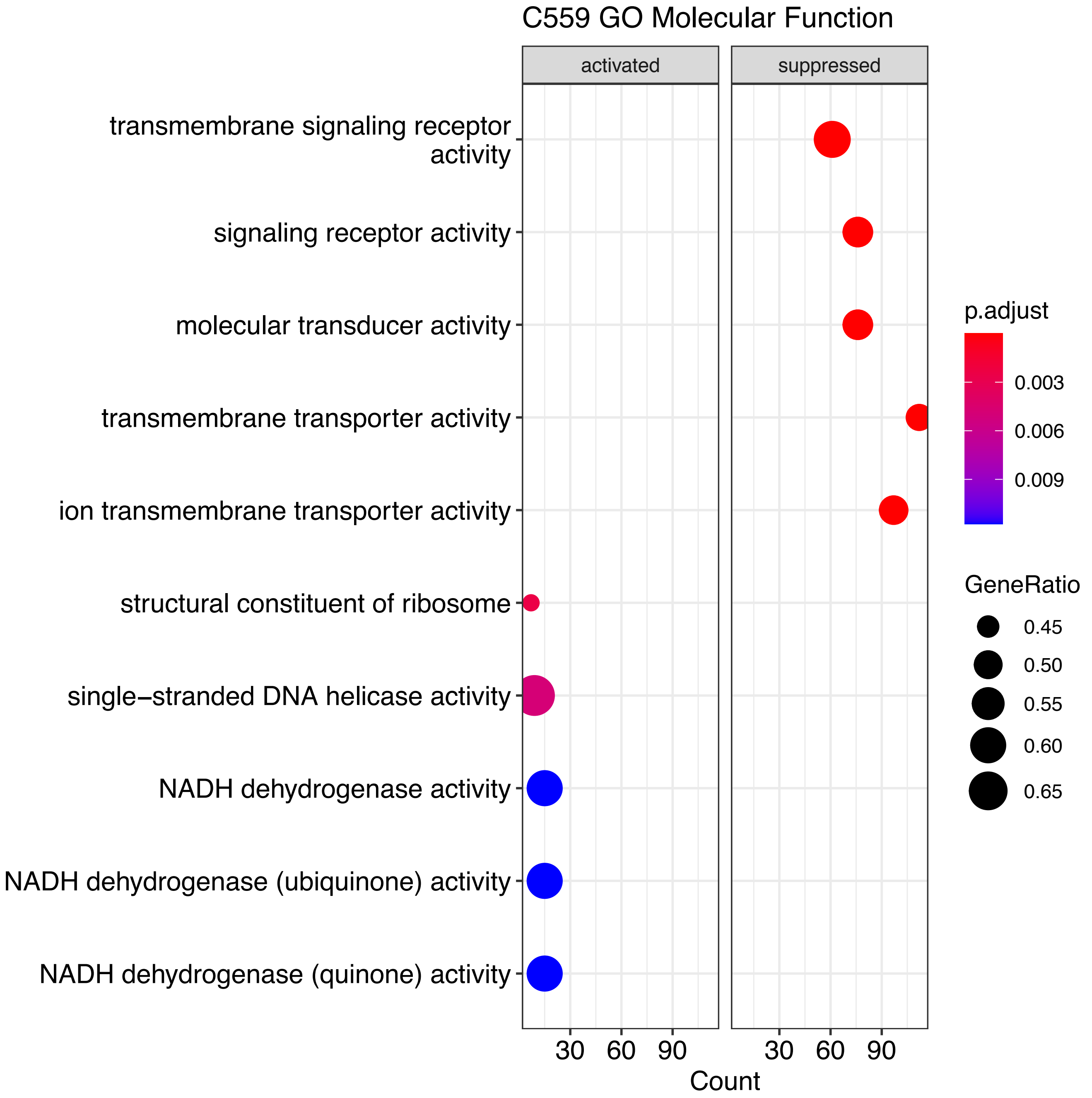

C559 GO Molecular Function
activated
suppressed
transmembrane signaling receptor activity
signaling receptor activity
molecular transducer activity
transmembrane transporter activity
ion transmembrane transporter activity
structural constituent of ribosome
single−stranded DNA helicase activity
NADH dehydrogenase activity
NADH dehydrogenase (ubiquinone) activity
NADH dehydrogenase (quinone) activity
30 60 90
30 60 90
Count
p.adjust
0.003
0.006
0.009
GeneRatio
0.45
0.50
0.55
0.60
0.65

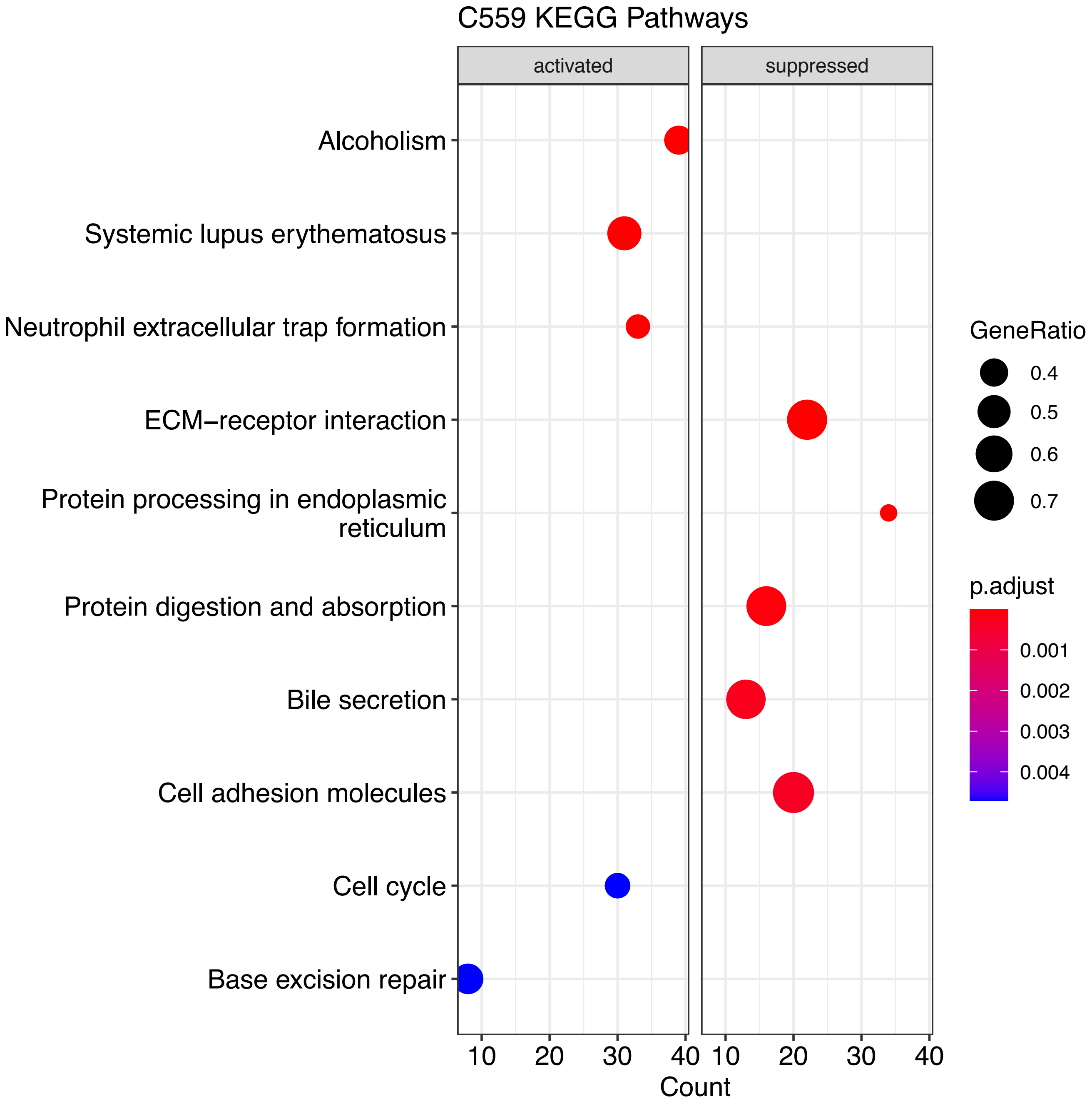
C559 KEGG Pathways
activated
suppressed
Alcoholism
Systemic lupus erythematosus
Neutrophil extracellular trap formation
ECM−receptor interaction
Protein processing in endoplasmic reticulum
Protein digestion and absorption
Bile secretion
Cell adhesion molecules
Cell cycle
Base excision repair
10
20
30
40
Count
GeneRatio
0.4
0.5
0.6
0.7
p.adjust
0.001
0.002
0.003
0.004

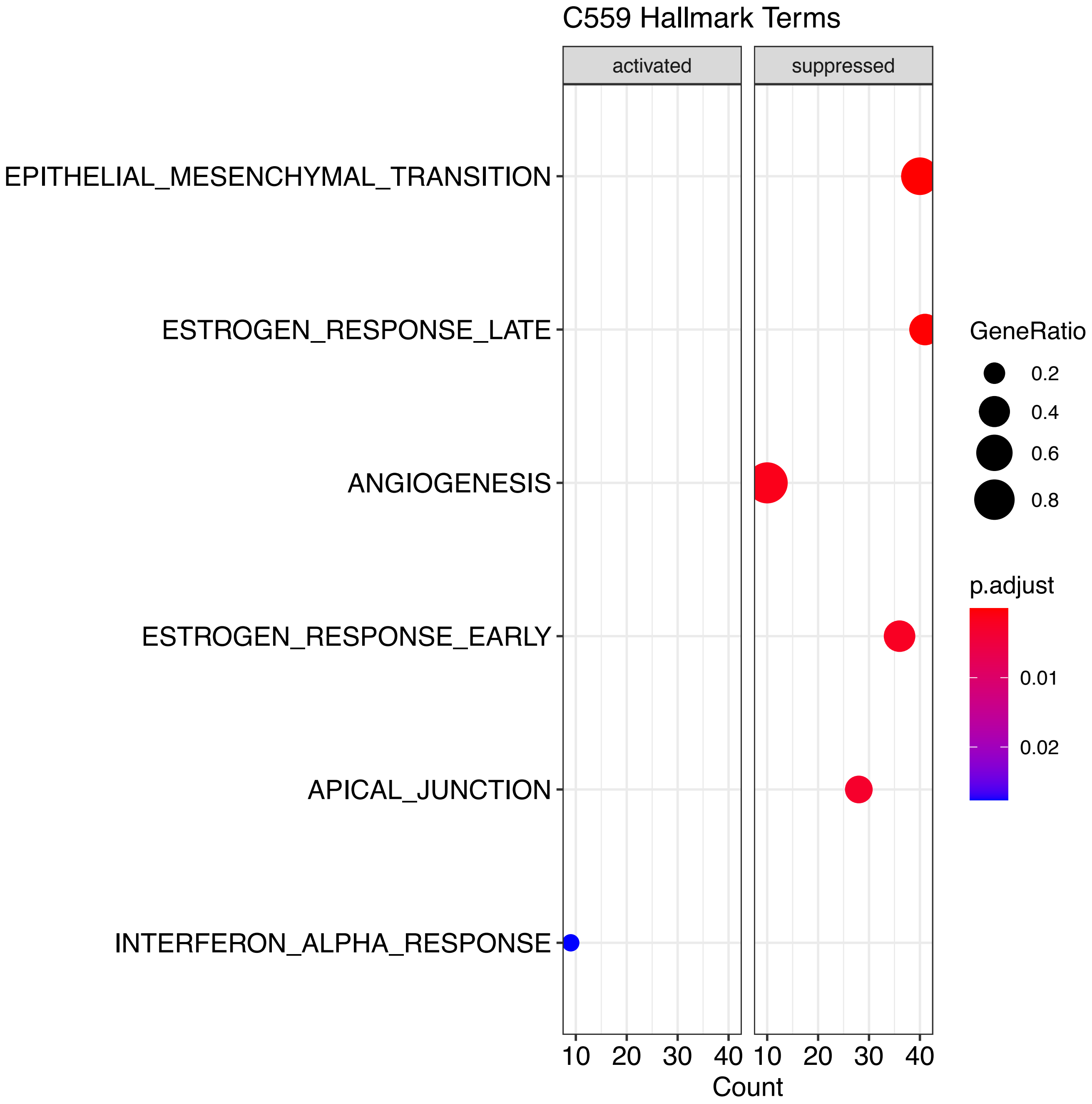
C559 Hallmark Terms
activated
suppressed
EPITHELIAL_MESENCHYMAL_TRANSITION
ESTROGEN_RESPONSE_LATE
ANGIOGENESIS
ESTROGEN_RESPONSE_EARLY
APICAL_JUNCTION
INTERFERON_ALPHA_RESPONSE
10 20 30 40
10 20 30 40
Count
GeneRatio
0.2
0.4
0.6
0.8
p.adjust
0.01
0.02
