## Supplementary figures and images for "Assessment of the evolutionary consequence of putative driver mutations in colorectal cancer with spatial multiomic data"

### figS1.png

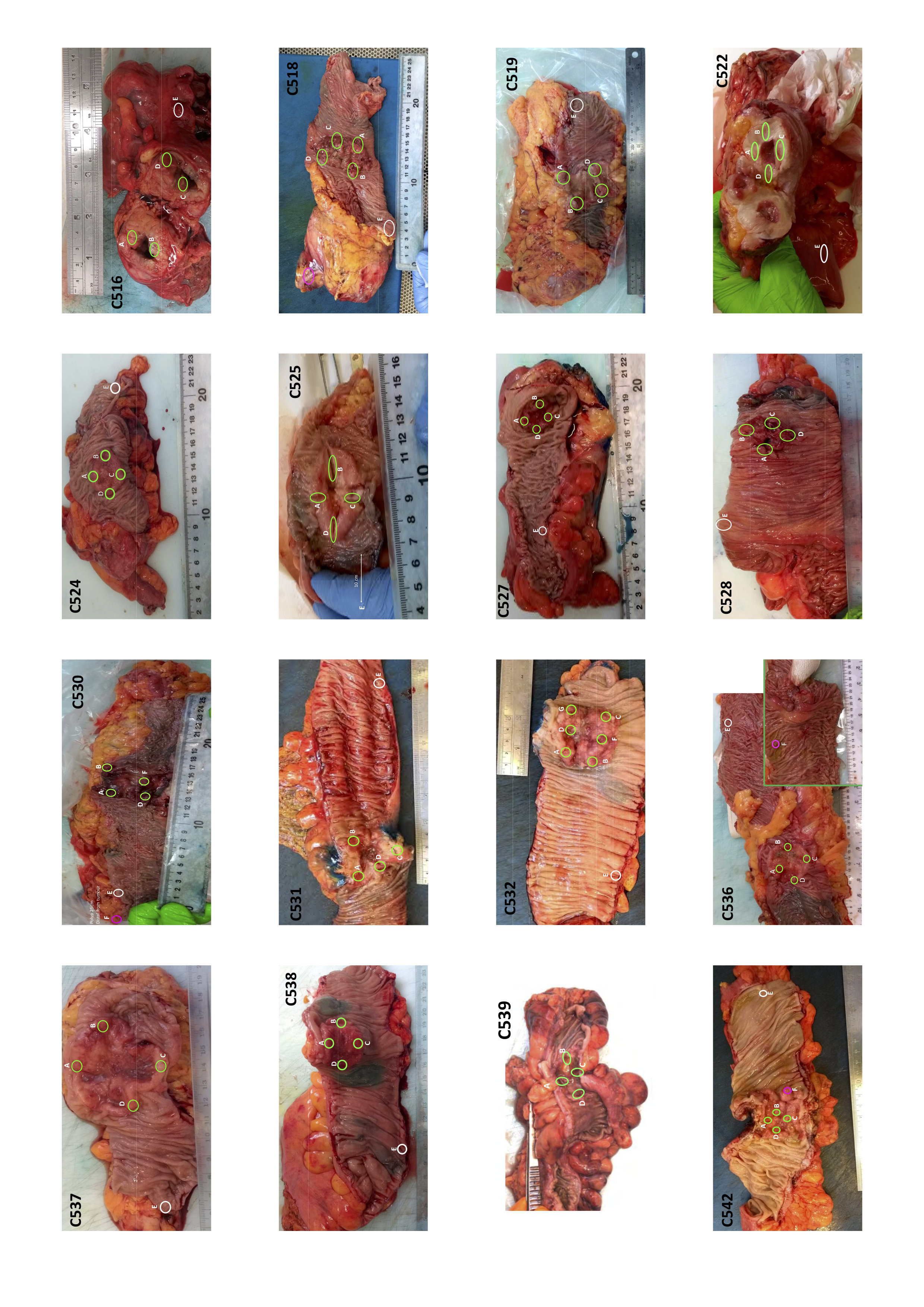

### figS3.pdf

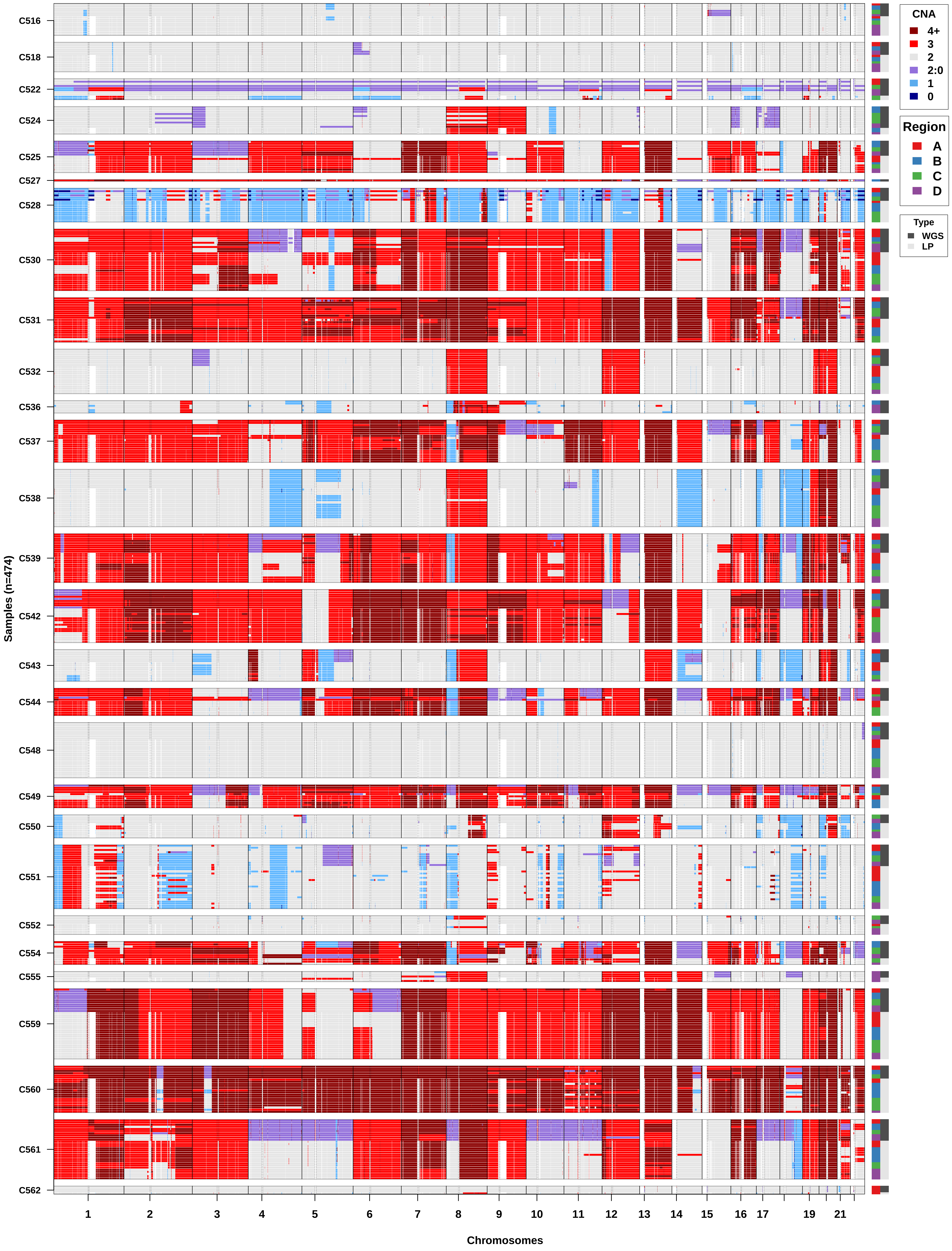

### figS6.pdf

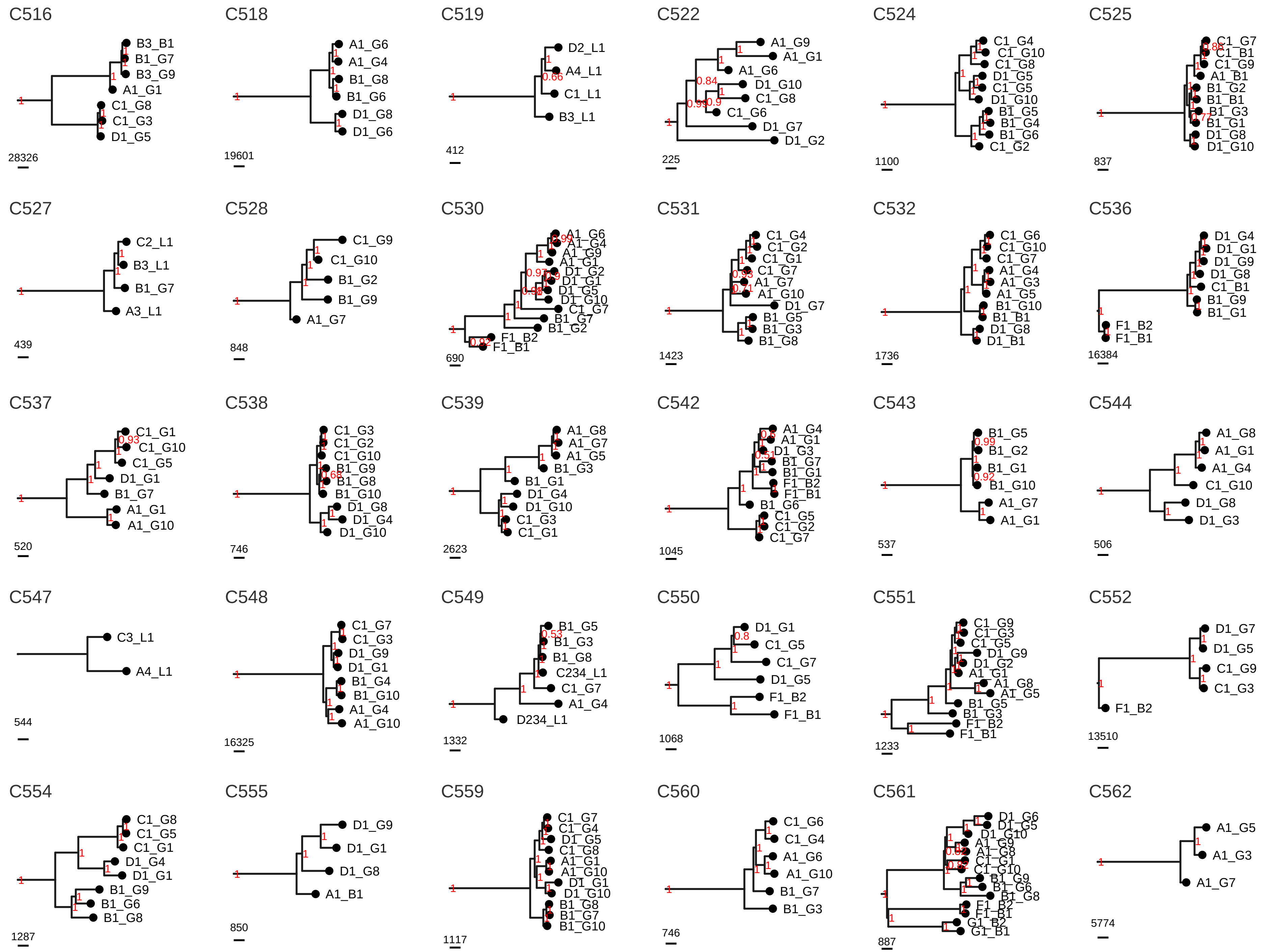

### figS7.pdf

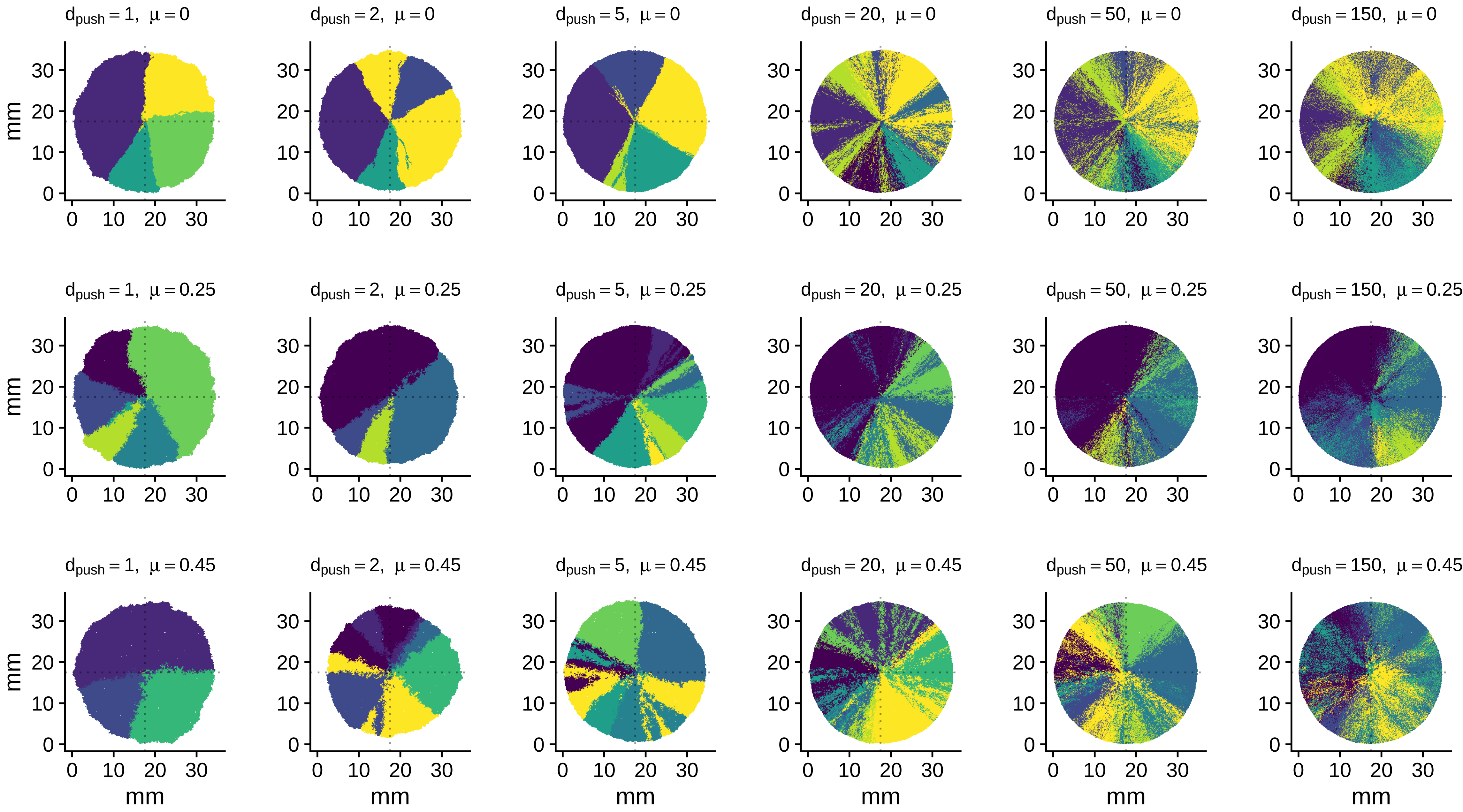

### figS8.pdf

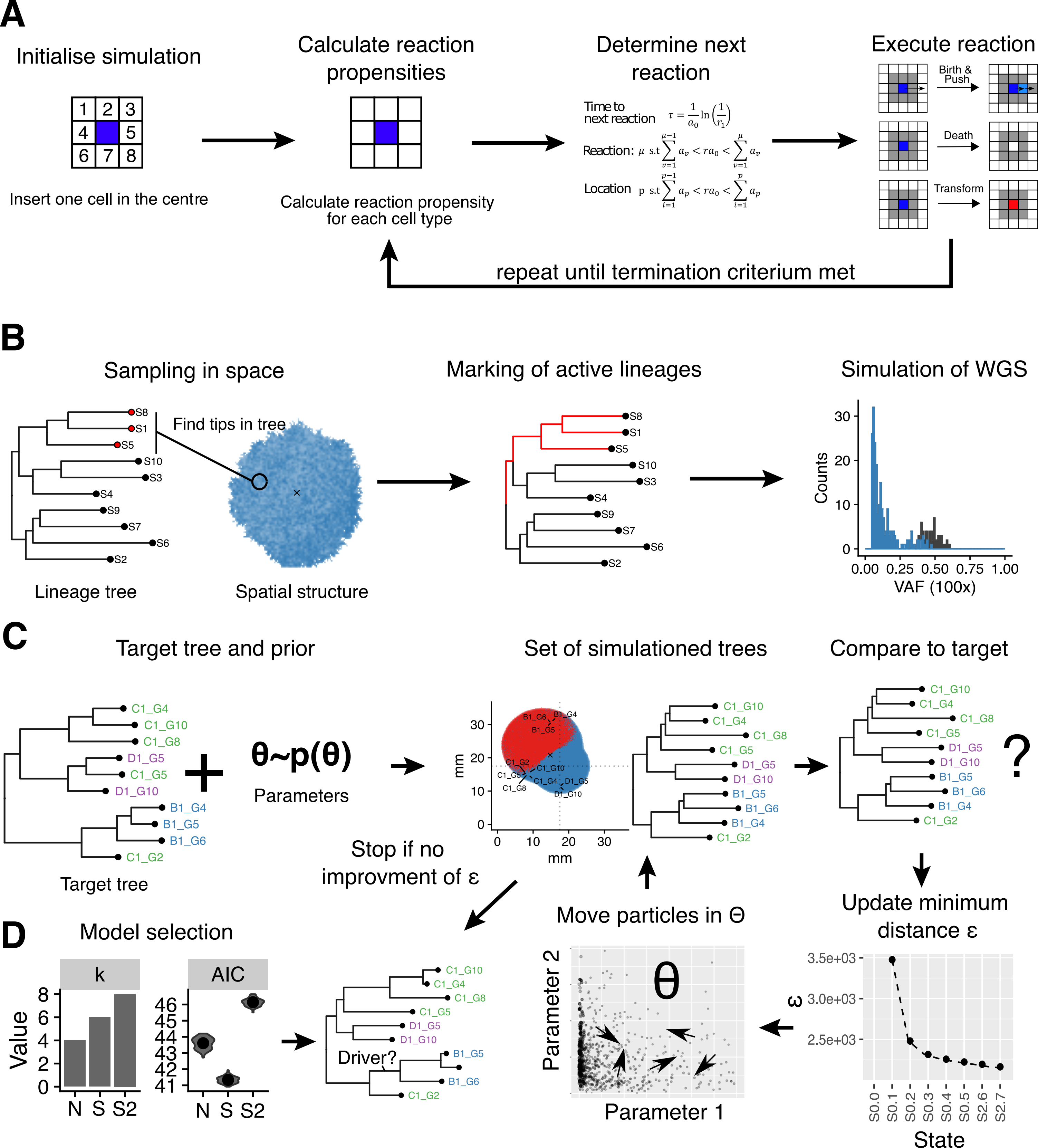

### figS9.png

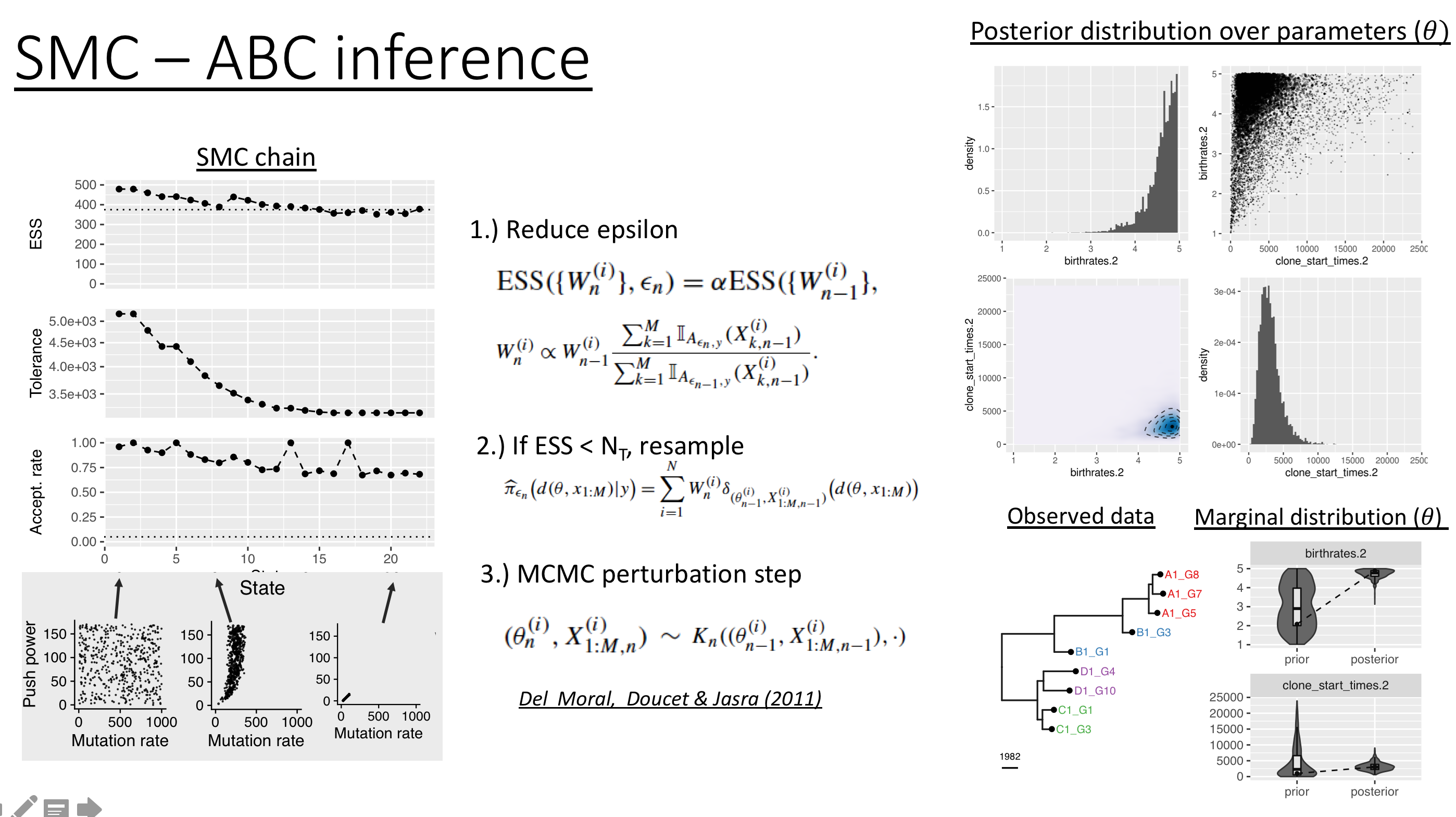
